# Cereblon-related mild intellectual disability disrupts response inhibition and uniformity of group–individual strategies

**DOI:** 10.1101/2025.09.19.676948

**Authors:** Chan Hee Kim, Kyogu Lee, Jeong-Eun Seo, Yoon Jeong Heo, Se-Young Choi

## Abstract

Temporal processing, including duration, is essential for survival and communication across species. Intellectual disability (ID), which has diverse causes, including Cereblon (CRBN), impairs duration discrimination. CRBN-related ID, link to abnormal cognitive behaviors, may disrupt both perception and behavior during duration discrimination. However, cross-species behavioral strategies, their variation with ID, and associated behavioral indices remain unclear. Here, humans and wild-type (WT) mice with typical intelligence, and CRBN knockout (KO) mice with ID, performed an auditory duration discrimination task with Long (10 s) and Short (2 s) cues. All groups distinguished stimulus durations, but latency-based strategies diverged. KO mice showed impulsivity and divergent responses at both individual and group levels, whereas WT mice and humans consistently delayed their responses by ∼2 s to the Short cue length, reflecting inhibition and convergent responses. In typical intelligence models, latencies for both stimuli clustered between 2–5 s, while in ID model depended on stimulus duration. Duration perception is a conserved cross-species capacity, while task-specific cognitive strategies are intrinsically preserved though variation emerges with ID. We suggest that latency indices dissociated inhibition from impulsivity, and convergence from divergence. We further suggest that CRBN-related ID preserves perceptual understanding but disrupts the shared behavioral language of typical intelligence.

## Introduction

Duration is a fundamental element for interpreting external stimuli and enabling communication. Auditory signals are organized into discrete units—such as phonemes or tones—whose durations convey meaning in contexts ranging from animal vocalizations to human speech and music. Humans can discriminate duration [1], and animals exhibit similar abilities [2-5]. Although they do not share the same language, these abilities can elicit comparable behavioral responses in tasks distinguishing two tones of different lengths [6]. Sensory discrimination is closely linked to intelligence [7]; even mild intellectual disability (ID; formerly mental retardation) can impair fundamental cognitive abilities [8, 9], including duration discrimination, leading to delayed or poor responses compared with individuals of typical intelligence [10-12]. ID is defined by significant deficits in intellectual functioning and adaptive behavior across conceptual, social, and practical domains [13]. This study investigated whether humans and animals exhibit comparable behavioral strategies in auditory duration discrimination task and whether these strategies are influenced by intelligence.

Task-related behaviors often rely on cognitive strategies [14] that optimize learning, problem-solving, and performance, and variability in behavior may reflect differences in such strategies [15, 16]. Reaction time (RT), defined as response latency, and correct rate (CR), reflecting response accuracy, serve as essential behavioral indices in both humans and animals. RT, in particular, is informative for evaluating cognitive strategies [9, 17, 18] and may correlated with CR, while being influenced by both subject- and stimulus-related factors. In mice, RT may differ methodologically from human RT due to the differences in the apparatus, protocol, and the behavioral nature of repeated responses. Building on our previous work [19], which defined and categorized RT-related indices from mouse responses to conditioned stimuli (CSs) of varying durations, we selected 10-s (Long) and 2-s (Short) white noises for the present duration discrimination task. This contrast was chosen to maximize perceptual contrast and to elicit distinct RT patterns. We examined how RT-related indices reflected subject-specific characteristics and their relationship with CRs (Figure S1). RTs to Long and Short stimuli may reveal either shared or distinct cognitive strategies across species and intelligence levels.

Transgenic animal models with targeted gene suppression have been instrumental in overcoming the limitations of patient-based research and elucidating neural mechanisms underlying ID [20]. Here, we generated an ID model via mutations in the Cereblon (CRBN) gene, which cause mild ID [21, 22], representing the majority of ID cases [23]. CRBN is critical for cognitive processes and memory [24-28], and its dysfunction is associated with atypical cognitive functions [25-27] and autism spectrum disorder (ASD) [29], in which time perception deficits are common [30, 31]. These observations suggest that CRBN-related deficits may affect both perceptual and behavioral performance in duration discrimination tasks. However, whether duration discrimination is preserved in individuals with CRBN-related ID and how it differs from those with typical intelligence, remains poorly understood. Because ID is heterogeneous [32] and CRBN mutations cause autosomal recessive nonsyndromic ID [26], isolating CRBN-specific phenotypes in humans is challenging. CRBN knockout (KO) mice therefore provide a critical model for investigating gene-specific behavioral characteristics, and combined human-animal data can inform diagnostic and educational approaches.

The Go/No-Go paradigm is widely used to probe both perception (e.g., distinguishing Long and Short durations) and behavior (e.g., inhibition and impulsivity) [33-35]. In mice, this paradigm was adapted so that a reward (unconditioned stimulus, US) was delivered only in the Long-Go condition, whereas human performed a duration discrimination task to account for the mice’s tendency to respond even in No-Go trials (see Methods and Figure 1). Despite these methodological differences, both paradigms targeted duration evaluation and response inhibition. Three groups were tested: wild-type (WT) mice, CRBN KO mice, and healthy human participants. We investigated how cognitive strategies are shared between humans and mice, and how CRBN-related ID alters perceptual and behavioral responses to auditory cues of different durations. We further examined which behavioral indices in humans and mice differentiate between perception and behavior, and between inhibition and impulsivity. We hypothesized that humans would resemble with WT mice, whereas KO mice would show perceptual deficits leading to atypical behavioral responses. Because CRBN may affect both perception and behavior, its dysfunction was expected to impair duration discrimination, sound–reward associations, and the regulation of response inhibition and impulsivity. Since the ability to discriminate auditory duration is fundamental to survival [36] and communication [37], and is closely linked to intelligence and language [38], our findings can provide insights into how alters cognitive strategies—particularly those involving impulsivity and inhibition in decision-making [39]—contribute to communication difficulties in CRBN-related ID, especially in social contexts [40] (Figure S1).

**Figure 1.**
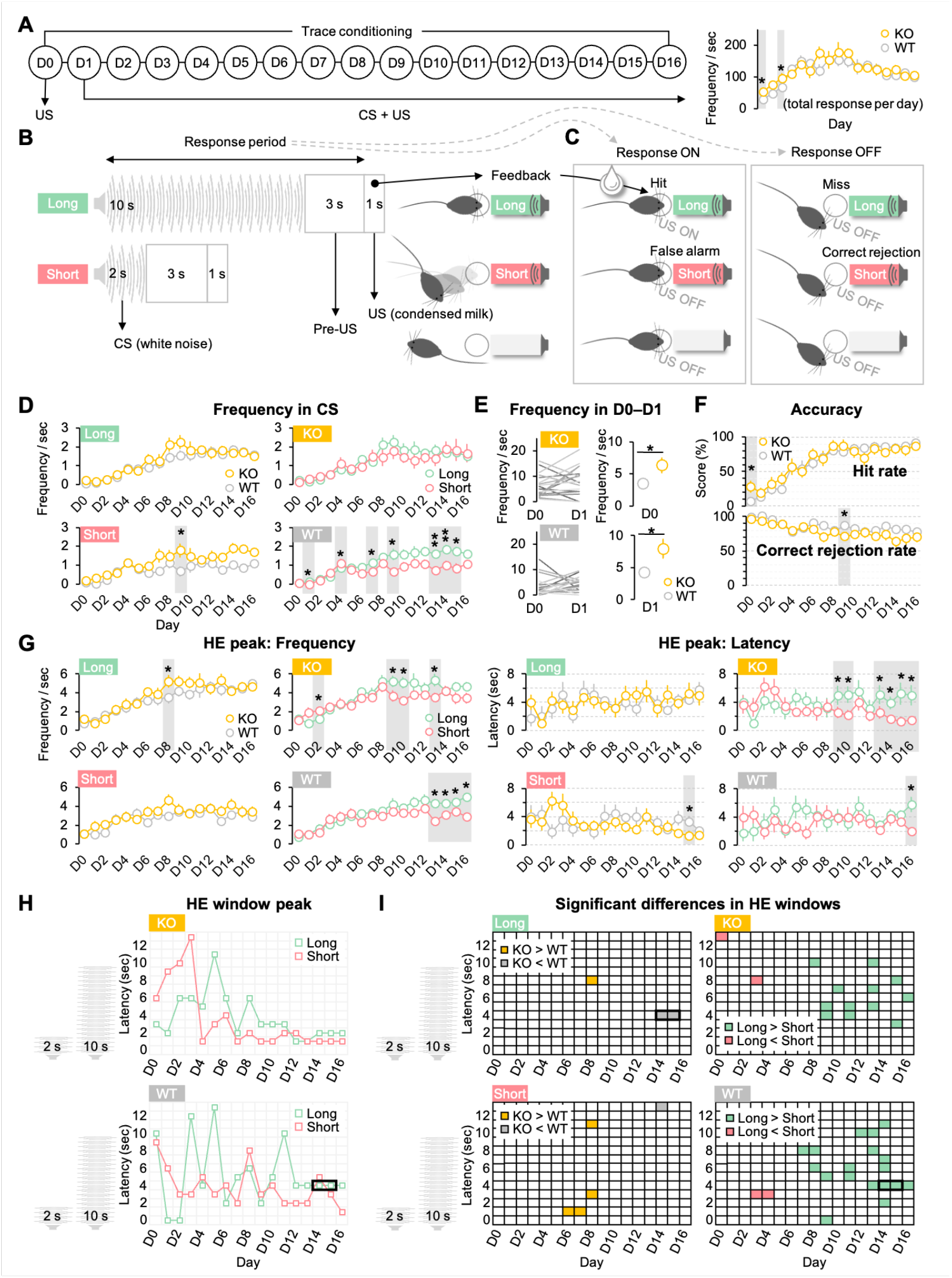
Mouse experiment and behavioral responses. (A) Trace conditioning over 16 days showed no difference in total HEs per day between KO and WT mice. (B) Mice learned to discriminate Long (Go) and Short (No-Go) trials. (C) In Long trials, US delivery followed an HE between CS onset and pre-US offset. A hit was an HE within 10 s of CS onset. (D) Significant group and condition differences in HE frequency appeared only during the CS period in late training days. (E) On D0–D1, KO mice had higher HE frequency than WT. (F) Accuracy (hits and correct rejections) during CS did not prominently differ across groups or conditions. (G) HE peak frequency varied by condition. HE peak latency was stable in WT mice, but it was variable in KO mice. (H) In late training, HE window peak latency was 1–3 s across conditions in KO mice; WT peaked after 2 s. (I) On D14–D15, the 4–5 s window showed group differences in Long trials and Long–Short differences in WT (Figure S7), overlapping with the WT HE window peak (panel H, Figure S6), suggesting a critical decision period for Long trials. See Figures S2–S7 and Tables S1–S5 for statistics. Statistical tests: Wilcoxon signed-rank, Mann–Whitney U; *p < 0.05, **p < 0.01 (N = 9). Error bars: 95% CI. Abbreviations: D, day; CS, conditioned stimulus; US, unconditioned stimulus; WT, wild-type; KO, CRBN KO; HE, head entry.

## Results

Behavioral data recorded response timing during task performance. In mice, head entries (HEs) into the food cup were variable, whereas humans showed a one-to-one response (see Methods). We analyzed classical indices—accuracy and latency—plus repetitive behavior in mice. To accommodate protocol differences, humans were measured using correct rate (CR) and reaction time (RT), and mice using HE accuracy and latency. Data were collected for Long (10 s) and Short (2 s) conditions in both mice and humans, using task designs adapted to their behavioral and cognitive profiles.

### Mouse behavioral data

Behavioral analyses for WT and KO mice included: (1) HE frequency, (2) HE accuracy (hit and correct rejection rates), (3) HE and fastest HE (FHE) peak frequency and latency, and (4) HE window peak frequency and latency derived from latency-window segmentation. These indices were used to assess behavior, discrimination, and temporal decision strategies. Indices (1)–(3) focused on individual CS periods, whereas (4) was based on parallel segments aligned to CS onset, independent of CS duration. Analyses of (4) were specifically conducted to identify the time point of decision-making within 0–14 s after CS onset, encompassing the CS, pre-US, and US periods.

### HE frequency: Differences occurred only during CS period

HE frequency was calculated for CS, pre-US, US, pre-CS, post-CS, and entire period per training day. Group differences were not prominent even during the CS period (Figure 1D). WT mice only showed condition-specific differences (Long vs. Short) on D1, D5, D7, D9 and D13–D15, indicating cue discrimination. Significant differences were rarely found in non-CS periods (Figure S2A), and total HEs per day did not differ between groups in late training days (Figure 1A), showing KO mice did not generally over-respond [26]. HE frequency thus did not reflect habitual behavior in either group. See Table S1 for complete statistics.

### HE frequency on initial days (D0 and D1): KO mice adapted faster

Group differences in HE frequencies of merged conditions were observed on D0 (US introduced) and D1 (CS and US introduced). KO mice showed greater engagement and less hesitation in novel environments (Mann–Whitney U test, D0, Z = –2.471, p = 0.013; D1, Z = –2.078, p = 0.037; Figure 1E). Within-group comparisons between D0 and D1 were not significant (Wilcoxon signed-ranks test, p > 0.05; Table S1), suggesting KO mice adapted faster, while WT mice were more cautious.

### HE Accuracy: Both groups learned the task

HE accuracy was based on HEs during the 10-s (Long) or 2-s (Short) CS. Both KO and WT mice showed high hit and correct rejection rates, with few significant differences (Figure 1F; Figure S2B). Early in training (D0–D3, D5/D6 in both groups), correct rejection rates exceeded hit rates (Wilcoxon signed-ranks test, p < 0.05), but these differences largely disappeared over time, except for a D16 hit rate advantage in WT mice (Figure S2C; Table S2).

### HE peak: WT and KO mice discriminated with different decision time

HE peak frequency and latency, extracted from individual HE distributions [19], indicate preferred decision times. Both groups showed more HEs for Long than Short conditions, reflecting successful discrimination (Figure 1G). HE peak frequency was higher for Long on D13–D16 (WT) and D2, D9, D10, D13 (KO), while latency was longer for Long on D16 (WT) and D9, D10, D13–D16 (KO) (Wilcoxon signed-ranks test, p < 0.05 in all cases). KO mice responded faster to the Short cue (1–2 s) than to Long cue (4–6 s), whereas WT mice postponed responses to both cues, typically around 3–5 s, suggesting KO mice initiated actions immediately at cue onset and then decided whether to continue or stop based on cue duration, while WT mice waited before deciding. Group differences were limited to D8 (Long, frequency: WT < KO) and D15 (Short, latency: WT > KO, Mann–Whitney U test, p < 0.05). See Table S3 for complete statistics.

### FHE peak: Response patterns were similar between groups

Latency and frequency of the first responses at CS onset did not differ between groups. FHE peak showed limited training- or condition-related changes (Figure S2D; see our previous study [19] for extraction details). Patterns were consistent, except for D0 (Short, frequency), and D4/D9 (Short, latency; Wilcoxon signed-ranks test, p < 0.05). Long and Short responses were largely similar, except WT latency on D16. KO mice did not respond immediately to cues. See Table S3 for statistical details.

### HE and FHE peak analyses: KO mice showed delayed responses to the Short cue

HE and FHE peak frequency patterns were similar across conditions in both groups (Figure S3A). Latency differences were generally absent, except for the Long condition in KO on D13, unlike previous findings where only the Short condition was non-significant [19]. Correlations between HE and FHE peak frequencies for Short during late training days (Figure S3B) were significant in KO mice but weaker in WT. Similarly, correlation between HE and FHE peak latencies for Short during late training were also significant in KO mice, indicating premature responses (Figure S3C). In HE peak latency, the proportion of KO mice responding within 2 s to Short increased across training, whereas WT mice showed no trend (Figure S3D), suggesting KO mice persisted in responding to the unrewarded Short cue. See Table S3 for full statistics.

### Latency window segmentation: WT mice showed delayed responses compared to KO mice

We analyzed group behavior by tracing individual HE windows over 14 s after CS onset. HEs were segmented into fixed latency windows (0–2 s, 0–10 s, and 1-s bins up to 14 s), regardless of condition (Figure S4). In the 2-s and 10-s latency window analyses (0–2 s and 0–10 s from CS onset), group differences were not prominent. Condition differences were significant in the 0–10 s window across training days (D1, D4–D6, D8, D9, and D11–D16 for WT mice; D2 and D4–D16 for KO mice; Wilcoxon signed-ranks test, p < 0.05 in all cases). In the 0–2 s window, differences occurred only on D13 (KO) and D8 (WT) (p < 0.05; Figure S4; Table S4). These results suggest cue effects—heard consistently during the 0–2 s window in both conditions and alternately during the 2–10 s window—were similar across groups.

For 1-s bin analyses, we extracted the peak bin from HE traces over 0–14 s for each training day (HE window peak; Figure S5 and S6), and assessed overlap with bins showing significant group or condition differences (Figure S7). WT mice showed HE window peaks at 4–5 s during Long cue trials, particularly on D14–D15, with significant differences overlapping for both between and within groups (Figure 1H and 1I). KO mice responded earlier, with peaks at 1–2 s for Short cues and 2–3 s for Long cues. The prominent HE window peaks for Long cue were due to lower responses compared with Short cues in WT mice and all conditions in KO mice (Figure S6). Across D12–D16, HE window peaks for Long cues in WT mice remained consistency in the 4–5 s bin, whereas Short cue peaks fluctuated. During D12–D16, latencies for both conditions were longer in WT mice than KO mice, except for Short cues on D12. These temporal distinctions indicate KO mice initiated responses faster and more persistently, whereas WT mice delayed responses, consistent with more deliberate decision-making within the 4–5 s window and fewer responses to Short cue. See Figure S6 for the frequency and latency of daily peaks, Figure S7 and Table S5 for the statistics for HE frequencies across 1-s bins (0–14 s).

### Human behavioral data

#### CR and RT: Humans paused responses until 2 s

In the duration discrimination task (Figure 2A), all participants rated it as “easy” or “very easy” on a 5-point Likert scale (very easy, easy, neutral, difficult, very difficult). Mean CR was high: 99.542 ± 0.924% for Long and 97.333 ± 3.393% for Short, with a small but significant difference favoring Long (Wilcoxon signed-rank test, Z = −4.107, p < 0.0001; Figure 2B). Mean RTs was significantly longer for Long (3.316 ± 0.739 s) than for Short (2.576 ± 0.111 s), with all responses exceeding 2 s (Wilcoxon signed-rank test, Z = −5.511, p < 0.0000001; Figure 2B). Only two subjects exhibited RTs >5 s, and none exhibited RTs within 2 s. Self-reports indicated participants typically clicked immediately after the offset of Short cue and mentally estimated the Short duration during Long trials. Consistently, RTs for Long and Short were positively correlated (Spearman’s ρ = 0.601, p < 0.0001; Table S6).

**Figure 2.**
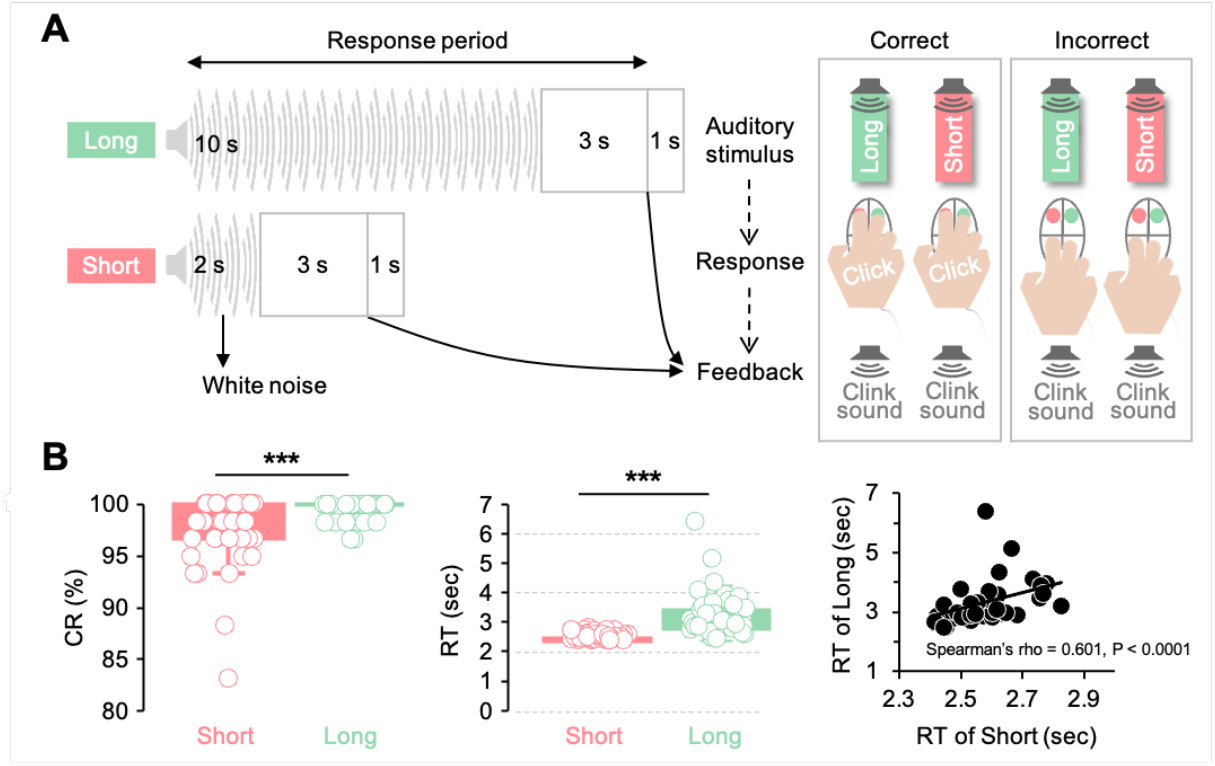
Human experiment and behavioral responses. (A) Participants performed a duration discrimination task with Long and Short trials. (B) Correct rate (CR) was higher in Long (99.54 ± 0.92%) than Short (97.33 ± 3.39%; Wilcoxon, Z = −4.107, p < 0.0001). Reaction time (RT) was longer in Long than Short (Z = −5.511, p < 0.0000001). Long and Short RTs were positively correlated (Spearman’s ρ = 0.601, p < 0.0001). (C) No subject had a mean RT < 2 s. See Table S6 for full statistics. Statistical tests: Wilcoxon signed-rank, Spearman’s rank correlation coefficient; ***p < 0.001 (N = 40). In each box plot, the box spans the interquartile range, the line marks the median, and whiskers indicate the most extreme non-outlier values. Abbreviations: CR, correct rate; RT, reaction time.

### Mouse–human comparison

#### RT and latencies: Humans and WT mice showed similar pattern

HE peak latency (mean of individual HE peak latencies) and HE window peak latency (HE peak latency of grand average across subjects) were compared over 0–14 s window (Figure S5). WT mice showed convergence within 2–5 s latencies for Long and Short, paralleling human RTs over 2 s (Figure 3A). KO mice responded earlier (<2 s) and less uninformed HE window peak latencies compared to delayed HE peak latency for Long (Figure 3A and 3B), suggesting individual variability and group-level HE window peaks affected by impulsive responses.

**Figure 3.**
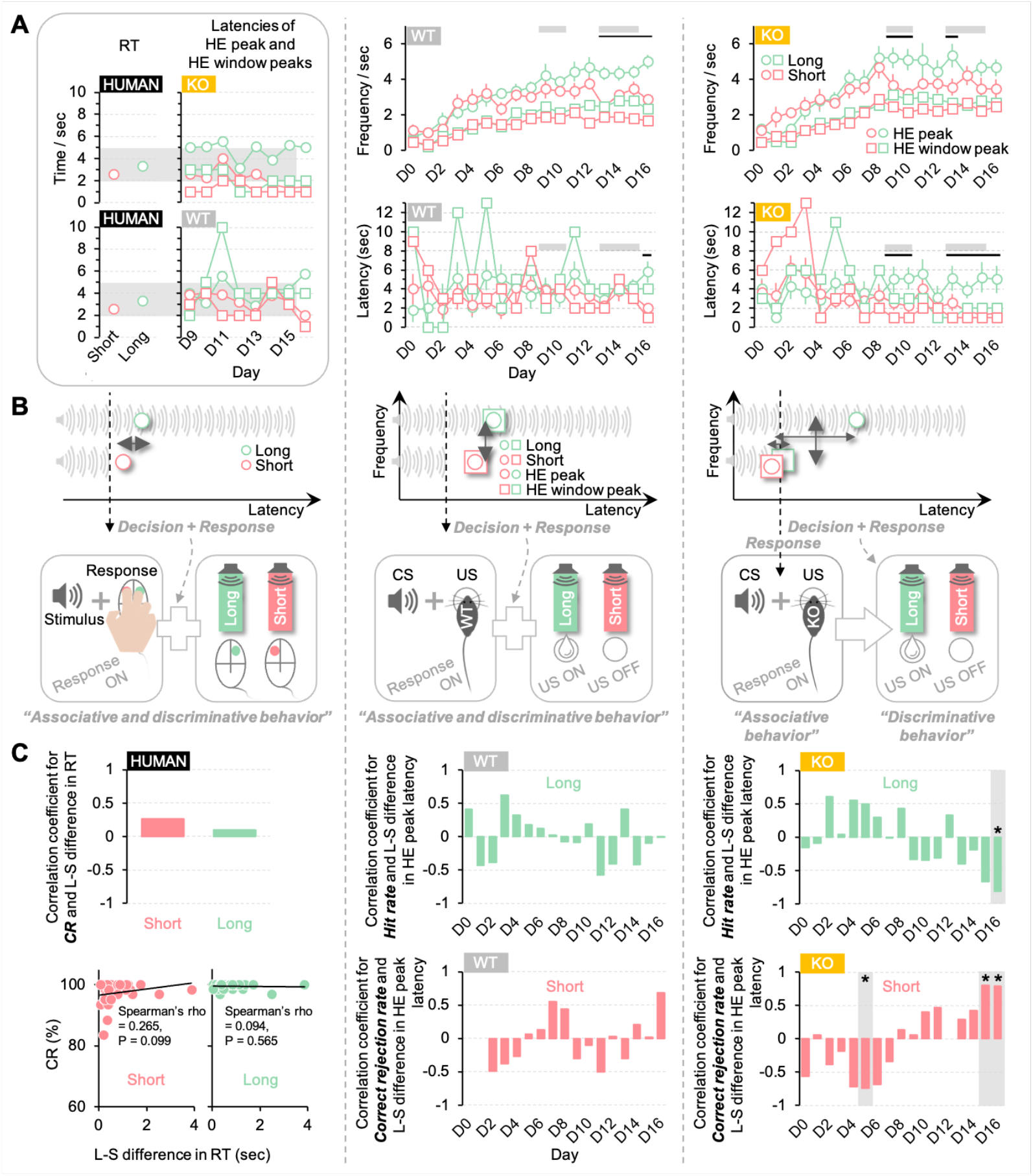
RT/latency and CR/accuracy in humans and mice. (A) Human RTs overlapped with WT but not KO peak latencies (gray areas, left). Gray bars (middle, right) mark comparable response periods; black bars mark training periods with significant HE peak differences. (B) WT mice showed consistent HE and HE window peaks (>2 s), similar to humans. KO mice showed inconsistent peaks: Long in HE peak delayed >2 s only. In WT/humans, associative and discriminative processes coincided; in KO, they dissociated into two stages. (C) L–S RT differences (HE latency in mice) correlated with CR only in KO (Spearman, N = 9, p < 0.05): latency negatively with hit rate (D16), positively with correct rejection (D5, D15, D16). No significant correlations in humans (N = 40) or WT (N = 9). See Figure S8 and Table S7. Error bars: 95% CI. Abbreviations: RT, reaction time; HE, head entry; D, training day; WT, wild-type; KO, CRBN KO; CS, conditioned stimulus; US, unconditioned stimulus.

#### Correlation between latency and accuracy: Humans and WT mice showed similar pattern

Humans and WT mice showed no significant correlation between latency (RT or HE peak latency) and accuracy (CR) (Spearman’s rank correlation coefficient, p > 0.05 in all cases). KO mice exhibited a negative correlation between the Long–Short difference in HE peak latency and hit rate (Spearman’s ρ = −0.808, p = 0.017 on D16), and a positive correlation with correct rejection rate (ρ = −0.735, p = 0.048 on D5; ρ = 0.798, p = 0.020 on D15; ρ = 0.788, p = 0.023 on D16) (Figure 3C). Further analyses with raw RT/latency and CR/accuracy values supported these findings exclusively in KO mice (Figure S8). These findings highlight that shorter latencies in KO mice improved No-Go accuracy but reduced Go accuracy, reflecting an impulsive bias. In contrast, humans and WT mice showed consistent delayed responses uncorrelated with task performance, reflecting more stable cognitive strategies. See Table S7 for full statistics.

## Discussion

### Cognitive strategy and RT/latency

This study investigated how cognitive strategies are shared across species with typical intelligence and how these strategies are affected by the ID phenotype caused by CRBN deficiency. All subjects, regardless of intelligence, discriminated between two auditory conditions, as reflected in accuracy (CR). However, behavioral responses differed between subjects with ID and those with typical intelligence. Behavioral data were interpreted within two frameworks: (1) response inhibition versus impulsive response, and (2) convergence versus divergence between group- and individual-level responses (Figures 3 and 4). CRBN KO mice exhibited impulsive responses and divergence, whereas WT mice and humans without ID showed response inhibition and convergence. RT and latencies of the HE peak and HE window peak differentiated behavioral patterns (Figure S1), indicating that cognitive strategies in auditory temporal discrimination are clearly reflected in RT and latency measures.

**Figure 4.**
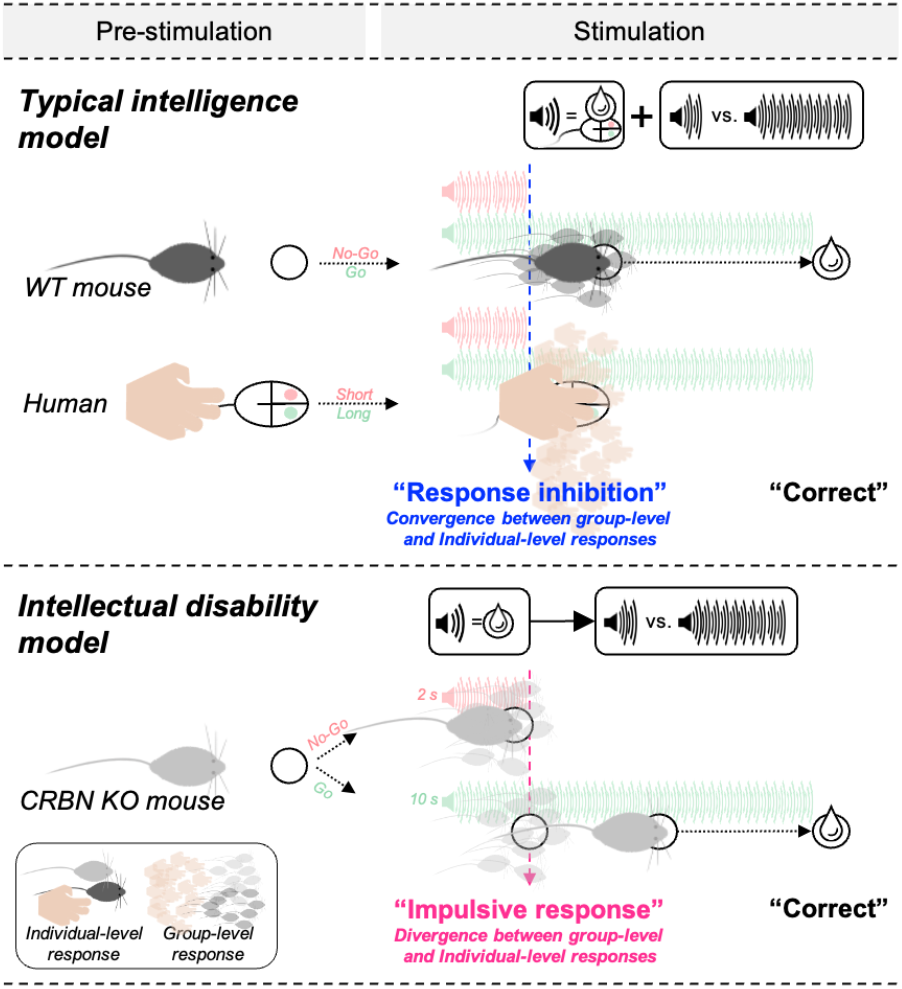
Intelligence-dependent behavioral responses and intelligence-independent perceptual responses. All groups were able to successfully discriminate the stimuli, regardless of intelligence level. However, WT mice and humans with typical intelligence exhibited response inhibition during decision-making until 2 s (the offset of the Short condition). In contrast, CRBN KO mice with ID showed impulsive responses around the 2-s mark, indicating a dissociation between associative and discriminative processes in KO mice. Furthermore, impulsivity in KO mice led to a divergence between the group-level response (HE window peak) and the individual-level response (HE peak). In contrast, humans and WT mice exhibited a convergence between group- and individual-level responses.

### Perception and behavior

Accuracy (CR) and HE peak frequency reflected recognition of the two conditions across all groups (Figures 1F and 1G). In mice, higher HE frequency generally indicates good task understanding. HE frequency were not consistent in WT and KO mice unlike HE peak frequency. Also, HE frequency during the CS period and HE window peak frequency did not clearly differentiate the two conditions in KO mice (Figure 1D and 3A). These results in KO mice were not due to over-responsiveness (Figure 1A). Behavioral indices were categorized as accuracy (CR), latency (RT), and frequency. In WT mice and humans, frequency and accuracy reflect stimulus perception, while latency reflects cognitive strategies. In KO mice, perception and behavior were mixed within the indices: perception appeared only in HE peak frequency, whereas HE frequency and HE window peak frequency, unrelated to perception, likely reflect artifacts (Figure S1C).

### Response inhibition and implicit response

Impulsive behaviors likely caused the artifacts in KO mice [41, 42]. The Go/No-Go paradigm assesses both perception and behavior, including impulsivity and inhibition [43, 44]. Mice learned both CS–US associations and Long–Short discrimination during training, but KO mice might initiate premature responses, followed by a decision to continue or inhibit responding. These behaviors disrupted the alignment between HE peak frequency and HE window peak frequency (Figures 3A), enhanced HE peak–FHE peak alignment, and increased responses within 2 s during the No-Go condition (Figures S3B and S3D). In a previous study using the same apparatus [19], freely moving C57BL/6NCrljOri mice reached the food cup within 2 s. In contrast, WT mice delayed responses until after 2 s, reflecting response inhibition [45, 46], and typically responded between 2–5 s post stimulus, allowing simultaneous processing of association and discrimination. Impulsivity has been documented in animals and humans [47]. Human studies have assessed impulsivity using self-report questionnaires [48] and meta-analyses [49, 50]. In our study, we employed a duration discrimination task instead of Go/No-Go paradigm to measure human responses. Like WT mice, humans paused ∼2 s after stimulus onset, implying duration comparison between Long and Short cues, before responding between 2 and 5 s (Figure 4). This 2-s point marks a critical threshold for demonstrating response inhibition.

### Group-level and individual-level responses

The “HE peak” reflects the average timing of individual responses, while the “HE window peak” reflects the group-level peak from averaged data (Figure S5). When individual and group peaks across 1-s bins (0–14 s) coincide, they align, indicating convergence between group- and individual-level responses. WT mice showed such alignment, suggesting shared decision timing, whereas, KO mice exhibited divergence, reflecting difference between individual and group strategies (Figure 4). The overlapping HE peak and HE window peak between 2–5 s in WT mice resembled human RTs, with both species delaying responses 2 s post-stimulus, indicative of response inhibition. Group averages often mask individual variability, as seen in insect learnings [51-54], and human tasks [55]. In our study, differences between group and individual patterns arose from analytical approaches (Figure S5), the HE window peak is derived from averaged data of all subjects, while the HE peak is the average of individual peaks. HE peak frequencies vary by latency across individuals, whereas HE window peak frequency is time-locked. Thus, HE peak and HE window peak serve as indicators of convergence or divergence between group- and individual-level behavioral responses.

### Human and mouse

Duration discrimination of auditory signals is an intrinsic ability in both humans and animals. However, species-specific differences complicate experimental design and interpretation. Even when healthy humans and mice show similar behaviors, methodological differences are unavoidable due to distinct repertoires and extended training in animals (Figure S1A). Humans entered the task with strategies shaped by prior experience, while mice were tested in a fixed cage environment. Humans completed three sessions in a single day, whereas mice underwent a 16-day training protocol, including pre-conditioning (Figure S1A). Despite different response modalities—head entry for mice versus mouse click for humans—both species converged on a common strategy within the 2–5 s time window (Figures 3 and 4). Human reported using the 2-s “Short” condition as a reference, and although mice did not report strategies, RTs and latencies in both humans and WT mice reflected comparable cognitive strategies. The ∼2 s pause indicates response inhibition, suggesting an intrinsic cognitive strategy [14] that generalizes across species with typical intelligence while allowing individual variation. These findings indicate that although strategies are learned, the ability to develop them is a conserved, cross-species trait. Behavioral indices reveal this shared strategy—not via identical paradigms, but through species-adapted variations.

### ID and CRBN

Previous studies linked CRBN to deficits in working memory [11], cognitive performance [56], stress [57], and depression [58], and to bipolar disorder, associated with impulsive and poor decision-making [39]. Our study confirmed impulsive behaviors prior to decision-making in CRBN KO mice, but perceptual discrimination, reflected in accuracy and HE peak frequency, was preserved, contrasting with delayed or poor responses in ID populations [10-12]. KO mice did not show abnormal repetitive behavior, consistent with prior findings [26] and unlike observations in ASD patients [59].

When first exposed to the US (milk) and CS (white noise, 75 dB), KO mice were more active than WT mice (Figure 1E). Their learning curve (HE peak frequency) indicated faster task acquisition (Figure 1G), possibly reflecting fearlessness linked to memory and learning deficits [25]. Only KO mice showed significant correlation between RT/latency and accuracy (Figure 3C). Delayed responses reduced hit rates, while faster responses decreased correct rejections, consistent with impulsive behavior. Their behaviors may also reflect reduced neophobia and heightened attention to reward cues, supporting robust CS–US associations. Despite impulsivity, perceptual accuracy was maintained, indicating the ID phenotype reflects alternative cognitive strategies rather than reduced intelligence. KO mice might simply respond in their own way.

However, these behavioral strategies were not uninform within the KO group, as seen in divergence between individual HE peak and group HE window peak. KO mice responded impatiently and excessively from the perspective of WT mice. The shared timing of response inhibition in humans and WT mice reflects a “common language” of cognitive strategy, which KO mice lacked. From the WT perspective, KO mice’s impulsivity may appear as unexpected noise. In human social communication, mutual predictability is essential, atypical responses of KO mice—and by extension, individuals with ID—may fall outside this implicit predictability. These findings suggest that understanding ID requires focusing not only on deficits or form of inferiority but on differences in cognitive strategy and behavior. Effective interventions [60] should start with a deeper understanding of how individuals approach tasks.

### Limitation

The Go/No-Go paradigm used—distinguishing 2- and 10-s cues—is relatively simple, stimulus-fixed task compared to prior studies. Food reward likely motivated the mice, and auditory sensitivity, essential for survival, appears intact in KO mice with mild ID, enabling accurate perception. Our findings indicate that duration processing strategies—a key aspect of auditory perception, including speech and music—are linked to intelligence. The paradigm was applied only to mice; humans were tested with the same stimuli but a different task. Despite these differences—WT mice running toward a food cup versus humans clicking— response latencies were similar (2–5 s). Future research should test both species using identical duration discrimination paradigm to confirm cross-species comparability. This study relied on female subjects, who are typically more sensitive to auditory cues; indicating males are necessary to examine sex differences. Findings in CRBN KO mice should also be replicated in other ID models [61-65] with varying severity [23].

## Conclusion

Time processing is essential for survival [36] and communication [37] and is impaired in disorders such as autism and schizophrenia [66], which involve cognitive and social-communication difficulties. ID affects time perception [67], duration judgment [12], and behaviors like RT [60]. Our findings suggest that CRBN mutations causing mild ID [21, 22] affect behavior and cognitive strategy rather than perception, potentially contributing to communication difficulties with neurotypical individuals. Behavioral indices were categorized into perception and behavior; in CRBN KO mice, these were intermixed, unlike WT mice where they were distinct. CRBN modulated response inhibition in associating sound with reward and discriminating durations. HE peak latency and HE window peak latency, and their harmonization, serve as indicators to distinguish response inhibition from impulsivity and to reflect unified cognitive strategies within a group. Latency and RT data also indicate a shared behavioral language of cognitive strategy between mice and humans with typical intelligence. Finally, duration perception is a fundamental capacity across species, and the ability to acquire cognitive strategies naturally during task performance is a conserved cross-species trait, though it varies with ID.

## Materials and Methods

### Mouse experiment

#### Subjects

The subjects were 18 mice divided into two groups: C57BL/6NCrljOri and CRBN KO mice (9 mice per group, aged 7 weeks). All subjects were female, based on evidence suggesting that females exhibit greater sensitivity to auditory signals [68-71], juxtaposed with human subjects. All animals were supplied by the Experimental Animal Center at Seoul National Dental University. Each mouse was housed individually and placed on a food restriction schedule to maintain its body weight at 85% of the normal free-feeding level throughout the experiment. Water was provided ad libitum. The housing environment was maintained on a 12-hour light–dark cycle, with lights on from 8:00 a.m. to 8:00 p.m. Behavioral training was conducted between 8 and 10 weeks of age. All procedures involving animals were reviewed and approved by the Institutional Animal Care and Use Committee of Seoul National University (SNU IACUC).

#### Apparatus

All experimental procedures were programmed and controlled using MED-PC V software (model SOF-735). Behavioral experiments were carried out in operant conditioning chambers (model EBV-307W-CT, Med Associates; interior dimensions: 21.6 × 17.8 × 12.7 cm). Each chamber featured a stainless steel grid floor composed of 24 rods (diameter: 0.32 cm; model ENV-307W-GF; floor dimensions: 17.8 × 15.2 × 5.7 cm) and was housed within a sound-attenuating MDF cubicle (model ENV-022MD). A house light (28 V DC, 1000 mA; model ENV-315W) remained illuminated throughout each session of a day. Auditory cues were delivered at 75 dB [72] via a white noise amplifier (10–25,000 Hz; model ENV-325SW) and emitted through a speaker mounted inside the chamber (model ENV-324W). Background noise from cubicle ventilation fans (model ENV-025F) was measured at approximately 65 dB. As a reward, condensed milk was dispensed into a 0.5-cc stainless steel cup via a liquid pipe (model ENV-303LPHD) connected to a syringe pump (model PHM-100A-EURO). Head entries (HEs) were automatically detected whenever the mouse interrupted the infrared beam within the liquid pipe (detector model ENV-303HDW).

#### Stimulus

In the task, two types of stimuli were used: a conditioned stimulus (CS) and an unconditioned stimulus (US). The CS was white noise presented at two durations—10 s and 2 s (Figure 1B). These durations were selected based on our previous study (16), which found no significant differences in response frequency or accuracy between the two CS conditions. In the trace conditioning paradigm, the US consisted of a 1-s delivery of condensed milk. The US occurred only 14–15 s after the onset of the 10-s CS and was not delivered following the 2-s CS. No stimulation was presented during the pre-US interval (i.e., the period between CS offset and US onset). The 10-s CS paired with the US in the current paradigm was designed to test delayed responses in decision-making, as the CS—strongly associated with the expectation of a food reward—was presented for a longer duration compared to 2-s CS. For this reason, we did not include a reversed condition in which the 2-s CS was paired with the US and the 10-s CS was not.

#### Procedure

Given the ID phenotype in KO mice, we implemented a simplified Go/No-Go paradigm in which a reward of US was provided only in the Go (Long) condition, and no punishment was used (Figure 1C). Mice were required to decide whether to make a HE response into a food cup presented during Go trials, which would result in a food reward. This protocol also allowed for repeated responses, enabling the assessment of behavioral differences between WT and KO mice over the course of operant conditioning, particularly since CRBN is associated with ASD-like repetitive behaviors [59].

Prior to the main trace conditioning phase, during 8-day preconditioning period, food restriction was applied to reduce each mouse’s weight to 85% of its baseline. Mice were also acclimated to handling and to the US. On the final day of preconditioning, a session was conducted in the experimental chamber in which the US was delivered without presentation of the CS. The main experiment was conducted over 16 consecutive days (Figure 1A). Each daily session consisted of each of 10 trials of 10 and 2 s, with a total session duration of approximately 1 hour per mouse, including preparation. The intertrial intervals were fixed as 90 s. During each trial, the US—a 1-s delivery of condensed milk—was administered at the offset of the Pre-US, triggered by the mice’s HE response except for D0–D2 of the adaptation period. Even though the US was contingent on a behavioral response, no punishments were applied for missed or premature responses. Mice were allowed to move freely within the chamber. The US was dispensed via syringe pumps connected to liquid pipes and food cups, with the pump activated only during the US interval. In this study, the behavioral response directed toward the food cup to consume the reward, defined as a “head entry” (HE), was recorded with a temporal resolution of 10 ms.

#### Analysis

Data were analyzed across two groups of 18 mice (9 CRBN KO and 9 WT mice), corresponding to the two CS durations: 10 and 2 s. First, HE frequencies were calculated for individual time windows. Total HE frequency were computed across each session. The HE frequencies before and after the CS onset, were computed for CS, pre-US, US, pre-CS, and post-CS periods, respectively. Pre-CS period was from -30 to 0 s after the onset of CS. Post-CS was 30 to 0 s after the offset of CS. The number of HEs for each subject was normalized to frequency per second. Second, HE accuracy was evaluated by the occurrence of HEs during CS period of 10 and 2 s, which was calculated on a per-trial basis and expressed as a percentage (with one correct trial equal to 10%). Hit and miss were estimated based on the HE for the CS duration of 10 s (Long), and correct rejection and false alarm were based on the CS duration of 10 s 2 s (Short). Third, the values of frequency and latency for the HE and first HE (FHE) peaks in individual subjects were extracted in accordance with our previous study [19]. The time window for both HE and FHE peaks was set from 0 to 10 s after CS onset across all groups. The HE and FHE peak revealed the most frequent response timing of individual subjects. Fourth, to trace changes in the HE density merged by all subjects, we estimated both frequency and latency of individual peaks per 1-s latency bin from 0 to 14 s encompassing CS, pre-US, and US periods. From these frequency and latency data, we extracted HE window peaks across 16 training days and the time windows from 0 to 14 s (see Figure S5 for the extraction of HE window peak and HE peak). Unlike the HE peak, the HE window peak showed a time locked change in HE frequency. Except for the 1-s window, the HE frequencies for two time windows from 0 to 2 s and from 0 to 10 s after the CS onset were also estimated to confirm the effect of decision time point discriminating between Long and Short. Using 1-s windows, we identified bins with overlapping statistically significant differences both between and within groups across all comparisons. Finally, the correlation tests were conducted for individual indices of latency and accuracy.

Behavioral data were non-normally distributed; therefore, all analyses used nonparametric tests. The analysis focused on group differences by genotype and condition differences within groups, assessed separately and independently, without considering genotype × condition interaction as in AVNOA. Within-genotype comparisons (Long vs. Short) were tested with the Wilcoxon signed-rank test, while between-genotype comparisons (KO vs. WT) were tested with the Mann–Whitney U test. Correlations were examined using Spearman’s rank correlation coefficient (α < 0.05). Multiple comparisons for two conditions in the correlation analyses and HE vs. HHE peaks analyses were corrected using the Bonferroni method. All analyses were conducted in MATLAB (version 9.12.0.2039608; MathWorks Inc., Natick, MA, USA) and SPSS (version 25.0; IBM, Armonk, NY, USA).

### Human experiment

#### Subjects

Forty healthy participants (mean age = 23.75 ± 3.09 years, range = 21–26 years) were all female, consistent with mouse subjects, considering sensitivity to auditory signals [68-71]. They recruited from Seoul National University. The age was based on criteria of mouse subjects [73]. All participants were right-handed and had normal hearing. On a 5-point Likert scale (very easy, easy, normal, hard, very hard), all participants rated the task as either “very easy” or “easy.” This study was approved by the Institutional Review Board of the Clinical Research Institute, Seoul National University School of Dentistry (Approval No. S-D20210011). All experimental procedures complied with ethical guidelines. Written informed consent was obtained from all participants prior to the experiment. Participants were compensated for their time after completing the task.

#### Procedure

The auditory stimuli consisted of two types of white noise, 10 s and 2 s in duration (referred to as Long and Short, respectively), identical to those used in the mouse experiment. The Go-No-Go paradigm from mouse experiment was not adapted. This approach was chosen to better align with the characteristics of the mouse experiment and to capture responses to the *Short* stimulus—even in contests where mice typically show No-Go behavior. To obtain RT data comparable to the mouse data, showing the HEs even in Short cue, participants were asked to discriminate between Long and Short durations. The experiment comprised three sessions. Each session included 20 trials per condition (Long and Short), presented with a 5 s inter-trial interval. In each trial, participants were instructed to respond by clicking the left mouse button for Long and the right mouse button for Short, using the index and/or middle fingers of their right hand (Figure 2A). They were required to make at least one click during the sound presentation and before hearing a subsequent “clink” sound. Before the main task, participants completed a training session consisting of approximately 10 practice trials per condition. No rewards (e.g., food) were provided, unlike in the mouse experiment. Instead, participants were encouraged to concentrate and minimize errors. Following the experiment, participants completed a questionnaire assessing task difficulty and the strategies employed. The total duration of the experiment, including preparation time, was approximately 1 hour. Auditory stimuli were generated using STIM2 software (Neuroscan, Charlotte, NC, United States) and presented at a sound pressure level of 60 dB through Sony stereo headphones (MDR-X50AP).

#### Analysis

Reaction time (RT) and correct rate (CR) for the Long and Short conditions were calculated for each participant. RT values were computed only from correct trials. For statistical analysis, the mean value across all trials for each subject was used. Because the dataset did not consistently follow a normal distribution, nonparametric tests were applied in line with the mouse data. These included the Wilcoxon signed-rank test, the Mann–Whitney U test, and Spearman’s rank correlation coefficient, with a significance threshold of α < 0.05. Bonferroni correction was applied to multiple comparisons in the correlation analyses. All analyses were conducted using SPSS (version 25.0; IBM, Armonk, NY, USA).

## Acknowledgments

This work was supported by Seoul National University Research Grant in 2021 and Basic Science Research Program through the National Research Foundation of Korea (NRF) funded by the Ministry of Education (RS-2022-NR075566).

## Author contributions

Conceptualization, C.H.K.; methodology, C.H.K. and S.Y.C.; investigation, C.H.K.; formal analysis, C.H.K.; visualization, C.H.K; writing – original draft, C.H.K.; writing – review & editing, C.H.K., K.L, J.E.S., Y.J.H., and S.Y.C.; resources, C.H.K. and S.Y.C.; supervision, C.H.K and S.Y.C.; funding acquisition, C.H.K and S.Y.C.

## Declaration of interests

The authors declare no competing interests.

## Notes

### Competing Interest Statement

The authors have declared no competing interest.

