## Supplementary figures for "Cereblon-related mild intellectual disability disrupts response inhibition and uniformity of group–individual strategies"

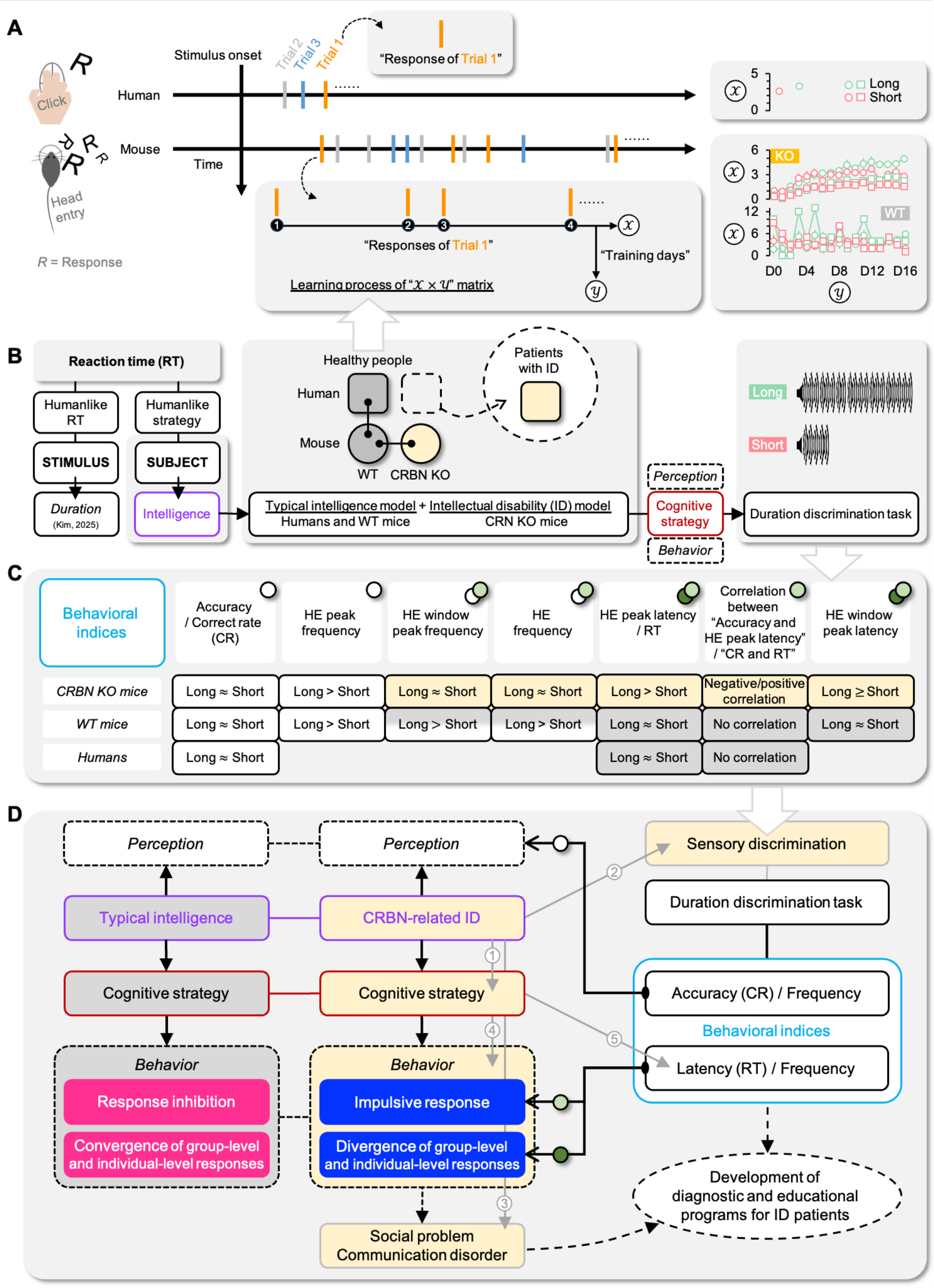


**Figure S1. Categorization of behavioral indices in the duration discrimination task.** (A) Human participants completed multiple trials within a single experimental day (x-axis). In contrast, mice underwent operant chamber conditioning over 16 days (x–y axis). Humans typically make a single response per trial, whereas mice exhibit repeated responses. Humans may have relied on pre-existing knowledge to develop cognitive strategies, whereas mice likely developed their strategies gradually through daily learning. (B) Based on previous study on human-like reaction times (RT) of mice in relation to stimulus duration [1], we designed a duration discrimination task that incorporates subject-dependent characteristics. We focused on human-like cognitive strategies in mice, as reflected in RT in relation to intelligence. Through these models, we expected to predict behavioral characteristics of patients with ID. (C) In this study, cognitive strategies and behavioral indices showed distinct characteristics depending on group. CRBN KO mice, a model of intellectual disability (ID), differed from WT mice and humans, who both represent typical intelligence. Behavioral indices include accuracy, frequency, and latency in mice, and CR and RT in humans. Same-colored boxes depict significant results with similar response patterns. White boxes indicate behavioral characteristics related to perception, common across all subjects regardless of ID. Yellow indicates characteristics specific to KO mice with ID, and gray indicates those specific to WT mice and humans without ID. White, green, and dark green circles correspond to the individual behavioral indices shown in panel D. During the duration discrimination task, differences between groups and conditions were observed. (D) Intelligence differences between subjects affects cognitive strategy (①) [2] and sensory discrimination (②) [3], and is connected to social problem and communication disorder (③) [4]. Cognitive strategy affects behavior (④) [5, 6] such as RT (⑤) [7-9], but not perception. Our data support classifying behavioral indices into two domains: “perception” and “behavior.” The former included accuracy (CR) and frequency, whereas the latter included latency (RT) and frequency. In our data, frequency measures—particularly HE window peak latency and HE frequency in WT mice—explained perceptual processes (similar to KO mice) and behavioral processes (different from KO mice). While ID affected behavioral responses, perceptual ability remained intact during task performance. Subject-dependent characteristics—such as intelligence and cognitive strategy—led to consistent patterns of response across various behavioral indices. Behavioral indices differentiated two properties: “response inhibition versus impulsive response” and “convergence versus divergence of group- and individual-level responses.” Finally, our finding suggest that CRBN-related ID phenotypes affects social problem and communication disorder.

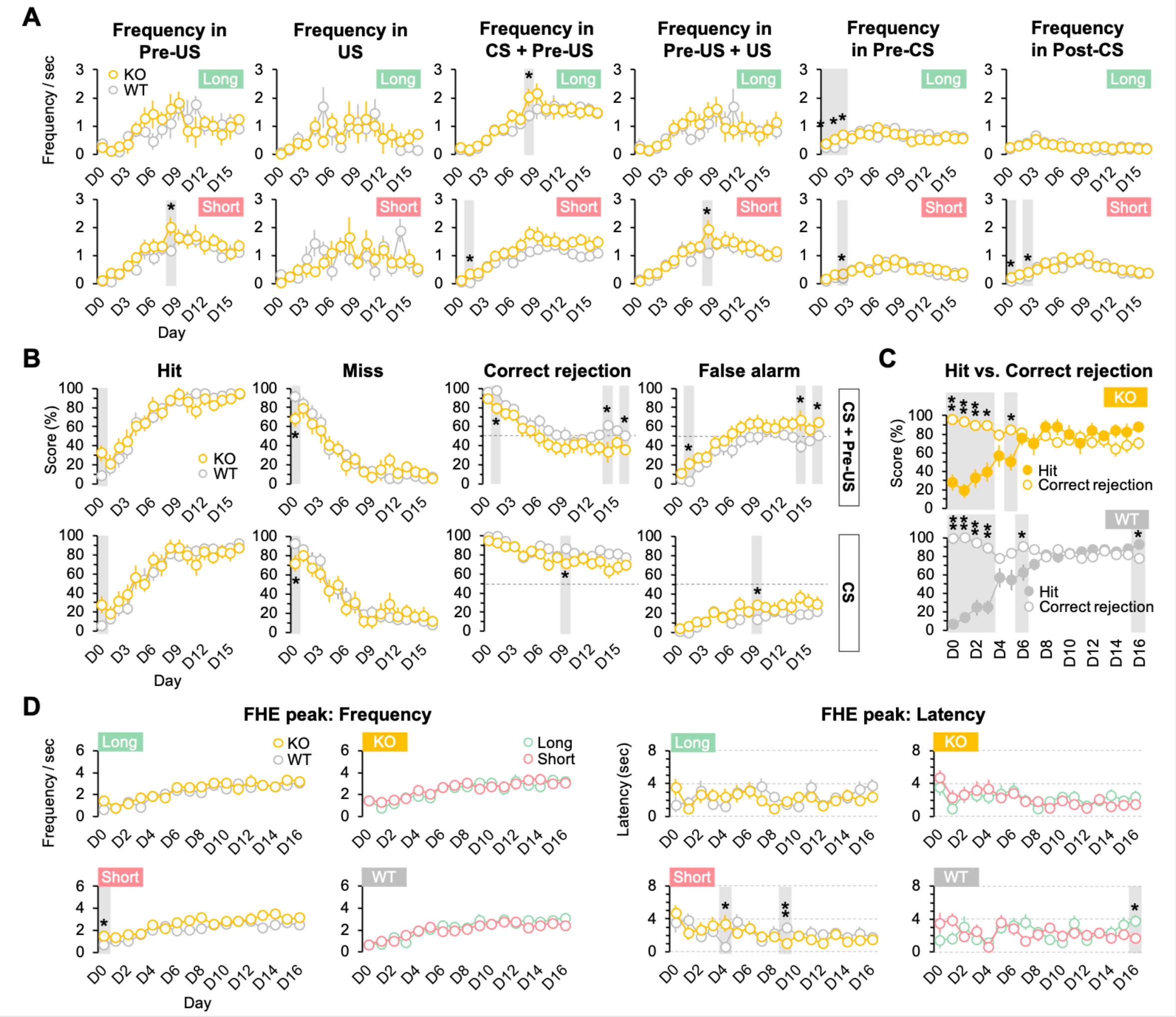


Figure S2. HE frequency and accuracy. (A) HE frequencies depending on the time windows. (B) Accuracy rate in the time windows of "CS + Pre-US" and "CS". (C) Difference between hit and correction rates. (D) Fastest HE (FHE) peak frequency and latency were comparable across groups and conditions. See Table S1–S3 for statistical details. Wilcoxon signed-ranks test and Mann–Whitney U test, *, $p < 0.05$; **, $p < 0.01$. Abbreviations: D, day; CS, conditioned stimulus; US, unconditioned stimulus; WT, wild-type mice; KO, CRBN KO mice HE, head entry; FHE, fastest HE.

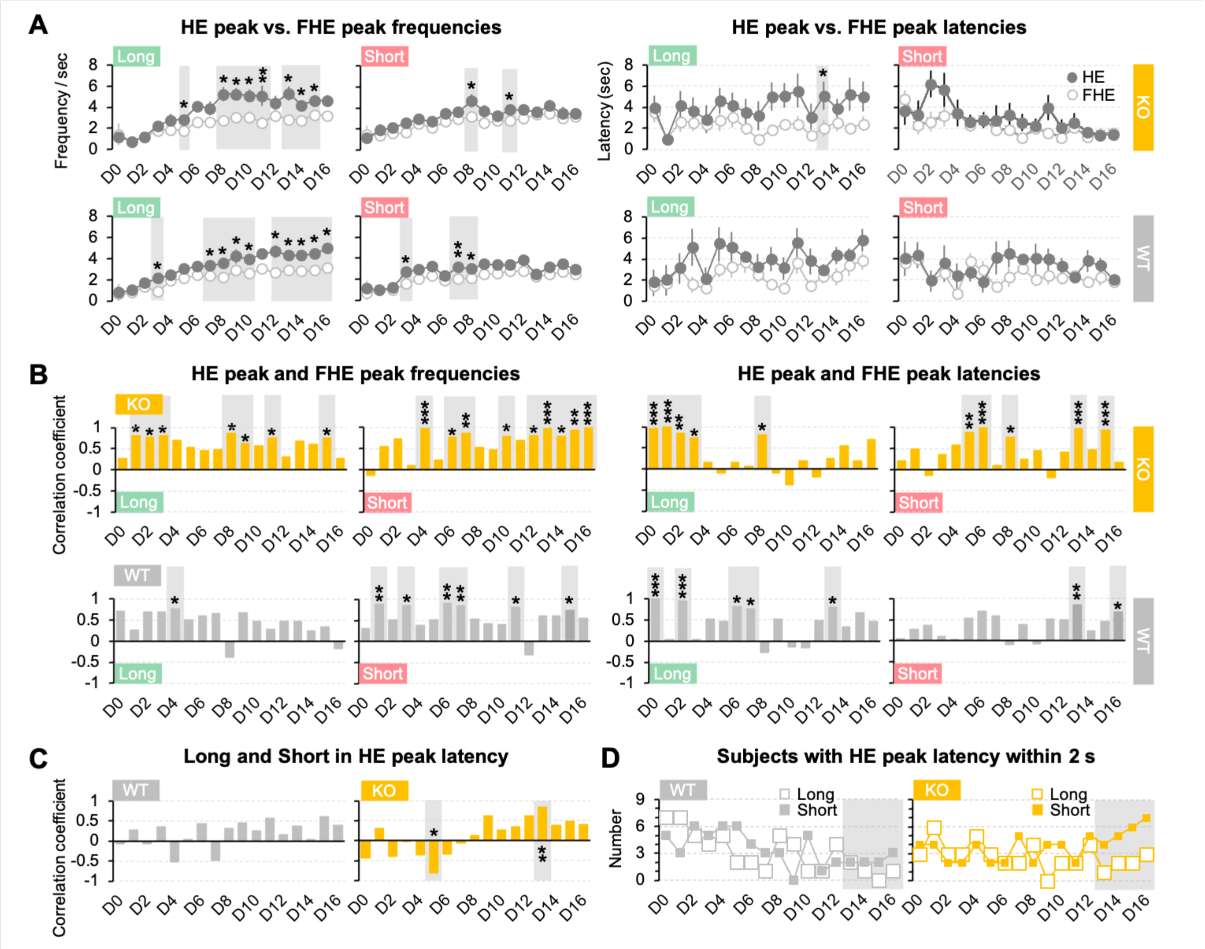


**Figure S3. HE and FHE peaks.** (A) Difference between the HE and FHE peaks were more prominent for frequency in both groups. (B) Correlation between the HE and FHE peaks in the late training days was more prominent in KO mice. (C) Correlations between HE peak latencies for Long and Short were not significant except for D5 and D13 in KO mice. (D) The number of KO mice with the HE peak latencies within 2 s increased linearly, from 4 subjects on D13 to 7 subjects on D16. See Table S3 for statistical details. Wilcoxon signed-ranks test, Mann–Whitney U test, and Spearman correlation test. *, $p < 0.05$; **, $p < 0.01$; ***, $p < 0.001$. Error bars denote 95% confidence intervals. Abbreviations: D, day; CS, conditioned stimulus; US, unconditioned stimulus; WT, wild-type mice; KO, CRBN KO mice; HE, head entry; FHE, fastest HE.

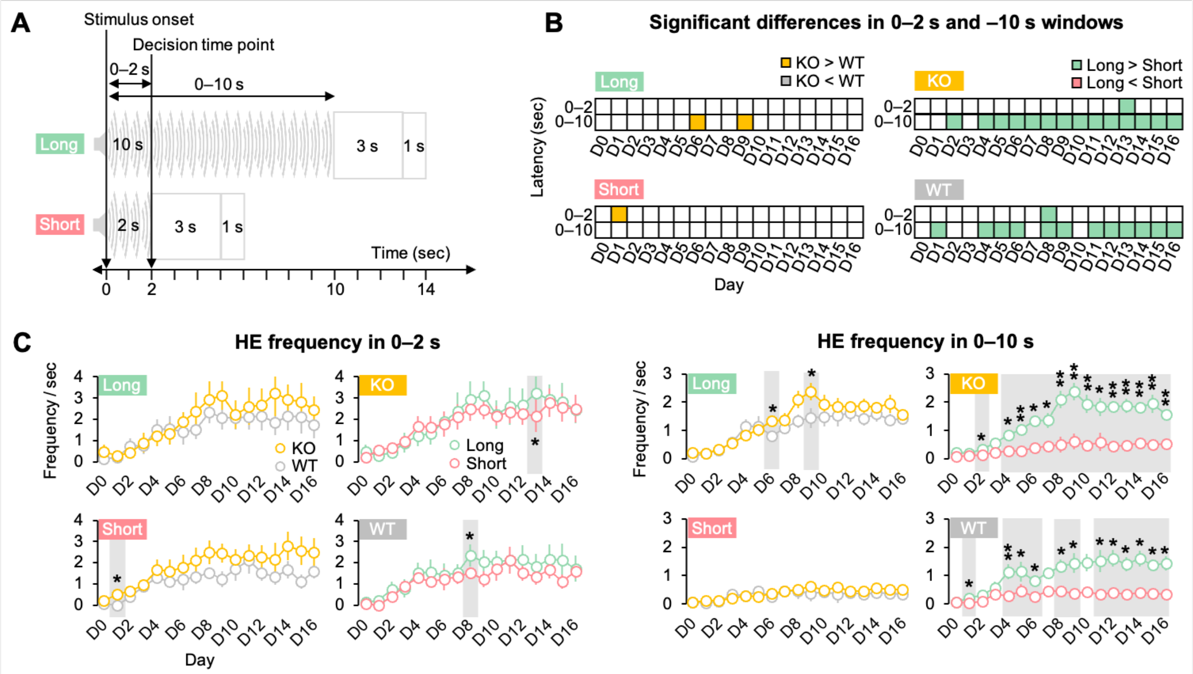


**Figure S4. HE frequencies in 0–2 and 0–14 after the stimulus onset.** (A) Analysis time windows were segmented into 0–2 s, 0–10 s, and successive 1-s bins (0–14 s). (B) Group and condition differences were rarely occurred in the 0–2 s window. In the 0–10 s window, both groups showed clear Long–Short differences. Significant differences are indicated by green- and orange-colored boxes. (C) The green- and orange-colored boxes in panel B were derived from the plots shown in panel C. See Table S4 for statistical details. Wilcoxon signed-ranks test and Mann–Whitney U test. *, $p < 0.05$; **, $p < 0.01$. Error bars denote 95% confidence intervals. Abbreviations: D, day; CS, conditioned stimulus; US, unconditioned stimulus; WT, wild-type mice; KO, CRBN KO mice; HE, head entry.

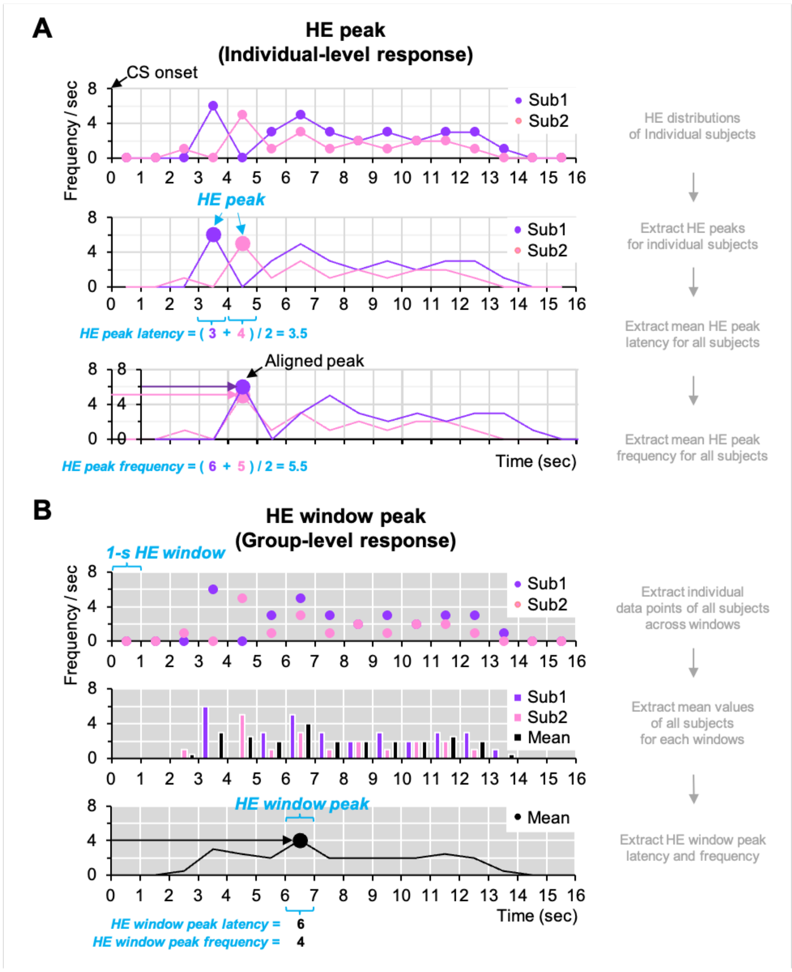


**Figure S5. Extraction of HE peak and HE window peak.** The HE peak latency may be not consistent with the HE window peak latency. Abbreviations: CS, conditioned stimulus; HE, head entry; Sub, subject.

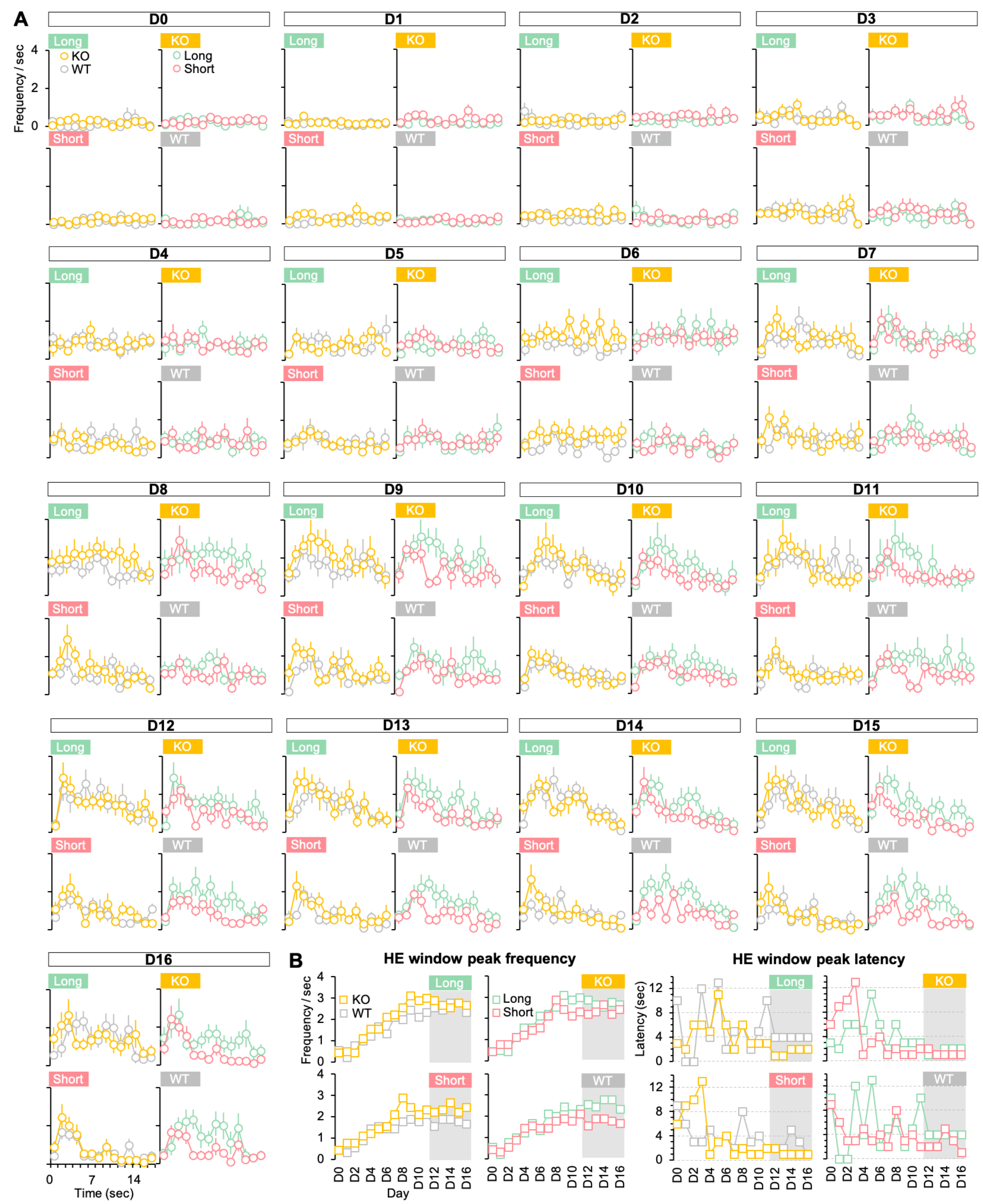


**Figure S6. Daily HE frequency change across 1-s latency bins and HE window peak.** (A) Distribution of HE frequency per 1-s bin was presented for individual training days. We defined a peak of frequency in each day as the HE window peak. (B) The frequency of HE window peak was lower in Short than in Long for WT mice in D12–D16 when HE frequency was saturated, which was different from the pattern in KO mice. In the latency of HE window peak, WT mice was relatively slower in Long, and KO mice responded quickly consistently for Short. HE window peak was exhibited within 2 s in KO mice, while it was consistent as 4–5 s for Long and was fluctuated from 2 s to 6 s for Short in WT mice. Error bars denote 95% confidence intervals. See also Figure 1H. Abbreviations: D, day; CS, conditioned stimulus; US, unconditioned stimulus; WT, wild-type mice; KO, KO mice; HE, head entry.

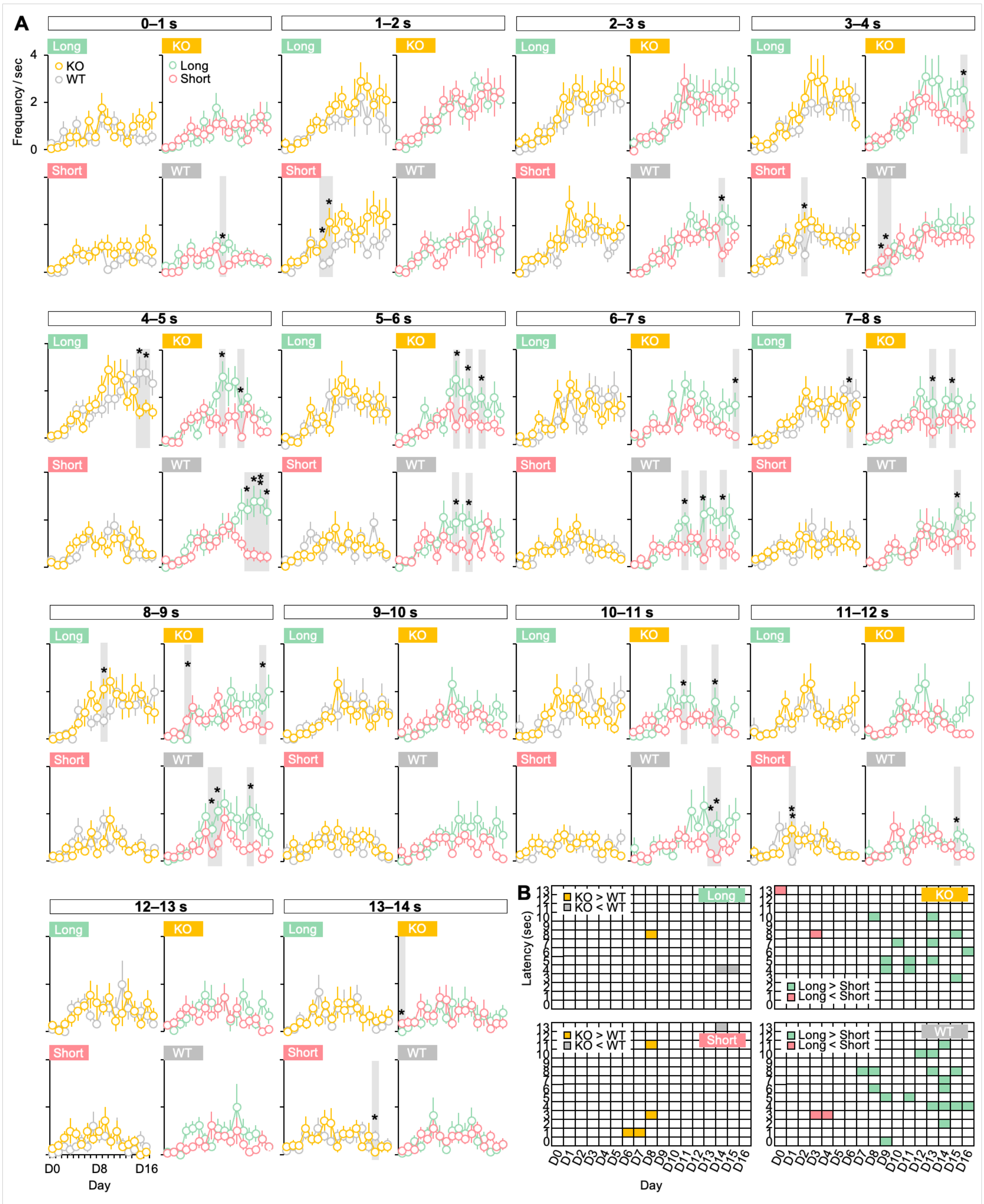


**Figure S7. Difference between groups and conditions for HE frequencies per 1-s latency bins across training days.** The green- and orange-colored boxes in panel B were derived from the plots shown in panel A. See also Figure 1I and Table S5 for full statistics. Wilcoxon signed-ranks test and Mann–Whitney U test, *, $p < 0.05$. Error bars denote 95% confidence intervals. Abbreviations: D, day; CS, conditioned stimulus; US, unconditioned stimulus; WT, wild-type mice; KO, CRBN KO mice.

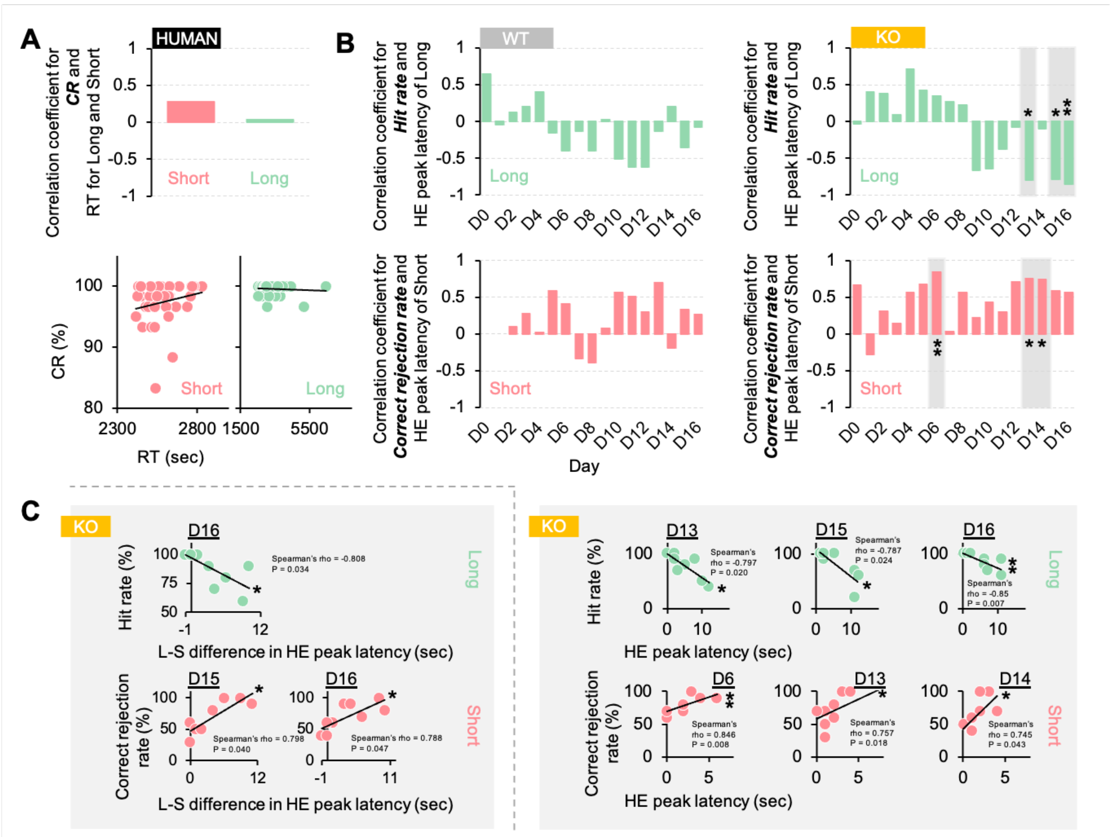


**Figure S8. Correlation between RT (latency) and CR (accuracy).** (A–B) Correlation between RT (HE peak latency) and CR (hit and correct rejection rates). (C) Correlation between the difference values between Long and Short (L-S difference) in HE peak latency and accuracy (hit and correct rejection rates), respectively. See Figure 3C and Table S7 for statistical details. Spearman correlation. *, $p < 0.05$; **, $p < 0.01$. Abbreviations: D, day; CS, conditioned stimulus; US, unconditioned stimulus; WT, wild-type mice; KO, CRBN KO mice; HE, head entry.
