## Supplementary tables for "Cereblon-related mild intellectual disability disrupts response inhibition and uniformity of group–individual strategies"

| Table number | Behavioral indices | | Contents |
|---|---|---|---|
| Table S1 | HE frequency | 1_1 | Total HE frequency |
| | | 1_2 | HE frequency in CS |
| | | 1_3 | HE frequency in pre-US |
| | | 1_4 | HE frequency in US |
| | | 1_5 | HE frequency in CS+pre-US |
| | | 1_6 | HE frequency in pre-US+US |
| | | 1_7 | HE frequency in pre-CS and post-CS |
| | | 1_8 | HE frequency in D0 and D1 |
| Table S2 | HE accuracy | 2_1 | Hit and correct rejection rate (CS) |
| | | 2_2 | Hit and correct rejection rate (CS+pre-US) |
| | | 2_3 | Hit rate vs. Correction rejection rate (CS) |
| Table S3 | HE and FHE peak | 3_1 | HE peak frequency |
| | | 3_2 | HE peak latency |
| | | 3_3 | FHE peak frequency |
| | | 3_4 | FHE peak latency |
| | | 3_5 | HE vs. FHE in frequencies |
| | | 3_6 | HE vs. FHE in latencies |
| | | 3_7 | Corrrelation betweeen HE and FHE peak frequencies |
| | | 3_8 | Corrrelation betweeen HE and FHE peak latencies |
| | | 3_9 | Correlation between Short and Long in HE peak latency |
| Table S4 | HE frequency for 2-s and 10-s latency windows | 4_1 | HE frequency in 0–2 s window |
| | | 4_2 | HE frequency in 0–10 s window |
| Table S5 | HE frequency for 1-s latency windows | 5_1 | HE frequency in 0-1 s |
| | | 5_2 | HE frequency in 1-2 s |
| | | 5_3 | HE frequency in 2-3 s |
| | | 5_4 | HE frequency in 3-4 s |
| | | 5_5 | HE frequency in 4-5 s |
| | | 5_6 | HE frequency in 5-6 s |
| | | 5_7 | HE frequency in 6-7 s |
| | | 5_8 | HE frequency in 7-8 s |
| | | 5_9 | HE frequency in 8-9 s |
| | | 5_10 | HE frequency in 9-10 s |
| | | 5_11 | HE frequency in 10-11 s |
| | | 5_12 | HE frequency in 11-12 s |
| | | 5_13 | HE frequency in 12-13 s |
| | | 5_14 | HE frequency in 13-14 s |
| Table S6 | CR and RT | 6_1 | CR |
| | | 6_2 | RT |
| | | 6_3 | Correlation between Long and Short in RT |
| Table S7 | Correlation between RT/Latency and CR/Accuracy | 7_1 | Human: Correlation between LATENCY (L–S difference in RT or RT) and ACCURACY (CR) |
| | | 7_2 | Mouse: Correlation between LATENCY (L–S difference in HE peak latency) and ACCURACY (hit and correct rejection rates) |
| | | 7_3 | Mouse: Correlation between LATENCY (HE peak latency) and ACCURACY (hit and correct rejection rates) |

Abbreviations: HE, head entry; CS, conditioned stimulus; US, unconditioned stimulus; CR, correct rate; RT, reaction time.

## 1_1

Total HE frequency

| Table | Related Figure | Day | Group | Mean | SEM | Stastistical test | Z | P | * |
|---|---|---|---|---|---|---|---|---|---|
| Table S1 | Figure 1D | D0 | KO | 54.722 | 8.545 | Mann-Whitney U Test | -1.988 | 0.047 | * |
| | | | WT | 30.000 | 10.141 | | | | |
| | | D1 | KO | 76.778 | 10.768 | | -1.634 | 0.102 | |
| | | | WT | 47.389 | 9.368 | | | | |
| | | D2 | KO | 95.222 | 10.613 | | -2.518 | 0.012 | * |
| | | | WT | 68.778 | 4.930 | | | | |
| | | D3 | KO | 108.778 | 16.707 | | -0.221 | 0.825 | |
| | | | WT | 125.778 | 17.680 | | | | |
| | | D4 | KO | 139.111 | 10.961 | | 0.000 | 1.000 | |
| | | | WT | 148.889 | 12.382 | | | | |
| | | D5 | KO | 119.222 | 10.801 | | -0.751 | 0.453 | |
| | | | WT | 142.889 | 18.461 | | | | |
| | | D6 | KO | 175.222 | 23.493 | | -1.810 | 0.070 | |
| | | | WT | 120.556 | 11.237 | | | | |
| | | D7 | KO | 153.333 | 19.258 | | -0.044 | 0.965 | |
| | | | WT | 153.667 | 18.066 | | | | |
| | | D8 | KO | 177.111 | 27.648 | | -0.442 | 0.659 | |
| | | | WT | 153.000 | 13.015 | | | | |
| | | D9 | KO | 175.556 | 32.384 | | -0.486 | 0.627 | |
| | | | WT | 149.111 | 18.530 | | | | |
| | | D10 | KO | 129.000 | 14.764 | | -0.221 | 0.825 | |
| | | | WT | 130.778 | 20.487 | | | | |
| | | D11 | KO | 129.111 | 15.123 | | -0.044 | 0.965 | |
| | | | WT | 137.222 | 23.956 | | | | |
| | | D12 | KO | 123.333 | 15.343 | | -0.132 | 0.895 | |
| | | | WT | 125.556 | 14.588 | | | | |
| | | D13 | KO | 115.889 | 19.062 | | 0.000 | 1.000 | |
| | | | WT | 106.111 | 9.398 | | | | |
| | | D14 | KO | 124.444 | 13.474 | | -0.575 | 0.566 | |
| | | | WT | 106.000 | 9.985 | | | | |
| | | D15 | KO | 103.111 | 13.007 | | -0.619 | 0.536 | |
| | | | WT | 114.111 | 14.496 | | | | |
| | | D16 | KO | 105.000 | 13.089 | | -0.044 | 0.965 | |
| | | | WT | 100.556 | 10.205 | | | | |

Abbreviations: HE, head entry; D, training day; KO, CRBN KO mice; WT, wild-type mice;  HE, head entry; CS, conditioned stimulus; US, unconditioned stimulus.

*, *p < 0.05, **p < 0.01, ***p < 0.001.

## 1_2

HE frequency in CS

| Table | Related Figure | Day | Conditon | Group | Mean | SEM | Stastistical test | Z | P | * |
|---|---|---|---|---|---|---|---|---|---|---|
| Table S1 | Figure 1D | D0 | Long | KO | 0.228 | 0.090 | Mann-Whitney U Test | -1.683 | 0.113 | |
| | | | | WT | 0.072 | 0.032 | | | | |
| | | D1 | Long | KO | 0.161 | 0.058 | | -0.544 | 0.605 | |
| | | | | WT | 0.111 | 0.045 | | | | |
| | | D2 | Long | KO | 0.272 | 0.082 | | -0.359 | 0.730 | |
| | | | | WT | 0.256 | 0.103 | | | | |
| | | D3 | Long | KO | 0.544 | 0.161 | | -0.671 | 0.546 | |
| | | | | WT | 0.400 | 0.133 | | | | |
| | | D4 | Long | KO | 0.811 | 0.176 | | -0.488 | 0.666 | |
| | | | | WT | 0.878 | 0.179 | | | | |
| | | D5 | Long | KO | 0.756 | 0.199 | | -0.490 | 0.666 | |
| | | | | WT | 0.856 | 0.209 | | | | |
| | | D6 | Long | KO | 1.356 | 0.209 | | -1.681 | 0.094 | |
| | | | | WT | 0.900 | 0.142 | | | | |
| | | D7 | Long | KO | 1.233 | 0.255 | | -0.312 | 0.796 | |
| | | | | WT | 1.144 | 0.166 | | | | |
| | | D8 | Long | KO | 2.133 | 0.378 | | -1.819 | 0.077 | |
| | | | | WT | 1.433 | 0.181 | | | | |
| | | D9 | Long | KO | 2.256 | 0.365 | | -1.459 | 0.161 | |
| | | | | WT | 1.567 | 0.297 | | | | |
| | | D10 | Long | KO | 1.833 | 0.370 | | -0.355 | 0.730 | |
| | | | | WT | 1.589 | 0.263 | | | | |
| | | D11 | Long | KO | 1.856 | 0.452 | | -0.177 | 0.863 | |
| | | | | WT | 1.689 | 0.337 | | | | |
| | | D12 | Long | KO | 1.678 | 0.213 | | -0.354 | 0.730 | |
| | | | | WT | 1.756 | 0.221 | | | | |
| | | D13 | Long | KO | 1.811 | 0.328 | | -0.354 | 0.730 | |
| | | | | WT | 1.622 | 0.179 | | | | |
| | | D14 | Long | KO | 1.700 | 0.260 | | -0.486 | 0.666 | |
| | | | | WT | 1.856 | 0.221 | | | | |
| | | D15 | Long | KO | 1.711 | 0.280 | | -0.267 | 0.796 | |
| | | | | WT | 1.744 | 0.221 | | | | |
| | | D16 | Long | KO | 1.522 | 0.146 | | -0.665 | 0.546 | |
| | | | | WT | 1.600 | 0.166 | | | | |
| Table S1 | Figure 1D | D0 | Short | KO | 0.139 | 0.084 | Mann-Whitney U Test | -1.032 | 0.489 | |
| | | | | WT | 0.056 | 0.056 | | | | |
| | | D1 | Short | KO | 0.306 | 0.116 | | -2.523 | 0.050 | |
| | | | | WT | 0.000 | 0.000 | | | | |
| | | D2 | Short | KO | 0.444 | 0.123 | | -1.518 | 0.161 | |
| | | | | WT | 0.194 | 0.069 | | | | |
| | | D3 | Short | KO | 0.556 | 0.176 | | -0.278 | 0.796 | |
| | | | | WT | 0.611 | 0.162 | | | | |
| | | D4 | Short | KO | 1.111 | 0.341 | | -0.495 | 0.666 | |
| | | | | WT | 1.111 | 0.217 | | | | |
| | | D5 | Short | KO | 0.833 | 0.186 | | -0.285 | 0.796 | |
| | | | | WT | 0.778 | 0.206 | | | | |
| | | D6 | Short | KO | 0.944 | 0.212 | | -1.423 | 0.190 | |
| | | | | WT | 0.556 | 0.212 | | | | |
| | | D7 | Short | KO | 1.500 | 0.354 | | -1.606 | 0.161 | |
| | | | | WT | 0.667 | 0.083 | | | | |
| | | D8 | Short | KO | 1.444 | 0.475 | | -0.270 | 0.796 | |
| | | | | WT | 1.111 | 0.247 | | | | |
| | | D9 | Short | KO | 1.778 | 0.426 | | -2.303 | 0.024 | * |
| | | | | WT | 0.667 | 0.144 | | | | |
| | | D10 | Short | KO | 1.444 | 0.306 | | -1.097 | 0.297 | |
| | | | | WT | 0.944 | 0.242 | | | | |
| | | D11 | Short | KO | 1.278 | 0.426 | | -0.136 | 0.931 | |
| | | | | WT | 1.111 | 0.217 | | | | |
| | | D12 | Short | KO | 1.444 | 0.327 | | -0.814 | 0.436 | |
| | | | | WT | 1.056 | 0.242 | | | | |
| | | D13 | Short | KO | 1.389 | 0.415 | | -1.078 | 0.297 | |
| | | | | WT | 0.722 | 0.147 | | | | |
| | | D14 | Short | KO | 1.889 | 0.423 | | -1.653 | 0.113 | |
| | | | | WT | 1.000 | 0.250 | | | | |
| | | D15 | Short | KO | 1.833 | 0.583 | | -1.033 | 0.340 | |
| | | | | WT | 0.833 | 0.264 | | | | |
| | | D16 | Short | KO | 1.667 | 0.486 | | -0.760 | 0.489 | |
| | | | | WT | 1.056 | 0.227 | | | | |

HE frequency in CS

| Table | Related Figure | Day | Group | Condition | Mean | SEM | Stastistical test | Z | P | * |
|---|---|---|---|---|---|---|---|---|---|---|
| Table S1 | Figure 1D | D0 | KO | Long | 0.228 | 0.090 | Wilcoxon Signed Ranks Test | -0.593 | 0.553 | |
| | | | | Short | 0.139 | 0.084 | | | | |
| | | D1 | KO | Long | 0.161 | 0.058 | | -1.014 | 0.310 | |
| | | | | Short | 0.306 | 0.116 | | | | |
| | | D2 | KO | Long | 0.272 | 0.082 | | -0.946 | 0.344 | |
| | | | | Short | 0.444 | 0.123 | | | | |
| | | D3 | KO | Long | 0.544 | 0.161 | | -0.070 | 0.944 | |
| | | | | Short | 0.556 | 0.176 | | | | |
| | | D4 | KO | Long | 0.811 | 0.176 | | -0.845 | 0.398 | |
| | | | | Short | 1.111 | 0.341 | | | | |
| | | D5 | KO | Long | 0.756 | 0.199 | | -0.560 | 0.575 | |
| | | | | Short | 0.833 | 0.186 | | | | |
| | | D6 | KO | Long | 1.356 | 0.209 | | -1.244 | 0.214 | |
| | | | | Short | 0.944 | 0.212 | | | | |
| | | D7 | KO | Long | 1.233 | 0.255 | | -0.772 | 0.440 | |
| | | | | Short | 1.500 | 0.354 | | | | |
| | | D8 | KO | Long | 2.133 | 0.378 | | -1.332 | 0.183 | |
| | | | | Short | 1.444 | 0.475 | | | | |
| | | D9 | KO | Long | 2.256 | 0.365 | | -1.122 | 0.262 | |
| | | | | Long | 1.778 | 0.426 | | | | |
| | | D10 | KO | Short | 1.833 | 0.370 | | -1.547 | 0.122 | |
| | | | | Long | 1.444 | 0.306 | | | | |
| | | D11 | KO | Short | 1.856 | 0.452 | | -1.542 | 0.123 | |
| | | | | Long | 1.278 | 0.426 | | | | |
| | | D12 | KO | Short | 1.678 | 0.213 | | -0.833 | 0.405 | |
| | | | | Long | 1.444 | 0.327 | | | | |
| | | D13 | KO | Short | 1.811 | 0.328 | | -1.483 | 0.138 | |
| | | | | Long | 1.389 | 0.415 | | | | |
| | | D14 | KO | Short | 1.700 | 0.260 | | -0.593 | 0.553 | |
| | | | | Long | 1.889 | 0.423 | | | | |
| | | D15 | KO | Short | 1.711 | 0.280 | | -0.296 | 0.767 | |
| | | | | Long | 1.833 | 0.583 | | | | |
| | | D16 | KO | Short | 1.522 | 0.146 | | 0.000 | 1.000 | |
| | | | | Long | 1.667 | 0.486 | | | | |
| Table S1 | Figure 1D | D0 | WT | Long | 0.072 | 0.032 | Wilcoxon Signed Ranks Test | -0.677 | 0.498 | |
| | | | | Short | 0.056 | 0.056 | | | | |
| | | D1 | WT | Long | 0.111 | 0.045 | | -2.214 | 0.027 | * |
| | | | | Short | 0.000 | 0.000 | | | | |
| | | D2 | WT | Long | 0.256 | 0.103 | | 0.000 | 1.000 | |
| | | | | Short | 0.194 | 0.069 | | | | |
| | | D3 | WT | Long | 0.400 | 0.133 | | -1.892 | 0.058 | |
| | | | | Short | 0.611 | 0.162 | | | | |
| | | D4 | WT | Long | 0.878 | 0.179 | | -2.120 | 0.034 | * |
| | | | | Short | 1.111 | 0.217 | | | | |
| | | D5 | WT | Long | 0.856 | 0.209 | | -0.416 | 0.677 | |
| | | | | Short | 0.778 | 0.206 | | | | |
| | | D6 | WT | Long | 0.900 | 0.142 | | -1.334 | 0.182 | |
| | | | | Short | 0.556 | 0.212 | | | | |
| | | D7 | WT | Long | 1.144 | 0.166 | | -2.077 | 0.038 | * |
| | | | | Short | 0.667 | 0.083 | | | | |
| | | D8 | WT | Long | 1.433 | 0.181 | | -0.912 | 0.362 | |
| | | | | Short | 1.111 | 0.247 | | | | |
| | | D9 | WT | Long | 1.567 | 0.297 | | -2.366 | 0.018 | * |
| | | | | Long | 0.667 | 0.144 | | | | |
| | | D10 | WT | Short | 1.589 | 0.263 | | -1.727 | 0.084 | |
| | | | | Long | 0.944 | 0.242 | | | | |
| | | D11 | WT | Short | 1.689 | 0.337 | | -1.365 | 0.172 | |
| | | | | Long | 1.111 | 0.217 | | | | |
| | | D12 | WT | Short | 1.756 | 0.221 | | -1.823 | 0.068 | |
| | | | | Long | 1.056 | 0.242 | | | | |
| | | D13 | WT | Short | 1.622 | 0.179 | | -2.524 | 0.012 | * |
| | | | | Long | 0.722 | 0.147 | | | | |
| | | D14 | WT | Short | 1.856 | 0.221 | | -2.429 | 0.015 | * |
| | | | | Long | 1.000 | 0.250 | | | | |
| | | D15 | WT | Short | 1.744 | 0.221 | | -2.312 | 0.021 | * |
| | | | | Long | 0.833 | 0.264 | | | | |
| | | D16 | WT | Short | 1.600 | 0.166 | | -1.755 | 0.079 | |
| | | | | Long | 1.056 | 0.227 | | | | |

## 1_3

HE frequency in pre-US

| Table | Related Figure | Day | Conditon | Group | Mean | SEM | Stastistical test | Z | P | * |
|---|---|---|---|---|---|---|---|---|---|---|
| Table S1 | Figure S2A | D0 | Long | KO | 0.222 | 0.073 | Mann-Whitney U Test | -1.415 | 0.222 | |
| | | | | WT | 0.370 | 0.331 | | | | |
| | | D1 | Long | KO | 0.093 | 0.049 | | 0.000 | 1.000 | |
| | | | | WT | 0.093 | 0.049 | | | | |
| | | D2 | Long | KO | 0.259 | 0.104 | | -1.222 | 0.297 | |
| | | | | WT | 0.074 | 0.040 | | | | |
| | | D3 | Long | KO | 0.333 | 0.147 | | -1.218 | 0.258 | |
| | | | | WT | 0.630 | 0.171 | | | | |
| | | D4 | Long | KO | 0.889 | 0.385 | | -0.585 | 0.605 | |
| | | | | WT | 0.815 | 0.158 | | | | |
| | | D5 | Long | KO | 1.259 | 0.265 | | -1.348 | 0.190 | |
| | | | | WT | 0.778 | 0.229 | | | | |
| | | D6 | Long | KO | 1.407 | 0.411 | | -1.717 | 0.094 | |
| | | | | WT | 0.444 | 0.124 | | | | |
| | | D7 | Long | KO | 1.222 | 0.503 | | -0.539 | 0.605 | |
| | | | | WT | 0.852 | 0.300 | | | | |
| | | D8 | Long | KO | 1.593 | 0.293 | | -1.249 | 0.222 | |
| | | | | WT | 1.074 | 0.318 | | | | |
| | | D9 | Long | KO | 1.815 | 0.545 | | -0.310 | 0.796 | |
| | | | | WT | 1.630 | 0.522 | | | | |
| | | D10 | Long | KO | 0.815 | 0.420 | | -0.722 | 0.489 | |
| | | | | WT | 1.222 | 0.423 | | | | |
| | | D11 | Long | KO | 0.778 | 0.397 | | -0.857 | 0.436 | |
| | | | | WT | 1.741 | 0.724 | | | | |
| | | D12 | Long | KO | 1.111 | 0.550 | | -0.405 | 0.730 | |
| | | | | WT | 1.000 | 0.329 | | | | |
| | | D13 | Long | KO | 1.000 | 0.272 | | -0.134 | 0.931 | |
| | | | | WT | 1.000 | 0.329 | | | | |
| | | D14 | Long | KO | 0.667 | 0.347 | | -1.419 | 0.190 | |
| | | | | WT | 1.037 | 0.312 | | | | |
| | | D15 | Long | KO | 0.963 | 0.349 | | -0.139 | 0.931 | |
| | | | | WT | 1.111 | 0.419 | | | | |
| | | D16 | Long | KO | 1.222 | 0.441 | | -0.319 | 0.796 | |
| | | | | WT | 0.889 | 0.324 | | | | |
| Table S1 | Figure S2A | D0 | Short | KO | 0.130 | 0.037 | Mann-Whitney U Test | -0.576 | 0.605 | |
| | | | | WT | 0.111 | 0.056 | | | | |
| | | D1 | Short | KO | 0.407 | 0.125 | | -2.146 | 0.050 | |
| | | | | WT | 0.074 | 0.049 | | | | |
| | | D2 | Short | KO | 0.370 | 0.117 | | -0.047 | 1.000 | |
| | | | | WT | 0.352 | 0.098 | | | | |
| | | D3 | Short | KO | 0.667 | 0.176 | | -0.181 | 0.863 | |
| | | | | WT | 0.630 | 0.161 | | | | |
| | | D4 | Short | KO | 0.963 | 0.171 | | -1.803 | 0.077 | |
| | | | | WT | 0.519 | 0.148 | | | | |
| | | D5 | Short | KO | 1.296 | 0.245 | | -0.179 | 0.863 | |
| | | | | WT | 1.222 | 0.351 | | | | |
| | | D6 | Short | KO | 1.333 | 0.266 | | -1.121 | 0.297 | |
| | | | | WT | 0.926 | 0.271 | | | | |
| | | D7 | Short | KO | 1.333 | 0.208 | | -0.134 | 0.931 | |
| | | | | WT | 1.259 | 0.253 | | | | |
| | | D8 | Short | KO | 2.037 | 0.321 | | -2.046 | 0.040 | * |
| | | | | WT | 1.185 | 0.201 | | | | |
| | | D9 | Short | KO | 1.667 | 0.289 | | -0.134 | 0.931 | |
| | | | | WT | 1.630 | 0.232 | | | | |
| | | D10 | Short | KO | 1.593 | 0.341 | | -0.089 | 0.931 | |
| | | | | WT | 1.630 | 0.275 | | | | |
| | | D11 | Short | KO | 1.704 | 0.406 | | -0.045 | 1.000 | |
| | | | | WT | 1.593 | 0.265 | | | | |
| | | D12 | Short | KO | 1.370 | 0.344 | | -0.223 | 0.863 | |
| | | | | WT | 1.370 | 0.280 | | | | |
| | | D13 | Short | KO | 1.556 | 0.248 | | -1.235 | 0.258 | |
| | | | | WT | 1.333 | 0.079 | | | | |
| | | D14 | Short | KO | 1.370 | 0.251 | | -0.947 | 0.387 | |
| | | | | WT | 0.963 | 0.232 | | | | |
| | | D15 | Short | KO | 1.074 | 0.241 | | -0.178 | 0.863 | |
| | | | | WT | 1.111 | 0.289 | | | | |
| | | D16 | Short | KO | 1.370 | 0.225 | | -0.458 | 0.666 | |
| | | | | WT | 1.148 | 0.137 | | | | |

HE frequency in pre-US

| Table | Related Figure | Day | Group | Condition | Mean | SEM | Stastistical test | Z | P | * |
|---|---|---|---|---|---|---|---|---|---|---|
| Table S1 | Figure S2A | D0 | KO | Long | 0.222 | 0.073 | Wilcoxon Signed Ranks Test | -0.850 | 0.395 | |
| | | | | Short | 0.130 | 0.037 | | | | |
| | | D1 | KO | Long | 0.093 | 0.049 | | -1.980 | 0.048 | |
| | | | | Short | 0.407 | 0.125 | | | | |
| | | D2 | KO | Long | 0.259 | 0.104 | | -0.948 | 0.343 | |
| | | | | Short | 0.370 | 0.117 | | | | |
| | | D3 | KO | Long | 0.333 | 0.147 | | -1.279 | 0.201 | |
| | | | | Short | 0.667 | 0.176 | | | | |
| | | D4 | KO | Long | 0.889 | 0.385 | | -0.943 | 0.345 | |
| | | | | Short | 0.963 | 0.171 | | | | |
| | | D5 | KO | Long | 1.259 | 0.265 | | -0.106 | 0.916 | |
| | | | | Short | 1.296 | 0.245 | | | | |
| | | D6 | KO | Long | 1.407 | 0.411 | | 0.000 | 1.000 | |
| | | | | Short | 1.333 | 0.266 | | | | |
| | | D7 | KO | Long | 1.222 | 0.503 | | -0.593 | 0.553 | |
| | | | | Short | 1.333 | 0.208 | | | | |
| | | D8 | KO | Long | 1.593 | 0.293 | | -1.126 | 0.260 | |
| | | | | Short | 2.037 | 0.321 | | | | |
| | | D9 | KO | Long | 1.815 | 0.545 | | -0.526 | 0.599 | |
| | | | | Long | 1.667 | 0.289 | | | | |
| | | D10 | KO | Short | 0.815 | 0.420 | | -1.550 | 0.121 | |
| | | | | Long | 1.593 | 0.341 | | | | |
| | | D11 | KO | Short | 0.778 | 0.397 | | -1.963 | 0.050 | |
| | | | | Long | 1.704 | 0.406 | | | | |
| | | D12 | KO | Short | 1.111 | 0.550 | | -1.193 | 0.233 | |
| | | | | Long | 1.370 | 0.344 | | | | |
| | | D13 | KO | Short | 1.000 | 0.272 | | -1.691 | 0.091 | |
| | | | | Long | 1.556 | 0.248 | | | | |
| | | D14 | KO | Short | 0.667 | 0.347 | | -1.548 | 0.122 | |
| | | | | Long | 1.370 | 0.251 | | | | |
| | | D15 | KO | Short | 0.963 | 0.349 | | -0.179 | 0.858 | |
| | | | | Long | 1.074 | 0.241 | | | | |
| | | D16 | KO | Short | 1.222 | 0.441 | | -0.178 | 0.858 | |
| | | | | Long | 1.370 | 0.225 | | | | |
| Table S1 | Figure S2A | D0 | WT | Long | 0.370 | 0.331 | Wilcoxon Signed Ranks Test | -0.447 | 0.655 | |
| | | | | Short | 0.111 | 0.056 | | | | |
| | | D1 | WT | Long | 0.093 | 0.049 | | -0.141 | 0.888 | |
| | | | | Short | 0.074 | 0.049 | | | | |
| | | D2 | WT | Long | 0.074 | 0.040 | | -2.214 | 0.027 | * |
| | | | | Short | 0.352 | 0.098 | | | | |
| | | D3 | WT | Long | 0.630 | 0.171 | | -0.086 | 0.931 | |
| | | | | Short | 0.630 | 0.161 | | | | |
| | | D4 | WT | Long | 0.815 | 0.158 | | -1.347 | 0.178 | |
| | | | | Short | 0.519 | 0.148 | | | | |
| | | D5 | WT | Long | 0.778 | 0.229 | | -1.897 | 0.058 | |
| | | | | Short | 1.222 | 0.351 | | | | |
| | | D6 | WT | Long | 0.444 | 0.124 | | -1.420 | 0.156 | |
| | | | | Short | 0.926 | 0.271 | | | | |
| | | D7 | WT | Long | 0.852 | 0.300 | | -1.606 | 0.108 | |
| | | | | Short | 1.259 | 0.253 | | | | |
| | | D8 | WT | Long | 1.074 | 0.318 | | -0.210 | 0.833 | |
| | | | | Short | 1.185 | 0.201 | | | | |
| | | D9 | WT | Long | 1.630 | 0.522 | | -0.059 | 0.953 | |
| | | | | Long | 1.630 | 0.232 | | | | |
| | | D10 | WT | Short | 1.222 | 0.423 | | -0.847 | 0.397 | |
| | | | | Long | 1.630 | 0.275 | | | | |
| | | D11 | WT | Short | 1.741 | 0.724 | | -0.070 | 0.944 | |
| | | | | Long | 1.593 | 0.265 | | | | |
| | | D12 | WT | Short | 1.000 | 0.329 | | -1.016 | 0.310 | |
| | | | | Long | 1.370 | 0.280 | | | | |
| | | D13 | WT | Short | 1.000 | 0.329 | | -1.018 | 0.309 | |
| | | | | Long | 1.333 | 0.079 | | | | |
| | | D14 | WT | Short | 1.037 | 0.312 | | -0.135 | 0.893 | |
| | | | | Long | 0.963 | 0.232 | | | | |
| | | D15 | WT | Short | 1.111 | 0.419 | | -0.059 | 0.953 | |
| | | | | Long | 1.111 | 0.289 | | | | |
| | | D16 | WT | Short | 0.889 | 0.324 | | -0.895 | 0.371 | |
| | | | | Long | 1.148 | 0.137 | | | | |

## 1_4

**HE frequency in US**

| Table | Related Figure | Day | Conditon | Group | Mean | SEM | Stastistical test | Z | P | * |
|---|---|---|---|---|---|---|---|---|---|---|
| Table S1 | Figure S2A | D0 | Long | KO | 0.000 | 0.000 | Mann-Whitney U Test | -1.000 | 0.730 | |
| | | | | WT | 0.056 | 0.056 | | | | |
| | | D1 | Long | KO | 0.167 | 0.118 | | -0.544 | 0.730 | |
| | | | | WT | 0.111 | 0.111 | | | | |
| | | D2 | Long | KO | 0.389 | 0.232 | | -0.503 | 0.666 | |
| | | | | WT | 0.556 | 0.256 | | | | |
| | | D3 | Long | KO | 0.333 | 0.236 | | 0.000 | 1.000 | |
| | | | | WT | 0.333 | 0.236 | | | | |
| | | D4 | Long | KO | 1.000 | 0.333 | | -0.095 | 0.931 | |
| | | | | WT | 0.889 | 0.261 | | | | |
| | | D5 | Long | KO | 0.444 | 0.242 | | -1.219 | 0.297 | |
| | | | | WT | 1.667 | 0.726 | | | | |
| | | D6 | Long | KO | 1.111 | 0.351 | | -1.046 | 0.340 | |
| | | | | WT | 0.778 | 0.465 | | | | |
| | | D7 | Long | KO | 0.556 | 0.242 | | -1.068 | 0.387 | |
| | | | | WT | 0.222 | 0.147 | | | | |
| | | D8 | Long | KO | 1.222 | 0.662 | | -0.190 | 0.863 | |
| | | | | WT | 1.000 | 0.408 | | | | |
| | | D9 | Long | KO | 0.889 | 0.512 | | -0.158 | 0.931 | |
| | | | | WT | 1.111 | 0.655 | | | | |
| | | D10 | Long | KO | 1.222 | 0.662 | | -0.252 | 0.863 | |
| | | | | WT | 0.889 | 0.455 | | | | |
| | | D11 | Long | KO | 1.000 | 0.373 | | -0.512 | 0.666 | |
| | | | | WT | 1.444 | 0.530 | | | | |
| | | D12 | Long | KO | 0.556 | 0.556 | | -0.910 | 0.546 | |
| | | | | WT | 0.556 | 0.338 | | | | |
| | | D13 | Long | KO | 0.667 | 0.441 | | -0.527 | 0.666 | |
| | | | | WT | 0.667 | 0.289 | | | | |
| | | D14 | Long | KO | 0.444 | 0.338 | | -0.680 | 0.666 | |
| | | | | WT | 0.111 | 0.111 | | | | |
| | | D15 | Long | KO | 0.556 | 0.338 | | -0.620 | 0.666 | |
| | | | | WT | 0.222 | 0.147 | | | | |
| | | D16 | Long | KO | 0.778 | 0.324 | | -0.591 | 0.605 | |
| | | | | WT | 0.444 | 0.176 | | | | |
| Table S1 | Figure S2A | D0 | Short | KO | 0.056 | 0.056 | Mann-Whitney U Test | -1.627 | 0.222 | |
| | | | | WT | 0.333 | 0.144 | | | | |
| | | D1 | Short | KO | 0.278 | 0.147 | | 0.000 | 1.000 | |
| | | | | WT | 0.278 | 0.147 | | | | |
| | | D2 | Short | KO | 0.500 | 0.204 | | -0.911 | 0.436 | |
| | | | | WT | 0.222 | 0.121 | | | | |
| | | D3 | Short | KO | 0.556 | 0.338 | | -0.264 | 0.863 | |
| | | | | WT | 0.889 | 0.455 | | | | |
| | | D4 | Short | KO | 0.444 | 0.176 | | -1.380 | 0.222 | |
| | | | | WT | 1.444 | 0.475 | | | | |
| | | D5 | Short | KO | 0.778 | 0.278 | | -1.101 | 0.340 | |
| | | | | WT | 1.222 | 0.278 | | | | |
| | | D6 | Short | KO | 1.111 | 0.423 | | -1.669 | 0.136 | |
| | | | | WT | 0.444 | 0.338 | | | | |
| | | D7 | Short | KO | 1.333 | 0.236 | | -0.047 | 1.000 | |
| | | | | WT | 1.333 | 0.333 | | | | |
| | | D8 | Short | KO | 1.667 | 0.707 | | -0.232 | 0.863 | |
| | | | | WT | 0.889 | 0.261 | | | | |
| | | D9 | Short | KO | 0.778 | 0.278 | | 0.000 | 1.000 | |
| | | | | WT | 0.778 | 0.278 | | | | |
| | | D10 | Short | KO | 1.444 | 0.475 | | -0.552 | 0.605 | |
| | | | | WT | 1.000 | 0.289 | | | | |
| | | D11 | Short | KO | 0.889 | 0.389 | | -0.949 | 0.436 | |
| | | | | WT | 0.444 | 0.338 | | | | |
| | | D12 | Short | KO | 1.222 | 0.364 | | -0.233 | 0.863 | |
| | | | | WT | 1.333 | 0.373 | | | | |
| | | D13 | Short | KO | 0.778 | 0.278 | | -0.687 | 0.546 | |
| | | | | WT | 0.556 | 0.294 | | | | |
| | | D14 | Short | KO | 0.778 | 0.222 | | -2.004 | 0.063 | |
| | | | | WT | 1.889 | 0.423 | | | | |
| | | D15 | Short | KO | 0.889 | 0.351 | | -0.191 | 0.863 | |
| | | | | WT | 0.778 | 0.324 | | | | |
| | | D16 | Short | KO | 0.556 | 0.176 | | -0.653 | 0.605 | |
| | | | | WT | 0.444 | 0.242 | | | | |

**HE frequency in US**

| Table | Related Figure | Day | Group | Condition | Mean | SEM | Stastistical test | Z | P | * |
|---|---|---|---|---|---|---|---|---|---|---|
| Table S1 | Figure S2A | D0 | KO | Long | 0.000 | 0.000 | Wilcoxon Signed Ranks Test | -1.000 | 0.317 | |
| | | | | Short | 0.056 | 0.056 | | | | |
| | | D1 | KO | Long | 0.167 | 0.118 | | -0.552 | 0.581 | |
| | | | | Short | 0.278 | 0.147 | | | | |
| | | D2 | KO | Long | 0.389 | 0.232 | | -0.557 | 0.577 | |
| | | | | Short | 0.500 | 0.204 | | | | |
| | | D3 | KO | Long | 0.333 | 0.236 | | -0.378 | 0.705 | |
| | | | | Short | 0.556 | 0.338 | | | | |
| | | D4 | KO | Long | 1.000 | 0.333 | | -1.186 | 0.236 | |
| | | | | Short | 0.444 | 0.176 | | | | |
| | | D5 | KO | Long | 0.444 | 0.242 | | -1.134 | 0.257 | |
| | | | | Short | 0.778 | 0.278 | | | | |
| | | D6 | KO | Long | 1.111 | 0.351 | | 0.000 | 1.000 | |
| | | | | Short | 1.111 | 0.423 | | | | |
| | | D7 | KO | Long | 0.556 | 0.242 | | -2.333 | 0.020 | * |
| | | | | Short | 1.333 | 0.236 | | | | |
| | | D8 | KO | Long | 1.222 | 0.662 | | -0.690 | 0.490 | |
| | | | | Short | 1.667 | 0.707 | | | | |
| | | D9 | KO | Long | 0.889 | 0.512 | | 0.000 | 1.000 | |
| | | | | Long | 0.778 | 0.278 | | | | |
| | | D10 | KO | Short | 1.222 | 0.662 | | -0.324 | 0.746 | |
| | | | | Long | 1.444 | 0.475 | | | | |
| | | D11 | KO | Short | 1.000 | 0.373 | | -0.184 | 0.854 | |
| | | | | Long | 0.889 | 0.389 | | | | |
| | | D12 | KO | Short | 0.556 | 0.556 | | -0.954 | 0.340 | |
| | | | | Long | 1.222 | 0.364 | | | | |
| | | D13 | KO | Short | 0.667 | 0.441 | | -0.171 | 0.864 | |
| | | | | Long | 0.778 | 0.278 | | | | |
| | | D14 | KO | Short | 0.444 | 0.338 | | -0.791 | 0.429 | |
| | | | | Long | 0.778 | 0.222 | | | | |
| | | D15 | KO | Short | 0.556 | 0.338 | | -0.647 | 0.518 | |
| | | | | Long | 0.889 | 0.351 | | | | |
| | | D16 | KO | Short | 0.778 | 0.324 | | -0.707 | 0.480 | |
| | | | | Long | 0.556 | 0.176 | | | | |
| Table S1 | Figure S2A | D0 | WT | Long | 0.056 | 0.056 | Wilcoxon Signed Ranks Test | -1.518 | 0.129 | |
| | | | | Short | 0.333 | 0.144 | | | | |
| | | D1 | WT | Long | 0.111 | 0.111 | | -0.756 | 0.450 | |
| | | | | Short | 0.278 | 0.147 | | | | |
| | | D2 | WT | Long | 0.556 | 0.256 | | -1.633 | 0.102 | |
| | | | | Short | 0.222 | 0.121 | | | | |
| | | D3 | WT | Long | 0.333 | 0.236 | | -1.289 | 0.197 | |
| | | | | Short | 0.889 | 0.455 | | | | |
| | | D4 | WT | Long | 0.889 | 0.261 | | -0.850 | 0.396 | |
| | | | | Short | 1.444 | 0.475 | | | | |
| | | D5 | WT | Long | 1.667 | 0.726 | | -0.514 | 0.607 | |
| | | | | Short | 1.222 | 0.278 | | | | |
| | | D6 | WT | Long | 0.778 | 0.465 | | -0.542 | 0.588 | |
| | | | | Short | 0.444 | 0.338 | | | | |
| | | D7 | WT | Long | 0.222 | 0.147 | | -2.271 | 0.023 | * |
| | | | | Short | 1.333 | 0.333 | | | | |
| | | D8 | WT | Long | 1.000 | 0.408 | | -0.108 | 0.914 | |
| | | | | Short | 0.889 | 0.261 | | | | |
| | | D9 | WT | Long | 1.111 | 0.655 | | -0.422 | 0.673 | |
| | | | | Short | 0.778 | 0.278 | | | | |
| | | D10 | WT | Long | 0.889 | 0.455 | | -0.172 | 0.863 | |
| | | | | Long | 1.000 | 0.289 | | | | |
| | | D11 | WT | Short | 1.444 | 0.530 | | -1.841 | 0.066 | |
| | | | | Long | 0.444 | 0.338 | | | | |
| | | D12 | WT | Short | 0.556 | 0.338 | | -1.063 | 0.288 | |
| | | | | Long | 1.333 | 0.373 | | | | |
| | | D13 | WT | Short | 0.667 | 0.289 | | -0.276 | 0.783 | |
| | | | | Long | 0.556 | 0.294 | | | | |
| | | D14 | WT | Short | 0.111 | 0.111 | | -2.555 | 0.011 | * |
| | | | | Long | 1.889 | 0.423 | | | | |
| | | D15 | WT | Short | 0.222 | 0.147 | | -1.265 | 0.206 | |
| | | | | Long | 0.778 | 0.324 | | | | |
| | | D16 | WT | Short | 0.444 | 0.176 | | 0.000 | 1.000 | |
| | | | | Long | 0.444 | 0.242 | | | | |

## 1_5

**HE frequency in CS + pre-US**

| Table | Related Figure | Day | Conditon | Group | Mean | SEM | Stastistical test | Z | P | * |
|---|---|---|---|---|---|---|---|---|---|---|
| Table S1 | Figure S2A | D0 | Long | KO | 0.226 | 0.084 | Mann-Whitney U Test | -1.656 | 0.113 | |
| | | | | WT | 0.141 | 0.099 | | | | |
| | | D1 | Long | KO | 0.145 | 0.054 | | -0.361 | 0.730 | |
| | | | | WT | 0.107 | 0.042 | | | | |
| | | D2 | Long | KO | 0.269 | 0.075 | | -0.535 | 0.605 | |
| | | | | WT | 0.214 | 0.078 | | | | |
| | | D3 | Long | KO | 0.487 | 0.130 | | -0.223 | 0.863 | |
| | | | | WT | 0.453 | 0.130 | | | | |
| | | D4 | Long | KO | 0.829 | 0.199 | | -0.133 | 0.931 | |
| | | | | WT | 0.863 | 0.152 | | | | |
| | | D5 | Long | KO | 0.872 | 0.187 | | -0.311 | 0.796 | |
| | | | | WT | 0.829 | 0.187 | | | | |
| | | D6 | Long | KO | 1.368 | 0.237 | | -1.642 | 0.113 | |
| | | | | WT | 0.795 | 0.113 | | | | |
| | | D7 | Long | KO | 1.231 | 0.268 | | -0.222 | 0.863 | |
| | | | | WT | 1.077 | 0.182 | | | | |
| | | D8 | Long | KO | 2.009 | 0.323 | | -2.210 | 0.024 | * |
| | | | | WT | 1.350 | 0.202 | | | | |
| | | D9 | Long | KO | 2.154 | 0.354 | | -1.284 | 0.222 | |
| | | | | WT | 1.581 | 0.329 | | | | |
| | | D10 | Long | KO | 1.598 | 0.299 | | -0.310 | 0.796 | |
| | | | | WT | 1.504 | 0.269 | | | | |
| | | D11 | Long | KO | 1.607 | 0.391 | | -0.133 | 0.931 | |
| | | | | WT | 1.701 | 0.404 | | | | |
| | | D12 | Long | KO | 1.538 | 0.248 | | 0.000 | 1.000 | |
| | | | | WT | 1.581 | 0.190 | | | | |
| | | D13 | Long | KO | 1.624 | 0.270 | | -0.089 | 0.931 | |
| | | | | WT | 1.479 | 0.185 | | | | |
| | | D14 | Long | KO | 1.462 | 0.231 | | -0.531 | 0.605 | |
| | | | | WT | 1.667 | 0.208 | | | | |
| | | D15 | Long | KO | 1.538 | 0.198 | | -0.177 | 0.863 | |
| | | | | WT | 1.598 | 0.220 | | | | |
| | | D16 | Long | KO | 1.453 | 0.158 | | -0.398 | 0.730 | |
| | | | | WT | 1.427 | 0.181 | | | | |
| Table S1 | Figure S2A | D0 | Short | KO | 0.133 | 0.041 | Mann-Whitney U Test | -1.122 | 0.297 | |
| | | | | WT | 0.089 | 0.048 | | | | |
| | | D1 | Short | KO | 0.367 | 0.109 | | -2.507 | 0.019 | * |
| | | | | WT | 0.044 | 0.029 | | | | |
| | | D2 | Short | KO | 0.400 | 0.099 | | -0.813 | 0.436 | |
| | | | | WT | 0.289 | 0.065 | | | | |
| | | D3 | Short | KO | 0.600 | 0.133 | | -0.091 | 0.931 | |
| | | | | WT | 0.622 | 0.122 | | | | |
| | | D4 | Short | KO | 1.000 | 0.186 | | -1.541 | 0.136 | |
| | | | | WT | 0.756 | 0.104 | | | | |
| | | D5 | Short | KO | 1.111 | 0.180 | | -0.621 | 0.546 | |
| | | | | WT | 1.044 | 0.262 | | | | |
| | | D6 | Short | KO | 1.156 | 0.194 | | -1.336 | 0.190 | |
| | | | | WT | 0.778 | 0.209 | | | | |
| | | D7 | Short | KO | 1.400 | 0.240 | | -0.905 | 0.387 | |
| | | | | WT | 1.000 | 0.149 | | | | |
| | | D8 | Short | KO | 1.778 | 0.261 | | -1.957 | 0.050 | |
| | | | | WT | 1.156 | 0.152 | | | | |
| | | D9 | Short | KO | 1.711 | 0.304 | | -1.378 | 0.190 | |
| | | | | WT | 1.244 | 0.172 | | | | |
| | | D10 | Short | KO | 1.533 | 0.269 | | -0.715 | 0.489 | |
| | | | | WT | 1.333 | 0.170 | | | | |
| | | D11 | Short | KO | 1.533 | 0.320 | | -0.133 | 0.931 | |
| | | | | WT | 1.400 | 0.231 | | | | |
| | | D12 | Short | KO | 1.400 | 0.240 | | -0.668 | 0.546 | |
| | | | | WT | 1.244 | 0.124 | | | | |
| | | D13 | Short | KO | 1.489 | 0.186 | | -1.935 | 0.063 | |
| | | | | WT | 1.089 | 0.075 | | | | |
| | | D14 | Short | KO | 1.578 | 0.209 | | -2.002 | 0.050 | |
| | | | | WT | 0.978 | 0.122 | | | | |
| | | D15 | Short | KO | 1.378 | 0.310 | | -1.201 | 0.258 | |
| | | | | WT | 1.000 | 0.167 | | | | |
| | | D16 | Short | KO | 1.489 | 0.214 | | -2.229 | 0.031 | |
| | | | | WT | 1.111 | 0.068 | | | | |

**HE frequency in CS + pre-US**

| Table | Related Figure | Day | Group | Condition | Mean | SEM | Stastistical test | Z | P | * |
|---|---|---|---|---|---|---|---|---|---|---|
| Table S1 | Figure S2A | D0 | KO | Long | 0.226 | 0.084 | Wilcoxon Signed Ranks Test | -1.008 | 0.314 | |
| | | | | Short | 0.133 | 0.041 | | | | |
| | | D1 | KO | Long | 0.145 | 0.054 | | -1.540 | 0.123 | |
| | | | | Short | 0.367 | 0.109 | | | | |
| | | D2 | KO | Long | 0.269 | 0.075 | | -1.823 | 0.068 | |
| | | | | Short | 0.400 | 0.099 | | | | |
| | | D3 | KO | Long | 0.487 | 0.130 | | -0.420 | 0.674 | |
| | | | | Short | 0.600 | 0.133 | | | | |
| | | D4 | KO | Long | 0.829 | 0.199 | | -1.352 | 0.176 | |
| | | | | Short | 1.000 | 0.186 | | | | |
| | | D5 | KO | Long | 0.872 | 0.187 | | -1.718 | 0.086 | |
| | | | | Short | 1.111 | 0.180 | | | | |
| | | D6 | KO | Long | 1.368 | 0.237 | | -0.770 | 0.441 | |
| | | | | Short | 1.156 | 0.194 | | | | |
| | | D7 | KO | Long | 1.231 | 0.268 | | -1.007 | 0.314 | |
| | | | | Short | 1.400 | 0.240 | | | | |
| | | D8 | KO | Long | 2.009 | 0.323 | | -1.007 | 0.314 | |
| | | | | Short | 1.778 | 0.261 | | | | |
| | | D9 | KO | Long | 2.154 | 0.354 | | -1.960 | 0.050 | |
| | | | | Long | 1.711 | 0.304 | | | | |
| | | D10 | KO | Short | 1.598 | 0.299 | | -0.059 | 0.953 | |
| | | | | Long | 1.533 | 0.269 | | | | |
| | | D11 | KO | Short | 1.607 | 0.391 | | -0.593 | 0.553 | |
| | | | | Long | 1.533 | 0.320 | | | | |
| | | D12 | KO | Short | 1.538 | 0.248 | | -0.356 | 0.722 | |
| | | | | Long | 1.400 | 0.240 | | | | |
| | | D13 | KO | Short | 1.624 | 0.270 | | -0.420 | 0.674 | |
| | | | | Long | 1.489 | 0.186 | | | | |
| | | D14 | KO | Short | 1.462 | 0.231 | | -0.474 | 0.635 | |
| | | | | Long | 1.578 | 0.209 | | | | |
| | | D15 | KO | Short | 1.538 | 0.198 | | -0.296 | 0.767 | |
| | | | | Long | 1.378 | 0.310 | | | | |
| | | D16 | KO | Short | 1.453 | 0.158 | | -0.178 | 0.859 | |
| | | | | Long | 1.489 | 0.214 | | | | |
| Table S1 | Figure S2A | D0 | WT | Long | 0.141 | 0.099 | Wilcoxon Signed Ranks Test | -0.135 | 0.893 | |
| | | | | Short | 0.089 | 0.048 | | | | |
| | | D1 | WT | Long | 0.107 | 0.042 | | -1.014 | 0.310 | |
| | | | | Short | 0.044 | 0.029 | | | | |
| | | D2 | WT | Long | 0.214 | 0.078 | | -1.532 | 0.125 | |
| | | | | Short | 0.289 | 0.065 | | | | |
| | | D3 | WT | Long | 0.453 | 0.130 | | -1.400 | 0.161 | |
| | | | | Short | 0.622 | 0.122 | | | | |
| | | D4 | WT | Long | 0.863 | 0.152 | | -0.771 | 0.441 | |
| | | | | Short | 0.756 | 0.104 | | | | |
| | | D5 | WT | Long | 0.829 | 0.187 | | -0.770 | 0.441 | |
| | | | | Short | 1.044 | 0.262 | | | | |
| | | D6 | WT | Long | 0.795 | 0.113 | | -0.178 | 0.859 | |
| | | | | Short | 0.778 | 0.209 | | | | |
| | | D7 | WT | Long | 1.077 | 0.182 | | -0.415 | 0.678 | |
| | | | | Short | 1.000 | 0.149 | | | | |
| | | D8 | WT | Long | 1.350 | 0.202 | | -0.840 | 0.401 | |
| | | | | Short | 1.156 | 0.152 | | | | |
| | | D9 | WT | Long | 1.581 | 0.329 | | -1.245 | 0.213 | |
| | | | | Short | 1.244 | 0.172 | | | | |
| | | D10 | WT | Short | 1.504 | 0.269 | | -0.280 | 0.779 | |
| | | | | Long | 1.333 | 0.170 | | | | |
| | | D11 | WT | Short | 1.701 | 0.404 | | -0.593 | 0.553 | |
| | | | | Long | 1.400 | 0.231 | | | | |
| | | D12 | WT | Short | 1.581 | 0.190 | | -1.362 | 0.173 | |
| | | | | Long | 1.244 | 0.124 | | | | |
| | | D13 | WT | Short | 1.479 | 0.185 | | -2.073 | 0.038 | |
| | | | | Long | 1.089 | 0.075 | | | | |
| | | D14 | WT | Short | 1.667 | 0.208 | | -2.310 | 0.021 | * |
| | | | | Long | 0.978 | 0.122 | | | | |
| | | D15 | WT | Short | 1.598 | 0.220 | | -1.836 | 0.066 | |
| | | | | Long | 1.000 | 0.167 | | | | |
| | | D16 | WT | Short | 1.427 | 0.181 | | -1.423 | 0.155 | |
| | | | | Long | 1.111 | 0.068 | | | | |

## 1_6

**HE frequency in pre-US + US**

| Table | Related Figure | Day | Conditon | Group | Mean | SEM | Stastistical test | Z | P | * |
|---|---|---|---|---|---|---|---|---|---|---|
| Table S1 | Figure S2A | D0 | Long | KO | 0.167 | 0.055 | Mann-Whitney U Test | -1.141 | 0.297 | |
| | | | | WT | 0.292 | 0.247 | | | | |
| | | D1 | Long | KO | 0.111 | 0.044 | | -0.201 | 0.863 | |
| | | | | WT | 0.100 | 0.040 | | | | |
| | | D2 | Long | KO | 0.292 | 0.114 | | -0.419 | 0.730 | |
| | | | | WT | 0.194 | 0.073 | | | | |
| | | D3 | Long | KO | 0.333 | 0.125 | | -1.103 | 0.297 | |
| | | | | WT | 0.556 | 0.149 | | | | |
| | | D4 | Long | KO | 0.917 | 0.315 | | -0.268 | 0.796 | |
| | | | | WT | 0.833 | 0.144 | | | | |
| | | D5 | Long | KO | 1.056 | 0.212 | | -0.178 | 0.863 | |
| | | | | WT | 1.000 | 0.306 | | | | |
| | | D6 | Long | KO | 1.333 | 0.351 | | -1.838 | 0.077 | |
| | | | | WT | 0.528 | 0.188 | | | | |
| | | D7 | Long | KO | 1.056 | 0.393 | | -0.809 | 0.436 | |
| | | | | WT | 0.694 | 0.246 | | | | |
| | | D8 | Long | KO | 1.500 | 0.344 | | -0.889 | 0.387 | |
| | | | | WT | 1.056 | 0.311 | | | | |
| | | D9 | Long | KO | 1.583 | 0.460 | | -0.664 | 0.546 | |
| | | | | WT | 1.500 | 0.550 | | | | |
| | | D10 | Long | KO | 0.917 | 0.470 | | -0.448 | 0.666 | |
| | | | | WT | 1.139 | 0.384 | | | | |
| | | D11 | Long | KO | 0.833 | 0.323 | | -0.535 | 0.605 | |
| | | | | WT | 1.667 | 0.664 | | | | |
| | | D12 | Long | KO | 0.972 | 0.539 | | -0.539 | 0.605 | |
| | | | | WT | 0.889 | 0.304 | | | | |
| | | D13 | Long | KO | 0.917 | 0.264 | | -0.133 | 0.931 | |
| | | | | WT | 0.917 | 0.309 | | | | |
| | | D14 | Long | KO | 0.611 | 0.334 | | -1.232 | 0.258 | |
| | | | | WT | 0.806 | 0.235 | | | | |
| | | D15 | Long | KO | 0.861 | 0.328 | | -0.182 | 0.863 | |
| | | | | WT | 0.889 | 0.326 | | | | |
| | | D16 | Long | KO | 1.111 | 0.398 | | -0.315 | 0.796 | |
| | | | | WT | 0.778 | 0.258 | | | | |
| Table S1 | Figure S2A | D0 | Short | KO | 0.111 | 0.033 | Mann-Whitney U Test | -0.094 | 0.931 | |
| | | | | WT | 0.167 | 0.075 | | | | |
| | | D1 | Short | KO | 0.375 | 0.108 | | -1.820 | 0.077 | |
| | | | | WT | 0.125 | 0.042 | | | | |
| | | D2 | Short | KO | 0.403 | 0.125 | | -0.453 | 0.666 | |
| | | | | WT | 0.319 | 0.081 | | | | |
| | | D3 | Short | KO | 0.639 | 0.157 | | -0.134 | 0.931 | |
| | | | | WT | 0.694 | 0.181 | | | | |
| | | D4 | Short | KO | 0.833 | 0.132 | | -0.540 | 0.605 | |
| | | | | WT | 0.750 | 0.156 | | | | |
| | | D5 | Short | KO | 1.167 | 0.220 | | -0.178 | 0.863 | |
| | | | | WT | 1.222 | 0.275 | | | | |
| | | D6 | Short | KO | 1.278 | 0.255 | | -1.251 | 0.222 | |
| | | | | WT | 0.806 | 0.199 | | | | |
| | | D7 | Short | KO | 1.333 | 0.144 | | -0.179 | 0.863 | |
| | | | | WT | 1.278 | 0.206 | | | | |
| | | D8 | Short | KO | 1.944 | 0.327 | | -2.087 | 0.040 | * |
| | | | | WT | 1.111 | 0.151 | | | | |
| | | D9 | Short | KO | 1.444 | 0.249 | | -0.357 | 0.730 | |
| | | | | WT | 1.417 | 0.177 | | | | |
| | | D10 | Short | KO | 1.556 | 0.311 | | -0.400 | 0.730 | |
| | | | | WT | 1.472 | 0.214 | | | | |
| | | D11 | Short | KO | 1.500 | 0.325 | | -0.312 | 0.796 | |
| | | | | WT | 1.306 | 0.242 | | | | |
| | | D12 | Short | KO | 1.333 | 0.298 | | -0.359 | 0.730 | |
| | | | | WT | 1.361 | 0.196 | | | | |
| | | D13 | Short | KO | 1.361 | 0.221 | | -1.449 | 0.161 | |
| | | | | WT | 1.139 | 0.073 | | | | |
| | | D14 | Short | KO | 1.222 | 0.202 | | -0.134 | 0.931 | |
| | | | | WT | 1.194 | 0.220 | | | | |
| | | D15 | Short | KO | 1.028 | 0.193 | | -0.403 | 0.730 | |
| | | | | WT | 1.028 | 0.226 | | | | |
| | | D16 | Short | KO | 1.167 | 0.161 | | -0.765 | 0.489 | |
| | | | | WT | 0.972 | 0.128 | | | | |

**HE frequency in pre-US + US**

| Table | Related Figure | Day | Group | Condition | Mean | SEM | Stastistical test | Z | P | * |
|---|---|---|---|---|---|---|---|---|---|---|
| Table S1 | Figure S2A | D0 | KO | Long | 0.167 | 0.055 | Wilcoxon Signed Ranks Test | -0.738 | 0.461 | |
| | | | | Short | 0.111 | 0.033 | | | | |
| | | D1 | KO | Long | 0.111 | 0.044 | | -2.201 | 0.028 | * |
| | | | | Short | 0.375 | 0.108 | | | | |
| | | D2 | KO | Long | 0.292 | 0.114 | | -1.300 | 0.194 | |
| | | | | Short | 0.403 | 0.125 | | | | |
| | | D3 | KO | Long | 0.333 | 0.125 | | -1.420 | 0.156 | |
| | | | | Short | 0.639 | 0.157 | | | | |
| | | D4 | KO | Long | 0.917 | 0.315 | | -0.344 | 0.731 | |
| | | | | Short | 0.833 | 0.132 | | | | |
| | | D5 | KO | Long | 1.056 | 0.212 | | -0.599 | 0.549 | |
| | | | | Short | 1.167 | 0.220 | | | | |
| | | D6 | KO | Long | 1.333 | 0.351 | | 0.000 | 1.000 | |
| | | | | Short | 1.278 | 0.255 | | | | |
| | | D7 | KO | Long | 1.056 | 0.393 | | -1.129 | 0.259 | |
| | | | | Short | 1.333 | 0.144 | | | | |
| | | D8 | KO | Long | 1.500 | 0.344 | | -1.476 | 0.140 | |
| | | | | Short | 1.944 | 0.327 | | | | |
| | | D9 | KO | Long | 1.583 | 0.460 | | -0.254 | 0.799 | |
| | | | | Long | 1.444 | 0.249 | | | | |
| | | D10 | KO | Short | 0.917 | 0.470 | | -1.402 | 0.161 | |
| | | | | Long | 1.556 | 0.311 | | | | |
| | | D11 | KO | Short | 0.833 | 0.323 | | -1.666 | 0.096 | |
| | | | | Long | 1.500 | 0.325 | | | | |
| | | D12 | KO | Short | 0.972 | 0.539 | | -1.424 | 0.154 | |
| | | | | Long | 1.333 | 0.298 | | | | |
| | | D13 | KO | Short | 0.917 | 0.264 | | -1.371 | 0.170 | |
| | | | | Long | 1.361 | 0.221 | | | | |
| | | D14 | KO | Short | 0.611 | 0.334 | | -1.602 | 0.109 | |
| | | | | Long | 1.222 | 0.202 | | | | |
| | | D15 | KO | Short | 0.861 | 0.328 | | -0.357 | 0.721 | |
| | | | | Long | 1.028 | 0.193 | | | | |
| | | D16 | KO | Short | 1.111 | 0.398 | | 0.000 | 1.000 | |
| | | | | Long | 1.167 | 0.161 | | | | |
| Table S1 | Figure S2A | D0 | WT | Long | 0.292 | 0.247 | Wilcoxon Signed Ranks Test | -0.271 | 0.786 | |
| | | | | Short | 0.167 | 0.075 | | | | |
| | | D1 | WT | Long | 0.097 | 0.040 | | -0.552 | 0.581 | |
| | | | | Short | 0.125 | 0.042 | | | | |
| | | D2 | WT | Long | 0.194 | 0.073 | | -1.289 | 0.197 | |
| | | | | Short | 0.319 | 0.081 | | | | |
| | | D3 | WT | Long | 0.556 | 0.149 | | -1.394 | 0.163 | |
| | | | | Short | 0.694 | 0.181 | | | | |
| | | D4 | WT | Long | 0.833 | 0.144 | | -0.285 | 0.776 | |
| | | | | Short | 0.750 | 0.156 | | | | |
| | | D5 | WT | Long | 1.000 | 0.306 | | -1.205 | 0.228 | |
| | | | | Short | 1.222 | 0.275 | | | | |
| | | D6 | WT | Long | 0.528 | 0.188 | | -0.775 | 0.438 | |
| | | | | Short | 0.806 | 0.199 | | | | |
| | | D7 | WT | Long | 0.694 | 0.246 | | -2.254 | 0.024 | * |
| | | | | Short | 1.278 | 0.206 | | | | |
| | | D8 | WT | Long | 1.056 | 0.311 | | -0.085 | 0.932 | |
| | | | | Short | 1.111 | 0.151 | | | | |
| | | D9 | WT | Long | 1.500 | 0.550 | | .000d | 1.000 | |
| | | | | Short | 1.417 | 0.177 | | | | |
| | | D10 | WT | Short | 1.139 | 0.384 | | -0.655 | 0.512 | |
| | | | | Long | 1.472 | 0.214 | | | | |
| | | D11 | WT | Short | 1.667 | 0.664 | | -0.356 | 0.722 | |
| | | | | Long | 1.306 | 0.242 | | | | |
| | | D12 | WT | Short | 0.889 | 0.304 | | -1.191 | 0.234 | |
| | | | | Long | 1.361 | 0.196 | | | | |
| | | D13 | WT | Short | 0.917 | 0.309 | | -0.775 | 0.438 | |
| | | | | Long | 1.139 | 0.073 | | | | |
| | | D14 | WT | Short | 0.806 | 0.235 | | -1.544 | 0.123 | |
| | | | | Long | 1.194 | 0.220 | | | | |
| | | D15 | WT | Short | 0.889 | 0.326 | | -0.535 | 0.592 | |
| | | | | Long | 1.028 | 0.226 | | | | |
| | | D16 | WT | Short | 0.778 | 0.258 | | -0.846 | 0.397 | |
| | | | | Long | 0.972 | 0.128 | | | | |

## 1_7

HE frequency in pre-CS

| Table | Related Figure | Day | Conditon | Group | Mean | SEM | Stastistical test | Z | P | * |
|---|---|---|---|---|---|---|---|---|---|---|
| Table S1 | Figure S2A | D0 | Long | KO | 0.369 | 0.046 | Mann-Whitney U Test | -2.121 | 0.031 | * |
| | | | | WT | 0.217 | 0.088 | | | | |
| | | D1 | Long | KO | 0.513 | 0.084 | | -2.123 | 0.031 | * |
| | | | | WT | 0.233 | 0.054 | | | | |
| | | D2 | Long | KO | 0.669 | 0.101 | | -2.255 | 0.024 | * |
| | | | | WT | 0.376 | 0.04 | | | | |
| | | D3 | Long | KO | 0.541 | 0.098 | | -0.708 | 0.489 | |
| | | | | WT | 0.637 | 0.08 | | | | |
| | | D4 | Long | KO | 0.737 | 0.101 | | -0.530 | 0.605 | |
| | | | | WT | 0.807 | 0.123 | | | | |
| | | D5 | Long | KO | 0.67 | 0.098 | | -0.886 | 0.387 | |
| | | | | WT | 0.856 | 0.091 | | | | |
| | | D6 | Long | KO | 0.941 | 0.157 | | -1.724 | 0.094 | |
| | | | | WT | 0.585 | 0.096 | | | | |
| | | D7 | Long | KO | 0.778 | 0.165 | | -1.106 | 0.297 | |
| | | | | WT | 0.856 | 0.091 | | | | |
| | | D8 | Long | KO | 0.693 | 0.151 | | -1.370 | 0.190 | |
| | | | | WT | 0.904 | 0.108 | | | | |
| | | D9 | Long | KO | 0.693 | 0.113 | | -0.133 | 0.931 | |
| | | | | WT | 0.622 | 0.087 | | | | |
| | | D10 | Long | KO | 0.437 | 0.067 | | -1.901 | 0.063 | |
| | | | | WT | 0.704 | 0.088 | | | | |
| | | D11 | Long | KO | 0.522 | 0.07 | | -1.510 | 0.136 | |
| | | | | WT | 0.667 | 0.064 | | | | |
| | | D12 | Long | KO | 0.5 | 0.052 | | -1.461 | 0.161 | |
| | | | | WT | 0.707 | 0.114 | | | | |
| | | D13 | Long | KO | 0.478 | 0.104 | | -0.887 | 0.387 | |
| | | | | WT | 0.559 | 0.073 | | | | |
| | | D14 | Long | KO | 0.615 | 0.084 | | -0.133 | 0.931 | |
| | | | | WT | 0.633 | 0.096 | | | | |
| | | D15 | Long | KO | 0.556 | 0.058 | | -1.105 | 0.297 | |
| | | | | WT | 0.652 | 0.088 | | | | |
| | | D16 | Long | KO | 0.556 | 0.078 | | -0.442 | 0.666 | |
| | | | | WT | 0.596 | 0.071 | | | | |
| Table S1 | Figure S2A | D0 | Short | KO | 0.193 | 0.051 | Mann-Whitney U Test | -1.245 | 0.222 | |
| | | | | WT | 0.119 | 0.035 | | | | |
| | | D1 | Short | KO | 0.335 | 0.068 | | -1.336 | 0.190 | |
| | | | | WT | 0.237 | 0.037 | | | | |
| | | D2 | Short | KO | 0.38 | 0.049 | | -2.435 | 0.014 | * |
| | | | | WT | 0.213 | 0.032 | | | | |
| | | D3 | Short | KO | 0.478 | 0.097 | | -0.133 | 0.931 | |
| | | | | WT | 0.515 | 0.1 | | | | |
| | | D4 | Short | KO | 0.596 | 0.077 | | -0.044 | 1.000 | |
| | | | | WT | 0.615 | 0.074 | | | | |
| | | D5 | Short | KO | 0.444 | 0.072 | | -0.620 | 0.546 | |
| | | | | WT | 0.519 | 0.064 | | | | |
| | | D6 | Short | KO | 0.859 | 0.142 | | -1.460 | 0.161 | |
| | | | | WT | 0.541 | 0.098 | | | | |
| | | D7 | Short | KO | 0.648 | 0.086 | | -0.884 | 0.387 | |
| | | | | WT | 0.781 | 0.125 | | | | |
| | | D8 | Short | KO | 0.859 | 0.161 | | -1.150 | 0.258 | |
| | | | | WT | 0.611 | 0.098 | | | | |
| | | D9 | Short | KO | 0.763 | 0.205 | | -0.354 | 0.730 | |
| | | | | WT | 0.781 | 0.159 | | | | |
| | | D10 | Short | KO | 0.53 | 0.067 | | -0.444 | 0.666 | |
| | | | | WT | 0.585 | 0.16 | | | | |
| | | D11 | Short | KO | 0.533 | 0.056 | | -0.265 | 0.796 | |
| | | | | WT | 0.633 | 0.173 | | | | |
| | | D12 | Short | KO | 0.574 | 0.102 | | -0.620 | 0.546 | |
| | | | | WT | 0.456 | 0.084 | | | | |
| | | D13 | Short | KO | 0.526 | 0.113 | | -1.061 | 0.297 | |
| | | | | WT | 0.348 | 0.061 | | | | |
| | | D14 | Short | KO | 0.485 | 0.089 | | -1.336 | 0.190 | |
| | | | | WT | 0.315 | 0.062 | | | | |
| | | D15 | Short | KO | 0.307 | 0.083 | | -0.709 | 0.489 | |
| | | | | WT | 0.396 | 0.085 | | | | |
| | | D16 | Short | KO | 0.407 | 0.056 | | -1.863 | 0.063 | |
| | | | | WT | 0.27 | 0.045 | | | | |

HE frequency in post-CS

| Table | Related Figure | Day | Conditon | Group | Mean | SEM | Stastistical test | Z | P | * |
|---|---|---|---|---|---|---|---|---|---|---|
| Table S1 | Figure S2A | D0 | Long | KO | 0.219 | 0.038 | Mann-Whitney U Test | -1.552 | 0.121 | |
| | | | | WT | 0.167 | 0.066 | | | | |
| | | D1 | Long | KO | 0.283 | 0.052 | | -0.620 | 0.535 | |
| | | | | WT | 0.319 | 0.116 | | | | |
| | | D2 | Long | KO | 0.320 | 0.070 | | -0.753 | 0.451 | |
| | | | | WT | 0.374 | 0.046 | | | | |
| | | D3 | Long | KO | 0.515 | 0.110 | | -0.712 | 0.476 | |
| | | | | WT | 0.648 | 0.097 | | | | |
| | | D4 | Long | KO | 0.352 | 0.082 | | -0.355 | 0.722 | |
| | | | | WT | 0.400 | 0.083 | | | | |
| | | D5 | Long | KO | 0.344 | 0.056 | | -0.089 | 0.929 | |
| | | | | WT | 0.330 | 0.057 | | | | |
| | | D6 | Long | KO | 0.267 | 0.065 | | -0.222 | 0.825 | |
| | | | | WT | 0.393 | 0.118 | | | | |
| | | D7 | Long | KO | 0.285 | 0.081 | | -0.177 | 0.859 | |
| | | | | WT | 0.267 | 0.084 | | | | |
| | | D8 | Long | KO | 0.204 | 0.063 | | -0.045 | 0.964 | |
| | | | | WT | 0.211 | 0.044 | | | | |
| | | D9 | Long | KO | 0.211 | 0.084 | | -0.178 | 0.859 | |
| | | | | WT | 0.204 | 0.064 | | | | |
| | | D10 | Long | KO | 0.263 | 0.124 | | 0.000 | 1.000 | |
| | | | | WT | 0.130 | 0.044 | | | | |
| | | D11 | Long | KO | 0.259 | 0.104 | | -0.672 | 0.502 | |
| | | | | WT | 0.141 | 0.052 | | | | |
| | | D12 | Long | KO | 0.178 | 0.097 | | -0.673 | 0.501 | |
| | | | | WT | 0.126 | 0.047 | | | | |
| | | D13 | Long | KO | 0.189 | 0.086 | | -1.243 | 0.214 | |
| | | | | WT | 0.256 | 0.098 | | | | |
| | | D14 | Long | KO | 0.215 | 0.094 | | -0.755 | 0.450 | |
| | | | | WT | 0.181 | 0.030 | | | | |
| | | D15 | Long | KO | 0.185 | 0.084 | | -1.064 | 0.287 | |
| | | | | WT | 0.230 | 0.034 | | | | |
| | | D16 | Long | KO | 0.178 | 0.086 | | -0.582 | 0.561 | |
| | | | | WT | 0.200 | 0.092 | | | | |
| Table S1 | Figure S2A | D0 | Short | KO | 0.200 | 0.050 | Mann-Whitney U Test | -1.994 | 0.046 | * |
| | | | | WT | 0.100 | 0.030 | | | | |
| | | D1 | Short | KO | 0.352 | 0.083 | | -1.283 | 0.199 | |
| | | | | WT | 0.198 | 0.051 | | | | |
| | | D2 | Short | KO | 0.430 | 0.056 | | -2.087 | 0.037 | * |
| | | | | WT | 0.291 | 0.043 | | | | |
| | | D3 | Short | KO | 0.526 | 0.097 | | -0.577 | 0.564 | |
| | | | | WT | 0.607 | 0.136 | | | | |
| | | D4 | Short | KO | 0.770 | 0.092 | | -0.531 | 0.595 | |
| | | | | WT | 0.741 | 0.100 | | | | |
| | | D5 | Short | KO | 0.552 | 0.057 | | -1.813 | 0.070 | |
| | | | | WT | 0.804 | 0.115 | | | | |
| | | D6 | Short | KO | 0.944 | 0.165 | | -1.769 | 0.077 | |
| | | | | WT | 0.578 | 0.098 | | | | |
| | | D7 | Short | KO | 0.800 | 0.111 | | -0.221 | 0.825 | |
| | | | | WT | 0.793 | 0.121 | | | | |
| | | D8 | Short | KO | 0.911 | 0.170 | | -0.265 | 0.791 | |
| | | | | WT | 0.881 | 0.124 | | | | |
| | | D9 | Short | KO | 1.015 | 0.289 | | -0.398 | 0.690 | |
| | | | | WT | 0.733 | 0.129 | | | | |
| | | D10 | Short | KO | 0.630 | 0.127 | | -0.310 | 0.757 | |
| | | | | WT | 0.670 | 0.127 | | | | |
| | | D11 | Short | KO | 0.596 | 0.128 | | -0.532 | 0.595 | |
| | | | | WT | 0.733 | 0.235 | | | | |
| | | D12 | Short | KO | 0.574 | 0.129 | | -0.133 | 0.894 | |
| | | | | WT | 0.559 | 0.103 | | | | |
| | | D13 | Short | KO | 0.489 | 0.116 | | -0.265 | 0.791 | |
| | | | | WT | 0.404 | 0.067 | | | | |
| | | D14 | Short | KO | 0.496 | 0.072 | | -0.487 | 0.626 | |
| | | | | WT | 0.430 | 0.066 | | | | |
| | | D15 | Short | KO | 0.433 | 0.098 | | -0.266 | 0.790 | |
| | | | | WT | 0.374 | 0.086 | | | | |
| | | D16 | Short | KO | 0.385 | 0.084 | | -0.133 | 0.894 | |
| | | | | WT | 0.400 | 0.067 | | | | |

## 1_8

HE frequency in D0 and D1

| Table | Related Figure | Condition/Group | | Mean | SEM | Stastistical test | Z | P | * |
|---|---|---|---|---|---|---|---|---|---|
| Table S1 | Figure 1E | D0 | KO | 6.361 | 1.354 | Mann-Whitney U Test | -2.471 | 0.013 | * |
| | | | WT | 3.444 | 0.919 | | | | |
| | | D1 | KO | 7.944 | 1.354 | | -2.078 | 0.037 | * |
| | | | WT | 4.250 | 0.646 | | | | |
| | | KO | D0 | 6.361 | 1.052 | Wilcoxon Signed Ranks Test | -1.090 | 0.276 | |
| | | | D1 | 7.944 | 1.354 | | | | |
| | | WT | D0 | 3.444 | 0.919 | | -1.091 | 0.275 | |
| | | | D1 | 4.250 | 0.646 | | | | |

## 2_1

Hit and correct rejection rates (CS)

| Table | Related Figure | Day | Condition | Group | Mean | SEM | Stastistical test | Z | P | * |
|---|---|---|---|---|---|---|---|---|---|---|
| Table S2 | Figure 1F | D0 | Hit | KO | 8.89 | 3.51 | Mann-Whitney U Test | -2.240 | 0.031 | * |
| | | | | WT | 32.22 | 9.83 | | | | |
| | | D1 | Hit | KO | 16.67 | 6.24 | | -0.460 | 0.666 | |
| | | | | WT | 21.11 | 8.24 | | | | |
| | | D2 | Hit | KO | 26.67 | 8.16 | | -0.315 | 0.796 | |
| | | | | WT | 37.78 | 9.97 | | | | |
| | | D3 | Hit | KO | 35.56 | 9.44 | | -1.126 | 0.297 | |
| | | | | WT | 44.44 | 10.56 | | | | |
| | | D4 | Hit | KO | 64.44 | 8.99 | | -0.045 | 1.000 | |
| | | | | WT | 60.00 | 12.91 | | | | |
| | | D5 | Hit | KO | 63.33 | 10.27 | | -0.314 | 0.796 | |
| | | | | WT | 63.33 | 9.86 | | | | |
| | | D6 | Hit | KO | 70.00 | 8.66 | | -1.027 | 0.340 | |
| | | | | WT | 81.11 | 7.90 | | | | |
| | | D7 | Hit | KO | 78.89 | 6.55 | | -0.268 | 0.796 | |
| | | | | WT | 74.44 | 10.29 | | | | |
| | | D8 | Hit | KO | 86.67 | 2.89 | | -1.697 | 0.113 | |
| | | | | WT | 87.78 | 7.95 | | | | |
| | | D9 | Hit | KO | 86.67 | 4.41 | | -1.855 | 0.077 | |
| | | | | WT | 93.33 | 5.53 | | | | |
| | | D10 | Hit | KO | 88.89 | 4.23 | | -0.417 | 0.730 | |
| | | | | WT | 85.56 | 10.42 | | | | |
| | | D11 | Hit | KO | 94.44 | 2.42 | | -0.722 | 0.489 | |
| | | | | WT | 75.56 | 11.07 | | | | |
| | | D12 | Hit | KO | 94.44 | 2.42 | | -0.419 | 0.730 | |
| | | | | WT | 88.89 | 7.72 | | | | |
| | | D13 | Hit | KO | 90.00 | 3.73 | | -0.904 | 0.387 | |
| | | | | WT | 82.22 | 6.62 | | | | |
| | | D14 | Hit | KO | 92.22 | 2.22 | | -0.637 | 0.546 | |
| | | | | WT | 87.78 | 6.41 | | | | |
| | | D15 | Hit | KO | 94.44 | 2.42 | | -0.233 | 0.863 | |
| | | | | WT | 88.89 | 6.96 | | | | |
| | | D16 | Hit | KO | 93.33 | 3.33 | | -0.472 | 0.666 | |
| | | | | WT | 94.44 | 3.77 | | | | |
| Table S2 | Figure 1F | D0 | Correctr ejecton | KO | 88.89 | 3.09 | Mann-Whitney U Test | -1.156 | 0.436 | |
| | | | | WT | 95.56 | 2.42 | | | | |
| | | D1 | Correctr ejecton | KO | 78.89 | 5.64 | | -2.535 | 0.050 | |
| | | | | WT | 97.78 | 1.47 | | | | |
| | | D2 | Correctr ejecton | KO | 73.33 | 6.45 | | -1.594 | 0.161 | |
| | | | | WT | 82.22 | 4.01 | | | | |
| | | D3 | Correctr ejecton | KO | 72.22 | 6.83 | | -0.140 | 0.931 | |
| | | | | WT | 77.78 | 4.34 | | | | |
| | | D4 | Correctr ejecton | KO | 57.78 | 6.83 | | -0.496 | 0.666 | |
| | | | | WT | 65.56 | 3.77 | | | | |
| | | D5 | Correctr ejecton | KO | 55.56 | 8.35 | | -0.331 | 0.796 | |
| | | | | WT | 61.11 | 6.11 | | | | |
| | | D6 | Correctr ejecton | KO | 47.78 | 7.41 | | -1.676 | 0.113 | |
| | | | | WT | 65.56 | 8.84 | | | | |
| | | D7 | Correctr ejecton | KO | 41.11 | 9.20 | | -1.606 | 0.161 | |
| | | | | WT | 55.56 | 7.09 | | | | |
| | | D8 | Correctr ejecton | KO | 37.78 | 7.78 | | -0.225 | 0.863 | |
| | | | | WT | 51.11 | 6.33 | | | | |
| | | D9 | Correctr ejecton | KO | 36.67 | 7.82 | | -2.553 | 0.011 | * |
| | | | | WT | 51.11 | 5.12 | | | | |
| | | D10 | Correctr ejecton | KO | 41.11 | 9.20 | | -1.147 | 0.297 | |
| | | | | WT | 43.33 | 8.16 | | | | |
| | | D11 | Correctr ejecton | KO | 42.22 | 9.09 | | -0.451 | 0.666 | |
| | | | | WT | 46.67 | 7.07 | | | | |
| | | D12 | Correctr ejecton | KO | 36.67 | 10.14 | | -0.814 | 0.436 | |
| | | | | WT | 48.89 | 4.55 | | | | |
| | | D13 | Correctr ejecton | KO | 37.78 | 7.60 | | -1.034 | 0.340 | |
| | | | | WT | 51.11 | 3.09 | | | | |
| | | D14 | Correctr ejecton | KO | 33.33 | 7.99 | | -1.793 | 0.077 | |
| | | | | WT | 61.11 | 4.23 | | | | |
| | | D15 | Correctr ejecton | KO | 43.33 | 10.93 | | -0.989 | 0.340 | |
| | | | | WT | 55.56 | 6.48 | | | | |
| | | D16 | Correctr ejecton | KO | 35.56 | 7.09 | | -0.637 | 0.546 | |
| | | | | WT | 50.00 | 2.89 | | | | |

## 2_2

Hit and correct rejection rates (CS + pre-US)

| Table | Related Figure | Day | Condition | Group | Mean | SEM | Stastistical test | Z | P | * |
|---|---|---|---|---|---|---|---|---|---|---|
| Table S2 | Figure S2B | D0 | Hit | KO | 27.78 | 8.78 | Mann-Whitney U Test | -2.176 | 0.031 | * |
| | | | | WT | 6.67 | 2.36 | | | | |
| | | D1 | Hit | KO | 18.89 | 7.72 | | -0.273 | 0.796 | |
| | | | | WT | 13.33 | 5.27 | | | | |
| | | D2 | Hit | KO | 32.22 | 9.83 | | -0.762 | 0.489 | |
| | | | | WT | 24.44 | 8.52 | | | | |
| | | D3 | Hit | KO | 38.89 | 10.06 | | -0.584 | 0.605 | |
| | | | | WT | 24.44 | 7.09 | | | | |
| | | D4 | Hit | KO | 56.67 | 12.02 | | -0.179 | 0.863 | |
| | | | | WT | 56.67 | 10.27 | | | | |
| | | D5 | Hit | KO | 50.00 | 9.13 | | -0.090 | 0.931 | |
| | | | | WT | 54.44 | 11.32 | | | | |
| | | D6 | Hit | KO | 75.56 | 7.66 | | -0.860 | 0.436 | |
| | | | | WT | 62.22 | 9.09 | | | | |
| | | D7 | Hit | KO | 68.89 | 11.48 | | -0.091 | 0.931 | |
| | | | | WT | 71.11 | 6.33 | | | | |
| | | D8 | Hit | KO | 87.78 | 7.95 | | -1.532 | 0.161 | |
| | | | | WT | 81.11 | 4.84 | | | | |
| | | D9 | Hit | KO | 87.78 | 8.62 | | -1.656 | 0.136 | |
| | | | | WT | 78.89 | 6.33 | | | | |
| | | D10 | Hit | KO | 80.00 | 10.41 | | -1.220 | 0.297 | |
| | | | | WT | 83.33 | 5.77 | | | | |
| | | D11 | Hit | KO | 70.00 | 11.55 | | -1.139 | 0.297 | |
| | | | | WT | 84.44 | 5.56 | | | | |
| | | D12 | Hit | KO | 83.33 | 9.13 | | -0.152 | 0.931 | |
| | | | | WT | 86.67 | 4.71 | | | | |
| | | D13 | Hit | KO | 77.78 | 7.03 | | -0.781 | 0.489 | |
| | | | | WT | 86.67 | 4.71 | | | | |
| | | D14 | Hit | KO | 83.33 | 7.64 | | -0.048 | 1.000 | |
| | | | | WT | 84.44 | 3.77 | | | | |
| | | D15 | Hit | KO | 82.22 | 9.25 | | -0.101 | 0.931 | |
| | | | | WT | 87.78 | 4.01 | | | | |
| | | D16 | Hit | KO | 87.78 | 4.94 | | -0.686 | 0.605 | |
| | | | | WT | 92.22 | 3.24 | | | | |
| Table S2 | Figure S2B | D0 | Correctr ejecton | KO | 95.56 | 2.42 | Mann-Whitney U Test | -1.719 | 0.113 | |
| | | | | WT | 98.89 | 1.11 | | | | |
| | | D1 | Correctr ejecton | KO | 93.33 | 2.36 | | -2.616 | 0.014 | * |
| | | | | WT | 100.00 | 0.00 | | | | |
| | | D2 | Correctr ejecton | KO | 88.89 | 2.61 | | -1.134 | 0.297 | |
| | | | | WT | 94.44 | 1.76 | | | | |
| | | D3 | Correctr ejecton | KO | 88.89 | 3.51 | | -0.762 | 0.489 | |
| | | | | WT | 88.89 | 2.61 | | | | |
| | | D4 | Correctr ejecton | KO | 78.89 | 6.11 | | -1.269 | 0.222 | |
| | | | | WT | 77.78 | 4.34 | | | | |
| | | D5 | Correctr ejecton | KO | 84.44 | 3.38 | | -0.670 | 0.546 | |
| | | | | WT | 83.33 | 2.36 | | | | |
| | | D6 | Correctr ejecton | KO | 81.11 | 4.23 | | -1.604 | 0.113 | |
| | | | | WT | 90.00 | 4.08 | | | | |
| | | D7 | Correctr ejecton | KO | 70.00 | 7.07 | | -0.936 | 0.387 | |
| | | | | WT | 86.67 | 1.67 | | | | |
| | | D8 | Correctr ejecton | KO | 77.78 | 5.47 | | -1.561 | 0.136 | |
| | | | | WT | 78.89 | 4.55 | | | | |
| | | D9 | Correctr ejecton | KO | 71.11 | 4.23 | | -1.914 | 0.063 | |
| | | | | WT | 86.67 | 2.89 | | | | |
| | | D10 | Correctr ejecton | KO | 73.33 | 5.00 | | -0.536 | 0.605 | |
| | | | | WT | 82.22 | 4.94 | | | | |
| | | D11 | Correctr ejecton | KO | 76.67 | 7.64 | | -0.534 | 0.605 | |
| | | | | WT | 77.78 | 3.64 | | | | |
| | | D12 | Correctr ejecton | KO | 71.11 | 6.55 | | -1.123 | 0.297 | |
| | | | | WT | 78.89 | 4.84 | | | | |
| | | D13 | Correctr ejecton | KO | 73.33 | 8.16 | | -1.350 | 0.190 | |
| | | | | WT | 85.56 | 2.94 | | | | |
| | | D14 | Correctr ejecton | KO | 63.33 | 7.99 | | -2.377 | 0.019 | * |
| | | | | WT | 81.11 | 4.55 | | | | |
| | | D15 | Correctr ejecton | KO | 67.78 | 8.46 | | -1.249 | 0.222 | |
| | | | | WT | 81.11 | 4.84 | | | | |
| | | D16 | Correctr ejecton | KO | 70.00 | 7.26 | | -2.163 | 0.031 | * |
| | | | | WT | 77.78 | 3.64 | | | | |

## 2_3

Hi ratet vs. Correct rejection rate (CS)

| Table | Related Figure | Day | Group | Condition | Mean | SEM | Stastistical test | Z | P | * |
|---|---|---|---|---|---|---|---|---|---|---|
| Table S3 | Figure S2C | D0 | KO | Long | 6.67 | 2.36 | Wilcoxon Signed Ranks Test | -2.682 | 0.007 | ** |
| | | | | Short | 98.89 | 1.11 | | | | |
| | | D1 | KO | Long | 13.33 | 5.27 | | -2.673 | 0.008 | ** |
| | | | | Short | 100.00 | 0.00 | | | | |
| | | D2 | KO | Long | 24.44 | 8.52 | | -2.673 | 0.008 | ** |
| | | | | Short | 94.44 | 1.76 | | | | |
| | | D3 | KO | Long | 24.44 | 7.09 | | -2.492 | 0.013 | * |
| | | | | Short | 88.89 | 2.61 | | | | |
| | | D4 | KO | Long | 56.67 | 10.27 | | -1.266 | 0.205 | |
| | | | | Short | 77.78 | 4.34 | | | | |
| | | D5 | KO | Long | 54.44 | 11.32 | | -2.320 | 0.020 | * |
| | | | | Short | 83.33 | 2.36 | | | | |
| | | D6 | KO | Long | 62.22 | 9.09 | | -0.060 | 0.952 | |
| | | | | Short | 90.00 | 4.08 | | | | |
| | | D7 | KO | Long | 71.11 | 6.33 | | -0.281 | 0.779 | |
| | | | | Short | 86.67 | 1.67 | | | | |
| | | D8 | KO | Long | 81.11 | 4.84 | | -1.194 | 0.233 | |
| | | | | Short | 78.89 | 4.55 | | | | |
| | | D9 | KO | Long | 78.89 | 6.33 | | -1.608 | 0.108 | |
| | | | | Long | 86.67 | 2.89 | | | | |
| | | D10 | KO | Short | 83.33 | 5.77 | | -0.718 | 0.473 | |
| | | | | Long | 82.22 | 4.94 | | | | |
| | | D11 | KO | Short | 84.44 | 5.56 | | -0.280 | 0.779 | |
| | | | | Long | 77.78 | 3.64 | | | | |
| | | D12 | KO | Short | 86.67 | 4.71 | | -1.130 | 0.258 | |
| | | | | Long | 78.89 | 4.84 | | | | |
| | | D13 | KO | Short | 86.67 | 4.71 | | -0.417 | 0.677 | |
| | | | | Long | 85.56 | 2.94 | | | | |
| | | D14 | KO | Short | 84.44 | 3.77 | | -1.196 | 0.232 | |
| | | | | Long | 81.11 | 4.55 | | | | |
| | | D15 | KO | Short | 87.78 | 4.01 | | -0.893 | 0.372 | |
| | | | | Long | 81.11 | 4.84 | | | | |
| | | D16 | KO | Short | 92.22 | 3.24 | | -1.411 | 0.158 | |
| | | | | Long | 77.78 | 3.64 | | | | |
| Table S3 | Figure S2C | D0 | WT | Long | 27.78 | 8.78 | Wilcoxon Signed Ranks Test | -2.701 | 0.007 | ** |
| | | | | Short | 95.56 | 2.42 | | | | |
| | | D1 | WT | Long | 18.89 | 7.72 | | -2.701 | 0.007 | ** |
| | | | | Short | 93.33 | 2.36 | | | | |
| | | D2 | WT | Long | 32.22 | 9.83 | | -2.694 | 0.007 | ** |
| | | | | Short | 88.89 | 2.61 | | | | |
| | | D3 | WT | Long | 38.89 | 10.06 | | -2.670 | 0.008 | ** |
| | | | | Short | 88.89 | 3.51 | | | | |
| | | D4 | WT | Long | 56.67 | 12.02 | | -1.436 | 0.151 | |
| | | | | Short | 78.89 | 6.11 | | | | |
| | | D5 | WT | Long | 50.00 | 9.13 | | -1.796 | 0.072 | |
| | | | | Short | 84.44 | 3.38 | | | | |
| | | D6 | WT | Long | 75.56 | 7.66 | | -2.066 | 0.039 | * |
| | | | | Short | 81.11 | 4.23 | | | | |
| | | D7 | WT | Long | 68.89 | 11.48 | | -1.904 | 0.057 | |
| | | | | Short | 70.00 | 7.07 | | | | |
| | | D8 | WT | Long | 87.78 | 7.95 | | -0.541 | 0.589 | |
| | | | | Short | 77.78 | 5.47 | | | | |
| | | D9 | WT | Long | 87.78 | 8.62 | | -0.954 | 0.340 | |
| | | | | Long | 71.11 | 4.23 | | | | |
| | | D10 | WT | Short | 80.00 | 10.41 | | -0.179 | 0.858 | |
| | | | | Long | 73.33 | 5.00 | | | | |
| | | D11 | WT | Short | 70.00 | 11.55 | | -1.029 | 0.303 | |
| | | | | Long | 76.67 | 7.64 | | | | |
| | | D12 | WT | Short | 83.33 | 9.13 | | -0.846 | 0.397 | |
| | | | | Long | 71.11 | 6.55 | | | | |
| | | D13 | WT | Short | 77.78 | 7.03 | | -0.085 | 0.932 | |
| | | | | Long | 73.33 | 8.16 | | | | |
| | | D14 | WT | Short | 83.33 | 7.64 | | -0.426 | 0.670 | |
| | | | | Long | 63.33 | 7.99 | | | | |
| | | D15 | WT | Short | 82.22 | 9.25 | | -0.877 | 0.380 | |
| | | | | Long | 67.78 | 8.46 | | | | |
| | | D16 | WT | Short | 87.78 | 4.94 | | -2.200 | 0.028 | * |
| | | | | Long | 70.00 | 7.26 | | | | |

Abbreviations: D, training day; KO, CRBN KO mice; WT, wild-type mice.
*, *p < 0.05, **p < 0.01, ***p < 0.001.

3_1

| HE peak frequency | | | | | | | | | | |
|---|---|---|---|---|---|---|---|---|---|---|
| Table | Related Figure | Day | Condition | Group | Mean | SEM | Stastistical test | Z | P | * |
| Table S3 | Figure 1G | D0 | Long | KO | 1.220 | 0.220 | Mann-Whitney U Test | -1.748 | 0.113 | |
| | | | | WT | 0.780 | 0.430 | | | | |
| | | D1 | Long | KO | 0.780 | 0.220 | | -0.458 | 0.730 | |
| | | | | WT | 1.000 | 0.290 | | | | |
| | | D2 | Long | KO | 1.220 | 0.280 | | -0.549 | 0.605 | |
| | | | | WT | 1.670 | 0.470 | | | | |
| | | D3 | Long | KO | 2.220 | 0.490 | | -0.276 | 0.796 | |
| | | | | WT | 2.110 | 0.350 | | | | |
| | | D4 | Long | KO | 2.780 | 0.620 | | -0.545 | 0.605 | |
| | | | | WT | 2.440 | 0.290 | | | | |
| | | D5 | Long | KO | 2.890 | 0.420 | | -0.182 | 0.863 | |
| | | | | WT | 3.000 | 0.530 | | | | |
| | | D6 | Long | KO | 4.110 | 0.420 | | -1.878 | 0.077 | |
| | | | | WT | 3.220 | 0.280 | | | | |
| | | D7 | Long | KO | 3.890 | 0.680 | | -0.586 | 0.605 | |
| | | | | WT | 3.330 | 0.410 | | | | |
| | | D8 | Long | KO | 5.220 | 0.680 | | -2.312 | 0.024 | * |
| | | | | WT | 3.560 | 0.340 | | | | |
| | | D9 | Long | KO | 5.220 | 0.600 | | -1.162 | 0.258 | |
| | | | | WT | 4.220 | 0.660 | | | | |
| | | D10 | Long | KO | 5.110 | 0.610 | | -1.691 | 0.113 | |
| | | | | WT | 3.890 | 0.260 | | | | |
| | | D11 | Long | KO | 5.110 | 0.960 | | -0.179 | 0.863 | |
| | | | | WT | 4.440 | 0.560 | | | | |
| | | D12 | Long | KO | 4.440 | 0.650 | | -0.091 | 0.931 | |
| | | | | WT | 4.670 | 0.580 | | | | |
| | | D13 | Long | KO | 5.330 | 0.670 | | -1.305 | 0.222 | |
| | | | | WT | 4.330 | 0.370 | | | | |
| | | D14 | Long | KO | 4.220 | 0.400 | | -0.091 | 0.931 | |
| | | | | WT | 4.330 | 0.410 | | | | |
| | | D15 | Long | KO | 4.670 | 0.470 | | -0.273 | 0.796 | |
| | | | | WT | 4.440 | 0.380 | | | | |
| | | D16 | Long | KO | 4.670 | 0.470 | | -0.872 | 0.436 | |
| | | | | WT | 5.000 | 0.410 | | | | |
| Table S3 | Figure 1G | D0 | Short | KO | 1.110 | 0.200 | Mann-Whitney U Test | -0.191 | 0.863 | |
| | | | | WT | 1.110 | 0.350 | | | | |
| | | D1 | Short | KO | 1.890 | 0.590 | | -1.189 | 0.297 | |
| | | | | WT | 1.000 | 0.240 | | | | |
| | | D2 | Short | KO | 2.110 | 0.310 | | -2.100 | 0.050 | |
| | | | | WT | 1.220 | 0.220 | | | | |
| | | D3 | Short | KO | 2.560 | 0.440 | | -0.047 | 1.000 | |
| | | | | WT | 2.670 | 0.500 | | | | |
| | | D4 | Short | KO | 2.890 | 0.590 | | -0.182 | 0.863 | |
| | | | | WT | 2.890 | 0.560 | | | | |
| | | D5 | Short | KO | 2.670 | 0.240 | | -0.599 | 0.605 | |
| | | | | WT | 3.220 | 0.550 | | | | |
| | | D6 | Short | KO | 3.440 | 0.530 | | -1.582 | 0.136 | |
| | | | | WT | 2.330 | 0.370 | | | | |
| | | D7 | Short | KO | 3.560 | 0.440 | | -0.731 | 0.489 | |
| | | | | WT | 3.110 | 0.350 | | | | |
| | | D8 | Short | KO | 4.670 | 0.620 | | -1.906 | 0.063 | |
| | | | | WT | 3.000 | 0.290 | | | | |
| | | D9 | Short | KO | 3.670 | 0.620 | | -0.275 | 0.796 | |
| | | | | WT | 3.440 | 0.500 | | | | |
| | | D10 | Short | KO | 3.220 | 0.280 | | -0.095 | 0.931 | |
| | | | | WT | 3.330 | 0.290 | | | | |
| | | D11 | Short | KO | 3.780 | 0.360 | | -0.911 | 0.387 | |
| | | | | WT | 3.330 | 0.470 | | | | |
| | | D12 | Short | KO | 3.780 | 0.550 | | -0.136 | 0.931 | |
| | | | | WT | 3.780 | 0.360 | | | | |
| | | D13 | Short | KO | 3.560 | 0.470 | | -1.780 | 0.094 | |
| | | | | WT | 2.440 | 0.180 | | | | |
| | | D14 | Short | KO | 4.220 | 0.550 | | -1.499 | 0.161 | |
| | | | | WT | 3.110 | 0.260 | | | | |
| | | D15 | Short | KO | 3.440 | 0.650 | | -0.592 | 0.605 | |
| | | | | WT | 3.440 | 0.440 | | | | |
| | | D16 | Short | KO | 3.440 | 0.560 | | -0.651 | 0.546 | |
| | | | | WT | 2.890 | 0.200 | | | | |

| HE peak frequency | | | | | | | | | | |
|---|---|---|---|---|---|---|---|---|---|---|
| Table | Related Figure | Day | Group | Condition | Mean | SEM | Stastistical test | Z | P | * |
| Table S3 | Figure 1G | D0 | KO | Long | 1.220 | 0.220 | Wilcoxon Signed Ranks Test | -0.447 | 0.655 | |
| | | | | Short | 1.110 | 0.200 | | | | |
| | | D1 | KO | Long | 0.780 | 0.220 | | -1.633 | 0.102 | |
| | | | | Short | 1.890 | 0.590 | | | | |
| | | D2 | KO | Long | 1.220 | 0.280 | | -2.271 | 0.023 | * |
| | | | | Short | 2.110 | 0.310 | | | | |
| | | D3 | KO | Long | 2.220 | 0.490 | | -0.707 | 0.480 | |
| | | | | Short | 2.560 | 0.440 | | | | |
| | | D4 | KO | Long | 2.780 | 0.620 | | -0.256 | 0.798 | |
| | | | | Short | 2.890 | 0.590 | | | | |
| | | D5 | KO | Long | 2.890 | 0.420 | | -0.649 | 0.516 | |
| | | | | Short | 2.670 | 0.240 | | | | |
| | | D6 | KO | Long | 4.110 | 0.420 | | -1.268 | 0.205 | |
| | | | | Short | 3.440 | 0.530 | | | | |
| | | D7 | KO | Long | 3.890 | 0.680 | | -1.000 | 0.317 | |
| | | | | Short | 3.560 | 0.440 | | | | |
| | | D8 | KO | Long | 5.220 | 0.680 | | -1.098 | 0.272 | |
| | | | | Short | 4.670 | 0.620 | | | | |
| | | D9 | KO | Long | 5.220 | 0.600 | | -2.354 | 0.019 | * |
| | | | | Long | 3.670 | 0.620 | | | | |
| | | D10 | KO | Short | 5.110 | 0.610 | | -2.379 | 0.017 | * |
| | | | | Long | 3.220 | 0.280 | | | | |
| | | D11 | KO | Short | 5.110 | 0.960 | | -1.538 | 0.124 | |
| | | | | Long | 3.780 | 0.360 | | | | |
| | | D12 | KO | Short | 4.440 | 0.650 | | -1.137 | 0.256 | |
| | | | | Long | 3.780 | 0.550 | | | | |
| | | D13 | KO | Short | 5.330 | 0.670 | | -2.129 | 0.033 | * |
| | | | | Long | 3.560 | 0.470 | | | | |
| | | D14 | KO | Short | 4.220 | 0.400 | | -0.333 | 0.739 | |
| | | | | Long | 4.220 | 0.550 | | | | |
| | | D15 | KO | Short | 4.670 | 0.470 | | -1.430 | 0.153 | |
| | | | | Long | 3.440 | 0.650 | | | | |
| | | D16 | KO | Short | 4.670 | 0.470 | | -1.612 | 0.107 | |
| | | | | Long | 3.440 | 0.560 | | | | |
| Table S3 | Figure 1G | D0 | WT | Long | 0.780 | 0.430 | Wilcoxon Signed Ranks Test | -0.750 | 0.453 | |
| | | | | Short | 1.110 | 0.350 | | | | |
| | | D1 | WT | Long | 1.000 | 0.290 | | 0.000 | 1.000 | |
| | | | | Short | 1.000 | 0.240 | | | | |
| | | D2 | WT | Long | 1.670 | 0.470 | | -1.300 | 0.194 | |
| | | | | Short | 1.220 | 0.220 | | | | |
| | | D3 | WT | Long | 2.110 | 0.350 | | -1.414 | 0.157 | |
| | | | | Short | 2.670 | 0.500 | | | | |
| | | D4 | WT | Long | 2.440 | 0.290 | | -0.707 | 0.480 | |
| | | | | Short | 2.890 | 0.560 | | | | |
| | | D5 | WT | Long | 3.000 | 0.530 | | -0.531 | 0.595 | |
| | | | | Short | 3.220 | 0.550 | | | | |
| | | D6 | WT | Long | 3.220 | 0.280 | | -1.480 | 0.139 | |
| | | | | Short | 2.330 | 0.370 | | | | |
| | | D7 | WT | Long | 3.330 | 0.410 | | -0.702 | 0.483 | |
| | | | | Short | 3.110 | 0.350 | | | | |
| | | D8 | WT | Long | 3.560 | 0.340 | | -1.406 | 0.160 | |
| | | | | Short | 3.000 | 0.290 | | | | |
| | | D9 | WT | Long | 4.220 | 0.660 | | -1.311 | 0.190 | |
| | | | | Long | 3.440 | 0.500 | | | | |
| | | D10 | WT | Short | 3.890 | 0.260 | | -1.155 | 0.248 | |
| | | | | Long | 3.330 | 0.290 | | | | |
| | | D11 | WT | Short | 4.440 | 0.560 | | -1.638 | 0.101 | |
| | | | | Long | 3.330 | 0.470 | | | | |
| | | D12 | WT | Short | 4.670 | 0.580 | | -1.121 | 0.262 | |
| | | | | Long | 3.780 | 0.360 | | | | |
| | | D13 | WT | Short | 4.330 | 0.370 | | -2.555 | 0.011 | * |
| | | | | Long | 2.440 | 0.180 | | | | |
| | | D14 | WT | Short | 4.330 | 0.410 | | -2.050 | 0.040 | * |
| | | | | Long | 3.110 | 0.260 | | | | |
| | | D15 | WT | Short | 4.440 | 0.380 | | -2.124 | 0.034 | * |
| | | | | Long | 3.440 | 0.440 | | | | |
| | | D16 | WT | Short | 5.000 | 0.410 | | -2.539 | 0.011 | * |
| | | | | Long | 2.890 | 0.200 | | | | |

Abbreviations: HE, head entry; FHE, fastest HE; D, training day; KO, CRBN KO mice; WT, wild-type mice.

*, *p < 0.05, **p < 0.01, ***p < 0.001.

3_2

| HE peak latency | | | | | | | | | | |
|---|---|---|---|---|---|---|---|---|---|---|
| Table | Related Figure | Day | Condition | Group | Mean | SEM | Stastistical test | Z | P | * |
| Table S3 | Figure 1G | D0 | Long | KO | 4.000 | 1.093 | Mann-Whitney U Test | -2.037 | 0.050 | |
| | | | | WT | 1.778 | 1.222 | | | | |
| | | D1 | Long | KO | 1.000 | 0.408 | | -0.252 | 0.863 | |
| | | | | WT | 2.000 | 1.443 | | | | |
| | | D2 | Long | KO | 4.222 | 1.310 | | -0.911 | 0.387 | |
| | | | | WT | 3.111 | 1.438 | | | | |
| | | D3 | Long | KO | 3.667 | 1.291 | | -0.542 | 0.605 | |
| | | | | WT | 5.111 | 1.728 | | | | |
| | | D4 | Long | KO | 2.889 | 1.296 | | -0.740 | 0.489 | |
| | | | | WT | 2.111 | 1.111 | | | | |
| | | D5 | Long | KO | 4.667 | 1.202 | | -0.267 | 0.796 | |
| | | | | WT | 5.444 | 1.608 | | | | |
| | | D6 | Long | KO | 4.222 | 1.164 | | -0.311 | 0.796 | |
| | | | | WT | 5.111 | 1.476 | | | | |
| | | D7 | Long | KO | 3.556 | 1.215 | | -1.211 | 0.258 | |
| | | | | WT | 4.222 | 0.572 | | | | |
| | | D8 | Long | KO | 3.222 | 1.382 | | -0.046 | 1.000 | |
| | | | | WT | 3.222 | 1.278 | | | | |
| | | D9 | Long | KO | 5.000 | 1.333 | | -0.489 | 0.666 | |
| | | | | WT | 4.000 | 1.179 | | | | |
| | | D10 | Long | KO | 5.111 | 1.172 | | -1.203 | 0.258 | |
| | | | | WT | 3.111 | 1.060 | | | | |
| | | D11 | Long | KO | 5.556 | 1.600 | | -0.045 | 1.000 | |
| | | | | WT | 5.556 | 1.365 | | | | |
| | | D12 | Long | KO | 3.111 | 1.252 | | -0.271 | 0.796 | |
| | | | | WT | 3.778 | 1.245 | | | | |
| | | D13 | Long | KO | 5.111 | 1.348 | | -1.069 | 0.297 | |
| | | | | WT | 2.889 | 0.633 | | | | |
| | | D14 | Long | KO | 3.889 | 0.889 | | -0.446 | 0.666 | |
| | | | | WT | 4.333 | 0.833 | | | | |
| | | D15 | Long | KO | 5.222 | 1.579 | | -0.403 | 0.730 | |
| | | | | WT | 4.333 | 0.667 | | | | |
| | | D16 | Long | KO | 5.000 | 1.414 | | -0.488 | 0.666 | |
| | | | | WT | 5.778 | 1.103 | | | | |
| Table S3 | Figure 1G | D0 | Short | KO | 3.667 | 1.312 | Mann-Whitney U Test | 0.000 | 1.000 | |
| | | | | WT | 4.000 | 1.616 | | | | |
| | | D1 | Short | KO | 3.333 | 1.312 | | -0.456 | 0.666 | |
| | | | | WT | 4.333 | 1.258 | | | | |
| | | D2 | Short | KO | 6.222 | 1.289 | | -2.051 | 0.050 | |
| | | | | WT | 1.889 | 1.020 | | | | |
| | | D3 | Short | KO | 5.667 | 1.607 | | -1.081 | 0.297 | |
| | | | | WT | 3.556 | 1.692 | | | | |
| | | D4 | Short | KO | 3.444 | 1.271 | | -0.498 | 0.666 | |
| | | | | WT | 2.333 | 1.202 | | | | |
| | | D5 | Short | KO | 2.667 | 0.645 | | -0.045 | 1.000 | |
| | | | | WT | 2.667 | 0.816 | | | | |
| | | D6 | Short | KO | 2.778 | 0.741 | | -1.161 | 0.297 | |
| | | | | WT | 1.778 | 0.969 | | | | |
| | | D7 | Short | KO | 2.778 | 1.392 | | -0.273 | 0.796 | |
| | | | | WT | 4.111 | 1.567 | | | | |
| | | D8 | Short | KO | 3.333 | 1.067 | | -0.357 | 0.730 | |
| | | | | WT | 4.444 | 1.271 | | | | |
| | | D9 | Short | KO | 2.556 | 0.801 | | -1.832 | 0.077 | |
| | | | | WT | 3.889 | 0.484 | | | | |
| | | D10 | Short | KO | 2.222 | 0.683 | | -0.555 | 0.605 | |
| | | | | WT | 4.000 | 1.404 | | | | |
| | | D11 | Short | KO | 4.000 | 1.190 | | 0.000 | 1.000 | |
| | | | | WT | 3.889 | 1.047 | | | | |
| | | D12 | Short | KO | 2.111 | 0.772 | | -1.124 | 0.297 | |
| | | | | WT | 3.222 | 0.862 | | | | |
| | | D13 | Short | KO | 2.556 | 0.899 | | -0.270 | 0.796 | |
| | | | | WT | 2.222 | 0.521 | | | | |
| | | D14 | Short | KO | 1.667 | 0.408 | | -1.399 | 0.190 | |
| | | | | WT | 3.778 | 1.064 | | | | |
| | | D15 | Short | KO | 1.333 | 0.577 | | -2.152 | 0.031 | * |
| | | | | WT | 3.333 | 0.726 | | | | |
| | | D16 | Short | KO | 1.444 | 0.377 | | -0.907 | 0.387 | |
| | | | | WT | 2.000 | 0.471 | | | | |

| HE peak latency | | | | | | | | | | |
|---|---|---|---|---|---|---|---|---|---|---|
| Table | Related Figure | Day | Group | Condition | Mean | SEM | Stastistical test | Z | P | * |
| Table S3 | Figure 1G | D0 | KO | Long | 4.000 | 1.093 | Wilcoxon Signed Ranks Test | -0.140 | 0.889 | |
| | | | | Short | 3.667 | 1.312 | | | | |
| | | D1 | KO | Long | 1.000 | 0.408 | | -1.706 | 0.088 | |
| | | | | Short | 3.333 | 1.312 | | | | |
| | | D2 | KO | Long | 4.222 | 1.310 | | -1.067 | 0.286 | |
| | | | | Short | 6.222 | 1.289 | | | | |
| | | D3 | KO | Long | 3.667 | 1.291 | | -0.844 | 0.398 | |
| | | | | Short | 5.667 | 1.607 | | | | |
| | | D4 | KO | Long | 2.889 | 1.296 | | -0.170 | 0.865 | |
| | | | | Short | 3.444 | 1.271 | | | | |
| | | D5 | KO | Long | 4.667 | 1.202 | | -1.077 | 0.282 | |
| | | | | Short | 2.667 | 0.645 | | | | |
| | | D6 | KO | Long | 4.222 | 1.164 | | -0.845 | 0.398 | |
| | | | | Short | 2.778 | 0.741 | | | | |
| | | D7 | KO | Long | 3.556 | 1.215 | | -0.702 | 0.483 | |
| | | | | Short | 2.778 | 1.392 | | | | |
| | | D8 | KO | Long | 3.222 | 1.382 | | -0.354 | 0.723 | |
| | | | | Short | 3.333 | 1.067 | | | | |
| | | D9 | KO | Long | 5.000 | 1.333 | | -2.252 | 0.024 | * |
| | | | | Long | 2.556 | 0.801 | | | | |
| | | D10 | KO | Short | 5.111 | 1.172 | | -2.184 | 0.029 | * |
| | | | | Long | 2.222 | 0.683 | | | | |
| | | D11 | KO | Short | 5.556 | 1.600 | | -1.268 | 0.205 | |
| | | | | Long | 4.000 | 1.190 | | | | |
| | | D12 | KO | Short | 3.111 | 1.252 | | -1.051 | 0.293 | |
| | | | | Long | 2.111 | 0.772 | | | | |
| | | D13 | KO | Short | 5.111 | 1.348 | | -2.393 | 0.017 | * |
| | | | | Long | 2.556 | 0.899 | | | | |
| | | D14 | KO | Short | 3.889 | 0.889 | | -2.113 | 0.035 | * |
| | | | | Long | 1.667 | 0.408 | | | | |
| | | D15 | KO | Short | 5.222 | 1.579 | | -2.371 | 0.018 | * |
| | | | | Long | 1.333 | 0.577 | | | | |
| | | D16 | KO | Short | 5.000 | 1.414 | | -2.117 | 0.034 | * |
| | | | | Long | 1.444 | 0.377 | | | | |
| Table S3 | Figure 1G | D0 | WT | Long | 1.778 | 1.222 | Wilcoxon Signed Ranks Test | -1.153 | 0.249 | |
| | | | | Short | 4.000 | 1.616 | | | | |
| | | D1 | WT | Long | 2.000 | 1.443 | | -1.781 | 0.075 | |
| | | | | Short | 4.333 | 1.258 | | | | |
| | | D2 | WT | Long | 3.111 | 1.438 | | -0.315 | 0.752 | |
| | | | | Short | 1.889 | 1.020 | | | | |
| | | D3 | WT | Long | 5.111 | 1.728 | | -0.845 | 0.398 | |
| | | | | Short | 3.556 | 1.692 | | | | |
| | | D4 | WT | Long | 2.111 | 1.111 | | 0.000 | 1.000 | |
| | | | | Short | 2.333 | 1.202 | | | | |
| | | D5 | WT | Long | 5.444 | 1.608 | | -1.187 | 0.235 | |
| | | | | Short | 2.667 | 0.816 | | | | |
| | | D6 | WT | Long | 5.111 | 1.476 | | -1.886 | 0.059 | |
| | | | | Short | 1.778 | 0.969 | | | | |
| | | D7 | WT | Long | 4.222 | 0.572 | | -0.070 | 0.944 | |
| | | | | Short | 4.111 | 1.567 | | | | |
| | | D8 | WT | Long | 3.222 | 1.278 | | -0.943 | 0.345 | |
| | | | | Short | 4.444 | 1.271 | | | | |
| | | D9 | WT | Long | 4.000 | 1.179 | | -0.060 | 0.952 | |
| | | | | Long | 3.889 | 0.484 | | | | |
| | | D10 | WT | Short | 3.111 | 1.060 | | -0.211 | 0.833 | |
| | | | | Long | 4.000 | 1.404 | | | | |
| | | D11 | WT | Short | 5.556 | 1.365 | | -0.953 | 0.341 | |
| | | | | Long | 3.889 | 1.047 | | | | |
| | | D12 | WT | Short | 3.778 | 1.245 | | -0.512 | 0.609 | |
| | | | | Long | 3.222 | 0.862 | | | | |
| | | D13 | WT | Short | 2.889 | 0.633 | | -0.991 | 0.322 | |
| | | | | Long | 2.222 | 0.521 | | | | |
| | | D14 | WT | Short | 4.333 | 0.833 | | -0.418 | 0.676 | |
| | | | | Long | 3.778 | 1.064 | | | | |
| | | D15 | WT | Short | 4.333 | 0.667 | | -1.549 | 0.121 | |
| | | | | Long | 3.333 | 0.726 | | | | |
| | | D16 | WT | Short | 5.778 | 1.103 | | -2.524 | 0.012 | * |
| | | | | Long | 2.000 | 0.471 | | | | |

3_3

| Table | Related Figure | Day | Condition | Group | Mean | SEM | Stastistical test | Z | P | * |
|---|---|---|---|---|---|---|---|---|---|---|
| | | | | | FHE peak frequency | | | | | |
| Table S3 | Figure S2D | D0 | Long | KO | 1.444 | 0.338 | Mann-Whitney U Test | -1.720 | 0.113 | |
| | | | | WT | 0.667 | 0.236 | | | | |
| | | D1 | Long | KO | 0.778 | 0.222 | | 0.000 | 1.000 | |
| | | | | WT | 0.778 | 0.222 | | | | |
| | | D2 | Long | KO | 1.222 | 0.278 | | -0.046 | 1.000 | |
| | | | | WT | 1.333 | 0.373 | | | | |
| | | D3 | Long | KO | 1.778 | 0.434 | | -1.609 | 0.136 | |
| | | | | WT | 0.889 | 0.261 | | | | |
| | | D4 | Long | KO | 1.889 | 0.389 | | -0.047 | 1.000 | |
| | | | | WT | 1.889 | 0.309 | | | | |
| | | D5 | Long | KO | 1.778 | 0.324 | | -0.752 | 0.489 | |
| | | | | WT | 2.111 | 0.351 | | | | |
| | | D6 | Long | KO | 2.667 | 0.333 | | -0.537 | 0.666 | |
| | | | | WT | 2.444 | 0.338 | | | | |
| | | D7 | Long | KO | 2.667 | 0.333 | | -0.691 | 0.546 | |
| | | | | WT | 2.333 | 0.333 | | | | |
| | | D8 | Long | KO | 2.778 | 0.401 | | -1.178 | 0.297 | |
| | | | | WT | 2.222 | 0.147 | | | | |
| | | D9 | Long | KO | 3.111 | 0.389 | | -0.726 | 0.489 | |
| | | | | WT | 2.889 | 0.564 | | | | |
| | | D10 | Long | KO | 3.111 | 0.455 | | -0.971 | 0.387 | |
| | | | | WT | 2.556 | 0.176 | | | | |
| | | D11 | Long | KO | 2.556 | 0.580 | | -1.187 | 0.297 | |
| | | | | WT | 3.000 | 0.167 | | | | |
| | | D12 | Long | KO | 3.222 | 0.619 | | -0.548 | 0.605 | |
| | | | | WT | 2.667 | 0.373 | | | | |
| | | D13 | Long | KO | 2.889 | 0.423 | | -0.092 | 0.931 | |
| | | | | WT | 2.889 | 0.261 | | | | |
| | | D14 | Long | KO | 2.778 | 0.324 | | -0.144 | 0.931 | |
| | | | | WT | 2.778 | 0.222 | | | | |
| | | D15 | Long | KO | 3.333 | 0.441 | | -0.867 | 0.436 | |
| | | | | WT | 2.889 | 0.309 | | | | |
| | | D16 | Long | KO | 3.222 | 0.364 | | -0.188 | 0.863 | |
| | | | | WT | 3.111 | 0.200 | | | | |
| Table S3 | Figure S2D | D0 | Short | KO | 1.444 | 0.176 | Mann-Whitney U Test | -2.585 | 0.024 | * |
| | | | | WT | 0.667 | 0.167 | | | | |
| | | D1 | Short | KO | 1.333 | 0.236 | | -1.222 | 0.297 | |
| | | | | WT | 1.000 | 0.289 | | | | |
| | | D2 | Short | KO | 1.556 | 0.242 | | -1.675 | 0.136 | |
| | | | | WT | 1.000 | 0.236 | | | | |
| | | D3 | Short | KO | 1.667 | 0.289 | | -0.372 | 0.730 | |
| | | | | WT | 1.556 | 0.338 | | | | |
| | | D4 | Short | KO | 2.444 | 0.444 | | -0.523 | 0.666 | |
| | | | | WT | 2.000 | 0.236 | | | | |
| | | D5 | Short | KO | 2.111 | 0.261 | | -0.595 | 0.605 | |
| | | | | WT | 2.333 | 0.167 | | | | |
| | | D6 | Short | KO | 2.667 | 0.289 | | -1.866 | 0.077 | |
| | | | | WT | 1.889 | 0.261 | | | | |
| | | D7 | Short | KO | 2.889 | 0.423 | | -1.509 | 0.161 | |
| | | | | WT | 2.000 | 0.289 | | | | |
| | | D8 | Short | KO | 3.111 | 0.484 | | -1.655 | 0.113 | |
| | | | | WT | 2.111 | 0.261 | | | | |
| | | D9 | Short | KO | 2.556 | 0.294 | | -0.241 | 0.863 | |
| | | | | WT | 2.444 | 0.242 | | | | |
| | | D10 | Short | KO | 2.778 | 0.222 | | -0.530 | 0.666 | |
| | | | | WT | 2.556 | 0.294 | | | | |
| | | D11 | Short | KO | 2.778 | 0.465 | | -0.046 | 1.000 | |
| | | | | WT | 2.778 | 0.278 | | | | |
| | | D12 | Short | KO | 3.000 | 0.601 | | -0.091 | 0.931 | |
| | | | | WT | 2.778 | 0.222 | | | | |
| | | D13 | Short | KO | 3.333 | 0.471 | | -2.062 | 0.063 | |
| | | | | WT | 2.222 | 0.147 | | | | |
| | | D14 | Short | KO | 3.444 | 0.475 | | -1.521 | 0.161 | |
| | | | | WT | 2.444 | 0.294 | | | | |
| | | D15 | Short | KO | 3.000 | 0.373 | | -1.011 | 0.340 | |
| | | | | WT | 2.667 | 0.471 | | | | |
| | | D16 | Short | KO | 3.111 | 0.423 | | -1.235 | 0.258 | |
| | | | | WT | 2.444 | 0.294 | | | | |

| Table | Related Figure | Day | Group | Condition | Mean | SEM | Stastistical test | Z | P | * |
|---|---|---|---|---|---|---|---|---|---|---|
| | | | | | FHE peak frequency | | | | | |
| Table S3 | Figure S2D | D0 | KO | Long | 1.444 | 0.338 | Wilcoxon Signed Ranks Test | -0.108 | 0.914 | |
| | | | | Short | 1.444 | 0.176 | | | | |
| | | D1 | KO | Long | 0.778 | 0.222 | | -1.667 | 0.096 | |
| | | | | Short | 1.333 | 0.236 | | | | |
| | | D2 | KO | Long | 1.222 | 0.278 | | -1.732 | 0.083 | |
| | | | | Short | 1.556 | 0.242 | | | | |
| | | D3 | KO | Long | 1.778 | 0.434 | | -0.276 | 0.783 | |
| | | | | Short | 1.667 | 0.289 | | | | |
| | | D4 | KO | Long | 1.889 | 0.389 | | -1.406 | 0.160 | |
| | | | | Short | 2.444 | 0.444 | | | | |
| | | D5 | KO | Long | 1.778 | 0.324 | | -1.342 | 0.180 | |
| | | | | Short | 2.111 | 0.261 | | | | |
| | | D6 | KO | Long | 2.667 | 0.333 | | 0.000 | 1.000 | |
| | | | | Short | 2.667 | 0.289 | | | | |
| | | D7 | KO | Long | 2.667 | 0.333 | | -0.632 | 0.527 | |
| | | | | Short | 2.889 | 0.423 | | | | |
| | | D8 | KO | Long | 2.778 | 0.401 | | -0.707 | 0.480 | |
| | | | | Short | 3.111 | 0.484 | | | | |
| | | D9 | KO | Long | 3.111 | 0.389 | | -1.633 | 0.102 | |
| | | | | Long | 2.556 | 0.294 | | | | |
| | | D10 | KO | Short | 3.111 | 0.455 | | -1.000 | 0.317 | |
| | | | | Long | 2.778 | 0.222 | | | | |
| | | D11 | KO | Short | 2.556 | 0.580 | | -0.520 | 0.603 | |
| | | | | Long | 2.778 | 0.465 | | | | |
| | | D12 | KO | Short | 3.222 | 0.619 | | -0.439 | 0.660 | |
| | | | | Long | 3.000 | 0.601 | | | | |
| | | D13 | KO | Short | 2.889 | 0.423 | | -1.027 | 0.305 | |
| | | | | Long | 3.333 | 0.471 | | | | |
| | | D14 | KO | Short | 2.778 | 0.324 | | -1.730 | 0.084 | |
| | | | | Long | 3.444 | 0.475 | | | | |
| | | D15 | KO | Short | 3.333 | 0.441 | | -0.780 | 0.435 | |
| | | | | Long | 3.000 | 0.373 | | | | |
| | | D16 | KO | Short | 3.222 | 0.364 | | -0.431 | 0.666 | |
| | | | | Long | 3.111 | 0.423 | | | | |
| Table S3 | Figure S2D | D0 | WT | Long | 0.667 | 0.236 | Wilcoxon Signed Ranks Test | 0.000 | 1.000 | |
| | | | | Short | 0.667 | 0.167 | | | | |
| | | D1 | WT | Long | 0.778 | 0.222 | | -1.000 | 0.317 | |
| | | | | Short | 1.000 | 0.289 | | | | |
| | | D2 | WT | Long | 1.333 | 0.373 | | -1.134 | 0.257 | |
| | | | | Short | 1.000 | 0.236 | | | | |
| | | D3 | WT | Long | 0.889 | 0.261 | | -1.897 | 0.058 | |
| | | | | Short | 1.556 | 0.338 | | | | |
| | | D4 | WT | Long | 1.889 | 0.309 | | -0.378 | 0.705 | |
| | | | | Short | 2.000 | 0.236 | | | | |
| | | D5 | WT | Long | 2.111 | 0.351 | | -0.632 | 0.527 | |
| | | | | Short | 2.333 | 0.167 | | | | |
| | | D6 | WT | Long | 2.444 | 0.338 | | -1.406 | 0.160 | |
| | | | | Short | 1.889 | 0.261 | | | | |
| | | D7 | WT | Long | 2.333 | 0.333 | | -0.749 | 0.454 | |
| | | | | Short | 2.000 | 0.289 | | | | |
| | | D8 | WT | Long | 2.222 | 0.147 | | -0.378 | 0.705 | |
| | | | | Short | 2.111 | 0.261 | | | | |
| | | D9 | WT | Long | 2.889 | 0.564 | | -0.597 | 0.551 | |
| | | | | Long | 2.444 | 0.242 | | | | |
| | | D10 | WT | Short | 2.556 | 0.176 | | 0.000 | 1.000 | |
| | | | | Long | 2.556 | 0.294 | | | | |
| | | D11 | WT | Short | 3.000 | 0.167 | | -1.000 | 0.317 | |
| | | | | Long | 2.778 | 0.278 | | | | |
| | | D12 | WT | Short | 2.667 | 0.373 | | -0.378 | 0.705 | |
| | | | | Long | 2.778 | 0.222 | | | | |
| | | D13 | WT | Short | 2.889 | 0.261 | | -1.730 | 0.084 | |
| | | | | Long | 2.222 | 0.147 | | | | |
| | | D14 | WT | Short | 2.778 | 0.222 | | -0.828 | 0.408 | |
| | | | | Long | 2.444 | 0.294 | | | | |
| | | D15 | WT | Short | 2.889 | 0.309 | | -0.702 | 0.483 | |
| | | | | Long | 2.667 | 0.471 | | | | |
| | | D16 | WT | Short | 3.111 | 0.200 | | -1.667 | 0.096 | |
| | | | | Long | 2.444 | 0.294 | | | | |

3_4

| Table | Related Figure | Day | Condition | Group | Mean | SEM | Stastistical test | Z | P | * |
|---|---|---|---|---|---|---|---|---|---|---|
| | | | | | FHE peak latency | | | | | |
| Table S3 | Figure S2D | D0 | Long | KO | 3.556 | 1.029 | Mann-Whitney U Test | -1.993 | 0.063 | |
| | | | | WT | 1.444 | 0.959 | | | | |
| | | D1 | Long | KO | 1.000 | 0.408 | | -0.389 | 0.730 | |
| | | | | WT | 1.667 | 0.764 | | | | |
| | | D2 | Long | KO | 2.667 | 1.027 | | -0.181 | 0.863 | |
| | | | | WT | 3.111 | 1.274 | | | | |
| | | D3 | Long | KO | 2.556 | 0.835 | | -0.975 | 0.387 | |
| | | | | WT | 1.556 | 0.747 | | | | |
| | | D4 | Long | KO | 2.444 | 0.930 | | -0.869 | 0.436 | |
| | | | | WT | 1.222 | 0.434 | | | | |
| | | D5 | Long | KO | 2.778 | 1.051 | | -0.677 | 0.546 | |
| | | | | WT | 3.000 | 0.782 | | | | |
| | | D6 | Long | KO | 3.111 | 1.006 | | -0.045 | 1.000 | |
| | | | | WT | 3.222 | 0.969 | | | | |
| | | D7 | Long | KO | 2.000 | 0.667 | | -1.083 | 0.297 | |
| | | | | WT | 3.667 | 0.972 | | | | |
| | | D8 | Long | KO | 1.000 | 0.373 | | -1.814 | 0.077 | |
| | | | | WT | 2.444 | 0.626 | | | | |
| | | D9 | Long | KO | 1.889 | 0.423 | | -0.547 | 0.605 | |
| | | | | WT | 1.556 | 0.412 | | | | |
| | | D10 | Long | KO | 2.444 | 0.412 | | -1.959 | 0.063 | |
| | | | | WT | 1.222 | 0.364 | | | | |
| | | D11 | Long | KO | 2.444 | 0.709 | | -0.723 | 0.489 | |
| | | | | WT | 3.556 | 0.915 | | | | |
| | | D12 | Long | KO | 1.333 | 0.167 | | -0.153 | 0.931 | |
| | | | | WT | 1.444 | 0.377 | | | | |
| | | D13 | Long | KO | 2.000 | 0.782 | | -0.790 | 0.489 | |
| | | | | WT | 2.333 | 0.577 | | | | |
| | | D14 | Long | KO | 2.556 | 0.801 | | -0.136 | 0.931 | |
| | | | | WT | 2.333 | 0.799 | | | | |
| | | D15 | Long | KO | 2.000 | 0.527 | | -0.895 | 0.387 | |
| | | | | WT | 3.333 | 0.957 | | | | |
| | | D16 | Long | KO | 2.444 | 0.689 | | -1.490 | 0.161 | |
| | | | | WT | 3.778 | 0.760 | | | | |
| Table S3 | Figure S2D | D0 | Short | KO | 4.778 | 0.862 | Mann-Whitney U Test | -0.808 | 0.436 | |
| | | | | WT | 3.556 | 1.226 | | | | |
| | | D1 | Short | KO | 2.333 | 0.913 | | -1.473 | 0.161 | |
| | | | | WT | 3.889 | 0.807 | | | | |
| | | D2 | Short | KO | 2.667 | 0.913 | | -0.495 | 0.666 | |
| | | | | WT | 1.889 | 0.676 | | | | |
| | | D3 | Short | KO | 3.222 | 1.038 | | -0.269 | 0.796 | |
| | | | | WT | 2.556 | 0.766 | | | | |
| | | D4 | Short | KO | 3.444 | 1.056 | | -2.139 | 0.040 | * |
| | | | | WT | 0.667 | 0.236 | | | | |
| | | D5 | Short | KO | 2.333 | 0.707 | | -1.072 | 0.297 | |
| | | | | WT | 3.667 | 0.882 | | | | |
| | | D6 | Short | KO | 2.889 | 0.696 | | -0.090 | 0.931 | |
| | | | | WT | 2.889 | 0.904 | | | | |
| | | D7 | Short | KO | 1.889 | 0.512 | | -0.725 | 0.489 | |
| | | | | WT | 1.333 | 0.441 | | | | |
| | | D8 | Short | KO | 1.889 | 0.389 | | -0.507 | 0.666 | |
| | | | | WT | 2.222 | 0.894 | | | | |
| | | D9 | Short | KO | 1.111 | 0.309 | | -2.653 | 0.008 | ** |
| | | | | WT | 3.000 | 0.471 | | | | |
| | | D10 | Short | KO | 2.000 | 0.553 | | -0.095 | 0.931 | |
| | | | | WT | 2.000 | 0.601 | | | | |
| | | D11 | Short | KO | 1.556 | 0.503 | | -0.633 | 0.546 | |
| | | | | WT | 2.333 | 0.764 | | | | |
| | | D12 | Short | KO | 1.111 | 0.261 | | -1.338 | 0.222 | |
| | | | | WT | 2.111 | 0.588 | | | | |
| | | D13 | Short | KO | 2.111 | 0.423 | | -0.596 | 0.605 | |
| | | | | WT | 2.333 | 0.441 | | | | |
| | | D14 | Short | KO | 1.222 | 0.364 | | -0.654 | 0.546 | |
| | | | | WT | 1.778 | 0.547 | | | | |
| | | D15 | Short | KO | 1.444 | 0.556 | | -0.683 | 0.546 | |
| | | | | WT | 2.222 | 0.795 | | | | |
| | | D16 | Short | KO | 1.556 | 0.294 | | -0.320 | 0.796 | |
| | | | | WT | 1.778 | 0.465 | | | | |

| Table | Related Figure | Day | Group | Condition | Mean | SEM | Stastistical test | Z | P | * |
|---|---|---|---|---|---|---|---|---|---|---|
| | | | | | FHE peak latency | | | | | |
| Table S3 | Figure S2D | D0 | KO | Long | 3.556 | 1.029 | Wilcoxon Signed Ranks Test | -0.704 | 0.481 | |
| | | | | Short | 4.778 | 0.862 | | | | |
| | | D1 | KO | Long | 1.000 | 0.408 | | -1.420 | 0.156 | |
| | | | | Short | 2.333 | 0.913 | | | | |
| | | D2 | KO | Long | 2.667 | 1.027 | | 0.000 | 1.000 | |
| | | | | Short | 2.667 | 0.913 | | | | |
| | | D3 | KO | Long | 2.556 | 0.835 | | -0.282 | 0.778 | |
| | | | | Short | 3.222 | 1.038 | | | | |
| | | D4 | KO | Long | 2.444 | 0.930 | | -0.422 | 0.673 | |
| | | | | Short | 3.444 | 1.056 | | | | |
| | | D5 | KO | Long | 2.778 | 1.051 | | -0.750 | 0.453 | |
| | | | | Short | 2.333 | 0.707 | | | | |
| | | D6 | KO | Long | 3.111 | 1.006 | | -0.170 | 0.865 | |
| | | | | Short | 2.889 | 0.696 | | | | |
| | | D7 | KO | Long | 2.000 | 0.667 | | -0.355 | 0.723 | |
| | | | | Short | 1.889 | 0.512 | | | | |
| | | D8 | KO | Long | 1.000 | 0.373 | | -1.730 | 0.084 | |
| | | | | Short | 1.889 | 0.389 | | | | |
| | | D9 | KO | Long | 1.889 | 0.423 | | -1.633 | 0.102 | |
| | | | | Long | 1.111 | 0.309 | | | | |
| | | D10 | KO | Short | 2.444 | 0.412 | | -0.660 | 0.509 | |
| | | | | Long | 2.000 | 0.553 | | | | |
| | | D11 | KO | Short | 2.444 | 0.709 | | -0.940 | 0.347 | |
| | | | | Long | 1.556 | 0.503 | | | | |
| | | D12 | KO | Short | 1.333 | 0.167 | | -0.707 | 0.480 | |
| | | | | Long | 1.111 | 0.261 | | | | |
| | | D13 | KO | Short | 2.000 | 0.782 | | -0.271 | 0.786 | |
| | | | | Long | 2.111 | 0.423 | | | | |
| | | D14 | KO | Short | 2.556 | 0.801 | | -1.450 | 0.147 | |
| | | | | Long | 1.222 | 0.364 | | | | |
| | | D15 | KO | Short | 2.000 | 0.527 | | -0.736 | 0.461 | |
| | | | | Long | 1.444 | 0.556 | | | | |
| | | D16 | KO | Short | 2.444 | 0.689 | | -0.780 | 0.436 | |
| | | | | Long | 1.556 | 0.294 | | | | |
| Table S3 | Figure S2D | D0 | WT | Long | 1.444 | 0.959 | Wilcoxon Signed Ranks Test | -1.153 | 0.249 | |
| | | | | Short | 3.556 | 1.226 | | | | |
| | | D1 | WT | Long | 1.667 | 0.764 | | -1.781 | 0.075 | |
| | | | | Short | 3.889 | 0.807 | | | | |
| | | D2 | WT | Long | 3.111 | 1.274 | | -0.933 | 0.351 | |
| | | | | Short | 1.889 | 0.676 | | | | |
| | | D3 | WT | Long | 1.556 | 0.747 | | -0.705 | 0.481 | |
| | | | | Short | 2.556 | 0.766 | | | | |
| | | D4 | WT | Long | 1.222 | 0.434 | | -1.222 | 0.222 | |
| | | | | Short | 0.667 | 0.236 | | | | |
| | | D5 | WT | Long | 3.000 | 0.782 | | -0.563 | 0.574 | |
| | | | | Short | 3.667 | 0.882 | | | | |
| | | D6 | WT | Long | 3.222 | 0.969 | | -0.426 | 0.670 | |
| | | | | Short | 2.889 | 0.904 | | | | |
| | | D7 | WT | Long | 3.667 | 0.972 | | -1.689 | 0.091 | |
| | | | | Short | 1.333 | 0.441 | | | | |
| | | D8 | WT | Long | 2.444 | 0.626 | | -0.181 | 0.856 | |
| | | | | Short | 2.222 | 0.894 | | | | |
| | | D9 | WT | Long | 1.556 | 0.412 | | -1.975 | 0.048 | |
| | | | | Long | 3.000 | 0.471 | | | | |
| | | D10 | WT | Short | 1.222 | 0.364 | | -0.962 | 0.336 | |
| | | | | Long | 2.000 | 0.601 | | | | |
| | | D11 | WT | Short | 3.556 | 0.915 | | -1.273 | 0.203 | |
| | | | | Long | 2.333 | 0.764 | | | | |
| | | D12 | WT | Short | 1.444 | 0.377 | | -0.948 | 0.343 | |
| | | | | Long | 2.111 | 0.588 | | | | |
| | | D13 | WT | Short | 2.333 | 0.577 | | -0.171 | 0.864 | |
| | | | | Long | 2.333 | 0.441 | | | | |
| | | D14 | WT | Short | 2.333 | 0.799 | | -0.877 | 0.380 | |
| | | | | Long | 1.778 | 0.547 | | | | |
| | | D15 | WT | Short | 3.333 | 0.957 | | -0.431 | 0.667 | |
| | | | | Long | 2.222 | 0.795 | | | | |
| | | D16 | WT | Short | 3.778 | 0.760 | | -2.401 | 0.016 | * |
| | | | | Long | 1.778 | 0.465 | | | | |

3_5

HE vs. FHE peak frequencies (KO)

| Table | Related Figure | Day | Condition | Pair | Mean | SEM | Stastistical test | Z | P | * |
|---|---|---|---|---|---|---|---|---|---|---|
| Table S3 | Figure S3A | D0 | Long | HE | 1.222 | 0.222 | Wilcoxon Signed Ranks Test | -1.414 | 0.315 | |
| | | | | FHE | 1.440 | 1.030 | | | | |
| | | D1 | Long | HE | 0.778 | 0.222 | | 0.000 | 1.000 | |
| | | | | FHE | 0.778 | 0.222 | | | | |
| | | D2 | Long | HE | 1.222 | 0.278 | | 0.000 | 1.000 | |
| | | | | FHE | 1.222 | 0.278 | | | | |
| | | D3 | Long | HE | 2.222 | 0.494 | | -1.633 | 0.205 | |
| | | | | FHE | 1.222 | 0.434 | | | | |
| | | D4 | Long | HE | 2.778 | 0.619 | | -1.841 | 0.131 | |
| | | | | FHE | 1.778 | 0.389 | | | | |
| | | D5 | Long | HE | 2.889 | 0.423 | | -2.456 | 0.028 | * |
| | | | | FHE | 1.778 | 0.324 | | | | |
| | | D6 | Long | HE | 4.111 | 0.423 | | -2.232 | 0.051 | |
| | | | | FHE | 2.667 | 0.333 | | | | |
| | | D7 | Long | HE | 3.889 | 0.676 | | -1.826 | 0.136 | |
| | | | | FHE | 2.667 | 0.333 | | | | |
| | | D8 | Long | HE | 5.222 | 0.683 | | -2.558 | 0.021 | * |
| | | | | FHE | 2.778 | 0.401 | | | | |
| | | D9 | Long | HE | 5.222 | 0.596 | | -2.555 | 0.021 | * |
| | | | | FHE | 3.111 | 0.389 | | | | |
| | | D10 | Long | HE | 5.111 | 0.611 | | -2.388 | 0.034 | * |
| | | | | FHE | 3.111 | 0.455 | | | | |
| | | D11 | Long | HE | 5.111 | 0.964 | | -2.692 | 0.014 | ** |
| | | | | FHE | 2.556 | 0.580 | | | | |
| | | D12 | Long | HE | 4.444 | 0.648 | | -2.060 | 0.079 | |
| | | | | FHE | 3.222 | 0.619 | | | | |
| | | D13 | Long | HE | 5.333 | 0.667 | | -2.539 | 0.022 | * |
| | | | | FHE | 2.889 | 0.423 | | | | |
| | | D14 | Long | HE | 4.222 | 0.401 | | -2.565 | 0.021 | * |
| | | | | FHE | 2.778 | 0.324 | | | | |
| | | D15 | Long | HE | 4.667 | 0.471 | | -2.401 | 0.033 | * |
| | | | | FHE | 3.333 | 0.441 | | | | |
| | | D16 | Long | HE | 4.667 | 0.471 | | -2.060 | 0.079 | |
| | | | | FHE | 3.222 | 0.364 | | | | |
| Table S3 | Figure S3A | D0 | Srhot | HE | 1.111 | 0.200 | Wilcoxon Signed Ranks Test | -1.732 | 0.167 | |
| | | | | FHE | 1.444 | 0.862 | | | | |
| | | D1 | Srhot | HE | 1.889 | 0.588 | | -0.816 | 0.828 | |
| | | | | FHE | 1.333 | 0.236 | | | | |
| | | D2 | Srhot | HE | 2.111 | 0.309 | | -1.890 | 0.118 | |
| | | | | FHE | 1.556 | 0.242 | | | | |
| | | D3 | Srhot | HE | 2.556 | 0.444 | | -1.841 | 0.131 | |
| | | | | FHE | 1.667 | 0.289 | | | | |
| | | D4 | Srhot | HE | 2.889 | 0.588 | | -2.000 | 0.091 | |
| | | | | FHE | 2.444 | 0.444 | | | | |
| | | D5 | Srhot | HE | 2.667 | 0.236 | | -1.633 | 0.205 | |
| | | | | FHE | 2.111 | 0.261 | | | | |
| | | D6 | Srhot | HE | 3.444 | 0.530 | | -1.841 | 0.131 | |
| | | | | FHE | 2.667 | 0.289 | | | | |
| | | D7 | Srhot | HE | 3.556 | 0.444 | | -2.121 | 0.068 | |
| | | | | FHE | 2.889 | 0.423 | | | | |
| | | D8 | Srhot | HE | 4.667 | 0.624 | | -2.410 | 0.032 | * |
| | | | | FHE | 3.111 | 0.484 | | | | |
| | | D9 | Srhot | HE | 3.667 | 0.624 | | -1.841 | 0.131 | |
| | | | | FHE | 2.556 | 0.294 | | | | |
| | | D10 | Srhot | HE | 3.222 | 0.278 | | -2.000 | 0.091 | |
| | | | | FHE | 2.778 | 0.222 | | | | |
| | | D11 | Srhot | HE | 3.778 | 0.364 | | -2.264 | 0.047 | * |
| | | | | FHE | 2.778 | 0.465 | | | | |
| | | D12 | Srhot | HE | 3.778 | 0.547 | | -1.890 | 0.118 | |
| | | | | FHE | 3.000 | 0.601 | | | | |
| | | D13 | Srhot | HE | 3.556 | 0.475 | | -1.414 | 0.315 | |
| | | | | FHE | 3.333 | 0.471 | | | | |
| | | D14 | Srhot | HE | 4.222 | 0.547 | | -1.841 | 0.131 | |
| | | | | FHE | 3.444 | 0.475 | | | | |
| | | D15 | Srhot | HE | 3.444 | 0.648 | | -1.342 | 0.359 | |
| | | | | FHE | 3.000 | 0.373 | | | | |
| | | D16 | Srhot | HE | 3.444 | 0.556 | | -1.732 | 0.167 | |
| | | | | FHE | 3.111 | 0.423 | | | | |

HE vs. FHE peak frequencies (WT)

| Table | Related Figure | Day | Condition | Pair | Mean | SEM | Stastistical test | Z | P | * |
|---|---|---|---|---|---|---|---|---|---|---|
| Table S3 | Figure S3A | D0 | Long | HE | 0.778 | 0.434 | Wilcoxon Signed Ranks Test | -0.447 | 1.000 | |
| | | | | FHE | 0.670 | 0.960 | | | | |
| | | D1 | Long | HE | 1.000 | 0.289 | | -0.707 | 0.959 | |
| | | | | FHE | 0.778 | 0.222 | | | | |
| | | D2 | Long | HE | 1.667 | 0.471 | | -0.966 | 0.668 | |
| | | | | FHE | 1.333 | 0.373 | | | | |
| | | D3 | Long | HE | 2.111 | 0.351 | | -2.428 | 0.030 | * |
| | | | | FHE | 0.889 | 0.261 | | | | |
| | | D4 | Long | HE | 2.444 | 0.294 | | -1.890 | 0.118 | |
| | | | | FHE | 1.889 | 0.309 | | | | |
| | | D5 | Long | HE | 3.000 | 0.527 | | -1.841 | 0.131 | |
| | | | | FHE | 2.111 | 0.351 | | | | |
| | | D6 | Long | HE | 3.222 | 0.278 | | -2.070 | 0.077 | |
| | | | | FHE | 2.444 | 0.338 | | | | |
| | | D7 | Long | HE | 3.333 | 0.408 | | -2.264 | 0.047 | * |
| | | | | FHE | 2.333 | 0.333 | | | | |
| | | D8 | Long | HE | 3.556 | 0.338 | | -2.414 | 0.032 | * |
| | | | | FHE | 2.222 | 0.147 | | | | |
| | | D9 | Long | HE | 4.222 | 0.662 | | -2.264 | 0.047 | * |
| | | | | FHE | 2.889 | 0.564 | | | | |
| | | D10 | Long | HE | 3.889 | 0.261 | | -2.585 | 0.019 | * |
| | | | | FHE | 2.556 | 0.176 | | | | |
| | | D11 | Long | HE | 4.444 | 0.556 | | -2.226 | 0.052 | |
| | | | | FHE | 3.000 | 0.167 | | | | |
| | | D12 | Long | HE | 4.667 | 0.577 | | -2.410 | 0.032 | * |
| | | | | FHE | 2.667 | 0.373 | | | | |
| | | D13 | Long | HE | 4.333 | 0.373 | | -2.414 | 0.032 | * |
| | | | | FHE | 2.889 | 0.261 | | | | |
| | | D14 | Long | HE | 4.333 | 0.408 | | -2.401 | 0.033 | * |
| | | | | FHE | 2.778 | 0.222 | | | | |
| | | D15 | Long | HE | 4.444 | 0.377 | | -2.585 | 0.019 | * |
| | | | | FHE | 2.889 | 0.309 | | | | |
| | | D16 | Long | HE | 5.000 | 0.408 | | -2.555 | 0.021 | * |
| | | | | FHE | 3.111 | 0.200 | | | | |
| Table S3 | Figure S3A | D0 | Srhot | HE | 1.111 | 0.351 | Wilcoxon Signed Ranks Test | -1.414 | 0.315 | |
| | | | | FHE | 0.667 | 1.226 | | | | |
| | | D1 | Srhot | HE | 1.000 | 0.236 | | 0.000 | 1.000 | |
| | | | | FHE | 1.000 | 0.289 | | | | |
| | | D2 | Srhot | HE | 1.222 | 0.222 | | -1.000 | 0.635 | |
| | | | | FHE | 1.000 | 0.236 | | | | |
| | | D3 | Srhot | HE | 2.667 | 0.500 | | -2.456 | 0.028 | * |
| | | | | FHE | 1.556 | 0.338 | | | | |
| | | D4 | Srhot | HE | 2.889 | 0.564 | | -1.604 | 0.218 | |
| | | | | FHE | 2.000 | 0.236 | | | | |
| | | D5 | Srhot | HE | 3.222 | 0.547 | | -1.633 | 0.205 | |
| | | | | FHE | 2.333 | 0.167 | | | | |
| | | D6 | Srhot | HE | 2.333 | 0.373 | | -2.000 | 0.091 | |
| | | | | FHE | 1.889 | 0.261 | | | | |
| | | D7 | Srhot | HE | 3.111 | 0.351 | | -2.640 | 0.017 | ** |
| | | | | FHE | 2.000 | 0.289 | | | | |
| | | D8 | Srhot | HE | 3.000 | 0.289 | | -2.271 | 0.046 | * |
| | | | | FHE | 2.111 | 0.261 | | | | |
| | | D9 | Srhot | HE | 3.444 | 0.503 | | -2.121 | 0.068 | |
| | | | | FHE | 2.444 | 0.242 | | | | |
| | | D10 | Srhot | HE | 3.333 | 0.289 | | -1.890 | 0.118 | |
| | | | | FHE | 2.556 | 0.294 | | | | |
| | | D11 | Srhot | HE | 3.333 | 0.471 | | -1.633 | 0.205 | |
| | | | | FHE | 2.778 | 0.278 | | | | |
| | | D12 | Srhot | HE | 3.778 | 0.364 | | -1.841 | 0.131 | |
| | | | | FHE | 2.778 | 0.222 | | | | |
| | | D13 | Srhot | HE | 2.444 | 0.176 | | -1.414 | 0.315 | |
| | | | | FHE | 2.222 | 0.147 | | | | |
| | | D14 | Srhot | HE | 3.111 | 0.261 | | -2.121 | 0.068 | |
| | | | | FHE | 2.444 | 0.294 | | | | |
| | | D15 | Srhot | HE | 3.444 | 0.444 | | -1.890 | 0.118 | |
| | | | | FHE | 2.667 | 0.471 | | | | |
| | | D16 | Srhot | HE | 2.889 | 0.200 | | -1.633 | 0.205 | |
| | | | | FHE | 2.444 | 0.294 | | | | |

3_6

HE vs. FHE peak latencies (KO)

| Table | Related Figure | Day | Condition | Pair | Mean | SEM | Stastistical test | Z | P | * |
|---|---|---|---|---|---|---|---|---|---|---|
| Table S3 | Figure S3A | D0 | Long | HE | 4.000 | 1.093 | Wilcoxon Signed Ranks Test | -1.342 | 0.359 | |
| | | | | FHE | 3.556 | 1.029 | | | | |
| | | D1 | Long | HE | 1.000 | 0.408 | | 0.000 | 1.000 | |
| | | | | FHE | 1.000 | 0.408 | | | | |
| | | D2 | Long | HE | 4.222 | 1.310 | | -1.604 | 0.218 | |
| | | | | FHE | 2.667 | 1.027 | | | | |
| | | D3 | Long | HE | 3.667 | 1.291 | | -0.447 | 1.000 | |
| | | | | FHE | 2.556 | 0.835 | | | | |
| | | D4 | Long | HE | 2.889 | 1.296 | | -0.447 | 1.000 | |
| | | | | FHE | 2.444 | 0.930 | | | | |
| | | D5 | Long | HE | 4.667 | 1.202 | | -0.813 | 0.833 | |
| | | | | FHE | 2.778 | 1.051 | | | | |
| | | D6 | Long | HE | 4.222 | 1.164 | | -0.730 | 0.930 | |
| | | | | FHE | 3.111 | 1.006 | | | | |
| | | D7 | Long | HE | 3.556 | 1.215 | | -1.342 | 0.359 | |
| | | | | FHE | 2.000 | 0.667 | | | | |
| | | D8 | Long | HE | 3.222 | 1.382 | | -1.826 | 0.136 | |
| | | | | FHE | 1.000 | 0.373 | | | | |
| | | D9 | Long | HE | 5.000 | 1.333 | | -1.841 | 0.131 | |
| | | | | FHE | 1.889 | 0.423 | | | | |
| | | D10 | Long | HE | 5.111 | 1.172 | | -1.753 | 0.159 | |
| | | | | FHE | 2.444 | 0.412 | | | | |
| | | D11 | Long | HE | 5.556 | 1.600 | | -1.378 | 0.336 | |
| | | | | FHE | 2.444 | 0.709 | | | | |
| | | D12 | Long | HE | 3.111 | 1.252 | | -1.095 | 0.547 | |
| | | | | FHE | 1.333 | 0.167 | | | | |
| | | D13 | Long | HE | 5.111 | 1.348 | | -2.252 | 0.049 | * |
| | | | | FHE | 2.000 | 0.782 | | | | |
| | | D14 | Long | HE | 3.889 | 0.889 | | -1.841 | 0.131 | |
| | | | | FHE | 2.556 | 0.801 | | | | |
| | | D15 | Long | HE | 5.222 | 1.579 | | -1.826 | 0.136 | |
| | | | | FHE | 2.000 | 0.527 | | | | |
| | | D16 | Long | HE | 5.000 | 1.414 | | -1.992 | 0.093 | |
| | | | | FHE | 2.444 | 0.689 | | | | |
| Table S3 | Figure S3A | D0 | Srhot | HE | 3.667 | 1.312 | Wilcoxon Signed Ranks Test | -0.813 | 0.833 | |
| | | | | FHE | 4.778 | 0.862 | | | | |
| | | D1 | Srhot | HE | 3.333 | 1.312 | | -0.730 | 0.930 | |
| | | | | FHE | 2.333 | 0.913 | | | | |
| | | D2 | Srhot | HE | 6.222 | 1.289 | | -1.992 | 0.093 | |
| | | | | FHE | 2.667 | 0.913 | | | | |
| | | D3 | Srhot | HE | 5.667 | 1.607 | | -1.342 | 0.359 | |
| | | | | FHE | 3.222 | 1.038 | | | | |
| | | D4 | Srhot | HE | 3.444 | 1.271 | | 0.000 | 1.000 | |
| | | | | FHE | 3.444 | 1.056 | | | | |
| | | D5 | Srhot | HE | 2.667 | 0.645 | | -1.000 | 0.635 | |
| | | | | FHE | 2.333 | 0.707 | | | | |
| | | D6 | Srhot | HE | 2.778 | 0.741 | | -1.000 | 0.635 | |
| | | | | FHE | 2.889 | 0.696 | | | | |
| | | D7 | Srhot | HE | 2.778 | 1.392 | | -0.136 | 1.000 | |
| | | | | FHE | 1.889 | 0.512 | | | | |
| | | D8 | Srhot | HE | 3.333 | 1.067 | | -1.841 | 0.131 | |
| | | | | FHE | 1.889 | 0.389 | | | | |
| | | D9 | Srhot | HE | 2.556 | 0.801 | | -1.633 | 0.205 | |
| | | | | FHE | 1.111 | 0.309 | | | | |
| | | D10 | Srhot | HE | 2.222 | 0.683 | | -0.447 | 1.000 | |
| | | | | FHE | 2.000 | 0.553 | | | | |
| | | D11 | Srhot | HE | 4.000 | 1.190 | | -1.625 | 0.208 | |
| | | | | FHE | 1.556 | 0.503 | | | | |
| | | D12 | Srhot | HE | 2.111 | 0.772 | | -1.069 | 0.570 | |
| | | | | FHE | 1.111 | 0.261 | | | | |
| | | D13 | Srhot | HE | 2.556 | 0.899 | | -0.447 | 1.000 | |
| | | | | FHE | 2.111 | 0.423 | | | | |
| | | D14 | Srhot | HE | 1.667 | 0.408 | | -1.069 | 0.570 | |
| | | | | FHE | 1.222 | 0.364 | | | | |
| | | D15 | Srhot | HE | 1.333 | 0.577 | | -1.000 | 0.635 | |
| | | | | FHE | 1.444 | 0.556 | | | | |
| | | D16 | Srhot | HE | 1.444 | 0.377 | | -0.368 | 1.000 | |
| | | | | FHE | 1.556 | 0.294 | | | | |

HE vs. FHE peak latencies (WT)

| Table | Related Figure | Day | Condition | Pair | Mean | SEM | Stastistical test | Z | P | * |
|---|---|---|---|---|---|---|---|---|---|---|
| Table S3 | Figure S3A | D0 | Long | HE | 1.778 | 1.222 | Wilcoxon Signed Ranks Test | -1.000 | 0.635 | |
| | | | | FHE | 1.444 | 0.959 | | | | |
| | | D1 | Long | HE | 2.000 | 1.443 | | 0.000 | 1.000 | |
| | | | | FHE | 1.667 | 0.764 | | | | |
| | | D2 | Long | HE | 3.111 | 1.438 | | 0.000 | 1.000 | |
| | | | | FHE | 3.111 | 1.274 | | | | |
| | | D3 | Long | HE | 5.111 | 1.728 | | -1.753 | 0.159 | |
| | | | | FHE | 1.556 | 0.747 | | | | |
| | | D4 | Long | HE | 2.111 | 1.111 | | -0.730 | 0.930 | |
| | | | | FHE | 1.222 | 0.434 | | | | |
| | | D5 | Long | HE | 5.444 | 1.608 | | -1.342 | 0.359 | |
| | | | | FHE | 3.000 | 0.782 | | | | |
| | | D6 | Long | HE | 5.111 | 1.476 | | -1.604 | 0.218 | |
| | | | | FHE | 3.222 | 0.969 | | | | |
| | | D7 | Long | HE | 4.222 | 0.572 | | -1.089 | 0.553 | |
| | | | | FHE | 3.667 | 0.972 | | | | |
| | | D8 | Long | HE | 3.222 | 1.278 | | -0.593 | 1.000 | |
| | | | | FHE | 2.444 | 0.626 | | | | |
| | | D9 | Long | HE | 4.000 | 1.179 | | -2.023 | 0.086 | |
| | | | | FHE | 1.556 | 0.412 | | | | |
| | | D10 | Long | HE | 3.111 | 1.060 | | -1.581 | 0.228 | |
| | | | | FHE | 1.222 | 0.364 | | | | |
| | | D11 | Long | HE | 5.556 | 1.365 | | -0.931 | 0.703 | |
| | | | | FHE | 3.556 | 0.915 | | | | |
| | | D12 | Long | HE | 3.778 | 1.245 | | -1.841 | 0.131 | |
| | | | | FHE | 1.444 | 0.377 | | | | |
| | | D13 | Long | HE | 2.889 | 0.633 | | -1.289 | 0.395 | |
| | | | | FHE | 2.333 | 0.577 | | | | |
| | | D14 | Long | HE | 4.333 | 0.833 | | -2.032 | 0.084 | |
| | | | | FHE | 2.333 | 0.799 | | | | |
| | | D15 | Long | HE | 4.333 | 0.667 | | -1.633 | 0.205 | |
| | | | | FHE | 3.333 | 0.957 | | | | |
| | | D16 | Long | HE | 5.778 | 1.103 | | -1.980 | 0.096 | |
| | | | | FHE | 3.778 | 0.760 | | | | |
| Table S3 | Figure S3A | D0 | Srhot | HE | 4.000 | 1.616 | Wilcoxon Signed Ranks Test | -0.405 | 1.000 | |
| | | | | FHE | 3.556 | 1.226 | | | | |
| | | D1 | Srhot | HE | 4.333 | 1.258 | | -0.535 | 1.000 | |
| | | | | FHE | 3.889 | 0.807 | | | | |
| | | D2 | Srhot | HE | 1.889 | 1.020 | | -0.368 | 1.000 | |
| | | | | FHE | 1.889 | 0.676 | | | | |
| | | D3 | Srhot | HE | 3.556 | 1.692 | | -0.736 | 0.923 | |
| | | | | FHE | 2.556 | 0.766 | | | | |
| | | D4 | Srhot | HE | 2.333 | 1.202 | | -1.069 | 0.570 | |
| | | | | FHE | 0.667 | 0.236 | | | | |
| | | D5 | Srhot | HE | 2.667 | 0.816 | | -1.342 | 0.359 | |
| | | | | FHE | 3.667 | 0.882 | | | | |
| | | D6 | Srhot | HE | 1.778 | 0.969 | | -1.604 | 0.218 | |
| | | | | FHE | 2.889 | 0.904 | | | | |
| | | D7 | Srhot | HE | 4.111 | 1.567 | | -1.753 | 0.159 | |
| | | | | FHE | 1.333 | 0.441 | | | | |
| | | D8 | Srhot | HE | 4.444 | 1.271 | | -1.476 | 0.280 | |
| | | | | FHE | 2.222 | 0.894 | | | | |
| | | D9 | Srhot | HE | 3.889 | 0.484 | | -1.342 | 0.359 | |
| | | | | FHE | 3.000 | 0.471 | | | | |
| | | D10 | Srhot | HE | 4.000 | 1.404 | | -1.214 | 0.450 | |
| | | | | FHE | 2.000 | 0.601 | | | | |
| | | D11 | Srhot | HE | 3.889 | 1.047 | | -1.604 | 0.218 | |
| | | | | FHE | 2.333 | 0.764 | | | | |
| | | D12 | Srhot | HE | 3.222 | 0.862 | | -1.461 | 0.288 | |
| | | | | FHE | 2.111 | 0.588 | | | | |
| | | D13 | Srhot | HE | 2.222 | 0.521 | | -0.447 | 1.000 | |
| | | | | FHE | 2.333 | 0.441 | | | | |
| | | D14 | Srhot | HE | 3.778 | 1.064 | | -1.633 | 0.205 | |
| | | | | FHE | 1.778 | 0.547 | | | | |
| | | D15 | Srhot | HE | 3.333 | 0.726 | | -1.841 | 0.131 | |
| | | | | FHE | 2.222 | 0.795 | | | | |
| | | D16 | Srhot | HE | 2.000 | 0.471 | | -0.447 | 1.000 | |
| | | | | FHE | 1.778 | 0.465 | | | | |

## 3_7

| Correlation between HE and FHE peak frequencies (KO) | | | | | | | | | | |
|---|---|---|---|---|---|---|---|---|---|---|
| Table | Related Figure | Day | Condition | Pair | Mean | SEM | Stastistical test | Spearman's ρ | P | * |
| Table S3 | Figure S3B | D0 | Long | HE | 1.222 | 0.222 | Spearman's rank correlation coefficient | 0.266 | 0.976 | |
| | | | | FHE | 1.440 | 1.030 | | | | |
| | | D1 | Long | HE | 0.778 | 0.222 | | 0.813 | 0.016 | * |
| | | | | FHE | 0.778 | 0.222 | | | | |
| | | D2 | Long | HE | 1.222 | 0.278 | | 0.767 | 0.032 | * |
| | | | | FHE | 1.222 | 0.278 | | | | |
| | | D3 | Long | HE | 2.222 | 0.494 | | 0.815 | 0.015 | * |
| | | | | FHE | 1.778 | 0.434 | | | | |
| | | D4 | Long | HE | 2.778 | 0.619 | | 0.700 | 0.071 | |
| | | | | FHE | 1.889 | 0.389 | | | | |
| | | D5 | Long | HE | 2.889 | 0.423 | | 0.523 | 0.298 | |
| | | | | FHE | 1.778 | 0.324 | | | | |
| | | D6 | Long | HE | 4.111 | 0.423 | | 0.454 | 0.440 | |
| | | | | FHE | 2.667 | 0.333 | | | | |
| | | D7 | Long | HE | 3.889 | 0.676 | | 0.467 | 0.409 | |
| | | | | FHE | 2.667 | 0.333 | | | | |
| | | D8 | Long | HE | 5.222 | 0.683 | | 0.860 | 0.006 | ** |
| | | | | FHE | 2.778 | 0.401 | | | | |
| | | D9 | Long | HE | 5.222 | 0.596 | | 0.624 | 0.145 | |
| | | | | FHE | 3.111 | 0.389 | | | | |
| | | D10 | Long | HE | 5.111 | 0.611 | | 0.557 | 0.238 | |
| | | | | FHE | 3.111 | 0.455 | | | | |
| | | D11 | Long | HE | 5.111 | 0.964 | | 0.749 | 0.040 | * |
| | | | | FHE | 2.556 | 0.580 | | | | |
| | | D12 | Long | HE | 4.444 | 0.648 | | 0.306 | 0.847 | |
| | | | | FHE | 3.222 | 0.619 | | | | |
| | | D13 | Long | HE | 5.333 | 0.667 | | 0.681 | 0.087 | |
| | | | | FHE | 2.889 | 0.423 | | | | |
| | | D14 | Long | HE | 4.222 | 0.401 | | 0.600 | 0.175 | |
| | | | | FHE | 2.778 | 0.324 | | | | |
| | | D15 | Long | HE | 4.667 | 0.471 | | 0.754 | 0.038 | * |
| | | | | FHE | 3.333 | 0.441 | | | | |
| | | D16 | Long | HE | 4.667 | 0.471 | | 0.258 | 1.000 | |
| | | | | FHE | 3.222 | 0.364 | | | | |
| Table S3 | Figure S3B | D0 | Srhot | HE | 1.111 | 0.200 | Spearman's rank correlation coefficient | -0.132 | 1.000 | |
| | | | | FHE | 1.444 | 0.862 | | | | |
| | | D1 | Srhot | HE | 1.889 | 0.588 | | 0.536 | 0.273 | |
| | | | | FHE | 1.333 | 0.236 | | | | |
| | | D2 | Srhot | HE | 2.111 | 0.309 | | 0.729 | 0.052 | |
| | | | | FHE | 1.556 | 0.242 | | | | |
| | | D3 | Srhot | HE | 2.556 | 0.444 | | 0.102 | 1.000 | |
| | | | | FHE | 1.667 | 0.289 | | | | |
| | | D4 | Srhot | HE | 2.889 | 0.588 | | 0.974 | 0.000 | *** |
| | | | | FHE | 2.444 | 0.444 | | | | |
| | | D5 | Srhot | HE | 2.667 | 0.236 | | 0.220 | 1.000 | |
| | | | | FHE | 2.111 | 0.261 | | | | |
| | | D6 | Srhot | HE | 3.444 | 0.530 | | 0.759 | 0.035 | * |
| | | | | FHE | 2.667 | 0.289 | | | | |
| | | D7 | Srhot | HE | 3.556 | 0.444 | | 0.862 | 0.006 | ** |
| | | | | FHE | 2.889 | 0.423 | | | | |
| | | D8 | Srhot | HE | 4.667 | 0.624 | | 0.522 | 0.299 | |
| | | | | FHE | 3.111 | 0.484 | | | | |
| | | D9 | Srhot | HE | 3.667 | 0.624 | | 0.466 | 0.411 | |
| | | | | FHE | 2.556 | 0.294 | | | | |
| | | D10 | Srhot | HE | 3.222 | 0.278 | | 0.787 | 0.024 | * |
| | | | | FHE | 2.778 | 0.222 | | | | |
| | | D11 | Srhot | HE | 3.778 | 0.364 | | 0.696 | 0.074 | |
| | | | | FHE | 2.778 | 0.465 | | | | |
| | | D12 | Srhot | HE | 3.778 | 0.547 | | 0.798 | 0.020 | * |
| | | | | FHE | 3.000 | 0.601 | | | | |
| | | D13 | Srhot | HE | 3.556 | 0.475 | | 0.964 | 0.000 | *** |
| | | | | FHE | 3.333 | 0.471 | | | | |
| | | D14 | Srhot | HE | 4.222 | 0.547 | | 0.781 | 0.026 | * |
| | | | | FHE | 3.444 | 0.475 | | | | |
| | | D15 | Srhot | HE | 3.444 | 0.648 | | 0.929 | 0.001 | ** |
| | | | | FHE | 3.000 | 0.373 | | | | |
| | | D16 | Srhot | HE | 3.444 | 0.556 | | 0.987 | 0.000 | *** |
| | | | | FHE | 3.111 | 0.423 | | | | |

| Correlation between HE and FHE peak frequencies (WT) | | | | | | | | | | |
|---|---|---|---|---|---|---|---|---|---|---|
| Table | Related Figure | Day | Condition | Pair | Mean | SEM | Stastistical test | Spearman's ρ | P | * |
| Table S3 | Figure S3B | D0 | Long | HE | 0.778 | 0.434 | Spearman's rank correlation coefficient | 0.714 | 0.061 | |
| | | | | FHE | 0.670 | 0.960 | | | | |
| | | D1 | Long | HE | 1.000 | 0.289 | | 0.256 | 1.000 | |
| | | | | FHE | 0.778 | 0.222 | | | | |
| | | D2 | Long | HE | 1.667 | 0.471 | | 0.694 | 0.076 | |
| | | | | FHE | 1.333 | 0.373 | | | | |
| | | D3 | Long | HE | 2.111 | 0.351 | | 0.682 | 0.086 | |
| | | | | FHE | 0.889 | 0.261 | | | | |
| | | D4 | Long | HE | 2.444 | 0.294 | | 0.768 | 0.031 | * |
| | | | | FHE | 1.889 | 0.309 | | | | |
| | | D5 | Long | HE | 3.000 | 0.527 | | 0.498 | 0.345 | |
| | | | | FHE | 2.111 | 0.351 | | | | |
| | | D6 | Long | HE | 3.222 | 0.278 | | 0.588 | 0.191 | |
| | | | | FHE | 2.444 | 0.338 | | | | |
| | | D7 | Long | HE | 3.333 | 0.408 | | 0.643 | 0.124 | |
| | | | | FHE | 2.333 | 0.333 | | | | |
| | | D8 | Long | HE | 3.556 | 0.338 | | -0.371 | 0.651 | |
| | | | | FHE | 2.222 | 0.147 | | | | |
| | | D9 | Long | HE | 4.222 | 0.662 | | 0.666 | 0.101 | |
| | | | | FHE | 2.889 | 0.564 | | | | |
| | | D10 | Long | HE | 3.889 | 0.261 | | 0.463 | 0.419 | |
| | | | | FHE | 2.556 | 0.176 | | | | |
| | | D11 | Long | HE | 4.444 | 0.556 | | 0.283 | 0.920 | |
| | | | | FHE | 3.000 | 0.167 | | | | |
| | | D12 | Long | HE | 4.667 | 0.577 | | 0.461 | 0.424 | |
| | | | | FHE | 2.667 | 0.373 | | | | |
| | | D13 | Long | HE | 4.333 | 0.373 | | 0.469 | 0.405 | |
| | | | | FHE | 2.889 | 0.261 | | | | |
| | | D14 | Long | HE | 4.333 | 0.408 | | 0.240 | 1.000 | |
| | | | | FHE | 2.778 | 0.222 | | | | |
| | | D15 | Long | HE | 4.444 | 0.377 | | 0.340 | 0.742 | |
| | | | | FHE | 2.889 | 0.309 | | | | |
| | | D16 | Long | HE | 5.000 | 0.408 | | -0.179 | 1.000 | |
| | | | | FHE | 3.111 | 0.200 | | | | |
| Table S3 | Figure S3B | D0 | Short | HE | 1.111 | 0.351 | Spearman's rank correlation coefficient | 0.297 | 0.877 | |
| | | | | FHE | 0.667 | 1.226 | | | | |
| | | D1 | Short | HE | 1.000 | 0.236 | | 0.887 | 0.003 | ** |
| | | | | FHE | 1.000 | 0.289 | | | | |
| | | D2 | Short | HE | 1.222 | 0.222 | | 0.505 | 0.331 | |
| | | | | FHE | 1.000 | 0.236 | | | | |
| | | D3 | Short | HE | 2.667 | 0.500 | | 0.838 | 0.010 | * |
| | | | | FHE | 1.556 | 0.338 | | | | |
| | | D4 | Short | HE | 2.889 | 0.564 | | 0.378 | 0.632 | |
| | | | | FHE | 2.000 | 0.236 | | | | |
| | | D5 | Short | HE | 3.222 | 0.547 | | 0.513 | 0.316 | |
| | | | | FHE | 2.333 | 0.167 | | | | |
| | | D6 | Short | HE | 2.333 | 0.373 | | 0.910 | 0.001 | ** |
| | | | | FHE | 1.889 | 0.261 | | | | |
| | | D7 | Short | HE | 3.111 | 0.351 | | 0.849 | 0.008 | ** |
| | | | | FHE | 2.000 | 0.289 | | | | |
| | | D8 | Short | HE | 3.000 | 0.289 | | 0.535 | 0.275 | |
| | | | | FHE | 2.111 | 0.261 | | | | |
| | | D9 | Short | HE | 3.444 | 0.503 | | 0.412 | 0.540 | |
| | | | | FHE | 2.444 | 0.242 | | | | |
| | | D10 | Short | HE | 3.333 | 0.289 | | 0.400 | 0.572 | |
| | | | | FHE | 2.556 | 0.294 | | | | |
| | | D11 | Short | HE | 3.333 | 0.471 | | 0.811 | 0.016 | * |
| | | | | FHE | 2.778 | 0.278 | | | | |
| | | D12 | Short | HE | 3.778 | 0.364 | | -0.318 | 0.809 | |
| | | | | FHE | 2.778 | 0.222 | | | | |
| | | D13 | Short | HE | 2.444 | 0.176 | | 0.598 | 0.178 | |
| | | | | FHE | 2.222 | 0.147 | | | | |
| | | D14 | Short | HE | 3.111 | 0.261 | | 0.611 | 0.160 | |
| | | | | FHE | 2.444 | 0.294 | | | | |
| | | D15 | Short | HE | 3.444 | 0.444 | | 0.734 | 0.049 | * |
| | | | | FHE | 2.667 | 0.471 | | | | |
| | | D16 | Short | HE | 2.889 | 0.200 | | 0.551 | 0.248 | |
| | | | | FHE | 2.444 | 0.294 | | | | |

## 3_8

| Correlation between HE and FHE peak latencies (KO) | | | | | | | | | | |
|---|---|---|---|---|---|---|---|---|---|---|
| Table | Related Figure | Day | Condition | Paiir | Mean | SEM | Stastistical test | Spearman's ρ | P | * |
| Table S3 | Figure S3B | D0 | Long | HE | 4.000 | 1.093 | Spearman's rank correlation coefficient | 0.953 | 0.000 | *** |
| | | | | FHE | 3.556 | 1.029 | | | | |
| | | D1 | Long | HE | 1.000 | 0.408 | | 1.000 | 0.000 | *** |
| | | | | FHE | 1.000 | 0.408 | | | | |
| | | D2 | Long | HE | 4.222 | 1.310 | | 0.843 | 0.009 | ** |
| | | | | FHE | 2.667 | 1.027 | | | | |
| | | D3 | Long | HE | 3.667 | 1.291 | | 0.734 | 0.049 | * |
| | | | | FHE | 2.556 | 0.835 | | | | |
| | | D4 | Long | HE | 2.889 | 1.296 | | 0.148 | 1.000 | |
| | | | | FHE | 2.444 | 0.930 | | | | |
| | | D5 | Long | HE | 4.667 | 1.202 | | -0.090 | 1.000 | |
| | | | | FHE | 2.778 | 1.051 | | | | |
| | | D6 | Long | HE | 4.222 | 1.164 | | 0.147 | 1.000 | |
| | | | | FHE | 3.111 | 1.006 | | | | |
| | | D7 | Long | HE | 3.556 | 1.215 | | 0.056 | 1.000 | |
| | | | | FHE | 2.000 | 0.667 | | | | |
| | | D8 | Long | HE | 3.222 | 1.382 | | 0.798 | 0.020 | * |
| | | | | FHE | 1.000 | 0.373 | | | | |
| | | D9 | Long | HE | 5.000 | 1.333 | | -0.087 | 1.000 | |
| | | | | FHE | 1.889 | 0.423 | | | | |
| | | D10 | Long | HE | 5.111 | 1.172 | | -0.364 | 0.671 | |
| | | | | FHE | 2.444 | 0.412 | | | | |
| | | D11 | Long | HE | 5.556 | 1.600 | | 0.194 | 1.000 | |
| | | | | FHE | 2.444 | 0.709 | | | | |
| | | D12 | Long | HE | 3.111 | 1.252 | | -0.189 | 1.000 | |
| | | | | FHE | 1.333 | 0.167 | | | | |
| | | D13 | Long | HE | 5.111 | 1.348 | | 0.252 | 1.000 | |
| | | | | FHE | 2.000 | 0.782 | | | | |
| | | D14 | Long | HE | 3.889 | 0.889 | | 0.537 | 0.273 | |
| | | | | FHE | 2.556 | 0.801 | | | | |
| | | D15 | Long | HE | 5.222 | 1.579 | | 0.186 | 1.000 | |
| | | | | FHE | 2.000 | 0.527 | | | | |
| | | D16 | Long | HE | 5.000 | 1.414 | | 0.698 | 0.073 | |
| | | | | FHE | 2.444 | 0.689 | | | | |
| Table S3 | Figure S3B | D0 | Srhot | HE | 3.667 | 1.312 | Spearman's rank correlation coefficient | 0.200 | 1.000 | |
| | | | | FHE | 4.778 | 0.862 | | | | |
| | | D1 | Srhot | HE | 3.333 | 1.312 | | 0.485 | 0.372 | |
| | | | | FHE | 2.333 | 0.913 | | | | |
| | | D2 | Srhot | HE | 6.222 | 1.289 | | -0.127 | 1.000 | |
| | | | | FHE | 2.667 | 0.913 | | | | |
| | | D3 | Srhot | HE | 5.667 | 1.607 | | 0.357 | 0.690 | |
| | | | | FHE | 3.222 | 1.038 | | | | |
| | | D4 | Srhot | HE | 3.444 | 1.271 | | 0.569 | 0.220 | |
| | | | | FHE | 3.444 | 1.056 | | | | |
| | | D5 | Srhot | HE | 2.667 | 0.645 | | 0.870 | 0.005 | ** |
| | | | | FHE | 2.333 | 0.707 | | | | |
| | | D6 | Srhot | HE | 2.778 | 0.741 | | 0.996 | 0.000 | *** |
| | | | | FHE | 2.889 | 0.696 | | | | |
| | | D7 | Srhot | HE | 2.778 | 1.392 | | 0.089 | 1.000 | |
| | | | | FHE | 1.889 | 0.512 | | | | |
| | | D8 | Srhot | HE | 3.333 | 1.067 | | 0.764 | 0.033 | * |
| | | | | FHE | 1.889 | 0.389 | | | | |
| | | D9 | Srhot | HE | 2.556 | 0.801 | | 0.244 | 1.000 | |
| | | | | FHE | 1.111 | 0.309 | | | | |
| | | D10 | Srhot | HE | 2.222 | 0.683 | | 0.450 | 0.448 | |
| | | | | FHE | 2.000 | 0.553 | | | | |
| | | D11 | Srhot | HE | 4.000 | 1.190 | | -0.181 | 1.000 | |
| | | | | FHE | 1.556 | 0.503 | | | | |
| | | D12 | Srhot | HE | 2.111 | 0.772 | | 0.418 | 0.527 | |
| | | | | FHE | 1.111 | 0.261 | | | | |
| | | D13 | Srhot | HE | 2.556 | 0.899 | | 0.969 | 0.000 | *** |
| | | | | FHE | 2.111 | 0.423 | | | | |
| | | D14 | Srhot | HE | 1.667 | 0.408 | | 0.474 | 0.395 | |
| | | | | FHE | 1.222 | 0.364 | | | | |
| | | D15 | Srhot | HE | 1.333 | 0.577 | | 0.932 | 0.000 | *** |
| | | | | FHE | 1.444 | 0.556 | | | | |
| | | D16 | Srhot | HE | 1.444 | 0.377 | | 0.160 | 1.000 | |
| | | | | FHE | 1.556 | 0.294 | | | | |

| Correlation between HE and FHE peak latencies (WT) | | | | | | | | | | |
|---|---|---|---|---|---|---|---|---|---|---|
| Table | Related Figure | Day | Condition | Pair | Mean | SEM | Stastistical test | Spearman's ρ | P | * |
| Table S3 | Figure S3B | D0 | Long | HE | 1.778 | 1.222 | Spearman's rank correlation coefficient | 1.000 | 0.000 | *** |
| | | | | FHE | 1.444 | 0.959 | | | | |
| | | D1 | Long | HE | 2.000 | 1.443 | | 0.033 | 1.000 | |
| | | | | FHE | 1.667 | 0.764 | | | | |
| | | D2 | Long | HE | 3.111 | 1.438 | | 0.948 | 0.000 | *** |
| | | | | FHE | 3.111 | 1.274 | | | | |
| | | D3 | Long | HE | 5.111 | 1.728 | | 0.037 | 1.000 | |
| | | | | FHE | 1.556 | 0.747 | | | | |
| | | D4 | Long | HE | 2.111 | 1.111 | | 0.520 | 0.303 | |
| | | | | FHE | 1.222 | 0.434 | | | | |
| | | D5 | Long | HE | 5.444 | 1.608 | | 0.457 | 0.433 | |
| | | | | FHE | 3.000 | 0.782 | | | | |
| | | D6 | Long | HE | 5.111 | 1.476 | | 0.822 | 0.013 | * |
| | | | | FHE | 3.222 | 0.969 | | | | |
| | | D7 | Long | HE | 4.222 | 0.572 | | 0.759 | 0.036 | * |
| | | | | FHE | 3.667 | 0.972 | | | | |
| | | D8 | Long | HE | 3.222 | 1.278 | | -0.270 | 0.964 | |
| | | | | FHE | 2.444 | 0.626 | | | | |
| | | D9 | Long | HE | 4.000 | 1.179 | | 0.518 | 0.306 | |
| | | | | FHE | 1.556 | 0.412 | | | | |
| | | D10 | Long | HE | 3.111 | 1.060 | | -0.137 | 1.000 | |
| | | | | FHE | 1.222 | 0.364 | | | | |
| | | D11 | Long | HE | 5.556 | 1.365 | | -0.154 | 1.000 | |
| | | | | FHE | 3.556 | 0.915 | | | | |
| | | D12 | Long | HE | 3.778 | 1.245 | | 0.476 | 0.391 | |
| | | | | FHE | 1.444 | 0.377 | | | | |
| | | D13 | Long | HE | 2.889 | 0.633 | | 0.803 | 0.018 | * |
| | | | | FHE | 2.333 | 0.577 | | | | |
| | | D14 | Long | HE | 4.333 | 0.833 | | 0.339 | 0.744 | |
| | | | | FHE | 2.333 | 0.799 | | | | |
| | | D15 | Long | HE | 4.333 | 0.667 | | 0.673 | 0.094 | |
| | | | | FHE | 3.333 | 0.957 | | | | |
| | | D16 | Long | HE | 5.778 | 1.103 | | 0.456 | 0.435 | |
| | | | | FHE | 3.778 | 0.760 | | | | |
| Table S3 | Figure S3B | D0 | Short | HE | 4.000 | 1.616 | Spearman's rank correlation coefficient | 0.045 | 1.000 | |
| | | | | FHE | 3.556 | 1.226 | | | | |
| | | D1 | Short | HE | 4.333 | 1.258 | | 0.271 | 0.962 | |
| | | | | FHE | 3.889 | 0.807 | | | | |
| | | D2 | Short | HE | 1.889 | 1.020 | | 0.354 | 0.699 | |
| | | | | FHE | 1.889 | 0.676 | | | | |
| | | D3 | Short | HE | 3.556 | 1.692 | | 0.101 | 1.000 | |
| | | | | FHE | 2.556 | 0.766 | | | | |
| | | D4 | Short | HE | 2.333 | 1.202 | | 0.019 | 1.000 | |
| | | | | FHE | 0.667 | 0.236 | | | | |
| | | D5 | Short | HE | 2.667 | 0.816 | | 0.521 | 0.300 | |
| | | | | FHE | 3.667 | 0.882 | | | | |
| | | D6 | Short | HE | 1.778 | 0.969 | | 0.692 | 0.078 | |
| | | | | FHE | 2.889 | 0.904 | | | | |
| | | D7 | Short | HE | 4.111 | 1.567 | | 0.591 | 0.188 | |
| | | | | FHE | 1.333 | 0.441 | | | | |
| | | D8 | Short | HE | 4.444 | 1.271 | | -0.090 | 1.000 | |
| | | | | FHE | 2.222 | 0.894 | | | | |
| | | D9 | Short | HE | 3.889 | 0.484 | | 0.372 | 0.649 | |
| | | | | FHE | 3.000 | 0.471 | | | | |
| | | D10 | Short | HE | 4.000 | 1.404 | | -0.064 | 1.000 | |
| | | | | FHE | 2.000 | 0.601 | | | | |
| | | D11 | Short | HE | 3.889 | 1.047 | | 0.508 | 0.326 | |
| | | | | FHE | 2.333 | 0.764 | | | | |
| | | D12 | Short | HE | 3.222 | 0.862 | | 0.487 | 0.367 | |
| | | | | FHE | 2.111 | 0.588 | | | | |
| | | D13 | Short | HE | 2.222 | 0.521 | | 0.849 | 0.008 | ** |
| | | | | FHE | 2.333 | 0.441 | | | | |
| | | D14 | Short | HE | 3.778 | 1.064 | | 0.228 | 1.000 | |
| | | | | FHE | 1.778 | 0.547 | | | | |
| | | D15 | Short | HE | 3.333 | 0.726 | | 0.461 | 0.423 | |
| | | | | FHE | 2.222 | 0.795 | | | | |
| | | D16 | Short | HE | 2.000 | 0.471 | | 0.678 | 0.089 | * |
| | | | | FHE | 1.778 | 0.465 | | | | |

3_9

| [illegible] | | | | | | | | | | |
|---|---|---|---|---|---|---|---|---|---|---|
| Table | Related Figure | Day | Group | Condition | Mean | SEM | Stastistical test | Spearman’s ρ | P | * |
| Table S3 | Figure S3C | D0 | KO | Long | 4.000 | 1.093 | Spearman's rank correlation coefficient | -0.434 | 0.244 | |
| | | | | Short | 3.667 | 1.312 | | | | |
| | | D1 | KO | Long | 1.000 | 0.408 | | -0.378 | 0.316 | |
| | | | | Short | 3.333 | 1.312 | | | | |
| | | D2 | KO | Long | 4.222 | 1.310 | | -0.352 | 0.353 | |
| | | | | Short | 6.222 | 1.289 | | | | |
| | | D3 | KO | Long | 3.667 | 1.291 | | -0.325 | 0.394 | |
| | | | | Short | 5.667 | 1.607 | | | | |
| | | D4 | KO | Long | 2.889 | 1.296 | | 0.116 | 0.767 | |
| | | | | Short | 3.444 | 1.271 | | | | |
| | | D5 | KO | Long | 4.667 | 1.202 | | 0.269 | 0.484 | |
| | | | | Short | 2.667 | 0.645 | | | | |
| | | D6 | KO | Long | 4.222 | 1.164 | | 0.622 | 0.074 | |
| | | | | Short | 2.778 | 0.741 | | | | |
| | | D7 | KO | Long | 3.556 | 1.215 | | 0.375 | 0.320 | |
| | | | | Short | 2.778 | 1.392 | | | | |
| | | D8 | KO | Long | 3.222 | 1.382 | | 0.391 | 0.298 | |
| | | | | Short | 3.333 | 1.067 | | | | |
| | | D9 | KO | Long | 5.000 | 1.333 | | 0.293 | 0.444 | |
| | | | | Long | 2.556 | 0.801 | | | | |
| | | D10 | KO | Short | 5.111 | 1.172 | | -0.004 | 0.991 | |
| | | | | Long | 2.222 | 0.683 | | | | |
| | | D11 | KO | Short | 5.556 | 1.600 | | -0.791 | 0.011 | * |
| | | | | Long | 4.000 | 1.190 | | | | |
| | | D12 | KO | Short | 3.111 | 1.252 | | -0.049 | 0.900 | |
| | | | | Long | 2.111 | 0.772 | | | | |
| | | D13 | KO | Short | 5.111 | 1.348 | | 0.617 | 0.076 | |
| | | | | Long | 2.556 | 0.899 | | | | |
| | | D14 | KO | Short | 3.889 | 0.889 | | 0.347 | 0.360 | |
| | | | | Long | 1.667 | 0.408 | | | | |
| | | D15 | KO | Short | 5.222 | 1.579 | | 0.829 | 0.006 | ** |
| | | | | Long | 1.333 | 0.577 | | | | |
| | | D16 | KO | Short | 5.000 | 1.414 | | 0.471 | 0.201 | |
| | | | | Long | 1.444 | 0.377 | | | | |
| Table S3 | Figure S3C | D0 | WT | Long | 1.778 | 1.222 | Spearman's rank correlation coefficient | -0.060 | 0.878 | |
| | | | | Short | 4.000 | 1.616 | | | | |
| | | D1 | WT | Long | 2.000 | 1.443 | | -0.060 | 0.878 | |
| | | | | Short | 4.333 | 1.258 | | | | |
| | | D2 | WT | Long | 3.111 | 1.438 | | -0.510 | 0.160 | |
| | | | | Short | 1.889 | 1.020 | | | | |
| | | D3 | WT | Long | 5.111 | 1.728 | | 0.432 | 0.246 | |
| | | | | Short | 3.556 | 1.692 | | | | |
| | | D4 | WT | Long | 2.111 | 1.111 | | 0.303 | 0.428 | |
| | | | | Short | 2.333 | 1.202 | | | | |
| | | D5 | WT | Long | 5.444 | 1.608 | | 0.253 | 0.511 | |
| | | | | Short | 2.667 | 0.816 | | | | |
| | | D6 | WT | Long | 5.111 | 1.476 | | 0.147 | 0.707 | |
| | | | | Short | 1.778 | 0.969 | | | | |
| | | D7 | WT | Long | 4.222 | 0.572 | | 0.025 | 0.948 | |
| | | | | Short | 4.111 | 1.567 | | | | |
| | | D8 | WT | Long | 3.222 | 1.278 | | 0.385 | 0.306 | |
| | | | | Short | 4.444 | 1.271 | | | | |
| | | D9 | WT | Long | 4.000 | 1.179 | | 0.278 | 0.469 | |
| | | | | Long | 3.889 | 0.484 | | | | |
| | | D10 | WT | Short | 3.111 | 1.060 | | 0.356 | 0.348 | |
| | | | | Long | 4.000 | 1.404 | | | | |
| | | D11 | WT | Short | 5.556 | 1.365 | | 0.026 | 0.947 | |
| | | | | Long | 3.889 | 1.047 | | | | |
| | | D12 | WT | Short | 3.778 | 1.245 | | -0.469 | 0.203 | |
| | | | | Long | 3.222 | 0.862 | | | | |
| | | D13 | WT | Short | 2.889 | 0.633 | | 0.444 | 0.232 | |
| | | | | Long | 2.222 | 0.521 | | | | |
| | | D14 | WT | Short | 4.333 | 0.833 | | 0.562 | 0.115 | |
| | | | | Long | 3.778 | 1.064 | | | | |
| | | D15 | WT | Short | 4.333 | 0.667 | | 0.376 | 0.319 | |
| | | | | Long | 3.333 | 0.726 | | | | |
| | | D16 | WT | Short | 5.778 | 1.103 | | 0.602 | 0.086 | |
| | | | | Long | 2.000 | 0.471 | | | | |

## 4_1

| Table | Related Figure | Day | Condition | Group | Mean | SEM | Stastistical test | Z | P | * |
|---|---|---|---|---|---|---|---|---|---|---|
| HE frequency in 0–2 s window | | | | | | | | | | |
| Table S4 | Figure S4 | D0 | Long | KO | 0.472 | 0.329 | Mann-Whitney U Test | -0.369 | 0.796 | |
| | | | | WT | 0.139 | 0.073 | | | | |
| | | D1 | Long | KO | 0.278 | 0.153 | | -0.158 | 0.931 | |
| | | | | WT | 0.222 | 0.121 | | | | |
| | | D2 | Long | KO | 0.417 | 0.144 | | -0.093 | 0.931 | |
| | | | | WT | 0.722 | 0.385 | | | | |
| | | D3 | Long | KO | 0.889 | 0.470 | | -0.094 | 0.931 | |
| | | | | WT | 0.778 | 0.426 | | | | |
| | | D4 | Long | KO | 1.222 | 0.480 | | -0.585 | 0.605 | |
| | | | | WT | 1.500 | 0.514 | | | | |
| | | D5 | Long | KO | 1.333 | 0.520 | | -0.045 | 1.000 | |
| | | | | WT | 1.556 | 0.586 | | | | |
| | | D6 | Long | KO | 1.889 | 0.455 | | -0.982 | 0.340 | |
| | | | | WT | 1.389 | 0.582 | | | | |
| | | D7 | Long | KO | 2.389 | 0.881 | | -0.358 | 0.730 | |
| | | | | WT | 1.500 | 0.433 | | | | |
| | | D8 | Long | KO | 2.944 | 0.843 | | -0.133 | 0.931 | |
| | | | | WT | 2.333 | 0.520 | | | | |
| | | D9 | Long | KO | 3.111 | 0.666 | | -1.113 | 0.297 | |
| | | | | WT | 2.056 | 0.530 | | | | |
| | | D10 | Long | KO | 2.222 | 0.683 | | -0.224 | 0.863 | |
| | | | | WT | 2.056 | 0.523 | | | | |
| | | D11 | Long | KO | 2.611 | 0.865 | | -0.045 | 1.000 | |
| | | | | WT | 2.111 | 0.594 | | | | |
| | | D12 | Long | KO | 2.667 | 0.791 | | -0.312 | 0.796 | |
| | | | | WT | 2.167 | 0.672 | | | | |
| | | D13 | Long | KO | 3.222 | 1.087 | | -0.761 | 0.489 | |
| | | | | WT | 1.833 | 0.577 | | | | |
| | | D14 | Long | KO | 2.944 | 0.684 | | -0.844 | 0.436 | |
| | | | | WT | 2.167 | 0.707 | | | | |
| | | D15 | Long | KO | 2.833 | 0.874 | | -0.671 | 0.546 | |
| | | | | WT | 2.111 | 0.790 | | | | |
| | | D16 | Long | KO | 2.444 | 0.621 | | -0.935 | 0.387 | |
| | | | | WT | 1.722 | 0.619 | | | | |
| Table S4 | Figure S4 | D0 | Short | KO | 0.222 | 0.114 | Mann-Whitney U Test | -1.455 | 0.258 | |
| | | | | WT | 0.056 | 0.056 | | | | |
| | | D1 | Short | KO | 0.528 | 0.193 | | -2.842 | 0.014 | * |
| | | | | WT | 0.000 | 0.000 | | | | |
| | | D2 | Short | KO | 0.667 | 0.161 | | -1.230 | 0.258 | |
| | | | | WT | 0.389 | 0.132 | | | | |
| | | D3 | Short | KO | 0.944 | 0.269 | | -0.182 | 0.863 | |
| | | | | WT | 0.889 | 0.247 | | | | |
| | | D4 | Short | KO | 1.667 | 0.486 | | -0.359 | 0.730 | |
| | | | | WT | 1.278 | 0.222 | | | | |
| | | D5 | Short | KO | 1.611 | 0.545 | | -0.274 | 0.796 | |
| | | | | WT | 1.111 | 0.274 | | | | |
| | | D6 | Short | KO | 1.778 | 0.596 | | -0.787 | 0.489 | |
| | | | | WT | 1.222 | 0.528 | | | | |
| | | D7 | Short | KO | 2.111 | 0.588 | | -0.871 | 0.436 | |
| | | | | WT | 1.333 | 0.354 | | | | |
| | | D8 | Short | KO | 2.500 | 0.565 | | -1.250 | 0.222 | |
| | | | | WT | 1.500 | 0.289 | | | | |
| | | D9 | Short | KO | 2.444 | 0.444 | | -1.958 | 0.050 | |
| | | | | WT | 1.222 | 0.364 | | | | |
| | | D10 | Short | KO | 2.167 | 0.456 | | -0.802 | 0.436 | |
| | | | | WT | 1.722 | 0.560 | | | | |
| | | D11 | Short | KO | 2.333 | 0.514 | | -0.445 | 0.666 | |
| | | | | WT | 2.111 | 0.633 | | | | |
| | | D12 | Short | KO | 2.278 | 0.547 | | -0.934 | 0.387 | |
| | | | | WT | 1.500 | 0.354 | | | | |
| | | D13 | Short | KO | 2.167 | 0.777 | | -0.356 | 0.730 | |
| | | | | WT | 1.333 | 0.363 | | | | |
| | | D14 | Short | KO | 2.778 | 0.678 | | -1.332 | 0.190 | |
| | | | | WT | 1.667 | 0.408 | | | | |
| | | D15 | Short | KO | 2.556 | 0.679 | | -1.341 | 0.190 | |
| | | | | WT | 1.111 | 0.351 | | | | |
| | | D16 | Short | KO | 2.500 | 0.651 | | -0.980 | 0.340 | |
| | | | | WT | 1.611 | 0.341 | | | | |

| Table | Related Figure | Day | Group | Condition | Mean | SEM | Stastistical test | Z | P | * |
|---|---|---|---|---|---|---|---|---|---|---|
| HE frequency in 0–2 s window | | | | | | | | | | |
| Table S4 | Figure S4 | D0 | KO | Long | 0.472 | 0.329 | Wilcoxon Signed Ranks Test | -0.184 | 0.854 | |
| | | | | Short | 0.222 | 0.114 | | | | |
| | | D1 | KO | Long | 0.278 | 0.153 | | -1.510 | 0.131 | |
| | | | | Short | 0.528 | 0.193 | | | | |
| | | D2 | KO | Long | 0.417 | 0.144 | | -1.579 | 0.114 | |
| | | | | Short | 0.667 | 0.161 | | | | |
| | | D3 | KO | Long | 0.889 | 0.470 | | -0.170 | 0.865 | |
| | | | | Short | 0.944 | 0.269 | | | | |
| | | D4 | KO | Long | 1.222 | 0.480 | | -0.985 | 0.325 | |
| | | | | Short | 1.667 | 0.486 | | | | |
| | | D5 | KO | Long | 1.333 | 0.520 | | -0.632 | 0.527 | |
| | | | | Short | 1.611 | 0.545 | | | | |
| | | D6 | KO | Long | 1.889 | 0.455 | | -0.060 | 0.952 | |
| | | | | Short | 1.778 | 0.596 | | | | |
| | | D7 | KO | Long | 2.389 | 0.881 | | -0.680 | 0.497 | |
| | | | | Short | 2.111 | 0.588 | | | | |
| | | D8 | KO | Long | 2.944 | 0.843 | | -0.517 | 0.605 | |
| | | | | Short | 2.500 | 0.565 | | | | |
| | | D9 | KO | Long | 3.111 | 0.666 | | -1.131 | 0.258 | |
| | | | | Long | 2.444 | 0.444 | | | | |
| | | D10 | KO | Short | 2.222 | 0.683 | | -0.071 | 0.943 | |
| | | | | Long | 2.167 | 0.456 | | | | |
| | | D11 | KO | Short | 2.611 | 0.865 | | -0.141 | 0.888 | |
| | | | | Long | 2.333 | 0.514 | | | | |
| | | D12 | KO | Short | 2.667 | 0.791 | | -0.938 | 0.348 | |
| | | | | Long | 2.278 | 0.547 | | | | |
| | | D13 | KO | Short | 3.222 | 1.087 | | -2.120 | 0.034 | * |
| | | | | Long | 2.167 | 0.777 | | | | |
| | | D14 | KO | Short | 2.944 | 0.684 | | -0.531 | 0.595 | |
| | | | | Long | 2.778 | 0.678 | | | | |
| | | D15 | KO | Short | 2.833 | 0.874 | | -0.085 | 0.932 | |
| | | | | Long | 2.556 | 0.679 | | | | |
| | | D16 | KO | Short | 2.444 | 0.621 | | -0.106 | 0.915 | |
| | | | | Long | 2.500 | 0.651 | | | | |
| Table S4 | Figure S4 | D0 | WT | Long | 0.139 | 0.073 | Wilcoxon Signed Ranks Test | -1.342 | 0.180 | |
| | | | | Short | 0.056 | 0.056 | | | | |
| | | D1 | WT | Long | 0.222 | 0.121 | | -1.633 | 0.102 | |
| | | | | Short | 0.000 | 0.000 | | | | |
| | | D2 | WT | Long | 0.722 | 0.385 | | -0.341 | 0.733 | |
| | | | | Short | 0.389 | 0.132 | | | | |
| | | D3 | WT | Long | 0.778 | 0.426 | | -0.970 | 0.332 | |
| | | | | Short | 0.889 | 0.247 | | | | |
| | | D4 | WT | Long | 1.500 | 0.514 | | -0.512 | 0.609 | |
| | | | | Short | 1.278 | 0.222 | | | | |
| | | D5 | WT | Long | 1.556 | 0.586 | | -0.923 | 0.356 | |
| | | | | Short | 1.111 | 0.274 | | | | |
| | | D6 | WT | Long | 1.389 | 0.582 | | -0.351 | 0.725 | |
| | | | | Short | 1.222 | 0.528 | | | | |
| | | D7 | WT | Long | 1.500 | 0.433 | | -0.849 | 0.396 | |
| | | | | Short | 1.333 | 0.354 | | | | |
| | | D8 | WT | Long | 2.333 | 0.520 | | -1.620 | 0.105 | |
| | | | | Short | 1.500 | 0.289 | | | | |
| | | D9 | WT | Long | 2.056 | 0.530 | | -2.132 | 0.033 | * |
| | | | | Long | 1.222 | 0.364 | | | | |
| | | D10 | WT | Short | 2.056 | 0.523 | | -1.289 | 0.197 | |
| | | | | Long | 1.722 | 0.560 | | | | |
| | | D11 | WT | Short | 2.111 | 0.594 | | 0.000 | 1.000 | |
| | | | | Long | 2.111 | 0.633 | | | | |
| | | D12 | WT | Short | 2.167 | 0.672 | | -1.527 | 0.127 | |
| | | | | Long | 1.500 | 0.354 | | | | |
| | | D13 | WT | Short | 1.833 | 0.577 | | -1.279 | 0.201 | |
| | | | | Long | 1.333 | 0.363 | | | | |
| | | D14 | WT | Short | 2.167 | 0.707 | | -0.319 | 0.750 | |
| | | | | Long | 1.667 | 0.408 | | | | |
| | | D15 | WT | Short | 2.111 | 0.790 | | -1.614 | 0.106 | |
| | | | | Long | 1.111 | 0.351 | | | | |
| | | D16 | WT | Short | 1.722 | 0.619 | | -0.085 | 0.932 | |
| | | | | Long | 1.611 | 0.341 | | | | |

Abbreviations: HE, head entry; D, training day; KO, CRBN KO mice; WT, wild-type mice.

*, *p < 0.05, **p < 0.01, ***p < 0.001.

## 4_2

| Table | Related Figure | Day | Condition | Group | Mean | SEM | Stastistical test | Z | P | * |
|---|---|---|---|---|---|---|---|---|---|---|
| HE frequency in 0–10 s window | | | | | | | | | | |
| Table S4 | Figure S4 | D0 | Long | KO | 0.228 | 0.090 | Mann-Whitney U Test | -1.683 | 0.113 | |
| | | | | WT | 0.072 | 0.032 | | | | |
| | | D1 | Long | KO | 0.194 | 0.063 | | -0.135 | 0.931 | |
| | | | | WT | 0.189 | 0.077 | | | | |
| | | D2 | Long | KO | 0.328 | 0.066 | | -0.535 | 0.605 | |
| | | | | WT | 0.300 | 0.110 | | | | |
| | | D3 | Long | KO | 0.544 | 0.161 | | -0.134 | 0.931 | |
| | | | | WT | 0.578 | 0.212 | | | | |
| | | D4 | Long | KO | 0.822 | 0.145 | | -0.975 | 0.340 | |
| | | | | WT | 1.133 | 0.271 | | | | |
| | | D5 | Long | KO | 1.033 | 0.190 | | -0.177 | 0.863 | |
| | | | | WT | 1.144 | 0.351 | | | | |
| | | D6 | Long | KO | 1.333 | 0.157 | | -2.083 | 0.040 | * |
| | | | | WT | 0.811 | 0.174 | | | | |
| | | D7 | Long | KO | 1.356 | 0.263 | | -0.843 | 0.436 | |
| | | | | WT | 1.089 | 0.199 | | | | |
| | | D8 | Long | KO | 2.100 | 0.299 | | -1.864 | 0.063 | |
| | | | | WT | 1.333 | 0.237 | | | | |
| | | D9 | Long | KO | 2.378 | 0.308 | | -2.035 | 0.040 | * |
| | | | | WT | 1.444 | 0.343 | | | | |
| | | D10 | Long | KO | 1.933 | 0.317 | | -0.666 | 0.546 | |
| | | | | WT | 1.467 | 0.315 | | | | |
| | | D11 | Long | KO | 1.833 | 0.326 | | -0.796 | 0.436 | |
| | | | | WT | 1.522 | 0.386 | | | | |
| | | D12 | Long | KO | 1.856 | 0.185 | | -0.798 | 0.436 | |
| | | | | WT | 1.611 | 0.294 | | | | |
| | | D13 | Long | KO | 1.878 | 0.258 | | -1.194 | 0.258 | |
| | | | | WT | 1.411 | 0.249 | | | | |
| | | D14 | Long | KO | 1.811 | 0.201 | | -0.619 | 0.546 | |
| | | | | WT | 1.600 | 0.292 | | | | |
| | | D15 | Long | KO | 1.944 | 0.313 | | -1.694 | 0.094 | |
| | | | | WT | 1.378 | 0.202 | | | | |
| | | D16 | Long | KO | 1.556 | 0.146 | | -0.089 | 0.931 | |
| | | | | WT | 1.433 | 0.244 | | | | |
| Table S4 | Figure S4 | D0 | Short | KO | 0.072 | 0.033 | Mann-Whitney U Test | -0.957 | 0.436 | |
| | | | | WT | 0.067 | 0.044 | | | | |
| | | D1 | Short | KO | 0.117 | 0.053 | | -1.610 | 0.161 | |
| | | | | WT | 0.044 | 0.034 | | | | |
| | | D2 | Short | KO | 0.122 | 0.043 | | -0.725 | 0.489 | |
| | | | | WT | 0.094 | 0.044 | | | | |
| | | D3 | Short | KO | 0.211 | 0.081 | | -0.091 | 0.931 | |
| | | | | WT | 0.333 | 0.157 | | | | |
| | | D4 | Short | KO | 0.289 | 0.073 | | -0.404 | 0.730 | |
| | | | | WT | 0.289 | 0.065 | | | | |
| | | D5 | Short | KO | 0.267 | 0.053 | | -1.066 | 0.340 | |
| | | | | WT | 0.456 | 0.236 | | | | |
| | | D6 | Short | KO | 0.378 | 0.144 | | -1.033 | 0.340 | |
| | | | | WT | 0.256 | 0.106 | | | | |
| | | D7 | Short | KO | 0.444 | 0.145 | | -0.925 | 0.387 | |
| | | | | WT | 0.444 | 0.211 | | | | |
| | | D8 | Short | KO | 0.500 | 0.267 | | -0.448 | 0.666 | |
| | | | | WT | 0.444 | 0.147 | | | | |
| | | D9 | Short | KO | 0.611 | 0.291 | | -1.539 | 0.136 | |
| | | | | WT | 0.367 | 0.163 | | | | |
| | | D10 | Short | KO | 0.478 | 0.149 | | -0.850 | 0.436 | |
| | | | | WT | 0.422 | 0.153 | | | | |
| | | D11 | Short | KO | 0.589 | 0.323 | | -0.401 | 0.730 | |
| | | | | WT | 0.344 | 0.073 | | | | |
| | | D12 | Short | KO | 0.456 | 0.082 | | -0.762 | 0.489 | |
| | | | | WT | 0.389 | 0.118 | | | | |
| | | D13 | Short | KO | 0.467 | 0.113 | | -1.335 | 0.190 | |
| | | | | WT | 0.333 | 0.120 | | | | |
| | | D14 | Short | KO | 0.567 | 0.118 | | -1.377 | 0.190 | |
| | | | | WT | 0.389 | 0.149 | | | | |
| | | D15 | Short | KO | 0.511 | 0.098 | | -1.066 | 0.297 | |
| | | | | WT | 0.378 | 0.136 | | | | |
| | | D16 | Short | KO | 0.522 | 0.120 | | -1.065 | 0.297 | |
| | | | | WT | 0.344 | 0.091 | | | | |

| Table | Related Figure | Day | Group | Condition | Mean | SEM | Stastistical test | Z | P | * |
|---|---|---|---|---|---|---|---|---|---|---|
| HE frequency in 0–10 s window | | | | | | | | | | |
| Table S4 | Figure S4 | D0 | KO | Long | 0.228 | 0.090 | Wilcoxon Signed Ranks Test | -1.616 | 0.106 | |
| | | | | Short | 0.072 | 0.033 | | | | |
| | | D1 | KO | Long | 0.194 | 0.063 | | -0.931 | 0.352 | |
| | | | | Short | 0.117 | 0.053 | | | | |
| | | D2 | KO | Long | 0.328 | 0.066 | | -2.079 | 0.038 | * |
| | | | | Short | 0.122 | 0.043 | | | | |
| | | D3 | KO | Long | 0.544 | 0.161 | | -1.825 | 0.068 | |
| | | | | Short | 0.211 | 0.081 | | | | |
| | | D4 | KO | Long | 0.822 | 0.145 | | -2.547 | 0.011 | * |
| | | | | Short | 0.289 | 0.073 | | | | |
| | | D5 | KO | Long | 1.033 | 0.190 | | -2.680 | 0.007 | ** |
| | | | | Short | 0.267 | 0.053 | | | | |
| | | D6 | KO | Long | 1.333 | 0.157 | | -2.547 | 0.011 | * |
| | | | | Short | 0.378 | 0.144 | | | | |
| | | D7 | KO | Long | 1.356 | 0.263 | | -2.521 | 0.012 | * |
| | | | | Short | 0.444 | 0.145 | | | | |
| | | D8 | KO | Long | 2.100 | 0.299 | | -2.666 | 0.008 | ** |
| | | | | Short | 0.500 | 0.267 | | | | |
| | | D9 | KO | Long | 2.378 | 0.308 | | -2.668 | 0.008 | ** |
| | | | | Long | 0.611 | 0.291 | | | | |
| | | D10 | KO | Short | 1.933 | 0.317 | | -2.666 | 0.008 | ** |
| | | | | Long | 0.478 | 0.149 | | | | |
| | | D11 | KO | Short | 1.833 | 0.326 | | -2.194 | 0.028 | * |
| | | | | Long | 0.589 | 0.323 | | | | |
| | | D12 | KO | Short | 1.856 | 0.185 | | -2.668 | 0.008 | ** |
| | | | | Long | 0.456 | 0.082 | | | | |
| | | D13 | KO | Short | 1.878 | 0.258 | | -2.668 | 0.008 | ** |
| | | | | Long | 0.467 | 0.113 | | | | |
| | | D14 | KO | Short | 1.811 | 0.201 | | -2.670 | 0.008 | ** |
| | | | | Long | 0.567 | 0.118 | | | | |
| | | D15 | KO | Short | 1.944 | 0.313 | | -2.675 | 0.007 | ** |
| | | | | Long | 0.511 | 0.098 | | | | |
| | | D16 | KO | Short | 1.556 | 0.146 | | -2.670 | 0.008 | ** |
| | | | | Long | 0.522 | 0.120 | | | | |
| Table S4 | Figure S4 | D0 | WT | Long | 0.072 | 0.032 | Wilcoxon Signed Ranks Test | -0.368 | 0.713 | |
| | | | | Short | 0.067 | 0.044 | | | | |
| | | D1 | WT | Long | 0.189 | 0.077 | | -2.379 | 0.017 | * |
| | | | | Short | 0.044 | 0.034 | | | | |
| | | D2 | WT | Long | 0.300 | 0.110 | | -1.753 | 0.080 | |
| | | | | Short | 0.094 | 0.044 | | | | |
| | | D3 | WT | Long | 0.578 | 0.212 | | -1.529 | 0.126 | |
| | | | | Short | 0.333 | 0.157 | | | | |
| | | D4 | WT | Long | 1.133 | 0.271 | | -2.670 | 0.008 | ** |
| | | | | Short | 0.289 | 0.065 | | | | |
| | | D5 | WT | Long | 1.144 | 0.351 | | -2.371 | 0.018 | * |
| | | | | Short | 0.456 | 0.236 | | | | |
| | | D6 | WT | Long | 0.811 | 0.174 | | -2.203 | 0.028 | * |
| | | | | Short | 0.256 | 0.106 | | | | |
| | | D7 | WT | Long | 1.089 | 0.199 | | -1.666 | 0.096 | |
| | | | | Short | 0.444 | 0.211 | | | | |
| | | D8 | WT | Long | 1.333 | 0.237 | | -2.561 | 0.010 | * |
| | | | | Short | 0.444 | 0.147 | | | | |
| | | D9 | WT | Long | 1.444 | 0.343 | | -2.439 | 0.015 | * |
| | | | | Long | 0.367 | 0.163 | | | | |
| | | D10 | WT | Short | 1.467 | 0.315 | | -1.904 | 0.057 | |
| | | | | Long | 0.422 | 0.153 | | | | |
| | | D11 | WT | Short | 1.522 | 0.386 | | -2.194 | 0.028 | * |
| | | | | Long | 0.344 | 0.073 | | | | |
| | | D12 | WT | Short | 1.611 | 0.294 | | -2.547 | 0.011 | * |
| | | | | Long | 0.389 | 0.118 | | | | |
| | | D13 | WT | Short | 1.411 | 0.249 | | -2.439 | 0.015 | * |
| | | | | Long | 0.333 | 0.120 | | | | |
| | | D14 | WT | Short | 1.600 | 0.292 | | -2.547 | 0.011 | * |
| | | | | Long | 0.389 | 0.149 | | | | |
| | | D15 | WT | Short | 1.378 | 0.202 | | -2.201 | 0.028 | * |
| | | | | Long | 0.378 | 0.136 | | | | |
| | | D16 | WT | Short | 1.433 | 0.244 | | -2.558 | 0.011 | * |
| | | | | Long | 0.344 | 0.091 | | | | |

5_1

| HE frequency in 0–1 s window | | | | | | | | | | |
|---|---|---|---|---|---|---|---|---|---|---|
| Table | Related Figure | Day | Condition | Group | Mean | SEM | Stastistical test | Z | P | * |
| Table S5 | Figure 1H,I / Figure S7 | D0 | Long | KO | 0.056 | 0.056 | Mann-Whitney U Test | -1.215 | 0.387 | |
| | | | | WT | 0.278 | 0.147 | | | | |
| | | D1 | Long | KO | 0.111 | 0.073 | | -0.243 | 0.863 | |
| | | | | WT | 0.222 | 0.147 | | | | |
| | | D2 | Long | KO | 0.167 | 0.118 | | -0.784 | 0.546 | |
| | | | | WT | 0.778 | 0.434 | | | | |
| | | D3 | Long | KO | 0.556 | 0.338 | | -0.160 | 0.931 | |
| | | | | WT | 0.333 | 0.167 | | | | |
| | | D4 | Long | KO | 0.556 | 0.444 | | -1.927 | 0.077 | |
| | | | | WT | 1.111 | 0.309 | | | | |
| | | D5 | Long | KO | 0.333 | 0.167 | | -0.338 | 0.796 | |
| | | | | WT | 0.333 | 0.236 | | | | |
| | | D6 | Long | KO | 1.222 | 0.434 | | -1.385 | 0.222 | |
| | | | | WT | 0.556 | 0.338 | | | | |
| | | D7 | Long | KO | 0.556 | 0.294 | | -0.318 | 0.796 | |
| | | | | WT | 0.333 | 0.167 | | | | |
| | | D8 | Long | KO | 1.778 | 0.619 | | -0.318 | 0.796 | |
| | | | | WT | 1.333 | 0.408 | | | | |
| | | D9 | Long | KO | 1.222 | 0.521 | | -0.140 | 0.931 | |
| | | | | WT | 1.000 | 0.289 | | | | |
| | | D10 | Long | KO | 0.556 | 0.176 | | -1.175 | 0.297 | |
| | | | | WT | 1.222 | 0.401 | | | | |
| | | D11 | Long | KO | 1.000 | 0.289 | | -0.992 | 0.387 | |
| | | | | WT | 0.778 | 0.434 | | | | |
| | | D12 | Long | KO | 0.333 | 0.236 | | -0.449 | 0.730 | |
| | | | | WT | 0.444 | 0.242 | | | | |
| | | D13 | Long | KO | 1.000 | 0.553 | | -0.239 | 0.863 | |
| | | | | WT | 0.778 | 0.278 | | | | |
| | | D14 | Long | KO | 1.333 | 0.408 | | -1.666 | 0.136 | |
| | | | | WT | 0.444 | 0.242 | | | | |
| | | D15 | Long | KO | 1.000 | 0.333 | | -1.467 | 0.190 | |
| | | | | WT | 0.444 | 0.294 | | | | |
| | | D16 | Long | KO | 1.444 | 0.580 | | -1.046 | 0.340 | |
| | | | | WT | 0.556 | 0.242 | | | | |
| Table S5A | Figure 1H,I / Figure S7 | D0 | Short | KO | 0.111 | 0.111 | Mann-Whitney U Test | -1.000 | 0.730 | |
| | | | | WT | 0.000 | 0.000 | | | | |
| | | D1 | Short | KO | 0.167 | 0.118 | | -1.455 | 0.436 | |
| | | | | WT | 0.000 | 0.000 | | | | |
| | | D2 | Short | KO | 0.444 | 0.227 | | -1.626 | 0.222 | |
| | | | | WT | 0.056 | 0.056 | | | | |
| | | D3 | Short | KO | 0.556 | 0.242 | | 0.000 | 1.000 | |
| | | | | WT | 0.556 | 0.242 | | | | |
| | | D4 | Short | KO | 1.000 | 0.408 | | -0.193 | 0.863 | |
| | | | | WT | 0.889 | 0.261 | | | | |
| | | D5 | Short | KO | 0.778 | 0.278 | | -0.579 | 0.605 | |
| | | | | WT | 0.556 | 0.242 | | | | |
| | | D6 | Short | KO | 0.667 | 0.373 | | -0.587 | 0.605 | |
| | | | | WT | 0.778 | 0.278 | | | | |
| | | D7 | Short | KO | 0.889 | 0.423 | | -0.743 | 0.546 | |
| | | | | WT | 0.889 | 0.200 | | | | |
| | | D8 | Short | KO | 1.111 | 0.261 | | -0.191 | 0.863 | |
| | | | | WT | 1.111 | 0.309 | | | | |
| | | D9 | Short | KO | 1.111 | 0.484 | | -2.013 | 0.094 | |
| | | | | WT | 0.111 | 0.111 | | | | |
| | | D10 | Short | KO | 0.778 | 0.278 | | -1.190 | 0.297 | |
| | | | | WT | 0.333 | 0.167 | | | | |
| | | D11 | Short | KO | 1.111 | 0.351 | | -1.414 | 0.222 | |
| | | | | WT | 0.556 | 0.377 | | | | |
| | | D12 | Short | KO | 1.111 | 0.351 | | -0.894 | 0.436 | |
| | | | | WT | 0.667 | 0.236 | | | | |
| | | D13 | Short | KO | 0.444 | 0.176 | | -0.201 | 0.863 | |
| | | | | WT | 0.667 | 0.333 | | | | |
| | | D14 | Short | KO | 1.111 | 0.351 | | -0.940 | 0.387 | |
| | | | | WT | 0.667 | 0.289 | | | | |
| | | D15 | Short | KO | 1.444 | 0.580 | | -0.943 | 0.387 | |
| | | | | WT | 0.556 | 0.176 | | | | |
| | | D16 | Short | KO | 0.889 | 0.309 | | -1.080 | 0.340 | |
| | | | | WT | 0.444 | 0.242 | | | | |

| HE frequency in 0–1 s window | | | | | | | | | | |
|---|---|---|---|---|---|---|---|---|---|---|
| Table | Related Figure | Day | Group | Condition | Mean | SEM | Stastistical test | Z | P | * |
| Table S5 | Figure 1H,I / Figure S7 | D0 | KO | Long | 0.056 | 0.056 | Wilcoxon Signed Ranks Test | -1.000 | 0.317 | |
| | | | | Short | 0.111 | 0.111 | | | | |
| | | D1 | KO | Long | 0.111 | 0.073 | | -0.577 | 0.564 | |
| | | | | Short | 0.167 | 0.118 | | | | |
| | | D2 | KO | Long | 0.167 | 0.118 | | -1.633 | 0.102 | |
| | | | | Short | 0.444 | 0.227 | | | | |
| | | D3 | KO | Long | 0.556 | 0.338 | | -0.378 | 0.705 | |
| | | | | Short | 0.556 | 0.242 | | | | |
| | | D4 | KO | Long | 0.556 | 0.444 | | -1.081 | 0.279 | |
| | | | | Short | 1.000 | 0.408 | | | | |
| | | D5 | KO | Long | 0.333 | 0.167 | | -1.633 | 0.102 | |
| | | | | Short | 0.778 | 0.278 | | | | |
| | | D6 | KO | Long | 1.222 | 0.434 | | -0.843 | 0.399 | |
| | | | | Short | 0.667 | 0.373 | | | | |
| | | D7 | KO | Long | 0.556 | 0.294 | | -0.647 | 0.518 | |
| | | | | Short | 0.889 | 0.423 | | | | |
| | | D8 | KO | Long | 1.778 | 0.619 | | -1.186 | 0.236 | |
| | | | | Short | 1.111 | 0.261 | | | | |
| | | D9 | KO | Long | 1.222 | 0.521 | | -0.175 | 0.861 | |
| | | | | Long | 1.111 | 0.484 | | | | |
| | | D10 | KO | Short | 0.556 | 0.176 | | -1.000 | 0.317 | |
| | | | | Long | 0.778 | 0.278 | | | | |
| | | D11 | KO | Short | 1.000 | 0.289 | | -0.302 | 0.763 | |
| | | | | Long | 1.111 | 0.351 | | | | |
| | | D12 | KO | Short | 0.333 | 0.236 | | -1.841 | 0.066 | |
| | | | | Long | 1.111 | 0.351 | | | | |
| | | D13 | KO | Short | 1.000 | 0.553 | | -0.647 | 0.518 | |
| | | | | Long | 0.444 | 0.176 | | | | |
| | | D14 | KO | Short | 1.333 | 0.408 | | -0.412 | 0.680 | |
| | | | | Long | 1.111 | 0.351 | | | | |
| | | D15 | KO | Short | 1.000 | 0.333 | | -0.632 | 0.527 | |
| | | | | Long | 1.444 | 0.580 | | | | |
| | | D16 | KO | Short | 1.444 | 0.580 | | -1.289 | 0.197 | |
| | | | | Long | 0.889 | 0.309 | | | | |
| Table S5 | Figure 1H,I / Figure S7 | D0 | WT | Long | 0.278 | 0.147 | Wilcoxon Signed Ranks Test | -1.633 | 0.102 | |
| | | | | Short | 0.000 | 0.000 | | | | |
| | | D1 | WT | Long | 0.222 | 0.147 | | -1.414 | 0.157 | |
| | | | | Short | 0.000 | 0.000 | | | | |
| | | D2 | WT | Long | 0.778 | 0.434 | | -1.473 | 0.141 | |
| | | | | Short | 0.056 | 0.056 | | | | |
| | | D3 | WT | Long | 0.333 | 0.167 | | -1.414 | 0.157 | |
| | | | | Short | 0.556 | 0.242 | | | | |
| | | D4 | WT | Long | 1.111 | 0.309 | | -0.632 | 0.527 | |
| | | | | Short | 0.889 | 0.261 | | | | |
| | | D5 | WT | Long | 0.333 | 0.236 | | -0.541 | 0.589 | |
| | | | | Short | 0.556 | 0.242 | | | | |
| | | D6 | WT | Long | 0.556 | 0.338 | | -0.541 | 0.589 | |
| | | | | Short | 0.778 | 0.278 | | | | |
| | | D7 | WT | Long | 0.333 | 0.167 | | -1.667 | 0.096 | |
| | | | | Short | 0.889 | 0.200 | | | | |
| | | D8 | WT | Long | 1.333 | 0.408 | | -0.574 | 0.566 | |
| | | | | Short | 1.111 | 0.309 | | | | |
| | | D9 | WT | Long | 1.000 | 0.289 | | -2.271 | 0.023 | * |
| | | | | Long | 0.111 | 0.111 | | | | |
| | | D10 | WT | Short | 1.222 | 0.401 | | -1.807 | 0.071 | |
| | | | | Long | 0.333 | 0.167 | | | | |
| | | D11 | WT | Short | 0.778 | 0.434 | | -0.707 | 0.480 | |
| | | | | Long | 0.556 | 0.377 | | | | |
| | | D12 | WT | Short | 0.444 | 0.242 | | -1.000 | 0.317 | |
| | | | | Long | 0.667 | 0.236 | | | | |
| | | D13 | WT | Short | 0.778 | 0.278 | | -0.276 | 0.783 | |
| | | | | Long | 0.667 | 0.333 | | | | |
| | | D14 | WT | Short | 0.444 | 0.242 | | -0.743 | 0.458 | |
| | | | | Long | 0.667 | 0.289 | | | | |
| | | D15 | WT | Short | 0.444 | 0.294 | | -0.176 | 0.860 | |
| | | | | Long | 0.556 | 0.176 | | | | |
| | | D16 | WT | Short | 0.556 | 0.242 | | -0.333 | 0.739 | |
| | | | | Long | 0.444 | 0.242 | | | | |

5_2

| HE frequency in 1–2 s window | | | | | | | | | | |
|---|---|---|---|---|---|---|---|---|---|---|
| Table | Related Figure | Day | Condition | Group | Mean | SEM | Stastistical test | Z | P | * |
| Table S5 | Figure 1H,I / Figure S7 | D0 | Long | KO | 0.278 | 0.222 | Mann-Whitney U Test | -1.455 | 0.436 | |
| | | | | WT | 0.000 | 0.000 | | | | |
| | | D1 | Long | KO | 0.056 | 0.056 | | -0.081 | 1.000 | |
| | | | | WT | 0.111 | 0.111 | | | | |
| | | D2 | Long | KO | 0.278 | 0.147 | | -0.910 | 0.546 | |
| | | | | WT | 0.444 | 0.444 | | | | |
| | | D3 | Long | KO | 0.333 | 0.167 | | -0.338 | 0.796 | |
| | | | | WT | 0.444 | 0.338 | | | | |
| | | D4 | Long | KO | 0.889 | 0.423 | | -0.143 | 0.931 | |
| | | | | WT | 0.778 | 0.278 | | | | |
| | | D5 | Long | KO | 1.222 | 0.324 | | -0.368 | 0.730 | |
| | | | | WT | 1.222 | 0.494 | | | | |
| | | D6 | Long | KO | 0.889 | 0.200 | | -0.335 | 0.796 | |
| | | | | WT | 1.111 | 0.351 | | | | |
| | | D7 | Long | KO | 1.667 | 0.373 | | -0.809 | 0.489 | |
| | | | | WT | 1.222 | 0.222 | | | | |
| | | D8 | Long | KO | 1.889 | 0.539 | | -0.775 | 0.489 | |
| | | | | WT | 1.333 | 0.333 | | | | |
| | | D9 | Long | KO | 2.222 | 0.465 | | -1.590 | 0.136 | |
| | | | | WT | 1.222 | 0.324 | | | | |
| | | D10 | Long | KO | 1.556 | 0.377 | | -0.183 | 0.863 | |
| | | | | WT | 1.444 | 0.412 | | | | |
| | | D11 | Long | KO | 2.000 | 0.986 | | -0.549 | 0.605 | |
| | | | | WT | 1.556 | 0.294 | | | | |
| | | D12 | Long | KO | 2.889 | 0.807 | | -0.454 | 0.666 | |
| | | | | WT | 2.222 | 0.572 | | | | |
| | | D13 | Long | KO | 2.667 | 0.601 | | -1.769 | 0.094 | |
| | | | | WT | 1.222 | 0.465 | | | | |
| | | D14 | Long | KO | 1.889 | 0.512 | | -0.090 | 0.931 | |
| | | | | WT | 2.000 | 0.577 | | | | |
| | | D15 | Long | KO | 2.444 | 0.729 | | -1.009 | 0.340 | |
| | | | | WT | 1.444 | 0.530 | | | | |
| | | D16 | Long | KO | 2.111 | 0.696 | | -1.262 | 0.258 | |
| | | | | WT | 0.889 | 0.455 | | | | |
| Table S5A | Figure 1H,I / Figure S7 | D0 | Short | KO | 0.167 | 0.083 | Mann-Whitney U Test | -0.913 | 0.546 | |
| | | | | WT | 0.111 | 0.111 | | | | |
| | | D1 | Short | KO | 0.444 | 0.242 | | -1.837 | 0.258 | |
| | | | | WT | 0.000 | 0.000 | | | | |
| | | D2 | Short | KO | 0.444 | 0.155 | | -0.528 | 0.666 | |
| | | | | WT | 0.333 | 0.144 | | | | |
| | | D3 | Short | KO | 0.556 | 0.176 | | -0.248 | 0.863 | |
| | | | | WT | 0.667 | 0.236 | | | | |
| | | D4 | Short | KO | 1.222 | 0.619 | | -0.876 | 0.436 | |
| | | | | WT | 1.333 | 0.373 | | | | |
| | | D5 | Short | KO | 0.889 | 0.200 | | 0.000 | 1.000 | |
| | | | | WT | 1.000 | 0.333 | | | | |
| | | D6 | Short | KO | 1.222 | 0.222 | | -2.466 | 0.019 | * |
| | | | | WT | 0.333 | 0.236 | | | | |
| | | D7 | Short | KO | 2.111 | 0.588 | | -2.233 | 0.031 | * |
| | | | | WT | 0.444 | 0.242 | | | | |
| | | D8 | Short | KO | 1.778 | 0.741 | | -0.369 | 0.730 | |
| | | | | WT | 1.111 | 0.309 | | | | |
| | | D9 | Short | KO | 2.444 | 0.475 | | -2.022 | 0.050 | |
| | | | | WT | 1.222 | 0.222 | | | | |
| | | D10 | Short | KO | 2.111 | 0.423 | | -0.859 | 0.436 | |
| | | | | WT | 1.556 | 0.503 | | | | |
| | | D11 | Short | KO | 1.444 | 0.648 | | -0.773 | 0.489 | |
| | | | | WT | 1.667 | 0.408 | | | | |
| | | D12 | Short | KO | 1.778 | 0.572 | | -0.181 | 0.863 | |
| | | | | WT | 1.444 | 0.412 | | | | |
| | | D13 | Short | KO | 2.333 | 0.764 | | -1.284 | 0.222 | |
| | | | | WT | 0.778 | 0.222 | | | | |
| | | D14 | Short | KO | 2.667 | 0.764 | | -1.219 | 0.258 | |
| | | | | WT | 1.333 | 0.333 | | | | |
| | | D15 | Short | KO | 2.222 | 0.830 | | -1.216 | 0.258 | |
| | | | | WT | 1.111 | 0.564 | | | | |
| | | D16 | Short | KO | 2.444 | 0.689 | | -0.816 | 0.436 | |
| | | | | WT | 1.667 | 0.373 | | | | |

| HE frequency in 1–2 s window | | | | | | | | | | |
|---|---|---|---|---|---|---|---|---|---|---|
| Table | Related Figure | Day | Group | Condition | Mean | SEM | Stastistical test | Z | P | * |
| Table S5 | Figure 1H,I / Figure S7 | D0 | KO | Long | 0.278 | 0.222 | Wilcoxon Signed Ranks Test | 0.000 | 1.000 | |
| | | | | Short | 0.167 | 0.083 | | | | |
| | | D1 | KO | Long | 0.056 | 0.056 | | -1.473 | 0.141 | |
| | | | | Short | 0.444 | 0.242 | | | | |
| | | D2 | KO | Long | 0.278 | 0.147 | | -0.780 | 0.435 | |
| | | | | Short | 0.444 | 0.155 | | | | |
| | | D3 | KO | Long | 0.333 | 0.167 | | -1.000 | 0.317 | |
| | | | | Short | 0.556 | 0.176 | | | | |
| | | D4 | KO | Long | 0.889 | 0.423 | | -0.319 | 0.750 | |
| | | | | Short | 1.222 | 0.619 | | | | |
| | | D5 | KO | Long | 1.222 | 0.324 | | -1.000 | 0.317 | |
| | | | | Short | 0.889 | 0.200 | | | | |
| | | D6 | KO | Long | 0.889 | 0.200 | | -0.966 | 0.334 | |
| | | | | Short | 1.222 | 0.222 | | | | |
| | | D7 | KO | Long | 1.667 | 0.373 | | -0.557 | 0.577 | |
| | | | | Short | 2.111 | 0.588 | | | | |
| | | D8 | KO | Long | 1.889 | 0.539 | | -0.259 | 0.796 | |
| | | | | Short | 1.778 | 0.741 | | | | |
| | | D9 | KO | Long | 2.222 | 0.465 | | -0.541 | 0.589 | |
| | | | | Long | 2.444 | 0.475 | | | | |
| | | D10 | KO | Short | 1.556 | 0.377 | | -1.186 | 0.236 | |
| | | | | Long | 2.111 | 0.423 | | | | |
| | | D11 | KO | Short | 2.000 | 0.986 | | -1.342 | 0.180 | |
| | | | | Long | 1.444 | 0.648 | | | | |
| | | D12 | KO | Short | 2.889 | 0.807 | | -1.558 | 0.119 | |
| | | | | Long | 1.778 | 0.572 | | | | |
| | | D13 | KO | Short | 2.667 | 0.601 | | -0.638 | 0.524 | |
| | | | | Long | 2.333 | 0.764 | | | | |
| | | D14 | KO | Short | 1.889 | 0.512 | | -0.938 | 0.348 | |
| | | | | Long | 2.667 | 0.764 | | | | |
| | | D15 | KO | Short | 2.444 | 0.729 | | -0.406 | 0.684 | |
| | | | | Long | 2.222 | 0.830 | | | | |
| | | D16 | KO | Short | 2.111 | 0.696 | | -0.722 | 0.470 | |
| | | | | Long | 2.444 | 0.689 | | | | |
| Table S5 | Figure 1H,I / Figure S7 | D0 | WT | Long | 0.000 | 0.000 | Wilcoxon Signed Ranks Test | -1.000 | 0.317 | |
| | | | | Short | 0.111 | 0.111 | | | | |
| | | D1 | WT | Long | 0.111 | 0.111 | | -1.000 | 0.317 | |
| | | | | Short | 0.000 | 0.000 | | | | |
| | | D2 | WT | Long | 0.444 | 0.444 | | -0.680 | 0.496 | |
| | | | | Short | 0.333 | 0.144 | | | | |
| | | D3 | WT | Long | 0.444 | 0.338 | | -1.000 | 0.317 | |
| | | | | Short | 0.667 | 0.236 | | | | |
| | | D4 | WT | Long | 0.778 | 0.278 | | -1.667 | 0.096 | |
| | | | | Short | 1.333 | 0.373 | | | | |
| | | D5 | WT | Long | 1.222 | 0.494 | | -0.552 | 0.581 | |
| | | | | Short | 1.000 | 0.333 | | | | |
| | | D6 | WT | Long | 1.111 | 0.351 | | -1.933 | 0.053 | |
| | | | | Short | 0.333 | 0.236 | | | | |
| | | D7 | WT | Long | 1.222 | 0.222 | | -1.823 | 0.068 | |
| | | | | Short | 0.444 | 0.242 | | | | |
| | | D8 | WT | Long | 1.333 | 0.333 | | -0.632 | 0.527 | |
| | | | | Short | 1.111 | 0.309 | | | | |
| | | D9 | WT | Long | 1.222 | 0.324 | | 0.000 | 1.000 | |
| | | | | Long | 1.222 | 0.222 | | | | |
| | | D10 | WT | Short | 1.444 | 0.412 | | -0.447 | 0.655 | |
| | | | | Long | 1.556 | 0.503 | | | | |
| | | D11 | WT | Short | 1.556 | 0.294 | | -0.431 | 0.666 | |
| | | | | Long | 1.667 | 0.408 | | | | |
| | | D12 | WT | Short | 2.222 | 0.572 | | -1.552 | 0.121 | |
| | | | | Long | 1.444 | 0.412 | | | | |
| | | D13 | WT | Short | 1.222 | 0.465 | | -1.134 | 0.257 | |
| | | | | Long | 0.778 | 0.222 | | | | |
| | | D14 | WT | Short | 2.000 | 0.577 | | -1.382 | 0.167 | |
| | | | | Long | 1.333 | 0.333 | | | | |
| | | D15 | WT | Short | 1.444 | 0.530 | | -0.414 | 0.679 | |
| | | | | Long | 1.111 | 0.564 | | | | |
| | | D16 | WT | Short | 0.889 | 0.455 | | -1.890 | 0.059 | |
| | | | | Long | 1.667 | 0.373 | | | | |

5_3

| HE frequency in 2–3 s window | | | | | | | | | | |
|---|---|---|---|---|---|---|---|---|---|---|
| Table | Related Figure | Day | Condition | Group | Mean | SEM | Stastistical test | Z | P | * |
| Table S5 | Figure 1H,I / Figure S7 | D0 | Long | KO | 0.333 | 0.220 | Mann-Whitney U Test | -1.837 | 0.258 | |
| | | | | WT | 0.000 | 0.000 | | | | |
| | | D1 | Long | KO | 0.500 | 0.236 | | -1.514 | 0.258 | |
| | | | | WT | 0.111 | 0.111 | | | | |
| | | D2 | Long | KO | 0.278 | 0.147 | | -1.035 | 0.489 | |
| | | | | WT | 0.444 | 0.444 | | | | |
| | | D3 | Long | KO | 0.778 | 0.434 | | -0.458 | 0.730 | |
| | | | | WT | 0.444 | 0.338 | | | | |
| | | D4 | Long | KO | 0.444 | 0.242 | | -0.843 | 0.489 | |
| | | | | WT | 0.778 | 0.278 | | | | |
| | | D5 | Long | KO | 0.778 | 0.434 | | -0.530 | 0.666 | |
| | | | | WT | 1.222 | 0.494 | | | | |
| | | D6 | Long | KO | 1.333 | 0.408 | | -0.279 | 0.796 | |
| | | | | WT | 1.111 | 0.351 | | | | |
| | | D7 | Long | KO | 2.222 | 0.662 | | -1.274 | 0.222 | |
| | | | | WT | 1.222 | 0.222 | | | | |
| | | D8 | Long | KO | 1.889 | 0.735 | | -0.273 | 0.796 | |
| | | | | WT | 1.333 | 0.333 | | | | |
| | | D9 | Long | KO | 2.667 | 0.553 | | -0.365 | 0.730 | |
| | | | | WT | 1.222 | 0.324 | | | | |
| | | D10 | Long | KO | 2.333 | 0.833 | | -0.045 | 1.000 | |
| | | | | WT | 1.444 | 0.412 | | | | |
| | | D11 | Long | KO | 1.778 | 0.760 | | -0.635 | 0.546 | |
| | | | | WT | 1.556 | 0.294 | | | | |
| | | D12 | Long | KO | 2.333 | 0.624 | | -0.855 | 0.436 | |
| | | | | WT | 2.222 | 0.572 | | | | |
| | | D13 | Long | KO | 2.667 | 0.782 | | -0.637 | 0.546 | |
| | | | | WT | 1.222 | 0.465 | | | | |
| | | D14 | Long | KO | 2.556 | 0.503 | | -0.093 | 0.931 | |
| | | | | WT | 2.000 | 0.577 | | | | |
| | | D15 | Long | KO | 2.778 | 0.760 | | -0.454 | 0.666 | |
| | | | | WT | 1.444 | 0.530 | | | | |
| | | D16 | Long | KO | 2.667 | 0.833 | | -0.408 | 0.730 | |
| | | | | WT | 0.889 | 0.455 | | | | |
| Table S5A | Figure 1H,I / Figure S7 | D0 | Short | KO | 0.263 | 0.088 | Mann-Whitney U Test | 0.000 | 1.000 | |
| | | | | WT | 0.000 | 0.000 | | | | |
| | | D1 | Short | KO | 1.120 | 0.456 | | -2.557 | 0.050 | |
| | | | | WT | 0.000 | 0.000 | | | | |
| | | D2 | Short | KO | 0.371 | 0.103 | | -1.389 | 0.222 | |
| | | | | WT | 0.222 | 0.121 | | | | |
| | | D3 | Short | KO | 0.751 | 0.149 | | -0.491 | 0.666 | |
| | | | | WT | 0.667 | 0.373 | | | | |
| | | D4 | Short | KO | 0.420 | 0.061 | | -0.356 | 0.796 | |
| | | | | WT | 0.556 | 0.338 | | | | |
| | | D5 | Short | KO | 0.961 | 0.136 | | -0.681 | 0.546 | |
| | | | | WT | 1.000 | 0.601 | | | | |
| | | D6 | Short | KO | 0.641 | 0.154 | | -1.459 | 0.222 | |
| | | | | WT | 0.444 | 0.294 | | | | |
| | | D7 | Short | KO | 1.179 | 0.313 | | -0.793 | 0.489 | |
| | | | | WT | 1.444 | 0.338 | | | | |
| | | D8 | Short | KO | 1.258 | 0.305 | | -1.173 | 0.258 | |
| | | | | WT | 1.667 | 0.441 | | | | |
| | | D9 | Short | KO | 1.528 | 0.452 | | -1.199 | 0.258 | |
| | | | | WT | 1.444 | 0.338 | | | | |
| | | D10 | Short | KO | 1.422 | 0.344 | | -0.045 | 1.000 | |
| | | | | WT | 1.444 | 0.294 | | | | |
| | | D11 | Short | KO | 1.330 | 0.335 | | -0.135 | 0.931 | |
| | | | | WT | 2.111 | 0.539 | | | | |
| | | D12 | Short | KO | 1.406 | 0.322 | | -0.413 | 0.730 | |
| | | | | WT | 1.556 | 0.444 | | | | |
| | | D13 | Short | KO | 1.542 | 0.400 | | -0.273 | 0.796 | |
| | | | | WT | 1.889 | 0.261 | | | | |
| | | D14 | Short | KO | 1.345 | 0.475 | | -1.944 | 0.063 | |
| | | | | WT | 0.778 | 0.324 | | | | |
| | | D15 | Short | KO | 1.507 | 0.422 | | -0.972 | 0.387 | |
| | | | | WT | 1.111 | 0.351 | | | | |
| | | D16 | Short | KO | 1.499 | 0.399 | | -0.772 | 0.489 | |
| | | | | WT | 1.556 | 0.338 | | | | |

| HE frequency in 2–3 s window | | | | | | | | | | |
|---|---|---|---|---|---|---|---|---|---|---|
| Table | Related Figure | Day | Group | Condition | Mean | SEM | Stastistical test | Z | P | * |
| Table S5 | Figure 1H,I / Figure S7 | D0 | KO | Long | 0.333 | 0.220 | Wilcoxon Signed Ranks Test | -1.633 | 0.102 | |
| | | | | Short | 0.263 | 0.088 | | | | |
| | | D1 | KO | Long | 0.500 | 0.236 | | -0.216 | 0.829 | |
| | | | | Short | 1.120 | 0.456 | | | | |
| | | D2 | KO | Long | 0.278 | 0.147 | | -1.633 | 0.102 | |
| | | | | Short | 0.371 | 0.103 | | | | |
| | | D3 | KO | Long | 0.778 | 0.434 | | 0.000 | 1.000 | |
| | | | | Short | 0.751 | 0.149 | | | | |
| | | D4 | KO | Long | 0.444 | 0.242 | | -0.333 | 0.739 | |
| | | | | Short | 0.420 | 0.061 | | | | |
| | | D5 | KO | Long | 0.778 | 0.434 | | -0.962 | 0.336 | |
| | | | | Short | 0.961 | 0.136 | | | | |
| | | D6 | KO | Long | 1.333 | 0.408 | | -0.213 | 0.832 | |
| | | | | Short | 0.641 | 0.154 | | | | |
| | | D7 | KO | Long | 2.222 | 0.662 | | -1.586 | 0.113 | |
| | | | | Short | 1.179 | 0.313 | | | | |
| | | D8 | KO | Long | 1.889 | 0.735 | | -0.954 | 0.340 | |
| | | | | Short | 1.258 | 0.305 | | | | |
| | | D9 | KO | Long | 2.667 | 0.553 | | -1.127 | 0.260 | |
| | | | | Long | 1.528 | 0.452 | | | | |
| | | D10 | KO | Short | 2.333 | 0.833 | | -0.862 | 0.389 | |
| | | | | Long | 1.422 | 0.344 | | | | |
| | | D11 | KO | Short | 1.778 | 0.760 | | -0.816 | 0.414 | |
| | | | | Long | 1.330 | 0.335 | | | | |
| | | D12 | KO | Short | 2.333 | 0.624 | | -0.531 | 0.595 | |
| | | | | Long | 1.406 | 0.322 | | | | |
| | | D13 | KO | Short | 2.667 | 0.782 | | -1.194 | 0.233 | |
| | | | | Long | 1.542 | 0.400 | | | | |
| | | D14 | KO | Short | 2.556 | 0.503 | | -1.294 | 0.196 | |
| | | | | Long | 1.345 | 0.475 | | | | |
| | | D15 | KO | Short | 2.778 | 0.760 | | -1.802 | 0.072 | |
| | | | | Long | 1.507 | 0.422 | | | | |
| | | D16 | KO | Short | 2.667 | 0.833 | | -1.023 | 0.306 | |
| | | | | Long | 1.499 | 0.399 | | | | |
| Table S5 | Figure 1H,I / Figure S7 | D0 | WT | Long | 0.000 | 0.000 | Wilcoxon Signed Ranks Test | 0.000 | 1.000 | |
| | | | | Short | 0.000 | 0.000 | | | | |
| | | D1 | WT | Long | 0.111 | 0.111 | | -1.000 | 0.317 | |
| | | | | Short | 0.000 | 0.000 | | | | |
| | | D2 | WT | Long | 0.444 | 0.444 | | -1.414 | 0.157 | |
| | | | | Short | 0.222 | 0.121 | | | | |
| | | D3 | WT | Long | 0.444 | 0.338 | | -0.552 | 0.581 | |
| | | | | Short | 0.667 | 0.373 | | | | |
| | | D4 | WT | Long | 0.778 | 0.278 | | -0.541 | 0.589 | |
| | | | | Short | 0.556 | 0.338 | | | | |
| | | D5 | WT | Long | 1.222 | 0.494 | | -0.085 | 0.932 | |
| | | | | Short | 1.000 | 0.601 | | | | |
| | | D6 | WT | Long | 1.111 | 0.351 | | -1.633 | 0.102 | |
| | | | | Short | 0.444 | 0.294 | | | | |
| | | D7 | WT | Long | 1.222 | 0.222 | | -0.965 | 0.335 | |
| | | | | Short | 1.444 | 0.338 | | | | |
| | | D8 | WT | Long | 1.333 | 0.333 | | -0.072 | 0.943 | |
| | | | | Short | 1.667 | 0.441 | | | | |
| | | D9 | WT | Long | 1.222 | 0.324 | | -1.310 | 0.190 | |
| | | | | Long | 1.444 | 0.338 | | | | |
| | | D10 | WT | Short | 1.444 | 0.412 | | -0.791 | 0.429 | |
| | | | | Long | 1.444 | 0.294 | | | | |
| | | D11 | WT | Short | 1.556 | 0.294 | | -0.568 | 0.570 | |
| | | | | Long | 2.111 | 0.539 | | | | |
| | | D12 | WT | Short | 2.222 | 0.572 | | -0.359 | 0.719 | |
| | | | | Long | 1.556 | 0.444 | | | | |
| | | D13 | WT | Short | 1.222 | 0.465 | | 0.000 | 1.000 | |
| | | | | Long | 1.889 | 0.261 | | | | |
| | | D14 | WT | Short | 2.000 | 0.577 | | -2.036 | 0.042 | * |
| | | | | Long | 0.778 | 0.324 | | | | |
| | | D15 | WT | Short | 1.444 | 0.530 | | -1.549 | 0.121 | |
| | | | | Long | 1.111 | 0.351 | | | | |
| | | D16 | WT | Short | 0.889 | 0.455 | | -1.134 | 0.257 | |
| | | | | Long | 1.556 | 0.338 | | | | |

Abbreviations: HE, head entry; D, training day; KO, CRBN KO mice; WT, wild-type mice.

*, *p < 0.05, **p < 0.01, ***p < 0.001.

## 5_4

HE frequency in 3–4 s window

| Table | Related Figure | Day | Condition | Group | Mean | SEM | Statistical test | Z | P | * |
|---|---|---|---|---|---|---|---|---|---|---|
| Table S5 | Figure 1H,I / Figure S7 | D0 | Long | KO | 0.444 | 0.212 | Mann-Whitney U Test | -2.187 | 0.113 | |
| | | | | WT | 0.000 | 0.000 | | | | |
| | | D1 | Long | KO | 0.167 | 0.118 | | 0.000 | 1.000 | |
| | | | | WT | 0.167 | 0.118 | | | | |
| | | D2 | Long | KO | 0.278 | 0.147 | | -1.215 | 0.387 | |
| | | | | WT | 0.056 | 0.056 | | | | |
| | | D3 | Long | KO | 0.556 | 0.242 | | -1.578 | 0.222 | |
| | | | | WT | 0.111 | 0.111 | | | | |
| | | D4 | Long | KO | 1.000 | 0.373 | | -0.372 | 0.730 | |
| | | | | WT | 1.111 | 0.309 | | | | |
| | | D5 | Long | KO | 0.667 | 0.289 | | -0.293 | 0.796 | |
| | | | | WT | 1.000 | 0.471 | | | | |
| | | D6 | Long | KO | 1.556 | 0.503 | | -1.269 | 0.258 | |
| | | | | WT | 0.778 | 0.222 | | | | |
| | | D7 | Long | KO | 1.333 | 0.645 | | -0.047 | 1.000 | |
| | | | | WT | 0.889 | 0.351 | | | | |
| | | D8 | Long | KO | 2.222 | 0.741 | | -0.454 | 0.666 | |
| | | | | WT | 1.667 | 0.333 | | | | |
| | | D9 | Long | KO | 3.111 | 0.935 | | -0.761 | 0.489 | |
| | | | | WT | 2.000 | 0.577 | | | | |
| | | D10 | Long | KO | 2.889 | 0.992 | | -0.227 | 0.863 | |
| | | | | WT | 1.778 | 0.401 | | | | |
| | | D11 | Long | KO | 3.000 | 1.067 | | -0.588 | 0.605 | |
| | | | | WT | 2.111 | 0.633 | | | | |
| | | D12 | Long | KO | 1.667 | 0.408 | | -0.281 | 0.796 | |
| | | | | WT | 1.889 | 0.200 | | | | |
| | | D13 | Long | KO | 2.444 | 0.626 | | -0.270 | 0.796 | |
| | | | | WT | 2.222 | 0.703 | | | | |
| | | D14 | Long | KO | 2.444 | 0.503 | | -0.988 | 0.340 | |
| | | | | WT | 1.889 | 0.655 | | | | |
| | | D15 | Long | KO | 2.556 | 0.530 | | -1.078 | 0.297 | |
| | | | | WT | 1.889 | 0.676 | | | | |
| | | D16 | Long | KO | 1.111 | 0.309 | | -1.365 | 0.190 | |
| | | | | WT | 2.222 | 0.619 | | | | |
| Table S5A | Figure 1H,I / Figure S7 | D0 | Short | KO | 0.167 | 0.083 | Mann-Whitney U Test | -1.844 | 0.258 | |
| | | | | WT | 0.000 | 0.000 | | | | |
| | | D1 | Short | KO | 0.556 | 0.227 | | -1.847 | 0.136 | |
| | | | | WT | 0.111 | 0.111 | | | | |
| | | D2 | Short | KO | 0.444 | 0.242 | | -0.403 | 0.730 | |
| | | | | WT | 0.556 | 0.256 | | | | |
| | | D3 | Short | KO | 0.556 | 0.242 | | -0.826 | 0.489 | |
| | | | | WT | 0.889 | 0.309 | | | | |
| | | D4 | Short | KO | 1.222 | 0.364 | | -1.363 | 0.222 | |
| | | | | WT | 0.556 | 0.242 | | | | |
| | | D5 | Short | KO | 1.444 | 0.444 | | -0.046 | 1.000 | |
| | | | | WT | 1.556 | 0.530 | | | | |
| | | D6 | Short | KO | 1.000 | 0.236 | | -0.328 | 0.796 | |
| | | | | WT | 0.889 | 0.309 | | | | |
| | | D7 | Short | KO | 2.000 | 0.500 | | -0.870 | 0.436 | |
| | | | | WT | 1.444 | 0.444 | | | | |
| | | D8 | Short | KO | 2.111 | 0.455 | | -2.332 | 0.031 | * |
| | | | | WT | 0.778 | 0.324 | | | | |
| | | D9 | Short | KO | 2.222 | 0.521 | | -0.499 | 0.666 | |
| | | | | WT | 1.889 | 0.655 | | | | |
| | | D10 | Short | KO | 1.889 | 0.455 | | -0.368 | 0.730 | |
| | | | | WT | 1.667 | 0.373 | | | | |
| | | D11 | Short | KO | 1.556 | 0.530 | | -0.092 | 0.931 | |
| | | | | WT | 1.333 | 0.236 | | | | |
| | | D12 | Short | KO | 1.556 | 0.444 | | -0.046 | 1.000 | |
| | | | | WT | 1.556 | 0.475 | | | | |
| | | D13 | Short | KO | 1.333 | 0.333 | | -0.458 | 0.666 | |
| | | | | WT | 1.556 | 0.377 | | | | |
| | | D14 | Short | KO | 1.222 | 0.434 | | -0.787 | 0.489 | |
| | | | | WT | 1.556 | 0.377 | | | | |
| | | D15 | Short | KO | 1.111 | 0.351 | | -0.731 | 0.489 | |
| | | | | WT | 1.778 | 0.572 | | | | |
| | | D16 | Short | KO | 1.556 | 0.530 | | 0.000 | 1.000 | |
| | | | | WT | 1.444 | 0.412 | | | | |

HE frequency in 3–4 s window

| Table | Related Figure | Day | Group | Condition | Mean | SEM | Statistical test | Z | P | * |
|---|---|---|---|---|---|---|---|---|---|---|
| Table S5 | Figure 1H,I / Figure S7 | D0 | KO | Long | 0.444 | 0.212 | Wilcoxon Signed Ranks Test | -1.186 | 0.236 | |
| | | | | Short | 0.167 | 0.083 | | | | |
| | | D1 | KO | Long | 0.167 | 0.118 | | -1.604 | 0.109 | |
| | | | | Short | 0.556 | 0.227 | | | | |
| | | D2 | KO | Long | 0.278 | 0.147 | | -1.089 | 0.276 | |
| | | | | Short | 0.444 | 0.242 | | | | |
| | | D3 | KO | Long | 0.556 | 0.242 | | 0.000 | 1.000 | |
| | | | | Short | 0.556 | 0.242 | | | | |
| | | D4 | KO | Long | 1.000 | 0.373 | | -0.333 | 0.739 | |
| | | | | Short | 1.222 | 0.364 | | | | |
| | | D5 | KO | Long | 0.667 | 0.289 | | -1.403 | 0.161 | |
| | | | | Short | 1.444 | 0.444 | | | | |
| | | D6 | KO | Long | 1.556 | 0.503 | | -1.265 | 0.206 | |
| | | | | Short | 1.000 | 0.236 | | | | |
| | | D7 | KO | Long | 1.333 | 0.645 | | -1.150 | 0.250 | |
| | | | | Short | 2.000 | 0.500 | | | | |
| | | D8 | KO | Long | 2.222 | 0.741 | | -0.284 | 0.776 | |
| | | | | Short | 2.111 | 0.455 | | | | |
| | | D9 | KO | Long | 3.111 | 0.935 | | -1.200 | 0.230 | |
| | | | | Short | 2.222 | 0.521 | | | | |
| | | D10 | KO | Short | 2.889 | 0.992 | | -1.109 | 0.268 | |
| | | | | Long | 1.889 | 0.455 | | | | |
| | | D11 | KO | Short | 3.000 | 1.067 | | -1.378 | 0.168 | |
| | | | | Long | 1.556 | 0.530 | | | | |
| | | D12 | KO | Short | 1.667 | 0.408 | | -0.447 | 0.655 | |
| | | | | Long | 1.556 | 0.444 | | | | |
| | | D13 | KO | Short | 2.444 | 0.626 | | -1.556 | 0.120 | |
| | | | | Long | 1.333 | 0.333 | | | | |
| | | D14 | KO | Short | 2.444 | 0.503 | | -1.620 | 0.105 | |
| | | | | Long | 1.222 | 0.434 | | | | |
| | | D15 | KO | Short | 2.556 | 0.530 | | -1.982 | 0.047 | * |
| | | | | Long | 1.111 | 0.351 | | | | |
| | | D16 | KO | Short | 1.111 | 0.309 | | -0.744 | 0.457 | |
| | | | | Long | 1.556 | 0.530 | | | | |
| Table S5 | Figure 1H,I / Figure S7 | D0 | WT | Long | 0.000 | 0.000 | Wilcoxon Signed Ranks Test | 0.000 | 1.000 | |
| | | | | Short | 0.000 | 0.000 | | | | |
| | | D1 | WT | Long | 0.167 | 0.118 | | -0.272 | 0.785 | |
| | | | | Short | 0.111 | 0.111 | | | | |
| | | D2 | WT | Long | 0.056 | 0.056 | | -1.841 | 0.066 | |
| | | | | Short | 0.556 | 0.256 | | | | |
| | | D3 | WT | Long | 0.111 | 0.111 | | -2.333 | 0.020 | * |
| | | | | Short | 0.889 | 0.309 | | | | |
| | | D4 | WT | Long | 1.111 | 0.309 | | -2.236 | 0.025 | * |
| | | | | Short | 0.556 | 0.242 | | | | |
| | | D5 | WT | Long | 1.000 | 0.471 | | -0.712 | 0.476 | |
| | | | | Short | 1.556 | 0.530 | | | | |
| | | D6 | WT | Long | 0.778 | 0.222 | | -0.333 | 0.739 | |
| | | | | Short | 0.889 | 0.309 | | | | |
| | | D7 | WT | Long | 0.889 | 0.351 | | -1.035 | 0.301 | |
| | | | | Short | 1.444 | 0.444 | | | | |
| | | D8 | WT | Long | 1.667 | 0.333 | | -1.282 | 0.200 | |
| | | | | Short | 0.778 | 0.324 | | | | |
| | | D9 | WT | Long | 2.000 | 0.577 | | -0.264 | 0.792 | |
| | | | | Short | 1.889 | 0.655 | | | | |
| | | D10 | WT | Short | 1.778 | 0.401 | | -0.289 | 0.773 | |
| | | | | Long | 1.667 | 0.373 | | | | |
| | | D11 | WT | Short | 2.111 | 0.633 | | -1.380 | 0.168 | |
| | | | | Long | 1.333 | 0.236 | | | | |
| | | D12 | WT | Short | 1.889 | 0.200 | | -0.548 | 0.584 | |
| | | | | Long | 1.556 | 0.475 | | | | |
| | | D13 | WT | Short | 2.222 | 0.703 | | -0.736 | 0.462 | |
| | | | | Long | 1.556 | 0.377 | | | | |
| | | D14 | WT | Short | 1.889 | 0.655 | | -0.283 | 0.777 | |
| | | | | Long | 1.556 | 0.377 | | | | |
| | | D15 | WT | Short | 1.889 | 0.676 | | -0.512 | 0.609 | |
| | | | | Long | 1.778 | 0.572 | | | | |
| | | D16 | WT | Short | 2.222 | 0.619 | | -0.718 | 0.473 | |
| | | | | Long | 1.444 | 0.412 | | | | |

## 5_5

HE frequency in 4–5 s window

| Table | Related Figure | Day | Condition | Group | Mean | SEM | Statistical test | Z | P | * |
|---|---|---|---|---|---|---|---|---|---|---|
| Table S5 | Figure 1H,I / Figure S7 | D0 | Long | KO | 0.111 | 0.073 | Mann-Whitney U Test | -1.458 | 0.436 | |
| | | | | WT | 0.000 | 0.000 | | | | |
| | | D1 | Long | KO | 0.167 | 0.118 | | 0.000 | 1.000 | |
| | | | | WT | 0.167 | 0.118 | | | | |
| | | D2 | Long | KO | 0.389 | 0.232 | | -0.616 | 0.666 | |
| | | | | WT | 0.167 | 0.118 | | | | |
| | | D3 | Long | KO | 0.778 | 0.364 | | -0.147 | 0.931 | |
| | | | | WT | 0.667 | 0.333 | | | | |
| | | D4 | Long | KO | 1.000 | 0.441 | | -0.648 | 0.546 | |
| | | | | WT | 1.222 | 0.364 | | | | |
| | | D5 | Long | KO | 0.444 | 0.242 | | -1.719 | 0.113 | |
| | | | | WT | 1.222 | 0.401 | | | | |
| | | D6 | Long | KO | 1.333 | 0.441 | | -0.839 | 0.436 | |
| | | | | WT | 0.889 | 0.423 | | | | |
| | | D7 | Long | KO | 1.444 | 0.377 | | -0.415 | 0.730 | |
| | | | | WT | 1.222 | 0.324 | | | | |
| | | D8 | Long | KO | 2.111 | 0.539 | | -1.300 | 0.222 | |
| | | | | WT | 1.222 | 0.364 | | | | |
| | | D9 | Long | KO | 2.889 | 0.633 | | -1.388 | 0.190 | |
| | | | | WT | 1.667 | 0.624 | | | | |
| | | D10 | Long | KO | 2.444 | 0.818 | | -0.274 | 0.796 | |
| | | | | WT | 1.667 | 0.471 | | | | |
| | | D11 | Long | KO | 2.667 | 0.667 | | -0.813 | 0.436 | |
| | | | | WT | 2.000 | 0.333 | | | | |
| | | D12 | Long | KO | 1.556 | 0.412 | | -0.913 | 0.387 | |
| | | | | WT | 2.556 | 0.766 | | | | |
| | | D13 | Long | KO | 2.000 | 0.850 | | -0.953 | 0.387 | |
| | | | | WT | 2.444 | 0.444 | | | | |
| | | D14 | Long | KO | 1.111 | 0.309 | | -2.137 | 0.040 | * |
| | | | | WT | 2.778 | 0.662 | | | | |
| | | D15 | Long | KO | 1.333 | 0.289 | | -2.320 | 0.024 | * |
| | | | | WT | 2.778 | 0.521 | | | | |
| | | D16 | Long | KO | 1.111 | 0.309 | | -1.799 | 0.094 | |
| | | | | WT | 2.333 | 0.527 | | | | |
| Table S5A | Figure 1H,I / Figure S7 | D0 | Short | KO | 0.222 | 0.121 | Mann-Whitney U Test | -0.318 | 0.796 | |
| | | | | WT | 0.333 | 0.167 | | | | |
| | | D1 | Short | KO | 0.111 | 0.111 | | 0.000 | 1.000 | |
| | | | | WT | 0.111 | 0.111 | | | | |
| | | D2 | Short | KO | 0.111 | 0.111 | | -1.035 | 0.489 | |
| | | | | WT | 0.278 | 0.147 | | | | |
| | | D3 | Short | KO | 0.778 | 0.324 | | -1.060 | 0.387 | |
| | | | | WT | 0.333 | 0.236 | | | | |
| | | D4 | Short | KO | 1.111 | 0.309 | | -1.585 | 0.161 | |
| | | | | WT | 0.444 | 0.242 | | | | |
| | | D5 | Short | KO | 1.222 | 0.324 | | -0.515 | 0.666 | |
| | | | | WT | 1.111 | 0.423 | | | | |
| | | D6 | Short | KO | 1.556 | 0.377 | | -0.229 | 0.863 | |
| | | | | WT | 1.444 | 0.412 | | | | |
| | | D7 | Short | KO | 0.889 | 0.455 | | -0.382 | 0.730 | |
| | | | | WT | 0.889 | 0.309 | | | | |
| | | D8 | Short | KO | 1.111 | 0.389 | | 0.000 | 1.000 | |
| | | | | WT | 1.111 | 0.389 | | | | |
| | | D9 | Short | KO | 0.667 | 0.289 | | -1.617 | 0.136 | |
| | | | | WT | 1.556 | 0.412 | | | | |
| | | D10 | Short | KO | 1.222 | 0.364 | | -0.686 | 0.546 | |
| | | | | WT | 1.778 | 0.521 | | | | |
| | | D11 | Short | KO | 1.222 | 0.547 | | -0.689 | 0.546 | |
| | | | | WT | 1.333 | 0.333 | | | | |
| | | D12 | Short | KO | 0.333 | 0.167 | | -1.738 | 0.113 | |
| | | | | WT | 1.000 | 0.289 | | | | |
| | | D13 | Short | KO | 1.556 | 0.412 | | -1.781 | 0.094 | |
| | | | | WT | 0.556 | 0.242 | | | | |
| | | D14 | Short | KO | 1.111 | 0.633 | | -0.639 | 0.605 | |
| | | | | WT | 0.556 | 0.294 | | | | |
| | | D15 | Short | KO | 0.556 | 0.242 | | -0.407 | 0.730 | |
| | | | | WT | 0.444 | 0.242 | | | | |
| | | D16 | Short | KO | 0.556 | 0.176 | | -0.653 | 0.605 | |
| | | | | WT | 0.444 | 0.242 | | | | |

HE frequency in 4–5 s window

| Table | Related Figure | Day | Group | Condition | Mean | SEM | Statistical test | Z | P | * |
|---|---|---|---|---|---|---|---|---|---|---|
| Table S5 | Figure 1H,I / Figure S7 | D0 | KO | Long | 0.111 | 0.073 | Wilcoxon Signed Ranks Test | -0.816 | 0.414 | |
| | | | | Short | 0.222 | 0.121 | | | | |
| | | D1 | KO | Long | 0.167 | 0.118 | | -1.000 | 0.317 | |
| | | | | Short | 0.111 | 0.111 | | | | |
| | | D2 | KO | Long | 0.389 | 0.232 | | -0.921 | 0.357 | |
| | | | | Short | 0.111 | 0.111 | | | | |
| | | D3 | KO | Long | 0.778 | 0.364 | | 0.000 | 1.000 | |
| | | | | Short | 0.778 | 0.324 | | | | |
| | | D4 | KO | Long | 1.000 | 0.441 | | -0.276 | 0.783 | |
| | | | | Short | 1.111 | 0.309 | | | | |
| | | D5 | KO | Long | 0.444 | 0.242 | | -1.588 | 0.112 | |
| | | | | Short | 1.222 | 0.324 | | | | |
| | | D6 | KO | Long | 1.333 | 0.441 | | -0.439 | 0.660 | |
| | | | | Short | 1.556 | 0.377 | | | | |
| | | D7 | KO | Long | 1.444 | 0.377 | | -1.179 | 0.238 | |
| | | | | Short | 0.889 | 0.455 | | | | |
| | | D8 | KO | Long | 2.111 | 0.539 | | -1.562 | 0.118 | |
| | | | | Short | 1.111 | 0.389 | | | | |
| | | D9 | KO | Long | 2.889 | 0.633 | | -2.207 | 0.027 | * |
| | | | | Short | 0.667 | 0.289 | | | | |
| | | D10 | KO | Short | 2.444 | 0.818 | | -1.380 | 0.168 | |
| | | | | Long | 1.222 | 0.364 | | | | |
| | | D11 | KO | Short | 2.667 | 0.667 | | -1.370 | 0.171 | |
| | | | | Long | 1.222 | 0.547 | | | | |
| | | D12 | KO | Short | 1.556 | 0.412 | | -1.994 | 0.046 | * |
| | | | | Long | 0.333 | 0.167 | | | | |
| | | D13 | KO | Short | 2.000 | 0.850 | | -0.426 | 0.670 | |
| | | | | Long | 1.556 | 0.412 | | | | |
| | | D14 | KO | Short | 1.111 | 0.309 | | -0.531 | 0.595 | |
| | | | | Long | 1.111 | 0.633 | | | | |
| | | D15 | KO | Short | 1.333 | 0.289 | | -1.552 | 0.121 | |
| | | | | Long | 0.556 | 0.242 | | | | |
| | | D16 | KO | Short | 1.111 | 0.309 | | -1.406 | 0.160 | |
| | | | | Long | 0.556 | 0.176 | | | | |
| Table S5 | Figure 1H,I / Figure S7 | D0 | WT | Long | 0.000 | 0.000 | Wilcoxon Signed Ranks Test | -1.732 | 0.083 | |
| | | | | Short | 0.333 | 0.167 | | | | |
| | | D1 | WT | Long | 0.167 | 0.118 | | -0.272 | 0.785 | |
| | | | | Short | 0.111 | 0.111 | | | | |
| | | D2 | WT | Long | 0.167 | 0.118 | | -0.552 | 0.581 | |
| | | | | Short | 0.278 | 0.147 | | | | |
| | | D3 | WT | Long | 0.667 | 0.333 | | -0.707 | 0.480 | |
| | | | | Short | 0.333 | 0.236 | | | | |
| | | D4 | WT | Long | 1.222 | 0.364 | | -1.552 | 0.121 | |
| | | | | Short | 0.444 | 0.242 | | | | |
| | | D5 | WT | Long | 1.222 | 0.401 | | -0.108 | 0.914 | |
| | | | | Short | 1.111 | 0.423 | | | | |
| | | D6 | WT | Long | 0.889 | 0.423 | | -1.179 | 0.238 | |
| | | | | Short | 1.444 | 0.412 | | | | |
| | | D7 | WT | Long | 1.222 | 0.324 | | -0.680 | 0.496 | |
| | | | | Short | 0.889 | 0.309 | | | | |
| | | D8 | WT | Long | 1.222 | 0.364 | | -0.264 | 0.792 | |
| | | | | Short | 1.111 | 0.389 | | | | |
| | | D9 | WT | Long | 1.667 | 0.624 | | -0.141 | 0.888 | |
| | | | | Short | 1.556 | 0.412 | | | | |
| | | D10 | WT | Short | 1.667 | 0.471 | | -0.086 | 0.931 | |
| | | | | Long | 1.778 | 0.521 | | | | |
| | | D11 | WT | Short | 2.000 | 0.333 | | -1.121 | 0.262 | |
| | | | | Long | 1.333 | 0.333 | | | | |
| | | D12 | WT | Short | 2.556 | 0.766 | | -1.693 | 0.090 | |
| | | | | Long | 1.000 | 0.289 | | | | |
| | | D13 | WT | Short | 2.444 | 0.444 | | -2.530 | 0.011 | * |
| | | | | Long | 0.556 | 0.242 | | | | |
| | | D14 | WT | Short | 2.778 | 0.662 | | -2.536 | 0.011 | * |
| | | | | Long | 0.556 | 0.294 | | | | |
| | | D15 | WT | Short | 2.778 | 0.521 | | -2.694 | 0.007 | ** |
| | | | | Long | 0.444 | 0.242 | | | | |
| | | D16 | WT | Short | 2.333 | 0.527 | | -2.116 | 0.034 | * |
| | | | | Long | 0.444 | 0.242 | | | | |

## 5_6

HE frequency in 5–6 s window

| Table | Related Figure | Day | Condition | Group | Mean | SEM | Statistical test | Z | P | * |
|---|---|---|---|---|---|---|---|---|---|---|
| Table S5 | Figure 1H,I / Figure S7 | D0 | Long | KO | 0.333 | 0.144 | Mann-Whitney U Test | -2.187 | 0.113 | |
| | | | | WT | 0.000 | 0.000 | | | | |
| | | D1 | Long | KO | 0.167 | 0.118 | | -0.449 | 0.730 | |
| | | | | WT | 0.222 | 0.121 | | | | |
| | | D2 | Long | KO | 0.111 | 0.073 | | -0.243 | 0.863 | |
| | | | | WT | 0.278 | 0.188 | | | | |
| | | D3 | Long | KO | 0.556 | 0.294 | | -0.687 | 0.546 | |
| | | | | WT | 0.778 | 0.278 | | | | |
| | | D4 | Long | KO | 1.556 | 0.475 | | -1.405 | 0.190 | |
| | | | | WT | 0.667 | 0.289 | | | | |
| | | D5 | Long | KO | 0.889 | 0.423 | | -0.430 | 0.730 | |
| | | | | WT | 1.111 | 0.423 | | | | |
| | | D6 | Long | KO | 1.222 | 0.364 | | -0.417 | 0.730 | |
| | | | | WT | 1.000 | 0.289 | | | | |
| | | D7 | Long | KO | 0.667 | 0.333 | | -1.535 | 0.161 | |
| | | | | WT | 2.111 | 0.696 | | | | |
| | | D8 | Long | KO | 2.222 | 0.572 | | -0.882 | 0.436 | |
| | | | | WT | 1.556 | 0.412 | | | | |
| | | D9 | Long | KO | 2.778 | 0.778 | | -0.769 | 0.489 | |
| | | | | WT | 1.889 | 0.455 | | | | |
| | | D10 | Long | KO | 2.222 | 0.572 | | -0.180 | 0.863 | |
| | | | | WT | 2.111 | 0.564 | | | | |
| | | D11 | Long | KO | 2.333 | 0.471 | | -0.682 | 0.546 | |
| | | | | WT | 1.889 | 0.484 | | | | |
| | | D12 | Long | KO | 1.667 | 0.408 | | -0.092 | 0.931 | |
| | | | | WT | 1.667 | 0.471 | | | | |
| | | D13 | Long | KO | 2.000 | 0.441 | | -0.662 | 0.546 | |
| | | | | WT | 1.778 | 0.641 | | | | |
| | | D14 | Long | KO | 1.333 | 0.471 | | -1.000 | 0.340 | |
| | | | | WT | 1.889 | 0.423 | | | | |
| | | D15 | Long | KO | 2.000 | 0.667 | | -0.639 | 0.546 | |
| | | | | WT | 1.333 | 0.408 | | | | |
| | | D16 | Long | KO | 1.333 | 0.441 | | -0.184 | 0.863 | |
| | | | | WT | 1.444 | 0.626 | | | | |
| Table S5A | Figure 1H,I / Figure S7 | D0 | Short | KO | 0.056 | 0.056 | Mann-Whitney U Test | -1.627 | 0.222 | |
| | | | | WT | 0.333 | 0.144 | | | | |
| | | D1 | Short | KO | 0.278 | 0.147 | | 0.000 | 1.000 | |
| | | | | WT | 0.278 | 0.147 | | | | |
| | | D2 | Short | KO | 0.500 | 0.204 | | -0.911 | 0.436 | |
| | | | | WT | 0.222 | 0.121 | | | | |
| | | D3 | Short | KO | 0.556 | 0.338 | | -0.264 | 0.863 | |
| | | | | WT | 0.889 | 0.455 | | | | |
| | | D4 | Short | KO | 0.444 | 0.176 | | -1.380 | 0.222 | |
| | | | | WT | 1.444 | 0.475 | | | | |
| | | D5 | Short | KO | 0.778 | 0.278 | | -1.101 | 0.340 | |
| | | | | WT | 1.222 | 0.278 | | | | |
| | | D6 | Short | KO | 1.111 | 0.423 | | -1.669 | 0.136 | |
| | | | | WT | 0.444 | 0.338 | | | | |
| | | D7 | Short | KO | 1.333 | 0.236 | | -0.047 | 1.000 | |
| | | | | WT | 1.333 | 0.333 | | | | |
| | | D8 | Short | KO | 1.667 | 0.707 | | -0.232 | 0.863 | |
| | | | | WT | 0.889 | 0.261 | | | | |
| | | D9 | Short | KO | 0.778 | 0.278 | | 0.000 | 1.000 | |
| | | | | WT | 0.778 | 0.278 | | | | |
| | | D10 | Short | KO | 1.444 | 0.475 | | -0.552 | 0.605 | |
| | | | | WT | 1.000 | 0.289 | | | | |
| | | D11 | Short | KO | 0.889 | 0.389 | | -0.949 | 0.436 | |
| | | | | WT | 0.444 | 0.338 | | | | |
| | | D12 | Short | KO | 1.222 | 0.364 | | -0.233 | 0.863 | |
| | | | | WT | 1.333 | 0.373 | | | | |
| | | D13 | Short | KO | 0.778 | 0.278 | | -0.687 | 0.546 | |
| | | | | WT | 0.556 | 0.294 | | | | |
| | | D14 | Short | KO | 0.778 | 0.222 | | -2.004 | 0.063 | |
| | | | | WT | 1.889 | 0.423 | | | | |
| | | D15 | Short | KO | 0.889 | 0.351 | | -0.191 | 0.863 | |
| | | | | WT | 0.778 | 0.324 | | | | |
| | | D16 | Short | KO | 0.556 | 0.176 | | -0.653 | 0.605 | |
| | | | | WT | 0.444 | 0.242 | | | | |

HE frequency in 5–6 s window

| Table | Related Figure | Day | Group | Condition | Mean | SEM | Statistical test | Z | P | * |
|---|---|---|---|---|---|---|---|---|---|---|
| Table S5 | Figure 1H,I / Figure S7 | D0 | KO | Long | 0.333 | 0.144 | Wilcoxon Signed Ranks Test | -1.890 | 0.059 | |
| | | | | Short | 0.056 | 0.056 | | | | |
| | | D1 | KO | Long | 0.167 | 0.118 | | -0.577 | 0.564 | |
| | | | | Short | 0.278 | 0.147 | | | | |
| | | D2 | KO | Long | 0.111 | 0.073 | | -1.841 | 0.066 | |
| | | | | Short | 0.500 | 0.204 | | | | |
| | | D3 | KO | Long | 0.556 | 0.294 | | 0.000 | 1.000 | |
| | | | | Short | 0.556 | 0.338 | | | | |
| | | D4 | KO | Long | 1.556 | 0.475 | | -1.802 | 0.072 | |
| | | | | Short | 0.444 | 0.176 | | | | |
| | | D5 | KO | Long | 0.889 | 0.423 | | -0.378 | 0.705 | |
| | | | | Short | 0.778 | 0.278 | | | | |
| | | D6 | KO | Long | 1.222 | 0.364 | | -0.312 | 0.755 | |
| | | | | Short | 1.111 | 0.423 | | | | |
| | | D7 | KO | Long | 0.667 | 0.333 | | -1.200 | 0.230 | |
| | | | | Short | 1.333 | 0.236 | | | | |
| | | D8 | KO | Long | 2.222 | 0.572 | | -1.667 | 0.096 | |
| | | | | Short | 1.667 | 0.707 | | | | |
| | | D9 | KO | Long | 2.778 | 0.778 | | -2.388 | 0.017 | * |
| | | | | Short | 0.778 | 0.278 | | | | |
| | | D10 | KO | Short | 2.222 | 0.572 | | -1.282 | 0.200 | |
| | | | | Long | 1.444 | 0.475 | | | | |
| | | D11 | KO | Short | 2.333 | 0.471 | | -1.975 | 0.048 | * |
| | | | | Long | 0.889 | 0.389 | | | | |
| | | D12 | KO | Short | 1.667 | 0.408 | | -0.933 | 0.351 | |
| | | | | Long | 1.222 | 0.364 | | | | |
| | | D13 | KO | Short | 2.000 | 0.441 | | -1.980 | 0.048 | * |
| | | | | Long | 0.778 | 0.278 | | | | |
| | | D14 | KO | Short | 1.333 | 0.471 | | -1.089 | 0.276 | |
| | | | | Long | 0.778 | 0.222 | | | | |
| | | D15 | KO | Short | 2.000 | 0.667 | | -1.581 | 0.114 | |
| | | | | Long | 0.889 | 0.351 | | | | |
| | | D16 | KO | Short | 1.333 | 0.441 | | -1.725 | 0.084 | |
| | | | | Long | 0.556 | 0.176 | | | | |
| Table S5 | Figure 1H,I / Figure S7 | D0 | WT | Long | 0.000 | 0.000 | Wilcoxon Signed Ranks Test | -1.857 | 0.063 | |
| | | | | Short | 0.333 | 0.144 | | | | |
| | | D1 | WT | Long | 0.222 | 0.121 | | -0.378 | 0.705 | |
| | | | | Short | 0.278 | 0.147 | | | | |
| | | D2 | WT | Long | 0.278 | 0.188 | | -0.378 | 0.705 | |
| | | | | Short | 0.222 | 0.121 | | | | |
| | | D3 | WT | Long | 0.778 | 0.278 | | 0.000 | 1.000 | |
| | | | | Short | 0.889 | 0.455 | | | | |
| | | D4 | WT | Long | 0.667 | 0.289 | | -1.633 | 0.102 | |
| | | | | Short | 1.444 | 0.475 | | | | |
| | | D5 | WT | Long | 1.111 | 0.423 | | -0.086 | 0.931 | |
| | | | | Short | 1.222 | 0.278 | | | | |
| | | D6 | WT | Long | 1.000 | 0.289 | | -1.179 | 0.238 | |
| | | | | Short | 0.444 | 0.338 | | | | |
| | | D7 | WT | Long | 2.111 | 0.696 | | -1.274 | 0.203 | |
| | | | | Short | 1.333 | 0.333 | | | | |
| | | D8 | WT | Long | 1.556 | 0.412 | | -1.300 | 0.194 | |
| | | | | Short | 0.889 | 0.261 | | | | |
| | | D9 | WT | Long | 1.889 | 0.455 | | -2.157 | 0.031 | * |
| | | | | Short | 0.778 | 0.278 | | | | |
| | | D10 | WT | Short | 2.111 | 0.564 | | -1.590 | 0.112 | |
| | | | | Long | 1.000 | 0.289 | | | | |
| | | D11 | WT | Short | 1.889 | 0.484 | | -2.392 | 0.017 | * |
| | | | | Long | 0.444 | 0.338 | | | | |
| | | D12 | WT | Short | 1.667 | 0.471 | | -0.426 | 0.670 | |
| | | | | Long | 1.333 | 0.373 | | | | |
| | | D13 | WT | Short | 1.778 | 0.641 | | -1.208 | 0.227 | |
| | | | | Long | 0.556 | 0.294 | | | | |
| | | D14 | WT | Short | 1.889 | 0.423 | | -0.137 | 0.891 | |
| | | | | Long | 1.889 | 0.423 | | | | |
| | | D15 | WT | Short | 1.333 | 0.408 | | -0.940 | 0.347 | |
| | | | | Long | 0.778 | 0.324 | | | | |
| | | D16 | WT | Short | 1.444 | 0.626 | | -1.930 | 0.054 | |
| | | | | Long | 0.444 | 0.242 | | | | |

5_7

HE frequency in 6–7 s window

| Table | Related Figure | Day | Condition | Group | Mean | SEM | Stastistical test | Z | P | * |
|---|---|---|---|---|---|---|---|---|---|---|
| Table S5 | Figure 1H,I / Figure S7 | D0 | Long | KO | 0.333 | 0.144 | Mann-Whitney U Test | -1.403 | 0.297 | |
| | | | | WT | 0.111 | 0.111 | | | | |
| | | D1 | Long | KO | 0.056 | 0.056 | | -0.081 | 1.000 | |
| | | | | WT | 0.111 | 0.111 | | | | |
| | | D2 | Long | KO | 0.444 | 0.227 | | -1.626 | 0.222 | |
| | | | | WT | 0.056 | 0.056 | | | | |
| | | D3 | Long | KO | 1.111 | 0.309 | | -1.973 | 0.094 | |
| | | | | WT | 0.333 | 0.333 | | | | |
| | | D4 | Long | KO | 0.667 | 0.289 | | -0.664 | 0.546 | |
| | | | | WT | 0.889 | 0.261 | | | | |
| | | D5 | Long | KO | 0.667 | 0.236 | | -0.646 | 0.605 | |
| | | | | WT | 0.444 | 0.176 | | | | |
| | | D6 | Long | KO | 2.111 | 0.611 | | -1.414 | 0.190 | |
| | | | | WT | 1.000 | 0.333 | | | | |
| | | D7 | Long | KO | 0.778 | 0.324 | | -1.894 | 0.077 | |
| | | | | WT | 1.667 | 0.333 | | | | |
| | | D8 | Long | KO | 2.556 | 0.530 | | -1.227 | 0.258 | |
| | | | | WT | 2.000 | 0.289 | | | | |
| | | D9 | Long | KO | 2.111 | 0.633 | | -1.796 | 0.094 | |
| | | | | WT | 1.111 | 0.696 | | | | |
| | | D10 | Long | KO | 1.444 | 0.338 | | -1.633 | 0.136 | |
| | | | | WT | 0.667 | 0.289 | | | | |
| | | D11 | Long | KO | 2.111 | 0.564 | | -0.225 | 0.863 | |
| | | | | WT | 2.222 | 0.778 | | | | |
| | | D12 | Long | KO | 1.556 | 0.444 | | -0.727 | 0.489 | |
| | | | | WT | 2.333 | 0.553 | | | | |
| | | D13 | Long | KO | 1.000 | 0.373 | | -1.324 | 0.222 | |
| | | | | WT | 1.889 | 0.512 | | | | |
| | | D14 | Long | KO | 1.667 | 0.687 | | -0.861 | 0.436 | |
| | | | | WT | 2.000 | 0.500 | | | | |
| | | D15 | Long | KO | 1.444 | 0.475 | | -0.691 | 0.546 | |
| | | | | WT | 2.333 | 0.745 | | | | |
| | | D16 | Long | KO | 1.667 | 0.500 | | -0.456 | 0.666 | |
| | | | | WT | 1.333 | 0.408 | | | | |
| Table S5A | Figure 1H,I / Figure S7 | D0 | Short | KO | 0.444 | 0.194 | Mann-Whitney U Test | -1.514 | 0.258 | |
| | | | | WT | 0.111 | 0.111 | | | | |
| | | D1 | Short | KO | 0.056 | 0.056 | | -1.214 | 0.387 | |
| | | | | WT | 0.333 | 0.186 | | | | |
| | | D2 | Short | KO | 0.611 | 0.182 | | -2.152 | 0.050 | |
| | | | | WT | 0.111 | 0.073 | | | | |
| | | D3 | Short | KO | 0.889 | 0.351 | | -0.287 | 0.796 | |
| | | | | WT | 0.889 | 0.455 | | | | |
| | | D4 | Short | KO | 0.667 | 0.289 | | -0.336 | 0.796 | |
| | | | | WT | 0.778 | 0.278 | | | | |
| | | D5 | Short | KO | 0.667 | 0.289 | | -0.145 | 0.931 | |
| | | | | WT | 0.667 | 0.236 | | | | |
| | | D6 | Short | KO | 1.444 | 0.626 | | -0.139 | 0.931 | |
| | | | | WT | 0.889 | 0.261 | | | | |
| | | D7 | Short | KO | 0.667 | 0.289 | | -0.098 | 0.931 | |
| | | | | WT | 0.889 | 0.455 | | | | |
| | | D8 | Short | KO | 1.333 | 0.373 | | -1.181 | 0.297 | |
| | | | | WT | 0.778 | 0.324 | | | | |
| | | D9 | Short | KO | 1.778 | 0.465 | | -0.732 | 0.489 | |
| | | | | WT | 1.222 | 0.364 | | | | |
| | | D10 | Short | KO | 0.889 | 0.351 | | -0.837 | 0.436 | |
| | | | | WT | 1.222 | 0.324 | | | | |
| | | D11 | Short | KO | 1.111 | 0.455 | | -1.464 | 0.222 | |
| | | | | WT | 0.333 | 0.236 | | | | |
| | | D12 | Short | KO | 0.778 | 0.364 | | -0.202 | 0.863 | |
| | | | | WT | 1.000 | 0.601 | | | | |
| | | D13 | Short | KO | 0.556 | 0.242 | | -0.960 | 0.387 | |
| | | | | WT | 1.000 | 0.333 | | | | |
| | | D14 | Short | KO | 0.778 | 0.222 | | -0.777 | 0.489 | |
| | | | | WT | 0.556 | 0.242 | | | | |
| | | D15 | Short | KO | 0.444 | 0.242 | | -0.607 | 0.605 | |
| | | | | WT | 0.889 | 0.455 | | | | |
| | | D16 | Short | KO | 0.333 | 0.167 | | -0.160 | 0.931 | |
| | | | | WT | 0.444 | 0.242 | | | | |

HE frequency in 6–7 s window

| Table | Related Figure | Day | Group | Condition | Mean | SEM | Stastistical test | Z | P | * |
|---|---|---|---|---|---|---|---|---|---|---|
| Table S5 | Figure 1H,I / Figure S7 | D0 | KO | Long | 0.333 | 0.144 | Wilcoxon Signed Ranks Test | -0.649 | 0.516 | |
| | | | | Short | 0.444 | 0.194 | | | | |
| | | D1 | KO | Long | 0.056 | 0.056 | | 0.000 | 1.000 | |
| | | | | Short | 0.056 | 0.056 | | | | |
| | | D2 | KO | Long | 0.444 | 0.227 | | -0.756 | 0.450 | |
| | | | | Short | 0.611 | 0.182 | | | | |
| | | D3 | KO | Long | 1.111 | 0.309 | | -0.531 | 0.595 | |
| | | | | Short | 0.889 | 0.351 | | | | |
| | | D4 | KO | Long | 0.667 | 0.289 | | -0.108 | 0.914 | |
| | | | | Short | 0.667 | 0.289 | | | | |
| | | D5 | KO | Long | 0.667 | 0.236 | | 0.000 | 1.000 | |
| | | | | Short | 0.667 | 0.289 | | | | |
| | | D6 | KO | Long | 2.111 | 0.611 | | -0.850 | 0.395 | |
| | | | | Short | 1.444 | 0.626 | | | | |
| | | D7 | KO | Long | 0.778 | 0.324 | | -0.108 | 0.914 | |
| | | | | Short | 0.667 | 0.289 | | | | |
| | | D8 | KO | Long | 2.556 | 0.530 | | -1.980 | 0.048 | |
| | | | | Short | 1.333 | 0.373 | | | | |
| | | D9 | KO | Long | 2.111 | 0.633 | | -0.345 | 0.730 | |
| | | | | Long | 1.778 | 0.465 | | | | |
| | | D10 | KO | Short | 1.444 | 0.338 | | -1.222 | 0.222 | |
| | | | | Long | 0.889 | 0.351 | | | | |
| | | D11 | KO | Short | 2.111 | 0.564 | | -1.807 | 0.071 | |
| | | | | Long | 1.111 | 0.455 | | | | |
| | | D12 | KO | Short | 1.556 | 0.444 | | -1.282 | 0.200 | |
| | | | | Long | 0.778 | 0.364 | | | | |
| | | D13 | KO | Short | 1.000 | 0.373 | | -0.973 | 0.330 | |
| | | | | Long | 0.556 | 0.242 | | | | |
| | | D14 | KO | Short | 1.667 | 0.687 | | -1.186 | 0.236 | |
| | | | | Long | 0.778 | 0.222 | | | | |
| | | D15 | KO | Short | 1.444 | 0.475 | | -1.807 | 0.071 | |
| | | | | Long | 0.444 | 0.242 | | | | |
| | | D16 | KO | Short | 1.667 | 0.500 | | -2.060 | 0.039 | * |
| | | | | Long | 0.333 | 0.167 | | | | |
| Table S5 | Figure 1H,I / Figure S7 | D0 | WT | Long | 0.111 | 0.111 | Wilcoxon Signed Ranks Test | 0.000 | 1.000 | |
| | | | | Short | 0.111 | 0.111 | | | | |
| | | D1 | WT | Long | 0.111 | 0.111 | | -0.921 | 0.357 | |
| | | | | Short | 0.333 | 0.186 | | | | |
| | | D2 | WT | Long | 0.056 | 0.056 | | -0.577 | 0.564 | |
| | | | | Short | 0.111 | 0.073 | | | | |
| | | D3 | WT | Long | 0.333 | 0.333 | | -0.921 | 0.357 | |
| | | | | Short | 0.889 | 0.455 | | | | |
| | | D4 | WT | Long | 0.889 | 0.261 | | -0.447 | 0.655 | |
| | | | | Short | 0.778 | 0.278 | | | | |
| | | D5 | WT | Long | 0.444 | 0.176 | | -0.707 | 0.480 | |
| | | | | Short | 0.667 | 0.236 | | | | |
| | | D6 | WT | Long | 1.000 | 0.333 | | -0.108 | 0.914 | |
| | | | | Short | 0.889 | 0.261 | | | | |
| | | D7 | WT | Long | 1.667 | 0.333 | | -1.329 | 0.184 | |
| | | | | Short | 0.889 | 0.455 | | | | |
| | | D8 | WT | Long | 2.000 | 0.289 | | -2.414 | 0.016 | * |
| | | | | Short | 0.778 | 0.324 | | | | |
| | | D9 | WT | Long | 1.111 | 0.696 | | -0.271 | 0.786 | |
| | | | | Long | 1.222 | 0.364 | | | | |
| | | D10 | WT | Short | 0.667 | 0.289 | | -1.311 | 0.190 | |
| | | | | Long | 1.222 | 0.324 | | | | |
| | | D11 | WT | Short | 2.222 | 0.778 | | -2.214 | 0.027 | * |
| | | | | Long | 0.333 | 0.236 | | | | |
| | | D12 | WT | Short | 2.333 | 0.553 | | -1.257 | 0.209 | |
| | | | | Long | 1.000 | 0.601 | | | | |
| | | D13 | WT | Short | 1.889 | 0.512 | | -1.098 | 0.272 | |
| | | | | Long | 1.000 | 0.333 | | | | |
| | | D14 | WT | Short | 2.000 | 0.500 | | -2.136 | 0.033 | * |
| | | | | Long | 0.556 | 0.242 | | | | |
| | | D15 | WT | Short | 2.333 | 0.745 | | -1.715 | 0.086 | |
| | | | | Long | 0.889 | 0.455 | | | | |
| | | D16 | WT | Short | 1.333 | 0.408 | | -1.552 | 0.121 | |
| | | | | Long | 0.444 | 0.242 | | | | |

5_8

HE frequency in 7–8 s window

| Table | Related Figure | Day | Condition | Group | Mean | SEM | Stastistical test | Z | P | * |
|---|---|---|---|---|---|---|---|---|---|---|
| Table S5 | Figure 1H,I / Figure S7 | D0 | Long | KO | 0.167 | 0.167 | Mann-Whitney U Test | 0.000 | 1.000 | |
| | | | | WT | 0.222 | 0.222 | | | | |
| | | D1 | Long | KO | 0.222 | 0.147 | | -1.000 | 0.730 | |
| | | | | WT | 0.000 | 0.000 | | | | |
| | | D2 | Long | KO | 0.389 | 0.232 | | -1.837 | 0.258 | |
| | | | | WT | 0.000 | 0.000 | | | | |
| | | D3 | Long | KO | 0.222 | 0.147 | | -1.055 | 0.387 | |
| | | | | WT | 0.333 | 0.167 | | | | |
| | | D4 | Long | KO | 0.889 | 0.200 | | -0.520 | 0.666 | |
| | | | | WT | 0.667 | 0.236 | | | | |
| | | D5 | Long | KO | 0.889 | 0.423 | | -0.646 | 0.605 | |
| | | | | WT | 0.444 | 0.176 | | | | |
| | | D6 | Long | KO | 1.000 | 0.333 | | -0.972 | 0.436 | |
| | | | | WT | 0.444 | 0.176 | | | | |
| | | D7 | Long | KO | 1.444 | 0.603 | | -0.627 | 0.605 | |
| | | | | WT | 0.889 | 0.351 | | | | |
| | | D8 | Long | KO | 2.222 | 0.662 | | -0.745 | 0.489 | |
| | | | | WT | 1.778 | 0.641 | | | | |
| | | D9 | Long | KO | 1.667 | 0.373 | | -0.138 | 0.931 | |
| | | | | WT | 1.333 | 0.471 | | | | |
| | | D10 | Long | KO | 1.778 | 0.662 | | -0.738 | 0.489 | |
| | | | | WT | 1.556 | 0.556 | | | | |
| | | D11 | Long | KO | 0.889 | 0.423 | | -0.320 | 0.796 | |
| | | | | WT | 1.778 | 0.683 | | | | |
| | | D12 | Long | KO | 1.889 | 0.564 | | -1.226 | 0.258 | |
| | | | | WT | 1.444 | 0.444 | | | | |
| | | D13 | Long | KO | 1.667 | 0.500 | | -0.749 | 0.489 | |
| | | | | WT | 1.333 | 0.471 | | | | |
| | | D14 | Long | KO | 1.889 | 0.512 | | -2.237 | 0.031 | * |
| | | | | WT | 2.333 | 0.373 | | | | |
| | | D15 | Long | KO | 0.889 | 0.389 | | -0.949 | 0.387 | |
| | | | | WT | 2.000 | 0.408 | | | | |
| | | D16 | Long | KO | 1.667 | 0.471 | | -1.660 | 0.113 | |
| | | | | WT | 2.111 | 0.633 | | | | |
| Table S5A | Figure 1H,I / Figure S7 | D0 | Short | KO | 0.333 | 0.118 | Mann-Whitney U Test | -1.642 | 0.190 | |
| | | | | WT | 0.222 | 0.222 | | | | |
| | | D1 | Short | KO | 0.389 | 0.200 | | -0.620 | 0.666 | |
| | | | | WT | 0.056 | 0.056 | | | | |
| | | D2 | Short | KO | 0.611 | 0.232 | | -0.879 | 0.436 | |
| | | | | WT | 0.222 | 0.121 | | | | |
| | | D3 | Short | KO | 0.444 | 0.242 | | -0.620 | 0.666 | |
| | | | | WT | 0.778 | 0.324 | | | | |
| | | D4 | Short | KO | 0.889 | 0.200 | | 0.000 | 1.000 | |
| | | | | WT | 1.000 | 0.373 | | | | |
| | | D5 | Short | KO | 0.889 | 0.261 | | -0.525 | 0.666 | |
| | | | | WT | 0.667 | 0.236 | | | | |
| | | D6 | Short | KO | 0.778 | 0.278 | | -0.474 | 0.666 | |
| | | | | WT | 0.222 | 0.147 | | | | |
| | | D7 | Short | KO | 1.333 | 0.527 | | -0.092 | 0.931 | |
| | | | | WT | 0.556 | 0.242 | | | | |
| | | D8 | Short | KO | 1.222 | 0.641 | | -1.321 | 0.222 | |
| | | | | WT | 1.333 | 0.687 | | | | |
| | | D9 | Short | KO | 1.000 | 0.667 | | -1.684 | 0.113 | |
| | | | | WT | 1.667 | 0.799 | | | | |
| | | D10 | Short | KO | 0.556 | 0.294 | | -1.639 | 0.136 | |
| | | | | WT | 0.889 | 0.261 | | | | |
| | | D11 | Short | KO | 1.000 | 0.333 | | -0.567 | 0.605 | |
| | | | | WT | 1.444 | 0.626 | | | | |
| | | D12 | Short | KO | 1.333 | 0.577 | | -0.869 | 0.436 | |
| | | | | WT | 0.778 | 0.324 | | | | |
| | | D13 | Short | KO | 1.000 | 0.553 | | -1.388 | 0.190 | |
| | | | | WT | 1.000 | 0.500 | | | | |
| | | D14 | Short | KO | 1.111 | 0.455 | | -1.139 | 0.297 | |
| | | | | WT | 1.111 | 0.423 | | | | |
| | | D15 | Short | KO | 1.000 | 0.333 | | -0.422 | 0.730 | |
| | | | | WT | 1.444 | 0.503 | | | | |
| | | D16 | Short | KO | 0.889 | 0.389 | | -1.286 | 0.222 | |
| | | | | WT | 0.889 | 0.455 | | | | |

HE frequency in 7–8 s window

| Table | Related Figure | Day | Group | Condition | Mean | SEM | Stastistical test | Z | P | * |
|---|---|---|---|---|---|---|---|---|---|---|
| Table S5 | Figure 1H,I / Figure S7 | D0 | KO | Long | 0.167 | 0.167 | Wilcoxon Signed Ranks Test | -1.342 | 0.180 | |
| | | | | Short | 0.333 | 0.118 | | | | |
| | | D1 | KO | Long | 0.222 | 0.147 | | -0.756 | 0.450 | |
| | | | | Short | 0.389 | 0.200 | | | | |
| | | D2 | KO | Long | 0.389 | 0.232 | | -0.816 | 0.414 | |
| | | | | Short | 0.611 | 0.232 | | | | |
| | | D3 | KO | Long | 0.222 | 0.147 | | -1.414 | 0.157 | |
| | | | | Short | 0.444 | 0.242 | | | | |
| | | D4 | KO | Long | 0.889 | 0.200 | | 0.000 | 1.000 | |
| | | | | Short | 0.889 | 0.200 | | | | |
| | | D5 | KO | Long | 0.889 | 0.423 | | 0.000 | 1.000 | |
| | | | | Short | 0.889 | 0.261 | | | | |
| | | D6 | KO | Long | 1.000 | 0.333 | | -0.816 | 0.414 | |
| | | | | Short | 0.778 | 0.278 | | | | |
| | | D7 | KO | Long | 1.444 | 0.603 | | -0.333 | 0.739 | |
| | | | | Short | 1.333 | 0.527 | | | | |
| | | D8 | KO | Long | 2.222 | 0.662 | | -1.807 | 0.071 | |
| | | | | Short | 1.222 | 0.641 | | | | |
| | | D9 | KO | Long | 1.667 | 0.373 | | -0.828 | 0.408 | |
| | | | | Long | 1.000 | 0.667 | | | | |
| | | D10 | KO | Short | 1.778 | 0.662 | | -2.060 | 0.039 | * |
| | | | | Long | 0.556 | 0.294 | | | | |
| | | D11 | KO | Short | 0.889 | 0.423 | | -0.276 | 0.783 | |
| | | | | Long | 1.000 | 0.333 | | | | |
| | | D12 | KO | Short | 1.889 | 0.564 | | -1.289 | 0.197 | |
| | | | | Long | 1.333 | 0.577 | | | | |
| | | D13 | KO | Short | 1.667 | 0.500 | | -2.121 | 0.034 | * |
| | | | | Long | 1.000 | 0.553 | | | | |
| | | D14 | KO | Short | 1.889 | 0.512 | | -1.200 | 0.230 | |
| | | | | Long | 1.111 | 0.455 | | | | |
| | | D15 | KO | Short | 0.889 | 0.389 | | -0.577 | 0.564 | |
| | | | | Long | 1.000 | 0.333 | | | | |
| | | D16 | KO | Short | 1.667 | 0.471 | | -1.890 | 0.059 | |
| | | | | Long | 0.889 | 0.389 | | | | |
| Table S5 | Figure 1H,I / Figure S7 | D0 | WT | Long | 0.222 | 0.222 | Wilcoxon Signed Ranks Test | 0.000 | 1.000 | |
| | | | | Short | 0.222 | 0.222 | | | | |
| | | D1 | WT | Long | 0.000 | 0.000 | | -1.000 | 0.317 | |
| | | | | Short | 0.056 | 0.056 | | | | |
| | | D2 | WT | Long | 0.000 | 0.000 | | -1.633 | 0.102 | |
| | | | | Short | 0.222 | 0.121 | | | | |
| | | D3 | WT | Long | 0.333 | 0.167 | | -1.633 | 0.102 | |
| | | | | Short | 0.778 | 0.324 | | | | |
| | | D4 | WT | Long | 0.667 | 0.236 | | -0.680 | 0.496 | |
| | | | | Short | 1.000 | 0.373 | | | | |
| | | D5 | WT | Long | 0.444 | 0.176 | | -0.707 | 0.480 | |
| | | | | Short | 0.667 | 0.236 | | | | |
| | | D6 | WT | Long | 0.444 | 0.176 | | -1.000 | 0.317 | |
| | | | | Short | 0.222 | 0.147 | | | | |
| | | D7 | WT | Long | 0.889 | 0.351 | | -0.750 | 0.453 | |
| | | | | Short | 0.556 | 0.242 | | | | |
| | | D8 | WT | Long | 1.778 | 0.641 | | -0.850 | 0.395 | |
| | | | | Short | 1.333 | 0.687 | | | | |
| | | D9 | WT | Long | 1.333 | 0.471 | | -0.378 | 0.705 | |
| | | | | Long | 1.667 | 0.799 | | | | |
| | | D10 | WT | Short | 1.556 | 0.556 | | -1.604 | 0.109 | |
| | | | | Long | 0.889 | 0.261 | | | | |
| | | D11 | WT | Short | 1.778 | 0.683 | | -0.816 | 0.414 | |
| | | | | Long | 1.444 | 0.626 | | | | |
| | | D12 | WT | Short | 1.444 | 0.444 | | -1.604 | 0.109 | |
| | | | | Long | 0.778 | 0.324 | | | | |
| | | D13 | WT | Short | 1.333 | 0.471 | | -1.134 | 0.257 | |
| | | | | Long | 1.000 | 0.500 | | | | |
| | | D14 | WT | Short | 2.333 | 0.373 | | -2.414 | 0.016 | * |
| | | | | Long | 1.111 | 0.423 | | | | |
| | | D15 | WT | Short | 2.000 | 0.408 | | -1.414 | 0.157 | |
| | | | | Long | 1.444 | 0.503 | | | | |
| | | D16 | WT | Short | 2.111 | 0.633 | | -1.807 | 0.071 | |
| | | | | Long | 0.889 | 0.455 | | | | |

5_9

HE frequency in 8–9 s window

| Table | Related Figure | Day | Condition | Group | Mean | SEM | Stastistical test | Z | P | * |
|---|---|---|---|---|---|---|---|---|---|---|
| Table S5 | Figure 1H,I / Figure S7 | D0 | Long | KO | 0.000 | 0.000 | Mann-Whitney U Test | -1.000 | 0.730 | |
| | | | | WT | 0.111 | 0.111 | | | | |
| | | D1 | Long | KO | 0.111 | 0.111 | | -1.000 | 0.730 | |
| | | | | WT | 0.000 | 0.000 | | | | |
| | | D2 | Long | KO | 0.167 | 0.118 | | -0.122 | 0.931 | |
| | | | | WT | 0.222 | 0.147 | | | | |
| | | D3 | Long | KO | 0.333 | 0.236 | | -1.455 | 0.436 | |
| | | | | WT | 0.000 | 0.000 | | | | |
| | | D4 | Long | KO | 0.667 | 0.236 | | -0.797 | 0.489 | |
| | | | | WT | 1.222 | 0.434 | | | | |
| | | D5 | Long | KO | 1.111 | 0.588 | | -0.846 | 0.436 | |
| | | | | WT | 1.111 | 0.309 | | | | |
| | | D6 | Long | KO | 1.889 | 0.539 | | -1.799 | 0.094 | |
| | | | | WT | 0.667 | 0.333 | | | | |
| | | D7 | Long | KO | 1.333 | 0.333 | | -1.101 | 0.340 | |
| | | | | WT | 0.889 | 0.351 | | | | |
| | | D8 | Long | KO | 2.111 | 0.423 | | -2.226 | 0.031 | * |
| | | | | WT | 0.778 | 0.278 | | | | |
| | | D9 | Long | KO | 2.444 | 0.556 | | -1.492 | 0.161 | |
| | | | | WT | 1.222 | 0.434 | | | | |
| | | D10 | Long | KO | 1.889 | 0.564 | | -0.187 | 0.863 | |
| | | | | WT | 2.000 | 0.601 | | | | |
| | | D11 | Long | KO | 1.778 | 0.954 | | -0.321 | 0.796 | |
| | | | | WT | 1.444 | 0.475 | | | | |
| | | D12 | Long | KO | 1.444 | 0.556 | | 0.000 | 1.000 | |
| | | | | WT | 1.444 | 0.556 | | | | |
| | | D13 | Long | KO | 2.111 | 0.655 | | -0.677 | 0.546 | |
| | | | | WT | 1.444 | 0.475 | | | | |
| | | D14 | Long | KO | 1.889 | 0.512 | | -0.284 | 0.796 | |
| | | | | WT | 1.778 | 0.521 | | | | |
| | | D15 | Long | KO | 1.222 | 0.596 | | -0.736 | 0.489 | |
| | | | | WT | 1.333 | 0.333 | | | | |
| | | D16 | Long | KO | 1.111 | 0.455 | | -1.059 | 0.340 | |
| | | | | WT | 2.000 | 0.707 | | | | |
| Table S5A | Figure 1H,I / Figure S7 | D0 | Short | KO | 0.222 | 0.121 | Mann-Whitney U Test | -0.620 | 0.666 | |
| | | | | WT | 0.111 | 0.073 | | | | |
| | | D1 | Short | KO | 0.167 | 0.118 | | -0.561 | 0.666 | |
| | | | | WT | 0.278 | 0.147 | | | | |
| | | D2 | Short | KO | 0.556 | 0.227 | | -1.464 | 0.222 | |
| | | | | WT | 0.167 | 0.118 | | | | |
| | | D3 | Short | KO | 0.222 | 0.147 | | -1.910 | 0.094 | |
| | | | | WT | 0.778 | 0.222 | | | | |
| | | D4 | Short | KO | 0.667 | 0.333 | | -0.903 | 0.436 | |
| | | | | WT | 1.333 | 0.500 | | | | |
| | | D5 | Short | KO | 0.444 | 0.176 | | -0.830 | 0.489 | |
| | | | | WT | 0.778 | 0.278 | | | | |
| | | D6 | Short | KO | 1.222 | 0.494 | | -0.472 | 0.666 | |
| | | | | WT | 0.778 | 0.324 | | | | |
| | | D7 | Short | KO | 0.333 | 0.236 | | -1.809 | 0.113 | |
| | | | | WT | 1.111 | 0.351 | | | | |
| | | D8 | Short | KO | 0.778 | 0.324 | | -1.937 | 0.063 | |
| | | | | WT | 1.778 | 0.401 | | | | |
| | | D9 | Short | KO | 1.778 | 0.521 | | -1.943 | 0.063 | |
| | | | | WT | 0.556 | 0.242 | | | | |
| | | D10 | Short | KO | 1.111 | 0.351 | | -0.186 | 0.863 | |
| | | | | WT | 1.111 | 0.423 | | | | |
| | | D11 | Short | KO | 0.778 | 0.324 | | -1.133 | 0.297 | |
| | | | | WT | 1.222 | 0.324 | | | | |
| | | D12 | Short | KO | 0.444 | 0.176 | | 0.000 | 1.000 | |
| | | | | WT | 0.444 | 0.176 | | | | |
| | | D13 | Short | KO | 0.556 | 0.242 | | -0.199 | 0.863 | |
| | | | | WT | 0.556 | 0.176 | | | | |
| | | D14 | Short | KO | 0.667 | 0.236 | | -0.096 | 0.931 | |
| | | | | WT | 0.778 | 0.364 | | | | |
| | | D15 | Short | KO | 0.111 | 0.111 | | -1.102 | 0.436 | |
| | | | | WT | 0.333 | 0.167 | | | | |
| | | D16 | Short | KO | 0.333 | 0.236 | | -1.132 | 0.340 | |
| | | | | WT | 0.556 | 0.176 | | | | |

HE frequency in 8–9 s window

| Table | Related Figure | Day | Group | Condition | Mean | SEM | Stastistical test | Z | P | * |
|---|---|---|---|---|---|---|---|---|---|---|
| Table S5 | Figure 1H,I / Figure S7 | D0 | KO | Long | 0.000 | 0.000 | Wilcoxon Signed Ranks Test | -1.633 | 0.102 | |
| | | | | Short | 0.222 | 0.121 | | | | |
| | | D1 | KO | Long | 0.111 | 0.111 | | -0.447 | 0.655 | |
| | | | | Short | 0.167 | 0.118 | | | | |
| | | D2 | KO | Long | 0.167 | 0.118 | | -1.511 | 0.131 | |
| | | | | Short | 0.556 | 0.227 | | | | |
| | | D3 | KO | Long | 0.333 | 0.236 | | -0.447 | 0.655 | |
| | | | | Short | 0.222 | 0.147 | | | | |
| | | D4 | KO | Long | 0.667 | 0.236 | | 0.000 | 1.000 | |
| | | | | Short | 0.667 | 0.333 | | | | |
| | | D5 | KO | Long | 1.111 | 0.588 | | -0.647 | 0.518 | |
| | | | | Short | 0.444 | 0.176 | | | | |
| | | D6 | KO | Long | 1.889 | 0.539 | | -1.730 | 0.084 | |
| | | | | Short | 1.222 | 0.494 | | | | |
| | | D7 | KO | Long | 1.333 | 0.333 | | -2.309 | 0.021 | * |
| | | | | Short | 0.333 | 0.236 | | | | |
| | | D8 | KO | Long | 2.111 | 0.423 | | -2.111 | 0.035 | * |
| | | | | Short | 0.778 | 0.324 | | | | |
| | | D9 | KO | Long | 2.444 | 0.556 | | -1.561 | 0.119 | |
| | | | | Long | 1.778 | 0.521 | | | | |
| | | D10 | KO | Short | 1.889 | 0.564 | | -0.781 | 0.435 | |
| | | | | Long | 1.111 | 0.351 | | | | |
| | | D11 | KO | Short | 1.778 | 0.954 | | -1.511 | 0.131 | |
| | | | | Long | 0.778 | 0.324 | | | | |
| | | D12 | KO | Short | 1.444 | 0.556 | | -1.807 | 0.071 | |
| | | | | Long | 0.444 | 0.176 | | | | |
| | | D13 | KO | Short | 2.111 | 0.655 | | -2.047 | 0.041 | * |
| | | | | Long | 0.556 | 0.242 | | | | |
| | | D14 | KO | Short | 1.889 | 0.512 | | -2.456 | 0.014 | * |
| | | | | Long | 0.667 | 0.236 | | | | |
| | | D15 | KO | Short | 1.222 | 0.596 | | -1.604 | 0.109 | |
| | | | | Long | 0.111 | 0.111 | | | | |
| | | D16 | KO | Short | 1.111 | 0.455 | | -1.841 | 0.066 | |
| | | | | Long | 0.333 | 0.236 | | | | |
| Table S5 | Figure 1H,I / Figure S7 | D0 | WT | Long | 0.111 | 0.111 | Wilcoxon Signed Ranks Test | 0.000 | 1.000 | |
| | | | | Short | 0.111 | 0.073 | | | | |
| | | D1 | WT | Long | 0.000 | 0.000 | | -1.633 | 0.102 | |
| | | | | Short | 0.278 | 0.147 | | | | |
| | | D2 | WT | Long | 0.222 | 0.147 | | -0.447 | 0.655 | |
| | | | | Short | 0.167 | 0.118 | | | | |
| | | D3 | WT | Long | 0.000 | 0.000 | | -2.333 | 0.020 | * |
| | | | | Short | 0.778 | 0.222 | | | | |
| | | D4 | WT | Long | 1.222 | 0.434 | | -0.333 | 0.739 | |
| | | | | Short | 1.333 | 0.500 | | | | |
| | | D5 | WT | Long | 1.111 | 0.309 | | -0.707 | 0.480 | |
| | | | | Short | 0.778 | 0.278 | | | | |
| | | D6 | WT | Long | 0.667 | 0.333 | | -0.276 | 0.783 | |
| | | | | Short | 0.778 | 0.324 | | | | |
| | | D7 | WT | Long | 0.889 | 0.351 | | -0.513 | 0.608 | |
| | | | | Short | 1.111 | 0.351 | | | | |
| | | D8 | WT | Long | 0.778 | 0.278 | | -1.913 | 0.056 | |
| | | | | Short | 1.778 | 0.401 | | | | |
| | | D9 | WT | Long | 1.222 | 0.434 | | -0.997 | 0.319 | |
| | | | | Long | 0.556 | 0.242 | | | | |
| | | D10 | WT | Short | 2.000 | 0.601 | | -1.613 | 0.107 | |
| | | | | Long | 1.111 | 0.423 | | | | |
| | | D11 | WT | Short | 1.444 | 0.475 | | -0.707 | 0.480 | |
| | | | | Long | 1.222 | 0.324 | | | | |
| | | D12 | WT | Short | 1.444 | 0.556 | | -1.807 | 0.071 | |
| | | | | Long | 0.444 | 0.176 | | | | |
| | | D13 | WT | Short | 1.444 | 0.475 | | -1.492 | 0.136 | |
| | | | | Long | 0.556 | 0.176 | | | | |
| | | D14 | WT | Short | 1.778 | 0.521 | | -1.567 | 0.117 | |
| | | | | Long | 0.778 | 0.364 | | | | |
| | | D15 | WT | Short | 1.333 | 0.333 | | -2.251 | 0.024 | * |
| | | | | Long | 0.333 | 0.167 | | | | |
| | | D16 | WT | Short | 2.000 | 0.707 | | -1.807 | 0.071 | |
| | | | | Long | 0.556 | 0.176 | | | | |

## 5_10

HE frequency in 9–10 s window

| Table | Related Figure | Day | Condition | Group | Mean | SEM | Stastistical test | Z | P | * |
|---|---|---|---|---|---|---|---|---|---|---|
| Table S5 | Figure 1H,I / Figure S7 | D0 | Long | KO | 0.222 | 0.147 | Mann-Whitney U Test | -1.458 | 0.436 | |
| | | | | WT | 0.000 | 0.000 | | | | |
| | | D1 | Long | KO | 0.056 | 0.056 | | -1.636 | 0.19 | |
| | | | | WT | 0.000 | 0.000 | | | | |
| | | D2 | Long | KO | 0.222 | 0.222 | | -0.910 | 0.546 | |
| | | | | WT | 0.444 | 0.155 | | | | |
| | | D3 | Long | KO | 0.222 | 0.147 | | -0.586 | 0.605 | |
| | | | | WT | 0.556 | 0.242 | | | | |
| | | D4 | Long | KO | 0.444 | 0.242 | | -1.628 | 0.136 | |
| | | | | WT | 0.333 | 0.333 | | | | |
| | | D5 | Long | KO | 0.556 | 0.242 | | -0.279 | 0.796 | |
| | | | | WT | 0.667 | 0.236 | | | | |
| | | D6 | Long | KO | 1.000 | 0.289 | | -0.185 | 0.863 | |
| | | | | WT | 1.222 | 0.324 | | | | |
| | | D7 | Long | KO | 0.889 | 0.309 | | -0.482 | 0.666 | |
| | | | | WT | 1.111 | 0.351 | | | | |
| | | D8 | Long | KO | 2.333 | 0.707 | | -0.049 | 1 | |
| | | | | WT | 1.000 | 0.667 | | | | |
| | | D9 | Long | KO | 1.444 | 0.444 | | -1.000 | 0.73 | |
| | | | | WT | 1.778 | 0.760 | | | | |
| | | D10 | Long | KO | 1.222 | 0.572 | | -1.068 | 0.387 | |
| | | | | WT | 1.444 | 0.530 | | | | |
| | | D11 | Long | KO | 1.000 | 0.441 | | -0.391 | 0.73 | |
| | | | | WT | 1.444 | 0.626 | | | | |
| | | D12 | Long | KO | 1.444 | 0.626 | | -0.419 | 0.73 | |
| | | | | WT | 1.778 | 0.778 | | | | |
| | | D13 | Long | KO | 0.556 | 0.294 | | -0.137 | 0.931 | |
| | | | | WT | 1.222 | 0.434 | | | | |
| | | D14 | Long | KO | 0.889 | 0.539 | | -0.327 | 0.796 | |
| | | | | WT | 1.000 | 0.441 | | | | |
| | | D15 | Long | KO | 1.444 | 0.475 | | -1.241 | 0.258 | |
| | | | | WT | 1.667 | 0.667 | | | | |
| | | D16 | Long | KO | 1.000 | 0.500 | | -0.046 | 1 | |
| | | | | WT | 1.111 | 0.588 | | | | |
| Table S5A | Figure 1H,I / Figure S7 | D0 | Short | KO | 0.278 | 0.147 | Mann-Whitney U Test | -0.168 | 0.931 | |
| | | | | WT | 0.444 | 0.306 | | | | |
| | | D1 | Short | KO | 0.778 | 0.334 | | -0.910 | 0.436 | |
| | | | | WT | 0.056 | 0.056 | | | | |
| | | D2 | Short | KO | 0.167 | 0.083 | | -0.819 | 0.489 | |
| | | | | WT | 0.556 | 0.256 | | | | |
| | | D3 | Short | KO | 0.556 | 0.294 | | -0.691 | 0.546 | |
| | | | | WT | 0.222 | 0.147 | | | | |
| | | D4 | Short | KO | 0.778 | 0.465 | | -1.866 | 0.094 | |
| | | | | WT | 0.778 | 0.222 | | | | |
| | | D5 | Short | KO | 0.778 | 0.278 | | -1.338 | 0.222 | |
| | | | | WT | 1.000 | 0.289 | | | | |
| | | D6 | Short | KO | 1.333 | 0.373 | | -1.735 | 0.113 | |
| | | | | WT | 1.000 | 0.373 | | | | |
| | | D7 | Short | KO | 0.667 | 0.167 | | -0.607 | 0.605 | |
| | | | | WT | 1.000 | 0.333 | | | | |
| | | D8 | Short | KO | 1.333 | 0.441 | | -0.615 | 0.73 | |
| | | | | WT | 0.333 | 0.167 | | | | |
| | | D9 | Short | KO | 0.889 | 0.423 | | -2.107 | 0.077 | |
| | | | | WT | 0.889 | 0.309 | | | | |
| | | D10 | Short | KO | 0.556 | 0.338 | | -0.729 | 0.605 | |
| | | | | WT | 1.111 | 0.351 | | | | |
| | | D11 | Short | KO | 1.000 | 0.441 | | -0.563 | 0.605 | |
| | | | | WT | 0.778 | 0.324 | | | | |
| | | D12 | Short | KO | 1.111 | 0.351 | | -0.595 | 0.605 | |
| | | | | WT | 0.333 | 0.167 | | | | |
| | | D13 | Short | KO | 1.000 | 0.441 | | -0.478 | 0.666 | |
| | | | | WT | 1.000 | 0.289 | | | | |
| | | D14 | Short | KO | 0.444 | 0.242 | | -0.239 | 0.863 | |
| | | | | WT | 0.889 | 0.455 | | | | |
| | | D15 | Short | KO | 0.556 | 0.176 | | -0.420 | 0.73 | |
| | | | | WT | 0.222 | 0.222 | | | | |
| | | D16 | Short | KO | 0.222 | 0.147 | | -1.655 | 0.19 | |
| | | | | WT | 0.111 | 0.111 | | | | |

HE frequency in 9–10 s window

| Table | Related Figure | Day | Group | Condition | Mean | SEM | Stastistical test | Z | P | * |
|---|---|---|---|---|---|---|---|---|---|---|
| Table S5 | Figure 1H,I / Figure S7 | D0 | KO | Long | 0.222 | 0.147 | Wilcoxon Signed Ranks Test | -0.141 | 0.888 | |
| | | | | Short | 0.278 | 0.147 | | | | |
| | | D1 | KO | Long | 0.056 | 0.056 | | 0 | 1.000 | |
| | | | | Short | 0.778 | 0.334 | | | | |
| | | D2 | KO | Long | 0.222 | 0.222 | | -0.378 | 0.705 | |
| | | | | Short | 0.167 | 0.083 | | | | |
| | | D3 | KO | Long | 0.222 | 0.147 | | -0.722 | 0.470 | |
| | | | | Short | 0.556 | 0.294 | | | | |
| | | D4 | KO | Long | 0.444 | 0.242 | | -1.604 | 0.109 | |
| | | | | Short | 0.778 | 0.465 | | | | |
| | | D5 | KO | Long | 0.556 | 0.242 | | -1.166 | 0.244 | |
| | | | | Short | 0.778 | 0.278 | | | | |
| | | D6 | KO | Long | 1.000 | 0.289 | | -0.535 | 0.593 | |
| | | | | Short | 1.333 | 0.373 | | | | |
| | | D7 | KO | Long | 0.889 | 0.309 | | -1.134 | 0.257 | |
| | | | | Short | 0.667 | 0.167 | | | | |
| | | D8 | KO | Long | 2.333 | 0.707 | | -1.841 | 0.066 | |
| | | | | Short | 1.333 | 0.441 | | | | |
| | | D9 | KO | Long | 1.444 | 0.444 | | -1.841 | 0.066 | |
| | | | | Long | 0.889 | 0.423 | | | | |
| | | D10 | KO | Short | 1.222 | 0.572 | | -1.134 | 0.257 | |
| | | | | Long | 0.556 | 0.338 | | | | |
| | | D11 | KO | Short | 1.000 | 0.441 | | -0.649 | 0.516 | |
| | | | | Long | 1.000 | 0.441 | | | | |
| | | D12 | KO | Short | 1.444 | 0.626 | | -0.816 | 0.414 | |
| | | | | Long | 1.111 | 0.351 | | | | |
| | | D13 | KO | Short | 0.556 | 0.294 | | -0.957 | 0.339 | |
| | | | | Long | 1.000 | 0.441 | | | | |
| | | D14 | KO | Short | 0.889 | 0.539 | | 0 | 1.000 | |
| | | | | Long | 0.444 | 0.242 | | | | |
| | | D15 | KO | Short | 1.444 | 0.475 | | -0.85 | 0.395 | |
| | | | | Long | 0.556 | 0.176 | | | | |
| | | D16 | KO | Short | 1.000 | 0.500 | | -1.947 | 0.052 | |
| | | | | Long | 0.222 | 0.147 | | | | |
| Table S5 | Figure 1H,I / Figure S7 | D0 | WT | Long | 0.000 | 0.000 | Wilcoxon Signed Ranks Test | -1.342 | 0.180 | |
| | | | | Short | 0.444 | 0.306 | | | | |
| | | D1 | WT | Long | 0.000 | 0.000 | | -0.345 | 0.730 | |
| | | | | Short | 0.056 | 0.056 | | | | |
| | | D2 | WT | Long | 0.444 | 0.155 | | -1.228 | 0.219 | |
| | | | | Short | 0.556 | 0.256 | | | | |
| | | D3 | WT | Long | 0.556 | 0.242 | | -0.425 | 0.671 | |
| | | | | Short | 0.222 | 0.147 | | | | |
| | | D4 | WT | Long | 0.333 | 0.333 | | -0.647 | 0.518 | |
| | | | | Short | 0.778 | 0.222 | | | | |
| | | D5 | WT | Long | 0.667 | 0.236 | | -0.638 | 0.524 | |
| | | | | Short | 1.000 | 0.289 | | | | |
| | | D6 | WT | Long | 1.222 | 0.324 | | -1.807 | 0.071 | |
| | | | | Short | 1.000 | 0.373 | | | | |
| | | D7 | WT | Long | 1.111 | 0.351 | | -0.264 | 0.792 | |
| | | | | Short | 1.000 | 0.333 | | | | |
| | | D8 | WT | Long | 1.000 | 0.667 | | -1.841 | 0.066 | |
| | | | | Short | 0.333 | 0.167 | | | | |
| | | D9 | WT | Long | 1.778 | 0.760 | | -1 | 0.317 | |
| | | | | Long | 0.889 | 0.309 | | | | |
| | | D10 | WT | Short | 1.444 | 0.530 | | -1.732 | 0.083 | |
| | | | | Long | 1.111 | 0.351 | | | | |
| | | D11 | WT | Short | 1.444 | 0.626 | | -0.905 | 0.366 | |
| | | | | Long | 0.778 | 0.324 | | | | |
| | | D12 | WT | Short | 1.778 | 0.778 | | -0.172 | 0.863 | |
| | | | | Long | 0.333 | 0.167 | | | | |
| | | D13 | WT | Short | 1.222 | 0.434 | | -0.954 | 0.340 | |
| | | | | Long | 1.000 | 0.289 | | | | |
| | | D14 | WT | Short | 1.000 | 0.441 | | -0.711 | 0.477 | |
| | | | | Long | 0.889 | 0.455 | | | | |
| | | D15 | WT | Short | 1.667 | 0.667 | | -0.412 | 0.680 | |
| | | | | Long | 0.222 | 0.222 | | | | |
| | | D16 | WT | Short | 1.111 | 0.588 | | -1.807 | 0.071 | |
| | | | | Long | 0.111 | 0.111 | | | | |

## 5_11

HE frequency in 10–11 s window

| Table | Related Figure | Day | Condition | Group | Mean | SEM | Stastistical test | Z | P | * |
|---|---|---|---|---|---|---|---|---|---|---|
| Table S5 | Figure 1H,I / Figure S7 | D0 | Long | KO | 0.111 | 0.073 | Mann-Whitney U Test | -0.243 | 0.863 | |
| | | | | WT | 0.556 | 0.444 | | | | |
| | | D1 | Long | KO | 0.111 | 0.111 | | -1.000 | 0.730 | |
| | | | | WT | 0.000 | 0.000 | | | | |
| | | D2 | Long | KO | 0.222 | 0.121 | | -1.837 | 0.258 | |
| | | | | WT | 0.000 | 0.000 | | | | |
| | | D3 | Long | KO | 0.222 | 0.222 | | -1.092 | 0.436 | |
| | | | | WT | 0.667 | 0.373 | | | | |
| | | D4 | Long | KO | 1.000 | 0.471 | | -0.605 | 0.605 | |
| | | | | WT | 0.667 | 0.441 | | | | |
| | | D5 | Long | KO | 1.111 | 0.389 | | -1.196 | 0.297 | |
| | | | | WT | 0.444 | 0.176 | | | | |
| | | D6 | Long | KO | 2.000 | 0.726 | | -1.862 | 0.113 | |
| | | | | WT | 0.222 | 0.147 | | | | |
| | | D7 | Long | KO | 1.333 | 0.527 | | -0.558 | 0.605 | |
| | | | | WT | 1.000 | 0.441 | | | | |
| | | D8 | Long | KO | 1.667 | 0.408 | | -1.278 | 0.222 | |
| | | | | WT | 1.000 | 0.441 | | | | |
| | | D9 | Long | KO | 1.667 | 0.707 | | -0.413 | 0.730 | |
| | | | | WT | 2.111 | 0.904 | | | | |
| | | D10 | Long | KO | 1.000 | 0.471 | | -0.792 | 0.489 | |
| | | | | WT | 1.444 | 0.475 | | | | |
| | | D11 | Long | KO | 0.778 | 0.364 | | -1.392 | 0.190 | |
| | | | | WT | 2.333 | 0.833 | | | | |
| | | D12 | Long | KO | 0.778 | 0.364 | | -1.249 | 0.258 | |
| | | | | WT | 1.333 | 0.333 | | | | |
| | | D13 | Long | KO | 1.556 | 0.603 | | -0.230 | 0.863 | |
| | | | | WT | 1.556 | 0.530 | | | | |
| | | D14 | Long | KO | 0.778 | 0.434 | | -0.733 | 0.546 | |
| | | | | WT | 1.111 | 0.423 | | | | |
| | | D15 | Long | KO | 0.444 | 0.242 | | -1.006 | 0.387 | |
| | | | | WT | 1.667 | 0.707 | | | | |
| | | D16 | Long | KO | 1.333 | 0.471 | | -0.279 | 0.796 | |
| | | | | WT | 1.889 | 0.772 | | | | |
| Table S5A | Figure 1H,I / Figure S7 | D0 | Short | KO | 0.278 | 0.121 | Mann-Whitney U Test | -1.068 | 0.387 | |
| | | | | WT | 0.111 | 0.073 | | | | |
| | | D1 | Short | KO | 0.389 | 0.200 | | -0.620 | 0.666 | |
| | | | | WT | 0.222 | 0.147 | | | | |
| | | D2 | Short | KO | 0.778 | 0.355 | | -1.896 | 0.113 | |
| | | | | WT | 0.111 | 0.111 | | | | |
| | | D3 | Short | KO | 0.778 | 0.364 | | -0.454 | 0.730 | |
| | | | | WT | 0.556 | 0.294 | | | | |
| | | D4 | Short | KO | 0.444 | 0.176 | | -0.646 | 0.605 | |
| | | | | WT | 0.667 | 0.236 | | | | |
| | | D5 | Short | KO | 0.333 | 0.236 | | -0.952 | 0.436 | |
| | | | | WT | 0.667 | 0.289 | | | | |
| | | D6 | Short | KO | 1.000 | 0.333 | | -0.756 | 0.489 | |
| | | | | WT | 0.667 | 0.289 | | | | |
| | | D7 | Short | KO | 1.111 | 0.423 | | -0.098 | 0.931 | |
| | | | | WT | 1.111 | 0.389 | | | | |
| | | D8 | Short | KO | 0.556 | 0.176 | | -0.049 | 1.000 | |
| | | | | WT | 1.000 | 0.500 | | | | |
| | | D9 | Short | KO | 0.778 | 0.364 | | -0.147 | 0.931 | |
| | | | | WT | 0.667 | 0.333 | | | | |
| | | D10 | Short | KO | 1.000 | 0.289 | | -1.196 | 0.297 | |
| | | | | WT | 0.778 | 0.434 | | | | |
| | | D11 | Short | KO | 0.778 | 0.324 | | -0.657 | 0.546 | |
| | | | | WT | 1.000 | 0.236 | | | | |
| | | D12 | Short | KO | 1.000 | 0.289 | | -1.738 | 0.113 | |
| | | | | WT | 0.333 | 0.167 | | | | |
| | | D13 | Short | KO | 0.333 | 0.236 | | -0.680 | 0.666 | |
| | | | | WT | 0.111 | 0.111 | | | | |
| | | D14 | Short | KO | 0.667 | 0.236 | | -0.096 | 0.931 | |
| | | | | WT | 0.778 | 0.364 | | | | |
| | | D15 | Short | KO | 0.333 | 0.167 | | -0.160 | 0.931 | |
| | | | | WT | 0.444 | 0.242 | | | | |
| | | D16 | Short | KO | 0.222 | 0.147 | | -1.982 | 0.077 | |
| | | | | WT | 1.000 | 0.333 | | | | |

HE frequency in 10–11 s window

| Table | Related Figure | Day | Group | Condition | Mean | SEM | Stastistical test | Z | P | * |
|---|---|---|---|---|---|---|---|---|---|---|
| Table S5 | Figure 1H,I / Figure S7 | D0 | KO | Long | 0.111 | 0.073 | Wilcoxon Signed Ranks Test | -1.000 | 0.317 | |
| | | | | Short | 0.278 | 0.121 | | | | |
| | | D1 | KO | Long | 0.111 | 0.111 | | -1.633 | 0.102 | |
| | | | | Short | 0.389 | 0.200 | | | | |
| | | D2 | KO | Long | 0.222 | 0.121 | | -1.355 | 0.176 | |
| | | | | Short | 0.778 | 0.355 | | | | |
| | | D3 | KO | Long | 0.222 | 0.222 | | -1.089 | 0.276 | |
| | | | | Short | 0.778 | 0.364 | | | | |
| | | D4 | KO | Long | 1.000 | 0.471 | | -1.225 | 0.221 | |
| | | | | Short | 0.444 | 0.176 | | | | |
| | | D5 | KO | Long | 1.111 | 0.389 | | -1.594 | 0.111 | |
| | | | | Short | 0.333 | 0.236 | | | | |
| | | D6 | KO | Long | 2.000 | 0.726 | | -1.518 | 0.129 | |
| | | | | Short | 1.000 | 0.333 | | | | |
| | | D7 | KO | Long | 1.333 | 0.527 | | -0.816 | 0.414 | |
| | | | | Short | 1.111 | 0.423 | | | | |
| | | D8 | KO | Long | 1.667 | 0.408 | | -1.983 | 0.047 | * |
| | | | | Short | 0.556 | 0.176 | | | | |
| | | D9 | KO | Long | 1.667 | 0.707 | | -1.160 | 0.246 | |
| | | | | Long | 0.778 | 0.364 | | | | |
| | | D10 | KO | Short | 1.000 | 0.471 | | 0.000 | 1.000 | |
| | | | | Long | 1.000 | 0.289 | | | | |
| | | D11 | KO | Short | 0.778 | 0.364 | | 0.000 | 1.000 | |
| | | | | Long | 0.778 | 0.324 | | | | |
| | | D12 | KO | Short | 0.778 | 0.364 | | -0.649 | 0.516 | |
| | | | | Long | 1.000 | 0.289 | | | | |
| | | D13 | KO | Short | 1.556 | 0.603 | | -2.232 | 0.026 | * |
| | | | | Long | 0.333 | 0.236 | | | | |
| | | D14 | KO | Short | 0.778 | 0.434 | | -0.378 | 0.705 | |
| | | | | Long | 0.667 | 0.236 | | | | |
| | | D15 | KO | Short | 0.444 | 0.242 | | -0.577 | 0.564 | |
| | | | | Long | 0.333 | 0.167 | | | | |
| | | D16 | KO | Short | 1.333 | 0.471 | | -1.715 | 0.086 | |
| | | | | Long | 0.222 | 0.147 | | | | |
| Table S5 | Figure 1H,I / Figure S7 | D0 | WT | Long | 0.556 | 0.444 | Wilcoxon Signed Ranks Test | -0.736 | 0.461 | |
| | | | | Short | 0.111 | 0.073 | | | | |
| | | D1 | WT | Long | 0.000 | 0.000 | | -1.414 | 0.157 | |
| | | | | Short | 0.222 | 0.147 | | | | |
| | | D2 | WT | Long | 0.000 | 0.000 | | -1.000 | 0.317 | |
| | | | | Short | 0.111 | 0.111 | | | | |
| | | D3 | WT | Long | 0.667 | 0.373 | | -0.184 | 0.854 | |
| | | | | Short | 0.556 | 0.294 | | | | |
| | | D4 | WT | Long | 0.667 | 0.441 | | -0.378 | 0.705 | |
| | | | | Short | 0.667 | 0.236 | | | | |
| | | D5 | WT | Long | 0.444 | 0.176 | | -0.707 | 0.480 | |
| | | | | Short | 0.667 | 0.289 | | | | |
| | | D6 | WT | Long | 0.222 | 0.147 | | -1.300 | 0.194 | |
| | | | | Short | 0.667 | 0.289 | | | | |
| | | D7 | WT | Long | 1.000 | 0.441 | | -0.447 | 0.655 | |
| | | | | Short | 1.111 | 0.389 | | | | |
| | | D8 | WT | Long | 1.000 | 0.441 | | -0.333 | 0.739 | |
| | | | | Short | 1.000 | 0.500 | | | | |
| | | D9 | WT | Long | 2.111 | 0.904 | | -1.913 | 0.056 | |
| | | | | Long | 0.667 | 0.333 | | | | |
| | | D10 | WT | Short | 1.444 | 0.475 | | -1.089 | 0.276 | |
| | | | | Long | 0.778 | 0.434 | | | | |
| | | D11 | WT | Short | 2.333 | 0.833 | | -1.294 | 0.196 | |
| | | | | Long | 1.000 | 0.236 | | | | |
| | | D12 | WT | Short | 1.333 | 0.333 | | -2.251 | 0.024 | * |
| | | | | Long | 0.333 | 0.167 | | | | |
| | | D13 | WT | Short | 1.556 | 0.530 | | -2.226 | 0.026 | * |
| | | | | Long | 0.111 | 0.111 | | | | |
| | | D14 | WT | Short | 1.111 | 0.423 | | -1.342 | 0.180 | |
| | | | | Long | 0.778 | 0.364 | | | | |
| | | D15 | WT | Short | 1.667 | 0.707 | | -1.581 | 0.114 | |
| | | | | Long | 0.444 | 0.242 | | | | |
| | | D16 | WT | Short | 1.889 | 0.772 | | -0.679 | 0.497 | |
| | | | | Long | 1.000 | 0.333 | | | | |

## 5_12

HE frequency in 11–12 s window

| Table | Related Figure | Day | Condition | Group | Mean | SEM | Stastistical test | Z | P | * |
|---|---|---|---|---|---|---|---|---|---|---|
| Table S5 | Figure 1H,I / Figure S7 | D0 | Long | KO | 0.278 | 0.188 | Mann-Whitney U Test | -0.476 | 0.796 | |
| | | | | WT | 0.444 | 0.444 | | | | |
| | | D1 | Long | KO | 0.111 | 0.073 | | -0.122 | 0.931 | |
| | | | | WT | 0.167 | 0.118 | | | | |
| | | D2 | Long | KO | 0.222 | 0.147 | | -0.281 | 0.863 | |
| | | | | WT | 0.222 | 0.121 | | | | |
| | | D3 | Long | KO | 0.222 | 0.147 | | 0.000 | 1.000 | |
| | | | | WT | 0.222 | 0.147 | | | | |
| | | D4 | Long | KO | 0.778 | 0.434 | | -0.098 | 0.931 | |
| | | | | WT | 0.667 | 0.289 | | | | |
| | | D5 | Long | KO | 1.556 | 0.475 | | -1.483 | 0.190 | |
| | | | | WT | 0.667 | 0.373 | | | | |
| | | D6 | Long | KO | 0.667 | 0.441 | | -0.053 | 1.000 | |
| | | | | WT | 0.444 | 0.242 | | | | |
| | | D7 | Long | KO | 1.111 | 0.351 | | -0.279 | 0.796 | |
| | | | | WT | 1.222 | 0.547 | | | | |
| | | D8 | Long | KO | 2.111 | 0.633 | | -1.231 | 0.258 | |
| | | | | WT | 1.111 | 0.423 | | | | |
| | | D9 | Long | KO | 2.333 | 0.833 | | -0.225 | 0.863 | |
| | | | | WT | 1.778 | 0.547 | | | | |
| | | D10 | Long | KO | 1.000 | 0.500 | | -0.476 | 0.666 | |
| | | | | WT | 1.333 | 0.471 | | | | |
| | | D11 | Long | KO | 0.778 | 0.364 | | -0.098 | 0.931 | |
| | | | | WT | 0.889 | 0.423 | | | | |
| | | D12 | Long | KO | 1.000 | 0.667 | | -0.252 | 0.863 | |
| | | | | WT | 0.667 | 0.289 | | | | |
| | | D13 | Long | KO | 0.556 | 0.294 | | -0.355 | 0.796 | |
| | | | | WT | 0.667 | 0.289 | | | | |
| | | D14 | Long | KO | 0.889 | 0.455 | | -0.713 | 0.546 | |
| | | | | WT | 1.111 | 0.351 | | | | |
| | | D15 | Long | KO | 1.222 | 0.547 | | -0.047 | 1.000 | |
| | | | | WT | 1.000 | 0.373 | | | | |
| | | D16 | Long | KO | 1.667 | 0.726 | | -1.211 | 0.340 | |
| | | | | WT | 0.444 | 0.338 | | | | |
| Table S5A | Figure 1H,I / Figure S7 | D0 | Short | KO | 0.389 | 0.232 | Mann-Whitney U Test | -1.835 | 0.258 | |
| | | | | WT | 0.000 | 0.000 | | | | |
| | | D1 | Short | KO | 0.167 | 0.118 | | -0.476 | 0.796 | |
| | | | | WT | 0.222 | 0.222 | | | | |
| | | D2 | Short | KO | 0.222 | 0.147 | | -0.122 | 0.931 | |
| | | | | WT | 0.167 | 0.118 | | | | |
| | | D3 | Short | KO | 0.333 | 0.167 | | -0.318 | 0.796 | |
| | | | | WT | 0.556 | 0.294 | | | | |
| | | D4 | Short | KO | 0.333 | 0.236 | | -1.568 | 0.190 | |
| | | | | WT | 1.444 | 0.648 | | | | |
| | | D5 | Short | KO | 0.889 | 0.200 | | -0.382 | 0.730 | |
| | | | | WT | 1.222 | 0.401 | | | | |
| | | D6 | Short | KO | 1.333 | 0.333 | | -3.181 | 0.004 | ** |
| | | | | WT | 0.000 | 0.000 | | | | |
| | | D7 | Short | KO | 0.889 | 0.351 | | -0.684 | 0.546 | |
| | | | | WT | 0.667 | 0.373 | | | | |
| | | D8 | Short | KO | 0.889 | 0.309 | | -0.927 | 0.387 | |
| | | | | WT | 1.333 | 0.333 | | | | |
| | | D9 | Short | KO | 1.000 | 0.553 | | -0.287 | 0.796 | |
| | | | | WT | 0.889 | 0.351 | | | | |
| | | D10 | Short | KO | 0.444 | 0.242 | | -1.350 | 0.222 | |
| | | | | WT | 1.000 | 0.333 | | | | |
| | | D11 | Short | KO | 1.222 | 0.434 | | -0.465 | 0.666 | |
| | | | | WT | 1.111 | 0.539 | | | | |
| | | D12 | Short | KO | 0.667 | 0.333 | | -0.199 | 0.863 | |
| | | | | WT | 0.556 | 0.176 | | | | |
| | | D13 | Short | KO | 0.444 | 0.176 | | -0.397 | 0.730 | |
| | | | | WT | 0.667 | 0.289 | | | | |
| | | D14 | Short | KO | 0.222 | 0.147 | | 0.000 | 1.000 | |
| | | | | WT | 0.222 | 0.147 | | | | |
| | | D15 | Short | KO | 0.222 | 0.147 | | -0.511 | 0.730 | |
| | | | | WT | 0.333 | 0.167 | | | | |
| | | D16 | Short | KO | 0.222 | 0.147 | | 0.000 | 1.000 | |
| | | | | WT | 0.222 | 0.147 | | | | |

HE frequency in 11–12 s window

| Table | Related Figure | Day | Group | Condition | Mean | SEM | Stastistical test | Z | P | * |
|---|---|---|---|---|---|---|---|---|---|---|
| Table S5 | Figure 1H,I / Figure S7 | D0 | KO | Long | 0.278 | 0.188 | Wilcoxon Signed Ranks Test | -0.271 | 0.786 | |
| | | | | Short | 0.389 | 0.232 | | | | |
| | | D1 | KO | Long | 0.111 | 0.073 | | -0.447 | 0.655 | |
| | | | | Short | 0.167 | 0.118 | | | | |
| | | D2 | KO | Long | 0.222 | 0.147 | | 0.000 | 1.000 | |
| | | | | Short | 0.222 | 0.147 | | | | |
| | | D3 | KO | Long | 0.222 | 0.147 | | -0.577 | 0.564 | |
| | | | | Short | 0.333 | 0.167 | | | | |
| | | D4 | KO | Long | 0.778 | 0.434 | | -1.633 | 0.102 | |
| | | | | Short | 0.333 | 0.236 | | | | |
| | | D5 | KO | Long | 1.556 | 0.475 | | -1.294 | 0.196 | |
| | | | | Short | 0.889 | 0.200 | | | | |
| | | D6 | KO | Long | 0.667 | 0.441 | | -1.387 | 0.165 | |
| | | | | Short | 1.333 | 0.333 | | | | |
| | | D7 | KO | Long | 1.111 | 0.351 | | -0.816 | 0.414 | |
| | | | | Short | 0.889 | 0.351 | | | | |
| | | D8 | KO | Long | 2.111 | 0.633 | | -1.420 | 0.156 | |
| | | | | Short | 0.889 | 0.309 | | | | |
| | | D9 | KO | Long | 2.333 | 0.833 | | -1.841 | 0.066 | |
| | | | | Long | 1.000 | 0.553 | | | | |
| | | D10 | KO | Short | 1.000 | 0.500 | | -0.736 | 0.461 | |
| | | | | Long | 0.444 | 0.242 | | | | |
| | | D11 | KO | Short | 0.778 | 0.364 | | -1.265 | 0.206 | |
| | | | | Long | 1.222 | 0.434 | | | | |
| | | D12 | KO | Short | 1.000 | 0.667 | | -0.318 | 0.750 | |
| | | | | Long | 0.667 | 0.333 | | | | |
| | | D13 | KO | Short | 0.556 | 0.294 | | -0.333 | 0.739 | |
| | | | | Long | 0.444 | 0.176 | | | | |
| | | D14 | KO | Short | 0.889 | 0.455 | | -1.604 | 0.109 | |
| | | | | Long | 0.222 | 0.147 | | | | |
| | | D15 | KO | Short | 1.222 | 0.547 | | -1.382 | 0.167 | |
| | | | | Long | 0.222 | 0.147 | | | | |
| | | D16 | KO | Short | 1.667 | 0.726 | | -1.753 | 0.080 | |
| | | | | Long | 0.222 | 0.147 | | | | |
| Table S5 | Figure 1H,I / Figure S7 | D0 | WT | Long | 0.444 | 0.444 | Wilcoxon Signed Ranks Test | -1.000 | 0.317 | |
| | | | | Short | 0.000 | 0.000 | | | | |
| | | D1 | WT | Long | 0.167 | 0.118 | | 0.000 | 1.000 | |
| | | | | Short | 0.222 | 0.222 | | | | |
| | | D2 | WT | Long | 0.222 | 0.121 | | -1.000 | 0.317 | |
| | | | | Short | 0.167 | 0.118 | | | | |
| | | D3 | WT | Long | 0.222 | 0.147 | | -1.089 | 0.276 | |
| | | | | Short | 0.556 | 0.294 | | | | |
| | | D4 | WT | Long | 0.667 | 0.289 | | -1.473 | 0.141 | |
| | | | | Short | 1.444 | 0.648 | | | | |
| | | D5 | WT | Long | 0.667 | 0.373 | | -0.940 | 0.347 | |
| | | | | Short | 1.222 | 0.401 | | | | |
| | | D6 | WT | Long | 0.444 | 0.242 | | -1.633 | 0.102 | |
| | | | | Short | 0.000 | 0.000 | | | | |
| | | D7 | WT | Long | 1.222 | 0.547 | | -0.962 | 0.336 | |
| | | | | Short | 0.667 | 0.373 | | | | |
| | | D8 | WT | Long | 1.111 | 0.423 | | -0.520 | 0.603 | |
| | | | | Short | 1.333 | 0.333 | | | | |
| | | D9 | WT | Long | 1.778 | 0.547 | | -1.511 | 0.131 | |
| | | | | Long | 0.889 | 0.351 | | | | |
| | | D10 | WT | Short | 1.333 | 0.471 | | -0.750 | 0.453 | |
| | | | | Long | 1.000 | 0.333 | | | | |
| | | D11 | WT | Short | 0.889 | 0.423 | | -0.707 | 0.480 | |
| | | | | Long | 1.111 | 0.539 | | | | |
| | | D12 | WT | Short | 0.667 | 0.289 | | -0.333 | 0.739 | |
| | | | | Long | 0.556 | 0.176 | | | | |
| | | D13 | WT | Short | 0.667 | 0.289 | | 0.000 | 1.000 | |
| | | | | Long | 0.667 | 0.289 | | | | |
| | | D14 | WT | Short | 1.111 | 0.351 | | -2.070 | 0.038 | * |
| | | | | Long | 0.222 | 0.147 | | | | |
| | | D15 | WT | Short | 1.000 | 0.373 | | -1.656 | 0.098 | |
| | | | | Long | 0.333 | 0.167 | | | | |
| | | D16 | WT | Short | 0.444 | 0.338 | | -0.378 | 0.705 | |
| | | | | Long | 0.222 | 0.147 | | | | |

5_13

| Table | Related Figure | Day | Condition | Group | Mean | SEM | Stastistical test | Z | P | * |
|---|---|---|---|---|---|---|---|---|---|---|
| HE frequency in 12–13 s window | | | | | | | | | | |
| Table S5 | Figure 1H,I / Figure S7 | D0 | Long | KO | 0.278 | 0.147 | Mann-Whitney U Test | -1.035 | 0.489 | |
| | | | | WT | 0.111 | 0.111 | | | | |
| | | D1 | Long | KO | 0.056 | 0.056 | | -0.615 | 0.730 | |
| | | | | WT | 0.111 | 0.073 | | | | |
| | | D2 | Long | KO | 0.333 | 0.236 | | -1.455 | 0.436 | |
| | | | | WT | 0.000 | 0.000 | | | | |
| | | D3 | Long | KO | 0.556 | 0.338 | | -1.253 | 0.258 | |
| | | | | WT | 1.000 | 0.333 | | | | |
| | | D4 | Long | KO | 0.889 | 0.423 | | -0.655 | 0.546 | |
| | | | | WT | 1.111 | 0.309 | | | | |
| | | D5 | Long | KO | 1.111 | 0.261 | | -0.185 | 0.863 | |
| | | | | WT | 1.222 | 0.364 | | | | |
| | | D6 | Long | KO | 1.556 | 0.580 | | -0.986 | 0.387 | |
| | | | | WT | 0.667 | 0.236 | | | | |
| | | D7 | Long | KO | 1.222 | 0.741 | | -1.055 | 0.387 | |
| | | | | WT | 0.333 | 0.167 | | | | |
| | | D8 | Long | KO | 1.000 | 0.289 | | -0.140 | 0.931 | |
| | | | | WT | 1.111 | 0.389 | | | | |
| | | D9 | Long | KO | 1.444 | 0.603 | | -0.418 | 0.730 | |
| | | | | WT | 1.000 | 0.373 | | | | |
| | | D10 | Long | KO | 0.444 | 0.444 | | -1.031 | 0.489 | |
| | | | | WT | 0.889 | 0.564 | | | | |
| | | D11 | Long | KO | 0.778 | 0.547 | | -1.122 | 0.340 | |
| | | | | WT | 2.000 | 0.986 | | | | |
| | | D12 | Long | KO | 1.556 | 0.689 | | -0.523 | 0.666 | |
| | | | | WT | 1.000 | 0.500 | | | | |
| | | D13 | Long | KO | 0.889 | 0.423 | | -0.098 | 0.931 | |
| | | | | WT | 0.778 | 0.364 | | | | |
| | | D14 | Long | KO | 0.333 | 0.236 | | -1.366 | 0.258 | |
| | | | | WT | 0.889 | 0.351 | | | | |
| | | D15 | Long | KO | 1.222 | 0.521 | | -0.535 | 0.666 | |
| | | | | WT | 0.667 | 0.333 | | | | |
| | | D16 | Long | KO | 0.667 | 0.333 | | -0.476 | 0.730 | |
| | | | | WT | 0.333 | 0.167 | | | | |
| Table S5A | Figure 1H,I / Figure S7 | D0 | Short | KO | 0.278 | 0.121 | Mann-Whitney U Test | -1.578 | 0.222 | |
| | | | | WT | 0.056 | 0.056 | | | | |
| | | D1 | Short | KO | 0.389 | 0.200 | | -1.156 | 0.436 | |
| | | | | WT | 0.111 | 0.111 | | | | |
| | | D2 | Short | KO | 0.722 | 0.278 | | -1.031 | 0.387 | |
| | | | | WT | 0.333 | 0.167 | | | | |
| | | D3 | Short | KO | 1.000 | 0.527 | | -1.748 | 0.161 | |
| | | | | WT | 0.222 | 0.222 | | | | |
| | | D4 | Short | KO | 0.889 | 0.261 | | -1.612 | 0.161 | |
| | | | | WT | 0.333 | 0.167 | | | | |
| | | D5 | Short | KO | 0.444 | 0.176 | | -0.051 | 1.000 | |
| | | | | WT | 0.556 | 0.294 | | | | |
| | | D6 | Short | KO | 1.000 | 0.333 | | -1.719 | 0.136 | |
| | | | | WT | 0.333 | 0.236 | | | | |
| | | D7 | Short | KO | 0.667 | 0.236 | | -0.191 | 0.863 | |
| | | | | WT | 1.222 | 0.596 | | | | |
| | | D8 | Short | KO | 1.000 | 0.441 | | -0.095 | 0.931 | |
| | | | | WT | 0.778 | 0.278 | | | | |
| | | D9 | Short | KO | 1.444 | 0.556 | | -0.660 | 0.546 | |
| | | | | WT | 0.778 | 0.222 | | | | |
| | | D10 | Short | KO | 0.556 | 0.338 | | -0.653 | 0.605 | |
| | | | | WT | 0.556 | 0.176 | | | | |
| | | D11 | Short | KO | 1.000 | 0.441 | | -0.146 | 0.931 | |
| | | | | WT | 1.000 | 0.553 | | | | |
| | | D12 | Short | KO | 0.333 | 0.167 | | -0.160 | 0.931 | |
| | | | | WT | 0.444 | 0.242 | | | | |
| | | D13 | Short | KO | 0.444 | 0.338 | | -0.680 | 0.666 | |
| | | | | WT | 0.111 | 0.111 | | | | |
| | | D14 | Short | KO | 0.556 | 0.242 | | -1.068 | 0.387 | |
| | | | | WT | 0.222 | 0.147 | | | | |
| | | D15 | Short | KO | 0.000 | 0.000 | | -2.191 | 0.113 | |
| | | | | WT | 0.667 | 0.333 | | | | |
| | | D16 | Short | KO | 0.111 | 0.111 | | -1.102 | 0.436 | |
| | | | | WT | 0.333 | 0.167 | | | | |

| Table | Related Figure | Day | Group | Condition | Mean | SEM | Stastistical test | Z | P | * |
|---|---|---|---|---|---|---|---|---|---|---|
| HE frequency in 12–13 s window | | | | | | | | | | |
| Table S5 | Figure 1H,I / Figure S7 | D0 | KO | Long | 0.278 | 0.147 | Wilcoxon Signed Ranks Test | 0.000 | 1.000 | |
| | | | | Short | 0.278 | 0.121 | | | | |
| | | D1 | KO | Long | 0.056 | 0.056 | | -1.604 | 0.109 | |
| | | | | Short | 0.389 | 0.200 | | | | |
| | | D2 | KO | Long | 0.333 | 0.236 | | -1.236 | 0.216 | |
| | | | | Short | 0.722 | 0.278 | | | | |
| | | D3 | KO | Long | 0.556 | 0.338 | | -0.736 | 0.461 | |
| | | | | Short | 1.000 | 0.527 | | | | |
| | | D4 | KO | Long | 0.889 | 0.423 | | 0.000 | 1.000 | |
| | | | | Short | 0.889 | 0.261 | | | | |
| | | D5 | KO | Long | 1.111 | 0.261 | | -1.897 | 0.058 | |
| | | | | Short | 0.444 | 0.176 | | | | |
| | | D6 | KO | Long | 1.556 | 0.580 | | -1.406 | 0.160 | |
| | | | | Short | 1.000 | 0.333 | | | | |
| | | D7 | KO | Long | 1.222 | 0.741 | | -0.535 | 0.593 | |
| | | | | Short | 0.667 | 0.236 | | | | |
| | | D8 | KO | Long | 1.000 | 0.289 | | 0.000 | 1.000 | |
| | | | | Short | 1.000 | 0.441 | | | | |
| | | D9 | KO | Long | 1.444 | 0.603 | | 0.000 | 1.000 | |
| | | | | Long | 1.444 | 0.556 | | | | |
| | | D10 | KO | Short | 0.444 | 0.444 | | -0.577 | 0.564 | |
| | | | | Long | 0.556 | 0.338 | | | | |
| | | D11 | KO | Short | 0.778 | 0.547 | | -0.368 | 0.713 | |
| | | | | Long | 1.000 | 0.441 | | | | |
| | | D12 | KO | Short | 1.556 | 0.689 | | -1.625 | 0.104 | |
| | | | | Long | 0.333 | 0.167 | | | | |
| | | D13 | KO | Short | 0.889 | 0.423 | | -0.816 | 0.414 | |
| | | | | Long | 0.444 | 0.338 | | | | |
| | | D14 | KO | Short | 0.333 | 0.236 | | -1.000 | 0.317 | |
| | | | | Long | 0.556 | 0.242 | | | | |
| | | D15 | KO | Short | 1.222 | 0.521 | | -1.841 | 0.066 | |
| | | | | Long | 0.000 | 0.000 | | | | |
| | | D16 | KO | Short | 0.667 | 0.333 | | -1.512 | 0.131 | |
| | | | | Long | 0.111 | 0.111 | | | | |
| Table S5 | Figure 1H,I / Figure S7 | D0 | WT | Long | 0.111 | 0.111 | Wilcoxon Signed Ranks Test | -0.447 | 0.655 | |
| | | | | Short | 0.056 | 0.056 | | | | |
| | | D1 | WT | Long | 0.111 | 0.073 | | 0.000 | 1.000 | |
| | | | | Short | 0.111 | 0.111 | | | | |
| | | D2 | WT | Long | 0.000 | 0.000 | | -1.732 | 0.083 | |
| | | | | Short | 0.333 | 0.167 | | | | |
| | | D3 | WT | Long | 1.000 | 0.333 | | -1.725 | 0.084 | |
| | | | | Short | 0.222 | 0.222 | | | | |
| | | D4 | WT | Long | 1.111 | 0.309 | | -1.823 | 0.068 | |
| | | | | Short | 0.333 | 0.167 | | | | |
| | | D5 | WT | Long | 1.222 | 0.364 | | -1.730 | 0.084 | |
| | | | | Short | 0.556 | 0.294 | | | | |
| | | D6 | WT | Long | 0.667 | 0.236 | | -1.134 | 0.257 | |
| | | | | Short | 0.333 | 0.236 | | | | |
| | | D7 | WT | Long | 0.333 | 0.167 | | -1.633 | 0.102 | |
| | | | | Short | 1.222 | 0.596 | | | | |
| | | D8 | WT | Long | 1.111 | 0.389 | | -1.000 | 0.317 | |
| | | | | Short | 0.778 | 0.278 | | | | |
| | | D9 | WT | Long | 1.000 | 0.373 | | -0.632 | 0.527 | |
| | | | | Long | 0.778 | 0.222 | | | | |
| | | D10 | WT | Short | 0.889 | 0.564 | | -0.378 | 0.705 | |
| | | | | Long | 0.556 | 0.176 | | | | |
| | | D11 | WT | Short | 2.000 | 0.986 | | -1.461 | 0.144 | |
| | | | | Long | 1.000 | 0.553 | | | | |
| | | D12 | WT | Short | 1.000 | 0.500 | | -1.518 | 0.129 | |
| | | | | Long | 0.444 | 0.242 | | | | |
| | | D13 | WT | Short | 0.778 | 0.364 | | -1.857 | 0.063 | |
| | | | | Long | 0.111 | 0.111 | | | | |
| | | D14 | WT | Short | 0.889 | 0.351 | | -1.511 | 0.131 | |
| | | | | Long | 0.222 | 0.147 | | | | |
| | | D15 | WT | Short | 0.667 | 0.333 | | -0.137 | 0.891 | |
| | | | | Long | 0.667 | 0.333 | | | | |
| | | D16 | WT | Short | 0.333 | 0.167 | | 0.000 | 1.000 | |
| | | | | Long | 0.333 | 0.167 | | | | |

5_14

| Table | Related Figure | Day | Condition | Group | Mean | SEM | Stastistical test | Z | P | * |
|---|---|---|---|---|---|---|---|---|---|---|
| HE frequency in 13–14 s window | | | | | | | | | | |
| Table S5 | Figure 1H,I / Figure S7 | D0 | Long | KO | 0.000 | 0.000 | Mann-Whitney U Test | -1.000 | 0.730 | |
| | | | | WT | 0.056 | 0.056 | | | | |
| | | D1 | Long | KO | 0.167 | 0.118 | | -0.544 | 0.730 | |
| | | | | WT | 0.111 | 0.111 | | | | |
| | | D2 | Long | KO | 0.389 | 0.232 | | -0.503 | 0.666 | |
| | | | | WT | 0.556 | 0.256 | | | | |
| | | D3 | Long | KO | 0.333 | 0.236 | | 0.000 | 1.000 | |
| | | | | WT | 0.333 | 0.236 | | | | |
| | | D4 | Long | KO | 1.000 | 0.333 | | -0.095 | 0.931 | |
| | | | | WT | 0.889 | 0.261 | | | | |
| | | D5 | Long | KO | 0.444 | 0.242 | | -1.219 | 0.297 | |
| | | | | WT | 1.667 | 0.726 | | | | |
| | | D6 | Long | KO | 1.111 | 0.351 | | -1.046 | 0.340 | |
| | | | | WT | 0.778 | 0.465 | | | | |
| | | D7 | Long | KO | 0.556 | 0.242 | | -1.068 | 0.387 | |
| | | | | WT | 0.222 | 0.147 | | | | |
| | | D8 | Long | KO | 1.222 | 0.662 | | -0.190 | 0.863 | |
| | | | | WT | 1.000 | 0.408 | | | | |
| | | D9 | Long | KO | 0.889 | 0.512 | | -0.158 | 0.931 | |
| | | | | WT | 1.111 | 0.655 | | | | |
| | | D10 | Long | KO | 1.222 | 0.662 | | -0.252 | 0.863 | |
| | | | | WT | 0.889 | 0.455 | | | | |
| | | D11 | Long | KO | 1.000 | 0.373 | | -0.512 | 0.666 | |
| | | | | WT | 1.444 | 0.530 | | | | |
| | | D12 | Long | KO | 0.556 | 0.556 | | -0.910 | 0.546 | |
| | | | | WT | 0.556 | 0.338 | | | | |
| | | D13 | Long | KO | 0.667 | 0.441 | | -0.527 | 0.666 | |
| | | | | WT | 0.667 | 0.289 | | | | |
| | | D14 | Long | KO | 0.444 | 0.338 | | -0.680 | 0.666 | |
| | | | | WT | 0.111 | 0.111 | | | | |
| | | D15 | Long | KO | 0.556 | 0.338 | | -0.620 | 0.666 | |
| | | | | WT | 0.222 | 0.147 | | | | |
| | | D16 | Long | KO | 0.778 | 0.324 | | -0.591 | 0.605 | |
| | | | | WT | 0.444 | 0.176 | | | | |
| Table S5A | Figure 1H,I / Figure S7 | D0 | Short | KO | 0.333 | 0.118 | Mann-Whitney U Test | -1.222 | 0.297 | |
| | | | | WT | 0.167 | 0.118 | | | | |
| | | D1 | Short | KO | 0.389 | 0.232 | | -0.252 | 0.863 | |
| | | | | WT | 0.333 | 0.144 | | | | |
| | | D2 | Short | KO | 0.389 | 0.162 | | -0.710 | 0.546 | |
| | | | | WT | 0.222 | 0.121 | | | | |
| | | D3 | Short | KO | 1.111 | 0.484 | | -0.094 | 0.931 | |
| | | | | WT | 0.889 | 0.261 | | | | |
| | | D4 | Short | KO | 0.667 | 0.289 | | -0.145 | 0.931 | |
| | | | | WT | 0.667 | 0.236 | | | | |
| | | D5 | Short | KO | 0.667 | 0.236 | | -0.520 | 0.666 | |
| | | | | WT | 1.111 | 0.455 | | | | |
| | | D6 | Short | KO | 1.444 | 0.444 | | -1.051 | 0.340 | |
| | | | | WT | 0.778 | 0.222 | | | | |
| | | D7 | Short | KO | 1.333 | 0.624 | | -0.486 | 0.666 | |
| | | | | WT | 0.556 | 0.176 | | | | |
| | | D8 | Short | KO | 0.333 | 0.236 | | -1.318 | 0.258 | |
| | | | | WT | 0.778 | 0.278 | | | | |
| | | D9 | Short | KO | 0.889 | 0.309 | | -0.050 | 1.000 | |
| | | | | WT | 0.778 | 0.222 | | | | |
| | | D10 | Short | KO | 0.889 | 0.261 | | -0.622 | 0.605 | |
| | | | | WT | 0.667 | 0.236 | | | | |
| | | D11 | Short | KO | 1.111 | 0.351 | | -0.561 | 0.605 | |
| | | | | WT | 0.889 | 0.389 | | | | |
| | | D12 | Short | KO | 0.333 | 0.167 | | -2.153 | 0.050 | |
| | | | | WT | 1.111 | 0.261 | | | | |
| | | D13 | Short | KO | 0.778 | 0.547 | | -0.505 | 0.730 | |
| | | | | WT | 0.333 | 0.236 | | | | |
| | | D14 | Short | KO | 0.111 | 0.111 | | -2.366 | 0.040 | * |
| | | | | WT | 0.778 | 0.222 | | | | |
| | | D15 | Short | KO | 0.333 | 0.236 | | 0.000 | 1.000 | |
| | | | | WT | 0.333 | 0.236 | | | | |
| | | D16 | Short | KO | 0.333 | 0.236 | | -0.449 | 0.730 | |
| | | | | WT | 0.444 | 0.242 | | | | |

| Table | Related Figure | Day | Group | Condition | Mean | SEM | Stastistical test | Z | P | * |
|---|---|---|---|---|---|---|---|---|---|---|
| HE frequency in 13–14 s window | | | | | | | | | | |
| Table S5 | Figure 1H,I / Figure S7 | D0 | KO | Long | 0.000 | 0.000 | Wilcoxon Signed Ranks Test | -2.121 | 0.034 | * |
| | | | | Short | 0.333 | 0.118 | | | | |
| | | D1 | KO | Long | 0.167 | 0.118 | | -0.680 | 0.496 | |
| | | | | Short | 0.389 | 0.232 | | | | |
| | | D2 | KO | Long | 0.389 | 0.232 | | 0.000 | 1.000 | |
| | | | | Short | 0.389 | 0.162 | | | | |
| | | D3 | KO | Long | 0.333 | 0.236 | | -1.361 | 0.174 | |
| | | | | Short | 1.111 | 0.484 | | | | |
| | | D4 | KO | Long | 1.000 | 0.333 | | -0.647 | 0.518 | |
| | | | | Short | 0.667 | 0.289 | | | | |
| | | D5 | KO | Long | 0.444 | 0.242 | | -0.707 | 0.480 | |
| | | | | Short | 0.667 | 0.236 | | | | |
| | | D6 | KO | Long | 1.111 | 0.351 | | -0.647 | 0.518 | |
| | | | | Short | 1.444 | 0.444 | | | | |
| | | D7 | KO | Long | 0.556 | 0.242 | | -0.689 | 0.491 | |
| | | | | Short | 1.333 | 0.624 | | | | |
| | | D8 | KO | Long | 1.222 | 0.662 | | -1.225 | 0.221 | |
| | | | | Short | 0.333 | 0.236 | | | | |
| | | D9 | KO | Long | 0.889 | 0.512 | | -0.106 | 0.916 | |
| | | | | Long | 0.889 | 0.309 | | | | |
| | | D10 | KO | Short | 1.222 | 0.662 | | -0.333 | 0.739 | |
| | | | | Long | 0.889 | 0.261 | | | | |
| | | D11 | KO | Short | 1.000 | 0.373 | | -0.276 | 0.783 | |
| | | | | Long | 1.111 | 0.351 | | | | |
| | | D12 | KO | Short | 0.556 | 0.556 | | -0.378 | 0.705 | |
| | | | | Long | 0.333 | 0.167 | | | | |
| | | D13 | KO | Short | 0.667 | 0.441 | | 0.000 | 1.000 | |
| | | | | Long | 0.778 | 0.547 | | | | |
| | | D14 | KO | Short | 0.444 | 0.338 | | -0.816 | 0.414 | |
| | | | | Long | 0.111 | 0.111 | | | | |
| | | D15 | KO | Short | 0.556 | 0.338 | | -0.412 | 0.680 | |
| | | | | Long | 0.333 | 0.236 | | | | |
| | | D16 | KO | Short | 0.778 | 0.324 | | -0.966 | 0.334 | |
| | | | | Long | 0.333 | 0.236 | | | | |
| Table S5 | Figure 1H,I / Figure S7 | D0 | WT | Long | 0.056 | 0.056 | Wilcoxon Signed Ranks Test | -0.816 | 0.414 | |
| | | | | Short | 0.167 | 0.118 | | | | |
| | | D1 | WT | Long | 0.111 | 0.111 | | -1.633 | 0.102 | |
| | | | | Short | 0.333 | 0.144 | | | | |
| | | D2 | WT | Long | 0.556 | 0.256 | | -1.289 | 0.197 | |
| | | | | Short | 0.222 | 0.121 | | | | |
| | | D3 | WT | Long | 0.333 | 0.236 | | -1.406 | 0.160 | |
| | | | | Short | 0.889 | 0.261 | | | | |
| | | D4 | WT | Long | 0.889 | 0.261 | | -0.707 | 0.480 | |
| | | | | Short | 0.667 | 0.236 | | | | |
| | | D5 | WT | Long | 1.667 | 0.726 | | -0.284 | 0.777 | |
| | | | | Short | 1.111 | 0.455 | | | | |
| | | D6 | WT | Long | 0.778 | 0.465 | | -0.431 | 0.666 | |
| | | | | Short | 0.778 | 0.222 | | | | |
| | | D7 | WT | Long | 0.222 | 0.147 | | -1.342 | 0.180 | |
| | | | | Short | 0.556 | 0.176 | | | | |
| | | D8 | WT | Long | 1.000 | 0.408 | | -0.425 | 0.671 | |
| | | | | Short | 0.778 | 0.278 | | | | |
| | | D9 | WT | Long | 1.111 | 0.655 | | -0.258 | 0.796 | |
| | | | | Long | 0.778 | 0.222 | | | | |
| | | D10 | WT | Short | 0.889 | 0.455 | | -0.412 | 0.680 | |
| | | | | Long | 0.667 | 0.236 | | | | |
| | | D11 | WT | Short | 1.444 | 0.530 | | -2.236 | 0.025 | |
| | | | | Long | 0.889 | 0.389 | | | | |
| | | D12 | WT | Short | 0.556 | 0.338 | | -1.207 | 0.227 | |
| | | | | Long | 1.111 | 0.261 | | | | |
| | | D13 | WT | Short | 0.667 | 0.289 | | -0.966 | 0.334 | |
| | | | | Long | 0.333 | 0.236 | | | | |
| | | D14 | WT | Short | 0.111 | 0.111 | | -1.897 | 0.058 | |
| | | | | Long | 0.778 | 0.222 | | | | |
| | | D15 | WT | Short | 0.222 | 0.147 | | -0.378 | 0.705 | |
| | | | | Long | 0.333 | 0.236 | | | | |
| | | D16 | WT | Short | 0.444 | 0.176 | | 0.000 | 1.000 | |
| | | | | Long | 0.444 | 0.242 | | | | |

## 6_1

| CR | | | | | | | | |
|---|---|---|---|---|---|---|---|---|
| Table | Related Figure | Condition | Mean | SEM | Stastistical test | Z | P | * |
| Table S6 | Figure 2B | Long | 3316.374 | 116.818 | Wilcoxon Signed Ranks Test | -4.107 | < 0.0001 | *** |
| | | Short | 2575.683 | 17.594 | | | | |

## 6_2

| RT | | | | | | | | |
|---|---|---|---|---|---|---|---|---|
| Table | Related Figure | Condition | Mean | SEM | Stastistical test | Z | P | * |
| Table S6 | Figure 2B | Long | 99.542 | 0.146 | Wilcoxon Signed Ranks Test | -5.511 | < 0.00000001 | *** |
| | | Short | 97.333 | 0.536 | | | | |

## 6_3

| Correlation between Long and Short in RT | | | | | | | | |
|---|---|---|---|---|---|---|---|---|
| Table | Related Figure | Condition | Mean | SEM | Stastistical test | Spearman's ρ | P | * |
| Table S6 | Figure 2B | Long | 99.542 | 0.146 | Spearman's rank correlation coefficient | 0.601 | < 0.0001 | *** |
| | | Short | 97.333 | 0.536 | | | | |

Abbreviations: CR, correct rate; RT, reaction time.
*, *p < 0.05, **p < 0.01, ***p < 0.001.

## 7_1

| Human: Correlation between LATENCY (L-S difference in RT) and ACCURACY (CR) | | | | | | | | |
|---|---|---|---|---|---|---|---|---|
| Table | Related Figure | Condition | Mean | SEM | Stastistical test | Spearman's ρ | Bonferroni corrected P | * |
| Table S7 | Figure 3C | RT difference | 740.69 | 110.733 | Spearman's rank correlation coefficient | 0.094 | 1.000 | |
| | | CR for Long | 99.542 | 0.146 | | | | |
| | | RT difference | 740.69 | 110.733 | | 0.265 | 0.198 | |
| | | CR for Short | 97.333 | 0.536 | | | | |

| Human: Correlation between LATENCY ( RT) and ACCURACY (CR) | | | | | | | | |
|---|---|---|---|---|---|---|---|---|
| Table | Related Figure | Condition | Mean | SEM | Stastistical test | Spearman's ρ | Bonferroni corrected P | * |
| Table S7 | Figure S8A | RT for Long | 3316.37 | 116.82 | Spearman's rank correlation coefficient | 0.032 | 1.0000 | |
| | | CR for Long | 99.542 | 0.146 | | | | |
| | | RT for Short | 2575.68 | 17.59 | | 0.275 | 1.0000 | |
| | | CR for Short | 97.333 | 0.536 | | | | |

Abbreviations: CR, correct rate; RT, reaction time; HE, head entry; D, training day; KO, CRBN KO mice; WT, wild-type mice.

*, *p < 0.05, **p < 0.01, ***p < 0.001.

## 7_2

| Mouse: Correlation between LATENCY (L-S difference in HE peak latency) and ACCURACY (hit and correct rejection rates) in KO mice | | | | | | | | | | |
|---|---|---|---|---|---|---|---|---|---|---|
| Table | Related Figure | Day | Condition | Pair | Mean | SEM | Stastistical test | Spearman's ρ | P | * |
| Table S7 | Figure 3C / Figure S8C | D0 | Long | LATENCY | 0.333 | 2.021 | Spearman's rank correlation coefficient | -0.157 | 1.000 | |
| | | | | ACCURACY | 27.780 | 8.780 | | | | |
| | | D1 | Long | LATENCY | -2.333 | 1.213 | | -0.095 | 1.000 | |
| | | | | ACCURACY | 18.890 | 7.720 | | | | |
| | | D2 | Long | LATENCY | -2.000 | 2.055 | | 0.613 | 0.159 | |
| | | | | ACCURACY | 32.220 | 9.830 | | | | |
| | | D3 | Long | LATENCY | -2.000 | 1.944 | | 0.047 | 1.000 | |
| | | | | ACCURACY | 38.890 | 10.060 | | | | |
| | | D4 | Long | LATENCY | -0.556 | 2.199 | | 0.551 | 0.248 | |
| | | | | ACCURACY | 56.670 | 12.020 | | | | |
| | | D5 | Long | LATENCY | 2.000 | 1.756 | | 0.494 | 0.354 | |
| | | | | ACCURACY | 50.000 | 9.130 | | | | |
| | | D6 | Long | LATENCY | 1.444 | 1.556 | | 0.302 | 0.859 | |
| | | | | ACCURACY | 75.560 | 7.660 | | | | |
| | | D7 | Long | LATENCY | 0.778 | 1.942 | | -0.017 | 1.000 | |
| | | | | ACCURACY | 68.890 | 11.480 | | | | |
| | | D8 | Long | LATENCY | -0.111 | 1.867 | | 0.433 | 0.489 | |
| | | | | ACCURACY | 87.780 | 7.950 | | | | |
| | | D9 | Long | LATENCY | 2.444 | 0.852 | | -0.337 | 0.751 | |
| | | | | ACCURACY | 87.780 | 8.620 | | | | |
| | | D10 | Long | LATENCY | 2.889 | 0.964 | | -0.345 | 0.726 | |
| | | | | ACCURACY | 80.000 | 10.410 | | | | |
| | | D11 | Long | LATENCY | 1.556 | 1.355 | | -0.309 | 0.837 | |
| | | | | ACCURACY | 70.000 | 11.550 | | | | |
| | | D12 | Long | LATENCY | 1.000 | 1.014 | | 0.334 | 0.759 | |
| | | | | ACCURACY | 83.330 | 9.130 | | | | |
| | | D13 | Long | LATENCY | 2.556 | 0.899 | | -0.397 | 0.581 | |
| | | | | ACCURACY | 77.780 | 7.030 | | | | |
| | | D14 | Long | LATENCY | 2.222 | 0.778 | | -0.196 | 1.000 | |
| | | | | ACCURACY | 83.330 | 7.640 | | | | |
| | | D15 | Long | LATENCY | 3.889 | 1.328 | | -0.672 | 0.095 | |
| | | | | ACCURACY | 82.220 | 9.250 | | | | |
| | | D16 | Long | LATENCY | 3.556 | 1.345 | | -0.808 | 0.017 | * |
| | | | | ACCURACY | 87.780 | 4.940 | | | | |
| Table S7 | Figure 3C / Figure S8C | D0 | Short | LATENCY | 0.333 | 2.021 | Spearman's rank correlation coefficient | -0.568 | 0.222 | |
| | | | | ACCURACY | 95.560 | 2.420 | | | | |
| | | D1 | Short | LATENCY | -2.333 | 1.213 | | 0.055 | 1.775 | |
| | | | | ACCURACY | 93.330 | 2.360 | | | | |
| | | D2 | Short | LATENCY | -2.000 | 2.055 | | -0.383 | 0.618 | |
| | | | | ACCURACY | 88.890 | 2.610 | | | | |
| | | D3 | Short | LATENCY | -2.000 | 1.944 | | -0.183 | 1.276 | |
| | | | | ACCURACY | 88.890 | 3.510 | | | | |
| | | D4 | Short | LATENCY | -0.556 | 2.199 | | -0.722 | 0.056 | |
| | | | | ACCURACY | 78.890 | 6.110 | | | | |
| | | D5 | Short | LATENCY | 2.000 | 1.756 | | -0.735 | 0.048 | * |
| | | | | ACCURACY | 84.440 | 3.380 | | | | |
| | | D6 | Short | LATENCY | 1.444 | 1.556 | | -0.690 | 0.080 | |
| | | | | ACCURACY | 81.110 | 4.230 | | | | |
| | | D7 | Short | LATENCY | 0.778 | 1.942 | | -0.337 | 0.752 | |
| | | | | ACCURACY | 70.000 | 7.070 | | | | |
| | | D8 | Short | LATENCY | -0.111 | 1.867 | | 0.139 | 1.442 | |
| | | | | ACCURACY | 77.780 | 5.470 | | | | |
| | | D9 | Short | LATENCY | 2.444 | 0.852 | | 0.056 | 1.773 | |
| | | | | ACCURACY | 71.110 | 4.230 | | | | |
| | | D10 | Short | LATENCY | 2.889 | 0.964 | | 0.407 | 0.553 | |
| | | | | ACCURACY | 73.330 | 5.000 | | | | |
| | | D11 | Short | LATENCY | 1.556 | 1.355 | | 0.470 | 0.403 | |
| | | | | ACCURACY | 76.670 | 7.640 | | | | |
| | | D12 | Short | LATENCY | 1.000 | 1.014 | | 0.000 | 2.000 | |
| | | | | ACCURACY | 71.110 | 6.550 | | | | |
| | | D13 | Short | LATENCY | 2.556 | 0.899 | | 0.291 | 0.894 | |
| | | | | ACCURACY | 73.330 | 8.160 | | | | |
| | | D14 | Short | LATENCY | 2.222 | 0.778 | | 0.419 | 0.523 | |
| | | | | ACCURACY | 63.330 | 7.990 | | | | |
| | | D15 | Short | LATENCY | 3.889 | 1.328 | | 0.798 | 0.020 | * |
| | | | | ACCURACY | 67.780 | 8.460 | | | | |
| | | D16 | Short | LATENCY | 3.556 | 1.345 | | 0.788 | 0.023 | * |
| | | | | ACCURACY | 70.000 | 7.260 | | | | |

| Mouse: Correlation between LATENCY (L-S difference in HE peak latency) and ACCURACY (hit and correct rejection rates) in WT mice | | | | | | | | | | |
|---|---|---|---|---|---|---|---|---|---|---|
| Table | Related Figure | Day | Condition | Pair | Mean | SEM | Stastistical test | Spearman's ρ | P | * |
| Table S7 | Figure 3C / Figure S8C | D0 | Long | LATENCY | -2.222 | 2.093 | Spearman's rank correlation coefficient | 0.409 | 0.550 | |
| | | | | ACCURACY | 6.670 | 2.360 | | | | |
| | | D1 | Long | LATENCY | -2.333 | 1.302 | | -0.431 | 0.493 | |
| | | | | ACCURACY | 13.330 | 5.270 | | | | |
| | | D2 | Long | LATENCY | 1.222 | 1.801 | | -0.389 | 0.603 | |
| | | | | ACCURACY | 24.440 | 8.520 | | | | |
| | | D3 | Long | LATENCY | 1.556 | 2.001 | | 0.621 | 0.149 | |
| | | | | ACCURACY | 24.440 | 7.090 | | | | |
| | | D4 | Long | LATENCY | -0.222 | 1.928 | | 0.321 | 0.800 | |
| | | | | ACCURACY | 56.670 | 10.270 | | | | |
| | | D5 | Long | LATENCY | 2.778 | 1.778 | | 0.178 | 1.000 | |
| | | | | ACCURACY | 54.440 | 11.320 | | | | |
| | | D6 | Long | LATENCY | 3.333 | 1.472 | | 0.127 | 1.000 | |
| | | | | ACCURACY | 62.220 | 9.090 | | | | |
| | | D7 | Long | LATENCY | 0.111 | 1.837 | | 0.025 | 1.000 | |
| | | | | ACCURACY | 71.110 | 6.330 | | | | |
| | | D8 | Long | LATENCY | -1.222 | 1.441 | | -0.070 | 1.000 | |
| | | | | ACCURACY | 81.110 | 4.840 | | | | |
| | | D9 | Long | LATENCY | 0.111 | 0.949 | | -0.083 | 1.000 | |
| | | | | ACCURACY | 78.890 | 6.330 | | | | |
| | | D10 | Long | LATENCY | -0.889 | 1.467 | | 0.191 | 1.000 | |
| | | | | ACCURACY | 83.330 | 5.770 | | | | |
| | | D11 | Long | LATENCY | 1.667 | 1.354 | | -0.569 | 0.220 | |
| | | | | ACCURACY | 84.440 | 5.560 | | | | |
| | | D12 | Long | LATENCY | 0.556 | 1.415 | | -0.411 | 0.545 | |
| | | | | ACCURACY | 86.670 | 4.710 | | | | |
| | | D13 | Long | LATENCY | 0.667 | 0.687 | | 0.411 | 0.545 | |
| | | | | ACCURACY | 86.670 | 4.710 | | | | |
| | | D14 | Long | LATENCY | 0.556 | 1.314 | | -0.416 | 0.532 | |
| | | | | ACCURACY | 84.440 | 3.770 | | | | |
| | | D15 | Long | LATENCY | 1.000 | 0.577 | | -0.101 | 1.000 | |
| | | | | ACCURACY | 87.780 | 4.010 | | | | |
| | | D16 | Long | LATENCY | 3.778 | 1.051 | | -0.009 | 1.000 | |
| | | | | ACCURACY | 92.220 | 3.240 | | | | |
| Table S7 | Figure 3C / Figure S8C | D0 | Short | LATENCY | -2.222 | 2.093 | Spearman's rank correlation coefficient | 0.000 | 1.000 | |
| | | | | ACCURACY | 98.890 | 1.110 | | | | |
| | | D1 | Short | LATENCY | -2.333 | 1.302 | | 0.000 | 0.000 | |
| | | | | ACCURACY | 100.000 | 0.000 | | | | |
| | | D2 | Short | LATENCY | 1.222 | 1.801 | | -0.487 | 0.368 | |
| | | | | ACCURACY | 94.440 | 1.760 | | | | |
| | | D3 | Short | LATENCY | 1.556 | 2.001 | | -0.377 | 0.634 | |
| | | | | ACCURACY | 88.890 | 2.610 | | | | |
| | | D4 | Short | LATENCY | -0.222 | 1.928 | | -0.272 | 0.957 | |
| | | | | ACCURACY | 77.780 | 4.340 | | | | |
| | | D5 | Short | LATENCY | 2.778 | 1.778 | | 0.064 | 1.000 | |
| | | | | ACCURACY | 83.330 | 2.360 | | | | |
| | | D6 | Short | LATENCY | 3.333 | 1.472 | | 0.134 | 1.000 | |
| | | | | ACCURACY | 90.000 | 4.080 | | | | |
| | | D7 | Short | LATENCY | 0.111 | 1.837 | | 0.550 | 0.250 | |
| | | | | ACCURACY | 86.670 | 1.670 | | | | |
| | | D8 | Short | LATENCY | -1.222 | 1.441 | | 0.438 | 0.477 | |
| | | | | ACCURACY | 78.890 | 4.550 | | | | |
| | | D9 | Short | LATENCY | 0.111 | 0.949 | | -0.305 | 0.849 | |
| | | | | ACCURACY | 86.670 | 2.890 | | | | |
| | | D10 | Short | LATENCY | -0.889 | 1.467 | | -0.106 | 1.000 | |
| | | | | ACCURACY | 82.220 | 4.940 | | | | |
| | | D11 | Short | LATENCY | 1.667 | 1.354 | | -0.507 | 0.328 | |
| | | | | ACCURACY | 77.780 | 3.640 | | | | |
| | | D12 | Short | LATENCY | 0.556 | 1.415 | | 0.026 | 1.000 | |
| | | | | ACCURACY | 78.890 | 4.840 | | | | |
| | | D13 | Short | LATENCY | 0.667 | 0.687 | | -0.301 | 0.863 | |
| | | | | ACCURACY | 85.560 | 2.940 | | | | |
| | | D14 | Short | LATENCY | 0.556 | 1.314 | | 0.207 | 1.000 | |
| | | | | ACCURACY | 81.110 | 4.550 | | | | |
| | | D15 | Short | LATENCY | 1.000 | 0.577 | | 0.017 | 1.000 | |
| | | | | ACCURACY | 81.110 | 4.840 | | | | |
| | | D16 | Short | LATENCY | 3.778 | 1.051 | | 0.681 | 0.087 | |
| | | | | ACCURACY | 77.780 | 3.640 | | | | |

## 7_3

| Mouse: Correlation between LATENCY (HE peak latency) and ACCURACY (hit and correct rejection rates) in KO mice | | | | | | | | | | |
|---|---|---|---|---|---|---|---|---|---|---|
| Table | Related Figure | Day | Condition | Pair | Mean | SEM | Stastistical test | Spearman's ρ | P | * |
| Table S7 | Figure S8B | D0 | Long | LATENCY | 4.000 | 1.093 | Spearman's rank correlation coefficient | -0.035 | 1.000 | |
| | | | | ACCURACY | 27.780 | 8.780 | | | | |
| | | D1 | Long | LATENCY | 1.000 | 0.408 | | 0.401 | 0.568 | |
| | | | | ACCURACY | 18.890 | 7.720 | | | | |
| | | D2 | Long | LATENCY | 4.222 | 1.310 | | 0.382 | 0.621 | |
| | | | | ACCURACY | 32.220 | 9.830 | | | | |
| | | D3 | Long | LATENCY | 3.667 | 1.291 | | 0.096 | 1.000 | |
| | | | | ACCURACY | 38.890 | 10.060 | | | | |
| | | D4 | Long | LATENCY | 2.889 | 1.296 | | 0.715 | 0.061 | |
| | | | | ACCURACY | 56.670 | 12.020 | | | | |
| | | D5 | Long | LATENCY | 4.667 | 1.202 | | 0.430 | 0.497 | |
| | | | | ACCURACY | 50.000 | 9.130 | | | | |
| | | D6 | Long | LATENCY | 4.222 | 1.164 | | 0.342 | 0.736 | |
| | | | | ACCURACY | 75.560 | 7.660 | | | | |
| | | D7 | Long | LATENCY | 3.556 | 1.215 | | 0.274 | 0.952 | |
| | | | | ACCURACY | 68.890 | 11.480 | | | | |
| | | D8 | Long | LATENCY | 3.222 | 1.382 | | 0.228 | 1.000 | |
| | | | | ACCURACY | 87.780 | 7.950 | | | | |
| | | D9 | Long | LATENCY | 5.000 | 1.333 | | -0.666 | 0.100 | |
| | | | | ACCURACY | 87.780 | 8.620 | | | | |
| | | D10 | Long | LATENCY | 5.111 | 1.172 | | -0.646 | 0.121 | |
| | | | | ACCURACY | 80.000 | 10.410 | | | | |
| | | D11 | Long | LATENCY | 5.556 | 1.600 | | -0.369 | 0.657 | |
| | | | | ACCURACY | 70.000 | 11.550 | | | | |
| | | D12 | Long | LATENCY | 3.111 | 1.252 | | -0.076 | 1.000 | |
| | | | | ACCURACY | 83.330 | 9.130 | | | | |
| | | D13 | Long | LATENCY | 5.111 | 1.348 | | -0.797 | 0.020 | * |
| | | | | ACCURACY | 77.780 | 7.030 | | | | |
| | | D14 | Long | LATENCY | 3.889 | 0.889 | | -0.101 | 1.000 | |
| | | | | ACCURACY | 83.330 | 7.640 | | | | |
| | | D15 | Long | LATENCY | 5.222 | 1.579 | | -0.787 | 0.024 | * |
| | | | | ACCURACY | 82.220 | 9.250 | | | | |
| | | D16 | Long | LATENCY | 5.000 | 1.414 | | -0.850 | 0.007 | * |
| | | | | ACCURACY | 87.780 | 4.940 | | | | |
| Table S7 | Figure S8B | D0 | Short | LATENCY | 3.667 | 1.312 | Spearman's rank correlation coefficient | 0.661 | 0.105 | |
| | | | | ACCURACY | 95.560 | 2.420 | | | | |
| | | D1 | Short | LATENCY | 3.333 | 1.312 | | -0.283 | 0.923 | |
| | | | | ACCURACY | 93.330 | 2.360 | | | | |
| | | D2 | Short | LATENCY | 6.222 | 1.289 | | 0.314 | 0.821 | |
| | | | | ACCURACY | 88.890 | 2.610 | | | | |
| | | D3 | Short | LATENCY | 5.667 | 1.607 | | 0.148 | 1.000 | |
| | | | | ACCURACY | 88.890 | 3.510 | | | | |
| | | D4 | Short | LATENCY | 3.444 | 1.271 | | 0.570 | 0.218 | |
| | | | | ACCURACY | 78.890 | 6.110 | | | | |
| | | D5 | Short | LATENCY | 2.667 | 0.645 | | 0.679 | 0.089 | |
| | | | | ACCURACY | 84.440 | 3.380 | | | | |
| | | D6 | Short | LATENCY | 2.778 | 0.741 | | 0.846 | 0.008 | ** |
| | | | | ACCURACY | 81.110 | 4.230 | | | | |
| | | D7 | Short | LATENCY | 2.778 | 1.392 | | 0.036 | 1.000 | |
| | | | | ACCURACY | 70.000 | 7.070 | | | | |
| | | D8 | Short | LATENCY | 3.333 | 1.067 | | 0.563 | 0.228 | |
| | | | | ACCURACY | 77.780 | 5.470 | | | | |
| | | D9 | Short | LATENCY | 2.556 | 0.801 | | 0.223 | 1.000 | |
| | | | | ACCURACY | 71.110 | 4.230 | | | | |
| | | D10 | Short | LATENCY | 2.222 | 0.683 | | 0.436 | 0.481 | |
| | | | | ACCURACY | 73.330 | 5.000 | | | | |
| | | D11 | Short | LATENCY | 4.000 | 1.190 | | 0.299 | 0.868 | |
| | | | | ACCURACY | 76.670 | 7.640 | | | | |
| | | D12 | Short | LATENCY | 2.111 | 0.772 | | 0.711 | 0.064 | |
| | | | | ACCURACY | 71.110 | 6.550 | | | | |
| | | D13 | Short | LATENCY | 2.556 | 0.899 | | 0.757 | 0.037 | * |
| | | | | ACCURACY | 73.330 | 8.160 | | | | |
| | | D14 | Short | LATENCY | 1.667 | 0.408 | | 0.745 | 0.043 | * |
| | | | | ACCURACY | 63.330 | 7.990 | | | | |
| | | D15 | Short | LATENCY | 1.333 | 0.577 | | 0.589 | 0.190 | |
| | | | | ACCURACY | 67.780 | 8.460 | | | | |
| | | D16 | Short | LATENCY | 1.444 | 0.377 | | 0.565 | 0.225 | |
| | | | | ACCURACY | 70.000 | 7.260 | | | | |

| Mouse: Correlation between LATENCY (HE peak latency) and ACCURACY (hit and correct rejection rates) in WT mice | | | | | | | | | | |
|---|---|---|---|---|---|---|---|---|---|---|
| Table | Related Figure | Day | Condition | Pair | Mean | SEM | Stastistical test | Spearman's ρ | P | * |
| Table S7 | Figure S8B | D0 | Long | LATENCY | 1.778 | 1.222 | Spearman's rank correlation coefficient | 0.650 | 0.116 | |
| | | | | ACCURACY | 6.670 | 2.360 | | | | |
| | | D1 | Long | LATENCY | 2.000 | 1.443 | | -0.042 | 1.000 | |
| | | | | ACCURACY | 13.330 | 5.270 | | | | |
| | | D2 | Long | LATENCY | 3.111 | 1.438 | | 0.122 | 1.000 | |
| | | | | ACCURACY | 24.440 | 8.520 | | | | |
| | | D3 | Long | LATENCY | 5.111 | 1.728 | | 0.205 | 1.000 | |
| | | | | ACCURACY | 24.440 | 7.090 | | | | |
| | | D4 | Long | LATENCY | 2.111 | 1.111 | | 0.405 | 0.559 | |
| | | | | ACCURACY | 56.670 | 10.270 | | | | |
| | | D5 | Long | LATENCY | 5.444 | 1.608 | | -0.154 | 1.000 | |
| | | | | ACCURACY | 54.440 | 11.320 | | | | |
| | | D6 | Long | LATENCY | 5.111 | 1.476 | | -0.399 | 0.574 | |
| | | | | ACCURACY | 62.220 | 9.090 | | | | |
| | | D7 | Long | LATENCY | 4.222 | 0.572 | | -0.125 | 1.000 | |
| | | | | ACCURACY | 71.110 | 6.330 | | | | |
| | | D8 | Long | LATENCY | 3.222 | 1.278 | | -0.396 | 0.582 | |
| | | | | ACCURACY | 81.110 | 4.840 | | | | |
| | | D9 | Long | LATENCY | 4.000 | 1.179 | | 0.022 | 1.000 | |
| | | | | ACCURACY | 78.890 | 6.330 | | | | |
| | | D10 | Long | LATENCY | 3.111 | 1.060 | | -0.502 | 0.337 | |
| | | | | ACCURACY | 83.330 | 5.770 | | | | |
| | | D11 | Long | LATENCY | 5.556 | 1.365 | | -0.618 | 0.152 | |
| | | | | ACCURACY | 84.440 | 5.560 | | | | |
| | | D12 | Long | LATENCY | 3.778 | 1.245 | | -0.614 | 0.157 | |
| | | | | ACCURACY | 86.670 | 4.710 | | | | |
| | | D13 | Long | LATENCY | 2.889 | 0.633 | | -0.130 | 1.000 | |
| | | | | ACCURACY | 86.670 | 4.710 | | | | |
| | | D14 | Long | LATENCY | 4.333 | 0.833 | | 0.204 | 1.000 | |
| | | | | ACCURACY | 84.440 | 3.770 | | | | |
| | | D15 | Long | LATENCY | 4.333 | 0.667 | | -0.351 | 0.708 | |
| | | | | ACCURACY | 87.780 | 4.010 | | | | |
| | | D16 | Long | LATENCY | 5.778 | 1.103 | | -0.069 | 1.000 | |
| | | | | ACCURACY | 92.220 | 3.240 | | | | |
| Table S7 | Figure S8B | D0 | Short | LATENCY | 4.000 | 1.616 | Spearman's rank correlation coefficient | 0.000 | 1.000 | |
| | | | | ACCURACY | 98.890 | 1.110 | | | | |
| | | D1 | Short | LATENCY | 4.333 | 1.258 | | 0.000 | 0.000 | |
| | | | | ACCURACY | 100.000 | 0.000 | | | | |
| | | D2 | Short | LATENCY | 1.889 | 1.020 | | 0.095 | 1.000 | |
| | | | | ACCURACY | 94.440 | 1.760 | | | | |
| | | D3 | Short | LATENCY | 3.556 | 1.692 | | 0.279 | 0.934 | |
| | | | | ACCURACY | 88.890 | 2.610 | | | | |
| | | D4 | Short | LATENCY | 2.333 | 1.202 | | 0.018 | 1.000 | |
| | | | | ACCURACY | 77.780 | 4.340 | | | | |
| | | D5 | Short | LATENCY | 2.667 | 0.816 | | 0.587 | 0.192 | |
| | | | | ACCURACY | 83.330 | 2.360 | | | | |
| | | D6 | Short | LATENCY | 1.778 | 0.969 | | 0.412 | 0.541 | |
| | | | | ACCURACY | 90.000 | 4.080 | | | | |
| | | D7 | Short | LATENCY | 4.111 | 1.567 | | -0.334 | 0.760 | |
| | | | | ACCURACY | 86.670 | 1.670 | | | | |
| | | D8 | Short | LATENCY | 4.444 | 1.271 | | -0.388 | 0.604 | |
| | | | | ACCURACY | 78.890 | 4.550 | | | | |
| | | D9 | Short | LATENCY | 3.889 | 0.484 | | 0.076 | 1.000 | |
| | | | | ACCURACY | 86.670 | 2.890 | | | | |
| | | D10 | Short | LATENCY | 4.000 | 1.404 | | 0.561 | 0.232 | |
| | | | | ACCURACY | 82.220 | 4.940 | | | | |
| | | D11 | Short | LATENCY | 3.889 | 1.047 | | 0.513 | 0.317 | |
| | | | | ACCURACY | 77.780 | 3.640 | | | | |
| | | D12 | Short | LATENCY | 3.222 | 0.862 | | 0.296 | 0.880 | |
| | | | | ACCURACY | 78.890 | 4.840 | | | | |
| | | D13 | Short | LATENCY | 2.222 | 0.521 | | 0.701 | 0.071 | |
| | | | | ACCURACY | 85.560 | 2.940 | | | | |
| | | D14 | Short | LATENCY | 3.778 | 1.064 | | -0.190 | 1.000 | |
| | | | | ACCURACY | 81.110 | 4.550 | | | | |
| | | D15 | Short | LATENCY | 3.333 | 0.726 | | 0.332 | 0.766 | |
| | | | | ACCURACY | 81.110 | 4.840 | | | | |
| | | D16 | Short | LATENCY | 2.000 | 0.471 | | 0.265 | 0.982 | |
| | | | | ACCURACY | 77.780 | 3.640 | | | | |
